# From Magnetosomes to Vesicles: LemA Proteins as Drivers of Bacterial Membrane Reorganization

**DOI:** 10.64898/2026.10.21.759321

**Authors:** José Manuel Borrero-de Acuña, Svea Holland, Rokas Juodeikis, Alejandro Arce-Rodríguez, Paula Ayala García, Mustafa Safa Karagöz, Joaquín Giner-Lamia, Gabriella Molinari, Gintare Deimantaviciute, Viktoria Otto, Carlos Azogue-Palma, Ian R. Brown, Katrin Müller, Martina Jahn, Kim Eileen Rennhack, Susanne Häussler, Manfred Rohde, Lothar Jänsch, Marco van Ham, Mathias Müsken, Evelyne Deery, C. Mark Smales, Meina Neumann-Schaal, Martin J. Warren, Dieter Jahn

**Affiliations:** Institute of Microbiology, Technische Universität Braunschweig, 38106 Braunschweig, Germany; Braunschweig Integrated Centre of Systems Biology (BRICS), Technische Universität Braunschweig, 38106 Braunschweig, Germany; Departamento de Microbiología, Facultad de Biología, Universidad de Sevilla, Sevilla, Spain; 4Leibniz-Institut DSMZ-Deutsche Sammlung von Mikroorganismen und Zellkulturen, 38124 Braunschweig, Germany; School of Biosciences, University of Kent, Canterbury, CT2 7NJ, UK; Quadram Institute Bioscience, Norwich Science Park, Norwich, NR4 7UQ, UK; Biochemistry and Biophysics Centre, National Heart, Lung, and Blood Institute, National Institutes of Health, Bethesda, Maryland, USA; Department of Medical Biochemistry and Biophysics, Karolinska Institutet, Tomtebodavägen 23A, 17165 Solna, Stockholm, Sweden; Instituto de Bioquímica Vegetal y Fotosíntesis, Universidad de Sevilla, Consejo Superior de Investigaciones Científicas, Américo Vespucio 49, Sevilla, 41092, Spain; Central Facility for Microscopy, Helmholtz Centre for Infection Research (HZI), 38124 Braunschweig, Germany; Institute for Molecular Bacteriology, TWINCORE GmbH, Centre of Clinical and Experimental Infection Research, a joint venture of the Hannover Medical School and the Helmholtz Centre for Infection Research, Hannover, Germany; Department of Molecular Bacteriology, Helmholtz Centre for Infection Research (HZI), 38124, Braunschweig, Germany; Department of Clinical Microbiology, Copenhagen University Hospital – Rigshospitalet, 2100 Copenhagen, Denmark; Cluster of Excellence RESIST (EXC 2155), Hannover Medical School, Hannover, 30625, Germany; Cellular Proteome Research, Helmholtz Centre for Infection Research (HZI), 38124 Braunschweig, Germany

**Keywords:** bacterial extracellular vesicles, cytoplasmic organelles, membrane invagination, *Pseudomonas aeruginosa*, bacterial cell division: PspA: LemA, MamQ, CpoB

## Abstract

The cellular machineries that remodel bacterial membranes during organelle and vesicle formation remain poorly defined. We identified MamQ as a determinant of the lipid invaginations enclosing the magnetosome. MamQ belongs to the conserved LemA protein family, whose members localize to the inner or outer membrane. Heterologous overproduction of 13 LemA proteins in *Escherichia coli* induced membrane restructuring and vesiculation, indicating conserved membrane-remodelling activity. In *Pseudomonas aeruginosa*, overproduction of the inner membrane protein LemA1 promoted cytoplasmic membrane vesicle formation, whereas the periplasmic, outer membrane-anchored LemA2 drove outer-membrane vesicle formation. Deletion of corresponding genes reduced extracellular vesicle production. Interactome analyses identified 256 partners of PaLemA1 and PaLemA2, including divisome components CpoB, FtsE, FtsI, FtsK, FtsN, FtsW, MinE, ParB, ZapG, and ZipA. Bacterial two-hybrid assays confirmed interactions between both LemA proteins and the cell division regulator CpoB, whose perturbation altered vesicle formation. Additional partners included PspA, an ESCRT-III-like protein implicated in membrane remodelling. Enriched proteins were involved in cell wall metabolism, membrane turnover, lipopolysaccharide biogenesis, protein export and maturation, energy metabolism, and stress-responsive gene regulation, suggesting that PaLemA proteins coordinate membrane-associated processes. LemA deletion mutants showed altered vesicle composition and gene expression. Furthermore, *lemA2* expression was induced during stationary phase and envelope stress via the extracytoplasmic sigma factor AlgU, itself a LemA interaction partner. Our findings support the hypothesis that the LemA family acts as a hub for assembling a conserved membrane-remodelling system that, with PspA, CpoB, and divisome components, organizes membrane-localized protein complexes required for bacterial organelle biogenesis and vesicle formation.

## Importance

Bacterial extracellular membrane vesicles and intracellular membrane-bound organelles play important roles in microbial physiology, communication, and virulence, yet the protein machineries underlying their formation remain poorly understood. Our findings reveal that members of the highly conserved LemA protein family interact closely with the cell division machinery, particularly the division regulator CpoB and the ESCRT-III-like protein PspA. These interactions support a model in which components of cell division complexes are repurposed to contribute to vesicle and organelle biogenesis. Furthermore, LemA appears to act as a central protein interaction hub linking membrane, cell wall, and lipopolysaccharide biogenesis with protein transport and gene regulation, potentially coordinating these processes within a larger multiprotein complex. Together, these findings provide new insights into bacterial membrane remodeling and may inform the engineering of bacterial vesicles for therapeutic delivery and vaccine development. Moreover, the identified proteins may represent promising candidates for future antimicrobial drug discovery.

## Introduction

The formation of intracellular membrane-bound structures and membrane vesicles in bacteria has been recognised for decades. Specialised organelles, such as chromatophores in photosynthetic bacteria, magnetosomes, and the membrane structures of methane- and ammonia-oxidizing bacteria, are central to their specialized biochemistry (1). Similarly, bacterial membrane vesicles (BMVs) play key roles in intercellular communication and microbial virulence (2, 3). Beyond their biological importance, the ability to control and engineer BMV and organelle formation holds considerable promise for biotechnology and biomedicine. Potential applications include vaccine development and immunotherapy, the targeted delivery of antimicrobial and therapeutic compounds, recombinant production platforms, cancer treatment, disease diagnostics and regenerative medicine. Moreover, engineered BMVs are being explored as platforms for delivering biomolecules, including gene-editing systems, highlighting their potential as versatile biological nanocarriers (4–7).

BMVs are commonly classified into outer membrane vesicles (OMVs), outer-inner membrane vesicles (OIMVs) and cytoplasmic membrane vesicles (CMVs). OMVs, which consist of a single membrane, are generally thought to arise from outer membrane blebbing of Gram-negative bacteria (8–10). OIMVs comprise two membranes and have been proposed to originate from coordinated blebbing or explosive cell lysis (11). In contrast, CMVs are single membrane vesicles normally produced by Gram-positive bacteria through a poorly understood mechanism (12). Several mechanisms of BMV biogenesis have been described. In Gram-negative bacteria, outer membrane vesicles (OMVs) can arise through membrane blebbing associated with envelope disturbances, including imbalances in peptidoglycan biosynthesis, the accumulation of misfolded proteins and the intercalation of hydrophobic compounds into the outer membrane. The *P. aeruginosa* signal PQS is a well-characterized example of the latter mechanism (13, 14). OIMVs can form when autolysin-mediated weakening of the peptidoglycan permits coordinated protrusion and release of the inner and outer membranes. Alternatively, phage-derived endolysins can trigger explosive cell lysis, after which membrane fragments reassemble into vesicles (15). In Gram-positive bacteria, endolysin-mediated cell-wall damage can induce ‘bubbling cell death’, while stress-induced autolysis and antibiotic-mediated inhibition of peptidoglycan synthesis can also promote cytoplasmic membrane vesicle release (16). In *Staphylococcus aureus*, phenol-soluble modulins disrupt the cytoplasmic membrane and, together with autolysin activity, contribute to vesicle release through the cell wall (17). However, all these mechanisms do not allow for a coordinated loading and defined release of vesicles as observed during various processes including infection. Moreover, while these established pathways rely largely on passive chemical blebbing, surfactant action, or destructive lytic events, the active protein machineries that orchestrate membrane remodelling during organelle and vesicle formation remain poorly defined.

To date, only limited components of bacterial membrane remodelling systems have been identified. The cyanobacterial phage shock protein PspA adopts an ESCRT-III- like fold and has been shown to promote positive membrane curvature, potentially serving as a lipid transfer platform that bridges separated membrane structures (18). In eukaryotes, the endosomal sorting complexes required for transport (ESCRTs) mediate diverse membrane remodelling processes, with the ESCRT-III subcomplex assembling into dynamic flat spiral structures that drive membrane invagination (19). Whether analogous systems operate more broadly in bacteria remains an open question.

Here, we identify the LemA family of proteins as a conserved bacterial remodelling system. We provide evidence showing that LemA proteins, in conjunction with the cell division machinery coordinated by CpoB and the ESCRT-III-like protein PspA, drive the formation of BMVs and intracellular organelles in the opportunistic pathogenic bacterium *Pseudomonas aeruginosa*.

## Results

### Initial discovery of a protein of the vesicle formation machinery: MamQ initiates inner membrane curvature during magnetosome formation

To identify proteins associated with BMV formation, we first examined magnetosome biogenesis in magnetotactic bacteria, where membrane invagination is known to precede magnetite deposition (20). We reasoned that this system could reveal factors responsible for initiating membrane curvature.

To identify such factors, we individually cloned and expressed genes previously implicated in magnetosome membrane invagination (21) (*mamB*, *I*, *L*, *Q* and *Y*). Among these, expression of *mamQ* from *Magnetospirillum gryphiswaldense* (MSR-1) in *Escherichia coli* produced a striking inner membrane phenotype, characterized by extensive invaginations and membrane ruffling, as visualised by transmission electron microscopy (TEM) of thin-sectioned cells (Fig. 1a).

**Figure 1.**
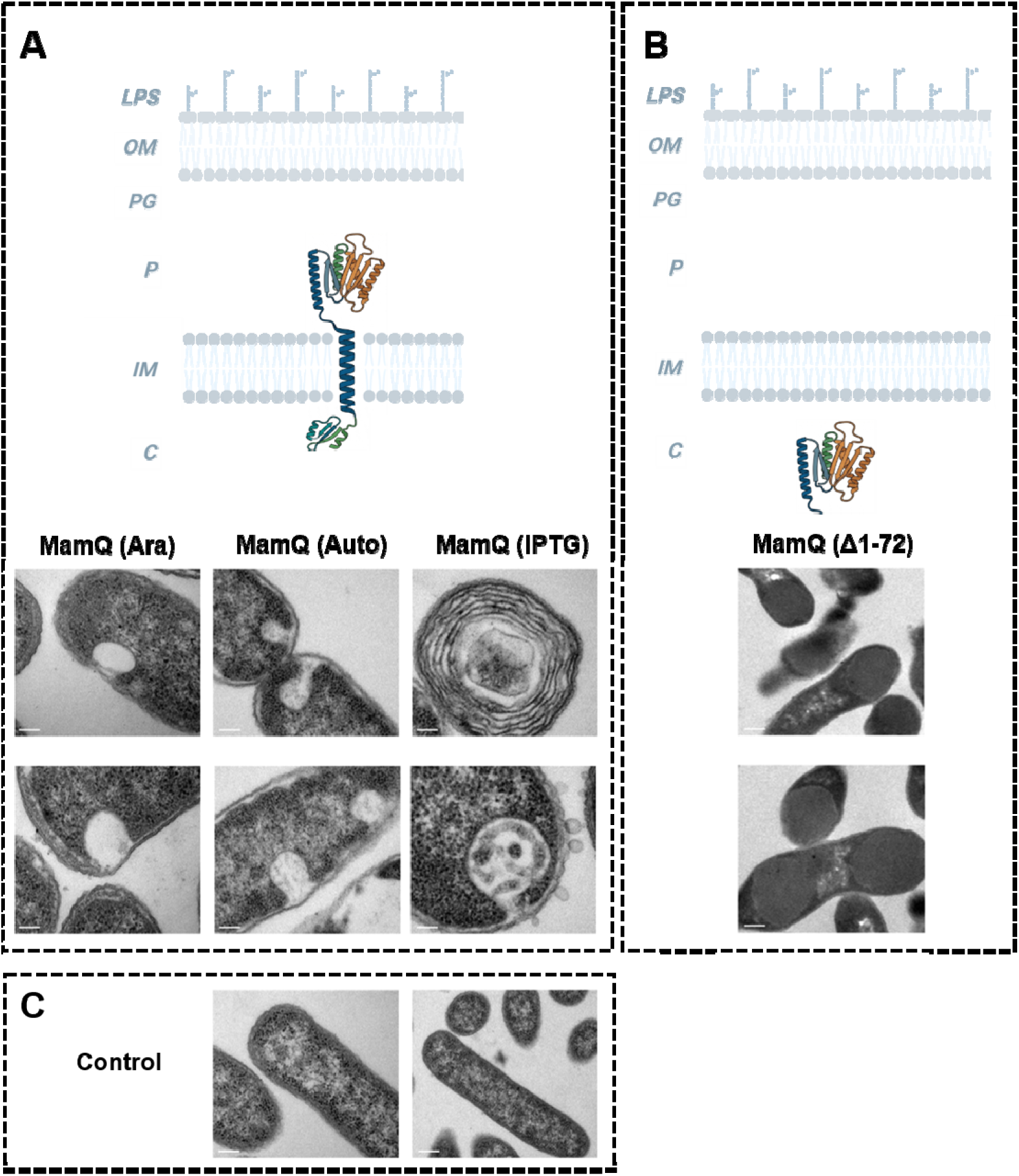
Transmission electron micrographs of recombinant *E. coli* cells producing *M. gryphiswaldense* MamQ. Electron micrographs of *E. coli* BL21 (DE3) Star cells producing (**A**) the *M. gryphiswaldense* MamQ protein using two different promoters to allow for differential gene expression and (**B**) the truncated version of MamQ. All images were taken after 16 hours of *E. coli* growth. The control strain is *E. coli* harbouring the empty pET3a plasmid (**C**). In the MamQ (Ara) panel (**A**) *E. coli* strains are shown producing MamQ under the control of an arabinose-inducible promoter, which results in a lower expression than the IPTG inducible promoter described below. Cells in the MamQ (Auto) marked panel (**A**) harbour the pET3a plasmid with an IPTG-inducible promoter grown in auto-inducible medium resulting in lower *mamQ* expression compared to induction with IPTG. MamQ (IPTG) shows the same cells fully induced with IPTG. Finally, in MamQ (Δ1-72) (**B**) the first 72 amino acid residues encoding a potential transmembrane helix were removed. The mutant protein was produced using the MamQ (IPTG) conditions. Schematic illustrations explaining the MamQ variants are shown in the upper parts of panels **A** and **B**. All scale bars are 200 nm. Abbreviations: LPS: Lipopolysaccharide; OM = Outer membrane; PG = Peptidoglycan; P = Periplasm; IM = Inner Membrane; C = Cytoplasm. The topology of MamQ: The N-terminus of MamQ faces the cytoplasm, followed by a transmembrane helix and a large periplasmic soluble domain. The PyMOL images were assembled in BioRender. The figure was created with BioRender.

Structural modelling using Alphafold (Extended Data chapter 1, Extended Data Fig. 1) predicted that MamQ comprises an N-terminal cytoplasmic domain (residues 1-46), a single transmembrane helix (residues 47 – 67) and a large C-terminal periplasmic domain (residues 68 – 272)(22, 23). Deletion of the N-terminal and the transmembrane regions resulted in protein aggregation and abolished the membrane remodelling phenotype (Fig. 1b). Consistent with these mutations in the periplasmic domain (E111A, E179A, Y181A, F242A), previously shown to impair formation, also eliminated membrane invagination when expressed in *E. coli* (22). No comparable phenotype was observed in the control strains (Fig. 1c). Together, these findings indicate that MamQ is sufficient to induce inner membrane invagination, likely through the induction of membrane curvature.

### MamQ is part of LemA family of bacterial proteins

Sequence analysis identified MamQ as a member of the LemA protein family (PF04011). To define the evolutionary relationships within this family, we constructed a maximum-likelihood phylogeny from 2,565 LemA homologues identified across 12,221 prokaryotic reference genomes in the proGenomes2 database (24) (Extended Data Tables 1-3). This analysis revealed a deep evolutionary division into two major clades, here designated LemA1 (n = 1,470) and LemA2 (n = 1,095) (Extended Data chapter 2, Extended Data Fig. 2a). Each clade further diversified into distinct sublineages, which we classify as LemA1.1, LemA1.2, LemA2.1 and LemA2.2.

LemA family members were detected across 36 bacterial phyla predominantly within Proteobacteria (46%), Firmicutes (20%), Bacteroidota (16%) and Actinobacteriota (9%) (Extended Data Fig. 2b). The ecological distribution of LemA homologs reveals an important deviation from the baseline habitat representation in the proGenomes2 database (24). While the reference dataset is heavily biased toward host-associated species (83%), LemA sequences are significantly enriched in environmental lineages (Fisher’s exact test, FDR-corrected). Specifically, although soil and aquatic habitats account for only ∼8% of the species in proGenomes2, they represent 20% of LemA occurrences in soil and 33% in aquatic environments (Extended Data Fig. 2c). Furthermore, the evolutionary retention of both homologs within a single genome is notably prevalent in solid environmental substrates, specifically soil (45%) and aquatic sediments (23%) (Extended Data Fig. 2c). In contrast, the presence of both LemA homologues is significantly lower in host- and disease-associated strains compared to the database baseline. These findings suggest that while LemA is maintained across diverse lifestyles, the conservation and duplication of these proteins likely confer a critical selective advantage for survival in fluctuating habitat such as soil and aquatic ecosystems.

### LemA proteins from diverse bacteria restructure *E. coli* membranes

Having identified MamQ as a LemA family member capable of inducing membrane invagination, and established the broad phylogenetic conservation of LemA proteins, we next asked whether this membrane-remodelling activity is a general feature of the family. To address this, we cloned and expressed thirteen different *lemA* homologues from phylogenetically diverse bacteria in *E. coli* (Extended Data Fig. 3 & 4). The selected proteins (Extended Data chapter 3 - Extended Data Fig. 3a) varied predicted membrane topology and originated from both Gram-negative (*Bacteroides*, *Brucella*, *Crinalium*, *Pseudomonas*, *Spirochaeta*, *Thioflavicoccus*) and Gram-positive genera (*Anaerobacillus*, *Bacillus*, *Exiguobacterium*, *Listeria*) (Extended Data Fig. 3b). Details of recombinant production, solubility, and impact on growth and cell integrity are given in Extended Data Figs. 3c, 4, 5). Figure 2 depicts the morphotypes of strains exhibiting robust LemA production, including some variants with slight deviations in electrophoretic mobility, while retaining normal growth characteristics and cellular integrity. In contrast, the strains presented in Extended Data Fig. 4 displayed aberrant electrophoretic migration patterns, negligible or undetectable protein production, and/or evidence of host toxicity. Although intracellular membrane proliferation was detected in all samples, the morphotypes shown in Extended Data Fig. 4 exhibited more pronounced intracellular protein aggregation together with signs of cellular disintegration. TEM of thin sectioned cells expressing LemA proteins revealed extensive intracellular membrane proliferation in all cases examined (Fig. 2 & Extended Data Fig. 4 & 6), consistent with the phenotype observed with MamQ. However, distinct morphological outcomes were observed depending on the LemA homologue expressed. These included (i) formation of CMVs (AaLemA1, BcLemA1, LmLemA1, BatLemA1B, BmLemA2), (ii) intracellular tubular membrane structures (ExcLemA1, ThmLemA2, CreLemA2), (iii) extrusion of the inner membrane into the periplasmic space (BrmLemA2) and (iv) cell lysis (BatLemA1A, StLemA2). Together, these results demonstrate that membrane remodelling is a conserved functional property of LemA proteins, while also revealing substantial diversity in the membrane architectures they generate.

**Figure 2.**
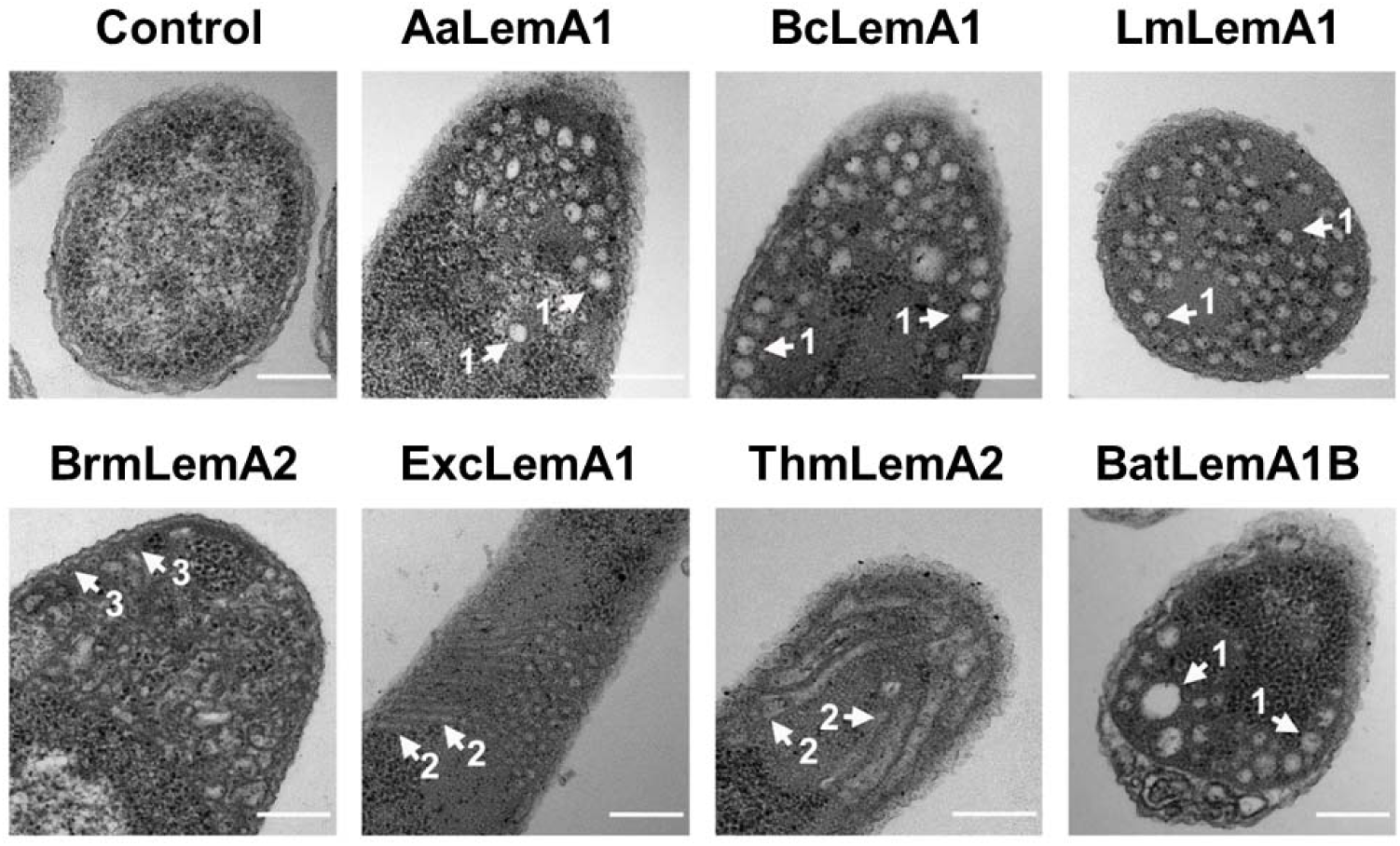
Transmission electron micrographs of recombinant *E. coli* cells producing 7 different bacterial LemA proteins reveal membrane restructuring. Recombinant *E. coli* BL21 (DE3) Star cells producing *Anaerobacillus alkalilacustris* LemA1 (AaLemA1), *Bacillus clausii* LemA1 (BcLemA1), *Listeria monocytogenes* (LmLemA1) *Brucella melitensis* LemA2 (BrmLemA2), *Exiguobacterium chinqhucha* LemA1 (ExcLemA1), *Thioflavicoccus mobilis* LemA2 (ThmLemA2) and *Bacteroides thetaiotaomicron* LemA1B (BatLemA1B) were analysed by TEM and photographed after 16 hours growth. LemA1 protein production leads predominantly to the generation of CMVs; LemA2 production leads to generation of a multitude of membranous structures; ExcLemA1 production generates tubular membranous structures. In the different sections the most characteristic membrane structures observed are indicated as 1 = CMVs, 2 = intracellular tubular membrane structures and 3 = extrusion of the inner membrane into the periplasmic space. The scale bars shown represent 200 nm.

### *P. aeruginosa* LemA1 and LemA2 differentially modulate membrane architecture and vesicle formation

Having established that LemA proteins broadly induce membrane remodelling, we next focused on two representative homologues from *Pseudomonas aeruginosa,* PaLemA1 and PaLemA2, to investigate their physiological roles. The *P. aeruginosa* PA14 genome encodes both proteins in distinct operons, suggesting potentially specialized functions. We first examined their effects following heterologous expression in *E. coli*. TEM and field emission scanning electron microscopy (FESEM) analysis (Extended Data chapter 4 & Extended Data Figs. 5 - 7) revealed distinct membrane phenotypes for each protein. Expression of PaLemA1 led predominantly to the formation of circular intracellular vesicles, in contrast to the lamellar membrane structures observed for several of the LemA1 homologues (Extended Data Figs. 6 & 7a). In comparison, PaLemA2 expression resulted in stacked membrane structures proximal to the inner membrane (Extended Data Figs. 6 & 7c) and the appearance of large vesicular protrusions at the cell surface (Extended Data Fig. 6), indicative of enhanced vesiculation. Prolonged expression of PaLemA1 led to intracellular protein aggregation and eventual cell lysis (Extended Data Fig. 5). Immunogold labelling localised PaLemA1 primarily to the inner membrane and to the inner membrane associated intracellular structures, whereas PaLemA2 was detected at both the inner and outer membranes (Extended Data Fig. 8), consistent with their distinct phenotypic outputs.

To assess the roles of these proteins in their native context, we generated single (ΔPa*lemA*1, ΔPa*lemA*2) and double (ΔPa*lemA*1+*A*2) deletion mutants in *P. aeruginosa*, along with corresponding complementation and overexpression strains.

Protein production was confirmed by immunoblotting (Extended Data Fig. 9). Electron microscopy analyses revealed no pronounced ultrastructural phenotypes when comparing wildtype and deletions strains, both harbouring the empty vector control (Extended Data Fig. 10 & 11). In contrast, overproduction of PaLemA1 induced intracellular tubular and vesicular membrane structures (Fig. 3), whereas PaLemA2 overproduction primarily enhanced outer membrane vesiculation and promoted cell aggregation, as visualized by FESEM (Fig. 3, Extended Data Figs. 7 & 10). A snapshot of the vesiculation process is shown in more detail in Extended Data Fig. 12.

**Figure 3:**
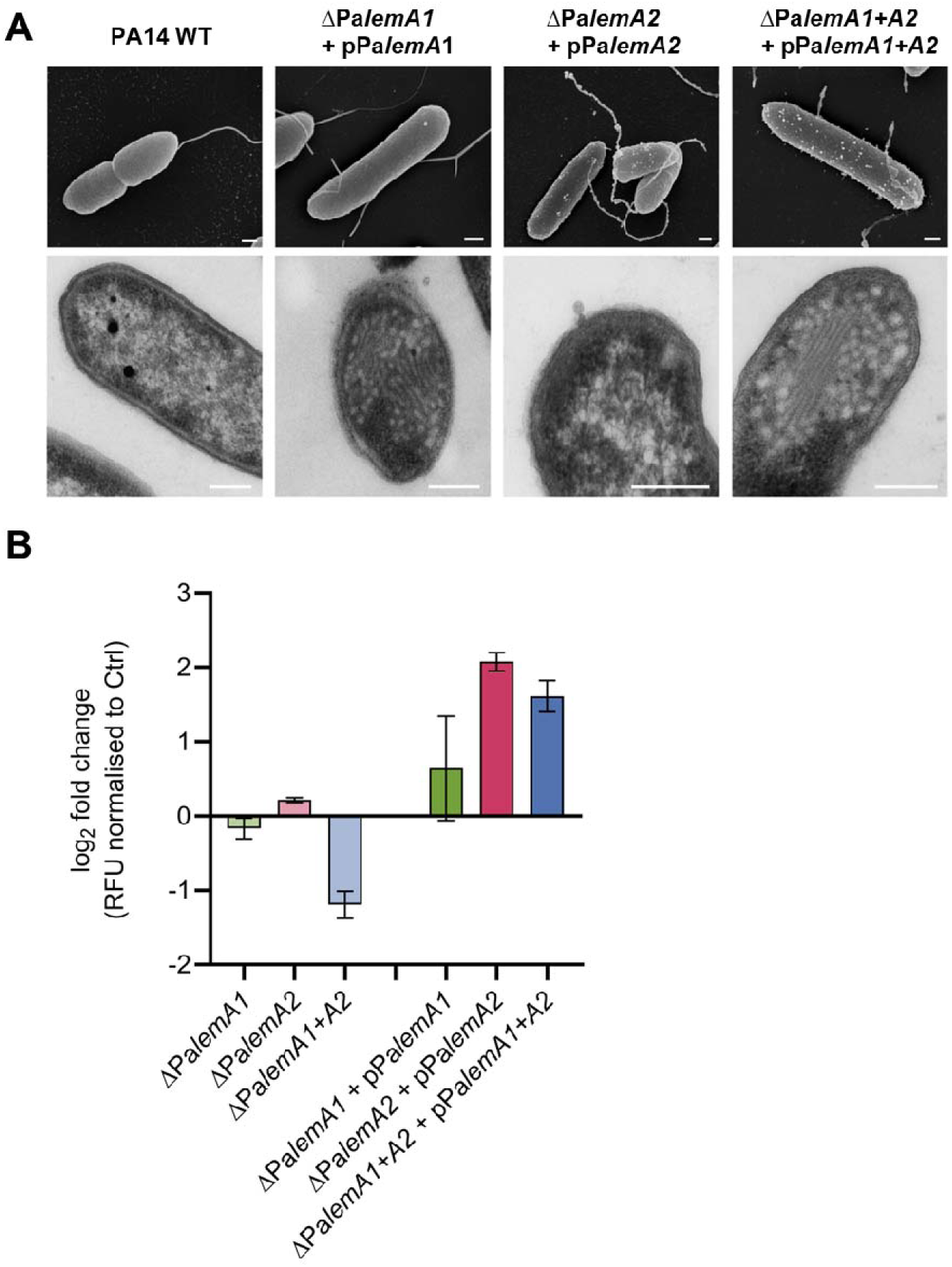
Impact of PaLemA proteins on BMV formation in *P. aeruginosa*. Both, Pa*lemA1* and Pa*lemA2* genes were individually (pPa*lemA1,* pPa*lemA2*) and collectively (pPa*lemA1+A2*) overexpressed in their corresponding *P. aeruginosa* PA14 knockout strains (_Δ_Pa*lemA1*, _Δ_Pa*lemA2*, _Δ_Pa*lemA1+A2*). **(A)** Field emission scanning electron microscopy (FESEM) and transmission electron microscopy (TEM) were used to visualise the effects of recombinant production of membrane-restructuring PaLemA proteins. Overproduction of PaLemA1 induced the formation of multiple intracellular globular structures as visualised by TEM (lower row), while FESEM (upper row) revealed membrane blebbing and BMV formation on the bacterial surface upon PaLemA2 overproduction. Strains overproducing both proteins displayed a combined phenotype. **(B)** BMV production of Pa*lemA* single and double knockout mutants, as well as BMVs of strains overexpressing the corresponding genes in the mutant background, were isolated. BMV quantification was performed using the fluorescent membrane-binding dye FM4-64 to determine lipid content by relative fluorescence (RFU). BMVs derived from knockout mutants were isolated in early stationary phase, while BMVs derived from strains overexpressing the genes of interest were isolated 2 hours after induction of gene expression. Data was normalized to the mean of the corresponding control samples: PA14 WT for knockout strains and the empty vector control (p*ctrl634*) for overexpressing strains. Data are represented as the log2 fold-change and show the mean ± SEM of three biological replicates. The graphs were assembled in GraphPad.

We next quantified BMV production using the lipophilic dye FM4-64 by measuring corresponding relative fluorescence units (RFU) (Fig. 3). To rule out non-specific cell lysis or cellular fragmentation, we confirmed via CFU counting and FESEM that cell integrity remained unaffected without cellular debris accumulation under our short-term induction conditions. This fluorescence-based approach builds on a robust, well- established method from our group ensuring reliable vesicle quantification independent of protein contaminants (25, 26). Thus, our controlled parameters confirm that the resulting signal reflects active, targeted membrane vesiculation rather than lytic breakdown. Deletion of both Pa*lemA* genes led to a marked reduction in vesicle production relative to wildtype, whereas single deletions had only minor effects, indicating partial functional redundancy. In contrast, overexpression of either protein increased vesicle yield, with PaLemA2 producing the strongest effect with 4.2 times more BMV compared to the wildtype control. Co-expression of both proteins also enhanced vesicle production (Fig. 3).

Together, these results suggest that PaLemA1 and PaLemA2 differentially remodel bacterial membranes and cooperatively promote vesicle biogenesis, linking LemA function to both intracellular membrane organization and extracellular vesicle production.

### Interactome analysis reveals a LemA-associated membrane-localised remodelling machinery

Having found evidence that PaLemA1 and PaLemA2 remodel membranes and promote vesicle biogenesis, we next sought to define the underlying protein machinery by identifying their interaction partners. To capture both stable and transient interactions, we performed *in vivo* formaldehyde-cross-linking in *P. aeruginosa* expressing Pa*lemA1* and Pa*lemA2* genes, followed by affinity-purification of Strep-tagged PaLemA1 and PaLemA2 complexes. Co-purifying proteins were identified by high-resolution tandem mass spectrometry. This approach enabled the detection of both direct interactors and proteins associated within crosslinked multiprotein complexes. Given the membrane localization of LemA proteins, the recovered complexes are expected to include components from both the cytoplasmic and periplasmic compartments.

Across three independent experiments, we identified 119 candidate proteins (from 1403 detected proteins) that were significantly enriched with PaLemA1, and 133 candidate proteins (from 2259 detected proteins) enriched with PaLemA2 (Extended Data Tables 4 & 5). Thirteen proteins were common to both datasets, indicating a shared interaction core (Extended Data Tables 4 & 5). Quantification and normalization were performed against the PA14 wildtype proteome. Down-stream analyses revealed a clear-cut separation of PaLemA1- and PaLemA2-associated protein clusters (Extended Data Fig. 13, in red), supporting the existence of both shared and distinct interaction networks for the two LemA proteins.

### The PaLemA interactome links vesicle biogenesis to the cell division machinery

Functional annotation of the PaLemA1- and PaLemA2-associated interactome revealed a striking enrichment of proteins involved in cell division (27) (Extended Data chapter 7, Extended Data Table 6). Although initially unexpected, this observation is consistent with the shared requirement for coordinated membrane remodelling during both cytokinesis and BMV formation.

Key components of the divisome were identified among the PaLemA interaction partners. Early steps in cell division involve the assembly of the FtsZ ring and its tethering to the membrane by FtsA and ZipA, the latter detected in association with PaLemA1. Spatial regulation of divisome positioning is mediated via the Min system, with MinE (PaLemA1 associated) playing a central role in rod-shaped bacteria. Additional factors linked to chromosome segregation and divisome coordination, including ParAB, were also enriched in the PaLemA1 interactome (28).

Subsequent stages of divisome assembly were similarly represented. The recruitment of the ABC-transporter-like FtsEX (FtsE bound to PaLemA1) and the translocase FtsK (bound to PaLemA1) was observed, components that interact with the preformed FtsZ-FtsA-ZipA complex (29). Late-stage division proteins are recruited and stabilized by FtsQLB at the midcell division site, including the recruitment of the septum-specific transpeptidase FtsI (bound to PaLemA2) and its partner the glycosyltransferase FtsW (30). Stabilisation of the Z-ring through Zap proteins, such as ZapG (bound to PaLemA1) (31), further supports a link between LemA proteins and divisome dynamics. In addition, proteins contributing to outer membrane integrity and constriction, including TolA of the Tol-Pal complex, were detected (32).

Beyond canonical divisome components, proteins involved in surface appendage assembly and cell envelope organization, such as FliF, FimV and PilP, were also enriched, suggesting broader integration of LemA proteins into envelope-associated processes.

Together, these findings indicate that LemA proteins interface extensively with the cell division machinery and associated envelope systems. We propose that PaLemA proteins co-opt or reorganize elements of the divisome to drive membrane curvature and vesicle formation. A working model illustrating the integration of the LemA proteins into cell division-associated membrane remodelling is presented in Fig. 4.

**Figure 4:**
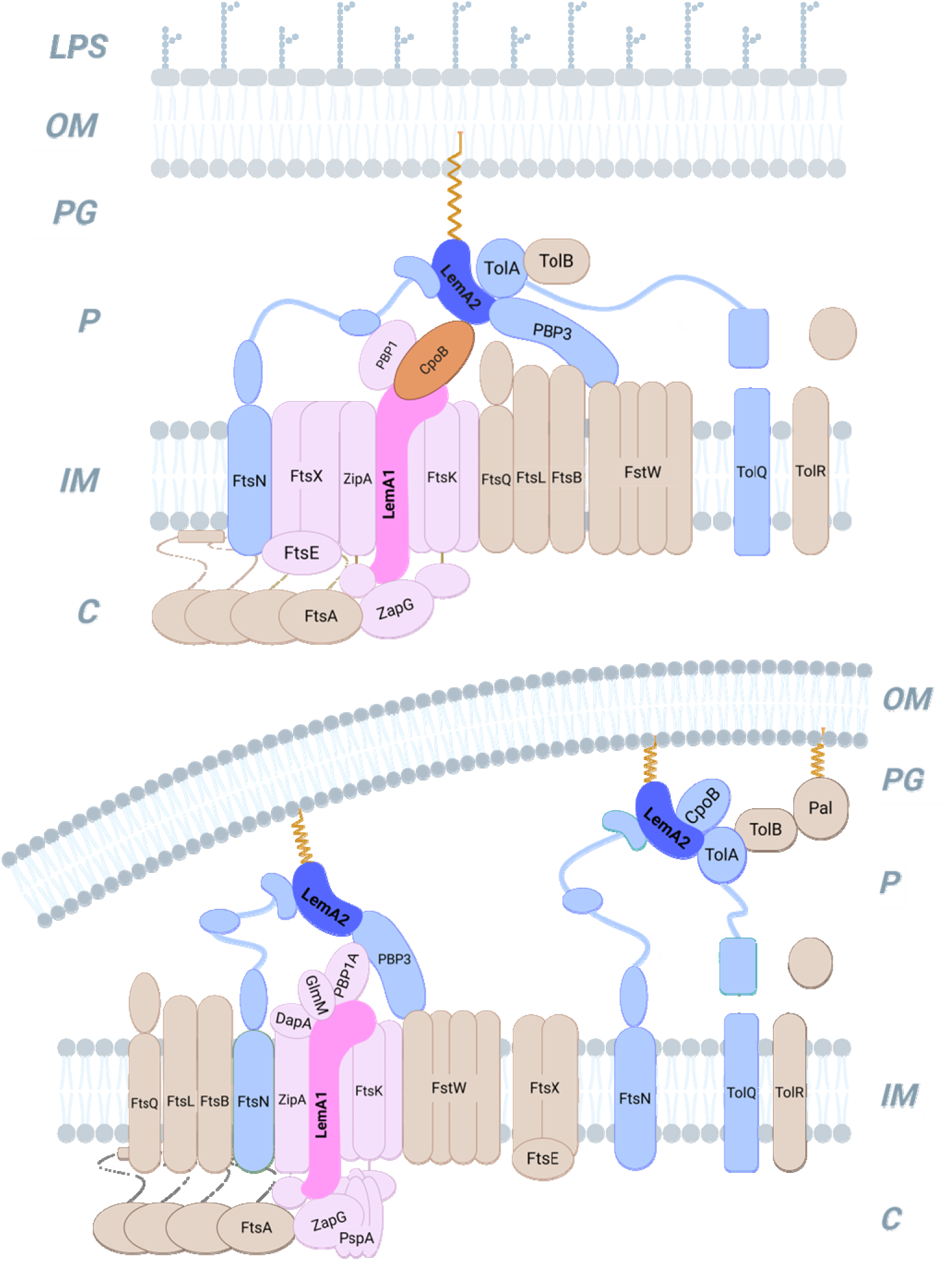
Protein complex model showing the major components of the cell division systems found attached to the PaLemAs. PaLemA1 and PaLemA2 interactome data for the interaction of various proteins of the know cell division system were visualized as a supercomplex. PaLemA1 is coloured in magenta while PaLemA2 is in blue. Displayed protein-protein interactions do not have to be necessarily of direct nature due to the crosslinking procedure. They rather represent the observed overall supercomplex. Shown are two stages of BMV formation. The upper panel shows the formation of the complex, while the lower panel shows the potential complex at work. Abbreviations: LPS = Lipopolysaccharide; OM = Outer membrane; P = Periplasm; PG = Peptidoglycan; IM = Inner Membrane; C = Cytoplasm. The figure was created with BioRender.

### The PaLemA interactome links vesicle formation to cell envelope biogenesis and lipid metabolism

In addition to divisome components, the PaLemA1 and PaLemA2 interactomes were strongly enriched involved in cell wall restructuring, membrane metabolism and lipopolysaccharide (LPS) assembly (Extended data chapter 8, Extended Data Table 6).

Cell division, and by extension vesicle formation, requires tightly coordinated peptidoglycan synthesis, remodelling and turnover. Central to these processes are penicillin-binding proteins (PBPs), which catalyse glycan strand polymerization and peptide cross-linking. We identified class A PBPs associated with PaLemA proteins, including PBP1A (PonA; interacting with both PaLemA1 and PaLemA2) and PBP1B (MrcB; enriched with PaLemA1), key enzymes that are essential for cell growth and division (33, 34). These PBPs are known to operate within dynamic complexes linked to the divisome, consistent with our observation that LemA proteins also associate with core division factors such as ZipA, FtsE and FtsN. In addition, the D,D-carboxypeptidase PBP5 (DacC), which localizes to sites of active peptidoglycan synthesis (35), was enriched with PaLemA2, further supporting a role for LemA proteins at sites of active cell wall remodelling.

Beyond PBPs, we detected enzymes involved in precursor biosynthesis and recycling of the cell wall, including GalU and GlmM, as well as the recycling enzyme NagZ. Components of the diaminopimelate (DAP) pathway (DapA, DapB and LysA), required for lysine and peptidoglycan synthesis, were also present, indicating that LemA-associated complexes are linked to both the synthesis and turnover of cell wall material (Extended Data Tables 4-6, detailed description of function in Extended Data chapters 7-8).

Consistent with a broader role in membrane dynamics, the PaLemA interactomes further included key enzymes of lipid biosynthesis and degradation. These encompassed glycerol-3-phosphate acyltransferase PlsB, acetyl-coenzyme A carboxylase carboxyl transferase (AccA) and enoyl-acyl-carrier protein reductase FabV, which contribute to phospholipid synthesis, as well as enzymes of β-oxidation including FadE (Acyl-coenzyme A dehydrogenase), FadB and the α-subunit of the multienzyme complex FadBA. The phospholipid-modifying enzyme PssA (CDP-diacylglycerol-serine O-phosphatidyltransferase) was also detected, suggesting active membrane remodelling at LemA-associated sites. In agreement with these interactions, PaLemA-dependent changes in lipid composition were observed, characterized by a decrease in long-chain and unsaturated lipids and a relative increase in mid-chain and hydroxylated lipid species (Extended Data Fig. 15, Extended Data chapter 9 with references).

Proteins involved in lipid transport and outer membrane assembly were likewise enriched. These included AsmA family proteins and components of the Ttg2ECBA (MlaBDEF) transport system, which mediate glycerophospholipids transport across the periplasm. In addition, multiple factors involved in LPS biosynthesis and transport were identified, including the heptosyltransferases WaaF and WaaC, the lipid A-modifying dioxygenase LpxO2, and the periplasmic subunit LptA. The lipoprotein chaperone LoIA, essential for outer membrane lipoprotein localization, was also associated with PaLemA2 (Extended Data chapter 11 with references). Notably, many of these components overlap with proteins previously identified in the purified divisome complex from *Salmonella enterica* serovar Typhimurium, including FtsE, FtsK, FtsN, CpoB, PBP3, PBP1A, PBP1B, TolA, RodZ as well as Zap and Min proteins (36), supporting the concept that LemA proteins engage a conserved membrane remodelling network. Collectively, these findings demonstrate that LemA proteins are embedded within a broad cell envelope biogenesis machinery, coordinating peptidoglycan remodelling, lipid metabolism and outer membrane assembly to drive membrane curvature and vesicle formation.

### The PaLemA1 interactome points towards suggests the ESCRT-III-like protein PaPspA as a modulator of vesicle formation

Among the PaLemA1 interactome partners, we identified PaPspA, an ESCRT-III-like protein previously implicated in bacterial membrane remodelling and vesicle formation (37, 38). This finding further supports a functional link between LemA proteins and conserved membrane-shaping systems.

To assess the role of PaPspA in LemA-dependent vesiculation, we analysed extracellular vesicle production in a *P. aeruginosa pspA* deletion strain. Gene deletion resulted in a modest reduction in vesicle production relative to the wild type, without severely compromising vesicle biogenesis (Extended Data Fig. 16), indicating that PaPspA contributes to, but is not essential for, vesicle formation under the conditions tested. However, overexpression of PaPspA in the mutant background resulted in a marked diminishment in extracellular BMV formation (Extended Data Fig. 16, Extended Data chapter 12 with references). A similar decrease was observed upon overexpression of either of the Pa*lemA* genes in the *P. aeruginosa* ΔPa*pspA* strain, with a more pronounced reduction observed when both genes were co-expressed (Extended Data Fig. 16).

Although speculative, these findings may suggest that vesicle formation depends on a balanced stoichiometry between LemA proteins and PaPspA, and that disruption of this balance impairs the function of the vesicle-forming machinery. Together, these results identify PaPspA as a non-essential but functionally important component of the LemA-associated vesiculation machinery, highlighting the plasticity of this system and its capacity to maintain membrane integrity under conditions of perturbation.

### The PaLemA interactome identifies PaCpoB as a key regulator of vesicle formation

Among the proteins shared between the PaLemA1 and PaLemA2 interactomes, the cell division protein CpoB (**C**oordinator of **P**eptidoglycan synthesis and **O**uter Membrane constriction, associated with PBP1**B**) emerged as a prominent candidate (Extended Data table 6). CpoB is known to associate with components of the Tol-Pal system and peptidoglycan synthesis machinery, linking cell wall remodelling to outer membrane dynamics (39).

To validate the interaction between LemA proteins and CpoB, we employed a bacterial two-hybrid system. Truncated variants of PaLemA1 (periplasmic domain) and PaLemA2 (lacking the signal peptide) as well as PaCpoB (lacking its leader sequence), were fused to complementary fragments of adenylate kinase. All tested combinations consistently demonstrated strong interactions between PaCpoB and both LemA proteins (Fig. 5a), confirming a direct association observed in the interactome dataset.

**Figure 5:**
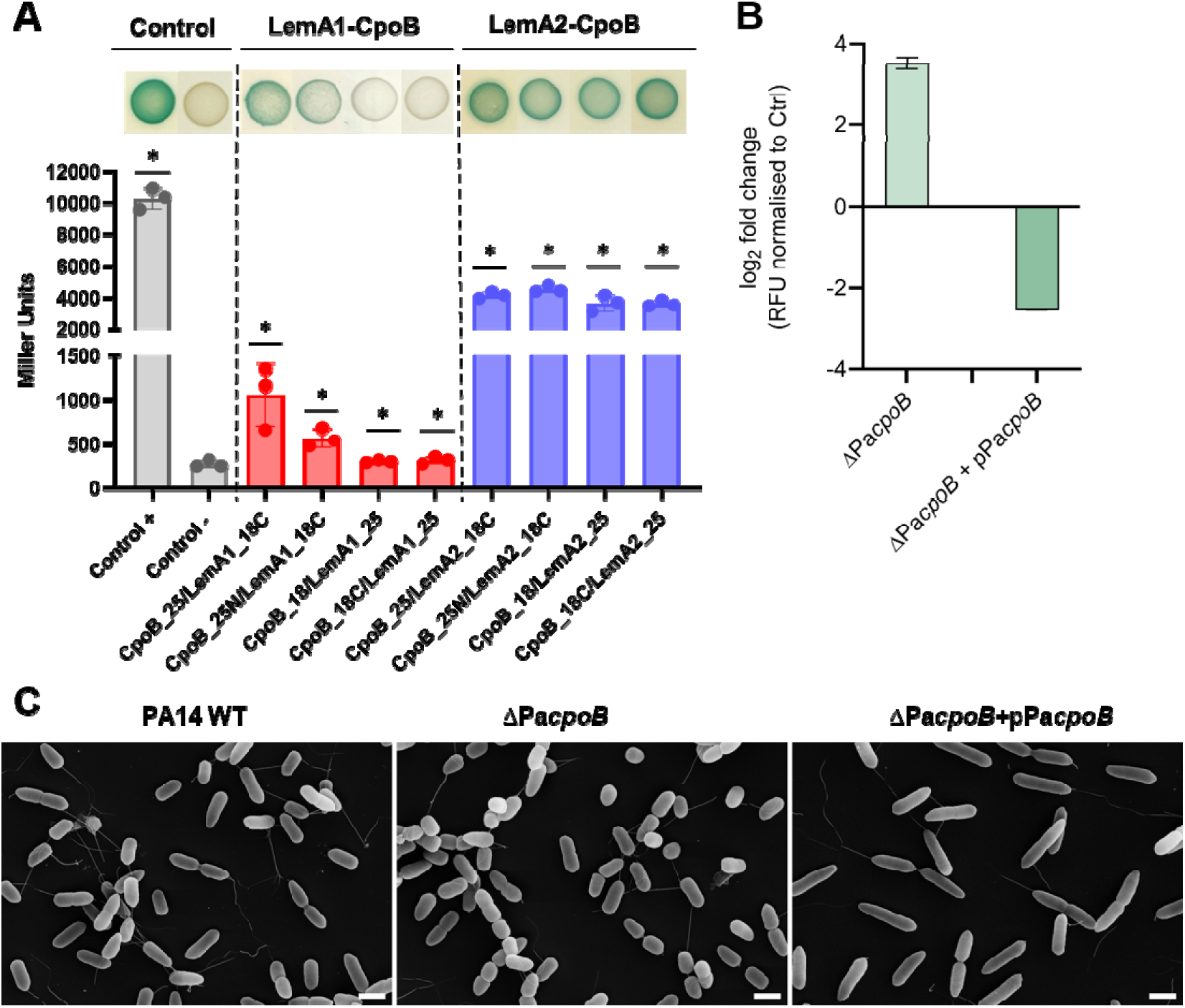
Function of the PaCpoB protein of cell division in BMV formation in *P. aeruginosa*. **(A)** Bacterial two-hybrid results for CpoB-LemA interactions. Levels of _β_-galactosidase activity of *E. coli* BTH101 cells co-transformed with the indicated T18 and T25 fusion constructs. The positive control (+) corresponds to leucine zipper fusions, whereas the negative control (-) corresponds to empty T18 and T25 vectors. The _β_-galactosidase activities were individually compared to the empty vector using a Mann-Whitney nonparametric test. Values tagged with asterisks (*) are significantly different at the level *p*-value = 0.05. At the top of the graph, representative colonies resulting from each co-transformation are shown. LacZ activity (blue) indicates protein-protein interaction. The graphs were assembled in GraphPad. **(B)** BMVs derived by the Pa*cpoB* knockout mutant (_Δ_Pa*cpoB*) and the Pa*cpoB*-deficient strain overexpressing Pa*cpoB* (_Δ_Pa*cpoB* +pPa*cpoB*) were isolated in early stationary phase or 2 hpi with 100 µM IPTG, respectively. BMV quantification was performed using the fluorescent membrane-binding dye FM4-64 to determine lipid content by relative fluorescence (RFU). All BMV samples were normalized to the mean RFU of the respective PA14 wildtype (WT) or the p*634ctrl* empty vector control. Data are represented as the log_2_ fold-change and show the mean ± SEM of three biological replicates. **(C)** FESEM analysis revealed that Pa*cpoB* deficiency compromised cell elongation and division, resulting in rounded nearly coccoid cells. This phenotype was largely rescued by complementation, which restored rod-shaped morphology, although the cells exhibited a sticky appearance and mild defects in cell fission. Scale bars are 1 µm.

We next assessed the functional role of CpoB in vesicle formation in *P. aeruginosa.* A CRISPR-Cas deletion of *cpoB* (ΔPa*cpoB*) resulted in a striking ∼11.5-fold increase in BMV production relative to wildtype, whereas complementation of the mutant (ΔPa*cpoB* + *cpoB*) markedly reduced vesicle formation (∼5.8-fold decrease compared to control) (Fig. 5b). In parallel, electron microscopy revealed pronounced morphological changes: the ΔPa*cpoB* mutant exhibited rounded, partially coccoid cells, while complementation restored the characteristic rod-shaped morphology (Fig. 5c).

These findings identify PaCpoB as a key negative regulator of vesicle formation and demonstrate that its interaction with PaLemA proteins is functionally relevant for maintaining membrane architecture and cell shape. Consistent with this, PaCpoB has been implicated in coordinating peptidoglycan synthesis with outer membrane constriction, suggesting that LemA proteins may modulate this process to promote membrane curvature and vesicle release.

Notably, bacterial two-hybrid analysis of PaLemA proteins with additional divisome-associated proteins, including FstN homologue and the bifunctional peptidoglycan synthase MrcB, showed weak or no direct interaction (Extended Data Fig.17). This suggests that these proteins are likely associated indirectly within larger LemA-associated complexes, consistent with the crosslinking-based interactomics approach.

### The PaLemA links vesicle biogenesis to protein export and cargo loading

In addition to cell envelope and division components, the PaLemA1 and PaLemA2 interactomes were strongly enriched in proteins involved in Sec- and TAT-dependent protein export, processes essential for the delivery of enzymes and structural factors to the periplasm outer membrane (40). We identified nearly the complete SEC translocon machinery among PaLemA-associated proteins, including SecA, SecB, SecD, SecF, SecY, PpiD and YajC, as well as accessory factors such as the signal peptidase LepB (41), the thiol disulfide oxidoreductases DsbC and DsbG and the periplasmic chaperone SurA (Extended Data Table 6 & 7, model in Extended Fig. 18). In addition, the TAT translocase component TatA was detected in the PaLemA1 interactome. Several known interactions within this network, including connections between the SecYEG translocon and FtsH and HflC, were also observed, supporting the integrity of the dataset (42).

These findings suggest that the LemA-associated complexes are closely linked to protein export pathways, raising the possibility that vesicle formation is coordinated with delivery and incorporation of specific cargo proteins. To test this, we analysed the protein composition of BMVs isolated from the wildtype and LemA deficient strains of *P. aeruginosa*. Proteomic analysis revealed 446 proteins in wildtype BMVs, whereas vesicles from ΔPa*lemA*1, ΔPa*lemA*2 and ΔPa*lemA*1+A2 mutants contained substantially fewer proteins (245, 234 and 247 proteins, respectively) (Extended Data Table 8). Notably, several abundant BMV-associated proteins were selectively depleted in the LemA mutant strains, indicating altered cargo composition (Extended Data chapter 14 with references). Comparison of the PaLemA interactomes with the BMV proteome further revealed a significant overlap, with multiple LemA-associated proteins also detected in wildtype vesicles. In contrast, these proteins were largely absent from vesicles derived from LemA-deficient strains (Extended Data Table 8), demonstrating that LemA proteins influence not only vesicle formation but also cargo selection. These results indicate that LemA proteins couple membrane remodelling to protein export pathways, thereby coordinating biogenesis with selective incorporation of protein cargo.

### The PaLemA interactome links vesicle formation to nutrient transport and metabolic adaptation

In addition to envelope biogenesis and protein export, the PaLemA1 and PaLemA2 interactomes were enriched in proteins involved in nutrient uptake, chemotaxis and amino acid metabolism (Extended Data Table 9 & Extended Data chapter 15 with references), suggesting that vesicle formation is coordinated with metabolic adaptation. A prominent feature of the dataset was the enrichment of transport systems dedicated to the uptake of reduced nitrogen sources. These included the di- and tripeptide transport system, with multiple substrate-binding proteins (SBP) (DppA1, DppA3-A5) and the ATP-binding component DppF, as well as polyamine transport proteins such as the spermidine SBP SpuD and SpuE and the ATP-binding proteins PotA and SpuF. Consistent with this, several amino acid transporters were identified, including systems for arginine (AotJ/AotP), branched chain amino acids (BraC) and methionine and related substrates (MetQ/YbeJ).

Strikingly, corresponding chemotaxis receptors (PctA, PctB, PctC), which sense amino acids and direct bacterial movement towards nutrient sources, were also enriched (43). This coordinated presence of transporters and chemosensory proteins suggests that LemA-associated complexes may spatially integrate nutrient sensing and uptake with membrane remodelling processes.

In addition, enzymes involved in amino acid biosynthesis and catabolism, including CysE, LeuB, PheA, PhhB/C, and GlyA2, among others, were detected, indicating that LemA-associated sites are linked to active metabolic flux. In contrast, transport systems for other substrates, such as sugars (GltF, GltJ) or inorganic ions (for example, CysP, Hit, FbpA, MgtE and PstS) were comparatively underrepresented, pointing to a preferential association with nitrogen-related metabolism.

Finally, proteins involved in membrane transport and stress adaptation, including the RND multidrug efflux system MexAB and mechanosensitive channels (MscK and YbdG), were also present (summarized in the model in Extended Data Fig.18), suggesting a role for LemA proteins in coordinating vesicle formation with envelope stress responses. These findings indicate that LemA-associated vesicle formation is functionally linked to nutrient acquisition and metabolic state, further supporting the concept that LemA proteins integrate membrane remodelling with core cellular physiology.

### The PaLemA interactome links vesicle formation to cellular energy metabolism

Membrane remodelling and vesicle biogenesis are energy-intensive processes. Consistent with this, the PaLemA1 and PaLemA2 interactomes were enriched in components of the respiratory chain and central energy metabolism (Extended Data Table 10), indicating that LemA-associated complexes are coupled to cellular energy generation (44). Multiple enzymes involved in electron transfer and redox metabolism were identified, including subunits of the NADH dehydrogenase (NuoB, NuoC, NuoE and NuoI), malate:quinone oxidoreductase (MqoB), glucose dehydrogenase (Gcd), and formate dehydrogenase-O (subunit FdnG), which were associated with PaLemA1. Additional components of the respiratory chain, such as two succinate dehydrogenase subunits (SdhC and SdhD), the periplasmic electron transferring protein azurin (Azu) and the iron-sulphur protein FixG, were also detected in the PaLemA interactome (Extended Data chapter 16 & Extended Data Table 10).

In *P. aeruginosa*, phenazines mediate extracellular electron transfer, and enzymes involved in their biosynthesis (PhzD1/D2) were found associated with PaLemA1(45). Furthermore, key elements of terminal electron transport were present, including subunits of the *cbb_3_*-type cytochrome c oxidase (CcoN and CcoP) and the cytochrome c-type biogenesis protein CycH. Consistent with these findings, enzymes and regulatory factors involved in the control and biosynthesis of essential redox cofactors were also enriched, including HemX (regulatory protein that helps control the levels of enzymes in the heme biosynthetic pathway), UbiE (ubiquinone biosynthesis) and ThiG (thiamine biosynthesis). Together these components suggest that LemA-associated complexes are positioned with metabolically active regions enriched in electron transport activity (summarized in the model in Extended Data Fig. 19).

### The PaLemA interactome links vesicle formation to gene regulation and stress response

In addition to membrane remodelling and metabolism, the PaLemA1 and PaLemA2 interactomes were enriched in proteins involved in DNA and RNA metabolism, gene regulation, protein synthesis and cellular stress response (Extended Data Table 11, Extended Data chapter 17 with references), indicating that vesicle formation is integrated with global cellular physiology.

Key components of nucleotide metabolism and DNA replication were identified, including enzymes such as GMP synthase (GuaA), Uracil phosphoribosyltransferase (Upp) as well as the DnaN sliding clamp of the replication machinery, suggesting coordination of vesicle formation with DNA replication and cellular proliferation. Proteins involved in transcriptional regulation were also enriched, including NusG and RapA, the stationary phase and stress-associated sigma factor RpoS, the redox regulator Anr, the catabolite repression protein Crc, the histidine kinase PhoQ of the PhoPQ two component regulatory system, and the quorum-sensing controlled nucleoside hydrolyse Nuh (46).

RNA metabolism and translation were similarly represented. Factors controlling mRNA stability and turnover, such as the poly(A) polymerase PcnB and the RNA helicase RhlE2, were detected alongside proteins required for ribosome assembly and function, including EngA, RimP and EttA. Multiple aminoacyl-tRNA synthetases (AspS, GlyQ, MetG, GltX, TrpS) and the tRNA-modifying enzyme MnmG were also present, indicating that LemA-associated complexes are linked to active protein synthesis. A broad range of stress-response proteins was identified, including RecA (SOS response), the catalase KatA, the major cold shock proteins CspA and CapB, the acid-tolerance protein FxsA, glutathione hydrolase Ggt, the periplasmic osmo-stress proteins OsmY and glucans biosynthesis protein OpgG in combination with the periplasmic cyclophilin type peptidyl-prolyl cis-trans isomerase PpiA. In addition, factors associated with host interaction and immune evasion, including the lysozyme inhibitor MliC and the protease inhibitor Ecotin, were detected.

These findings suggest that LemA-associated complexes are embedded within a highly dynamic cellular network that integrates membrane remodelling with gene expression, protein homeostasis and stress adaptation. A detailed description is given in Extended Data chapter 18.

### LemA-associated regulators may coordinate vesicle formation-related gene expression

To determine how LemA-associated transcription regulators influence gene expression during vesicle formation, we compared the transcriptomes of ΔPa*lemA*1 and ΔPa*lemA*2 mutants with wildtype *P. aeruginosa* (Extended Data Table 12-14, explained in Extended Data chapter 18; the raw transcriptomic data will be deposited in the Gene Expression Omnibus (GEO) public repository upon acceptance).

In the ΔPa*lemA*1 mutant, genes encoding multiple transport systems (including *dppA1*, *dppA3*, *dppA4*, *dppC*, *dppF*, *gltF*, *gltK* and *braCF*), as well as metabolic enzymes such as components of the glycine cleavage system (*gcvP2H2*) and phenylalanine hydroxylase (*phhA*), were upregulated. Many of these genes are associated with LemA-interacting proteins and fall within the regulon of the catabolite repression regulator Crc (47), identified in the PaLemA interactome. In contrast, genes encoding components of the type III secretion system (*pscNO*, *pcr12DHV*, *exoUY*, *popBDN*), also under Crc control (48), were repressed, alongside with several genes involved in energy metabolism (*atpDEH*, *fbr*, *aceEF*, *cyoABCE*).

In the ΔPa*lemA*2 mutant, a distinct transcriptional response revealed the strong induction of around 30 genes of the *P. aeruginosa* Anr regulon (i. e. *arcABD*, *adhA*, *ccoDN*, *oprG*, *hemN*, *nirS*, *dnr*, *uspA, narXL*, Extended Data Table 12-14) (49, 50). Consistent with this, the redox-responsive regulator Anr was identified as a PaLemA interaction partner. In parallel, a large number of genes (∼ 160) were downregulated, including approximately 30 transcriptional regulators such as *nusG*, *rpoH*, *mvaT*, *chpD*, *phoB*, *prtR*, *wspF*, *ntrB*, *dsmR*, *feaR*, *cueR*, indicating widespread rewiring of gene expression networks.

The transcriptome of the double mutant (ΔPa*lemA*1+*A*2) combined features of both single mutants, suggesting partially overlapping but distinct regulatory roles for the two LemA proteins. Notably, the *pqsABCDE* operon, which encodes enzymes required for quinolone signalling and biofilm formation, was consistently upregulated in all mutants (51). This pathway is linked to the alternative sigma factor RpoS, another PaLemA1-associated regulator, further supporting a role for LemA proteins in coordinating stress and quorum-sensing responses (52). Interestingly, established extracellular factors such as the *Pseudomonas* quinolone signal (PQS) drive outer-membrane curvature via hydrophobic chemical intercalation, and as a consequence lytic events trigger membrane fragmentation. Several lines of evidence establish the LemA-assembled complex as a distinct, protein-machinery-driven structural system. Unlike passive surfactant-like blebbing or destructive explosive lysis, LemA proteins function as platforms spanning across the cell envelope that physically interface with core divisome components, peptidoglycan synthases, and the negative regulator CpoB. The observed upregulation of the *pqsABCDE* operon in *PalemA* mutants might reflect a compensatory regulatory response as part of a general stress response. Future investigation will shed light on the connection of both systems.

Together, these findings demonstrate that LemA-associated transcriptional regulators, including Anr, Crc and RpoS, coordinate gene expression programmes linked to vesicle formation, metabolism and stress adaptation, further reinforcing the central role of LemA proteins in integrating membrane remodelling with global cellular regulation.

### PaLemA-associated networks are enriched in essential genes

To associate the physiological importance of LemA-associated protein networks, we compared the PalemA interactome with previously defined essential gene sets in *P. aeruginosa*. Prior studies have identified 364 essential genes in strain PA14 (6.18% of all protein-coding genes) (53) and 352 in strain PAO1 (6.32%) (54). Notably, 49 out of the 254 PaLemA-associated proteins (∼19.29%) correspond to proteins encoded by essential genes, representing a substantial enrichment compared to the genomic baseline (Extended Data Tables 4 & 5).

### The alternative sigma factor AlgU controls Pa*lemA2* expression in response to envelope stress and growth phase

To define the environmental and regulatory inputs controlling PaLemA production, we examined transcriptional regulation of the genes for Pa*lemA1* and Pa*lemA2* in *P. aeruginosa*. Several regulatory proteins in the PaLemA interactome, including AlgU, AlgW, Crc, Anr and RpoS, suggested that PaLemA production may be linked to stress and signalling pathways (details in the Extended Data chapter 19, summarised in Extended Data Fig. 20) (55).

We therefore analysed promoter activity using GFP reporter fusions under a range of environmental conditions. In parallel, we searched for transcription factor binding sites using the PRODORIC database and the Virtual Footprint prediction tool (56). While no clear regulatory pattern was observed for Pa*lemA1*, consistent with the absence of a defined physiological role under the test conditions, Pa*lemA2* displayed a distinct and inducible expression profile.

Expression of Pa*lemA2* was strongly induced during the transition to stationary phase (Extended Data Figs. 21-23). In addition, conditions that disturb the cell envelope, such as a treatment with sub-inhibitory concentrations of the β-lactam antimicrobial meropenem or the peptidoglycan synthesis inhibitor D-cycloserine, triggered activation of the Pa*lemA2* promoter (Extended Data Figs. 22-23). Importantly, this induction was abolished in Δ*algU* mutant, demonstrating that the extracytoplasmic function (ECF) sigma factor AlgU-binding site is required for PalemA2 transcription under these conditions. Consistent with this, a putative AlgU binding site was identified upstream of Pa*lemA2*, overlapping the predicted -10/-35 promoter elements (Extended Data Fig. 20).

Further support for stress-dependent regulation came from biofilm-associated conditions: PaLemA2 mRNA abundance was elevated in a Δ*pel* mutant defective in exopolysaccharide production (57), indicating that alterations in cell envelope structure modulate its transcription. Together, these findings establish PaLemA2 as part of an AlgU-dependent envelope stress response pathway that is activated during stationary phase and cell wall perturbation. We propose that LemA proteins are embedded within this regulatory network, linking environmental sensing via AlgU to membrane remodelling and vesicle formation (Extended Data Fig. 20).

## Discussion

Bacterial membrane vesicle (BMV) formation and intracellular membrane remodelling are widespread yet mechanistically poorly understood processes (17). Here, we provide experimental evidence of the LemA protein family, including MamQ (22, 58), as a conserved membrane-remodelling system that organizes a large, multi-protein machinery integrating BMV biogenesis with core cellular processes. Our data support a model in which LemA proteins act as central organizational hubs that coordinate membrane curvature generation, envelope remodelling, and BMV release by recruiting and repurposing existing cellular machineries (30).

A central finding of this study is the functional specialization of the two LemA paralogues in *P. aeruginosa*. PaLemA1, anchored in the inner membrane, seems to promote the formation of intracellular membrane invaginations and cytoplasmic membrane vesicles (CMVs), whereas PaLemA2, associated with the outer membrane and extending into the periplasm, appears to drive outer membrane vesicle (OMV) formation (17, 59). This spatial separation suggests that LemA proteins define membrane-specific platforms that coordinate distinct but interconnected steps of BMV biogenesis. Importantly, these two systems are functionally linked through the periplasmic regulator PaCpoB (60) -a known modulator of the Tol-Pal system, cell wall elongation and outer membrane constriction during division-which interacts with both PaLemA proteins and acts as a critical modulator of BMV formation.

The identification of PaCpoB as a shared interaction partner and negative regulator provides a key mechanistic insight. Deletion of Pa*cpoB* leads to a dramatic increase in BMV production, whereas its overexpression suppresses vesiculation, indicating that CpoB constrains membrane deformation under normal conditions. In this context, PaCpoB likely acts as a checkpoint that coordinates peptidoglycan remodelling with membrane curvature, preventing uncontrolled BMV release that would compromise envelope integrity.

### A structural model for the PaLemA-PaCpoB hub

*In silico* structural modelling further supports this functional architecture. Predictions using AlphaFold suggest that PaLemA1 and PaLemA2 engage PaCpoB through distinct, non-overlapping interfaces, allowing the formation of a ternary PaLemA1-PaLemA2-PaCpoB complex. PaLemA1 appears to form an extended interaction surface with PaCpoB, whereas PaLemA2 engages more localized contact sites. These differences imply divergent binding modes that could enable differential regulation of inner and outer membrane processes. Although these models require experimental validation, they support the concept of a tripartite scaffold bridging both membranes via PaCpoB (Fig. 6).

**Figure 6.**
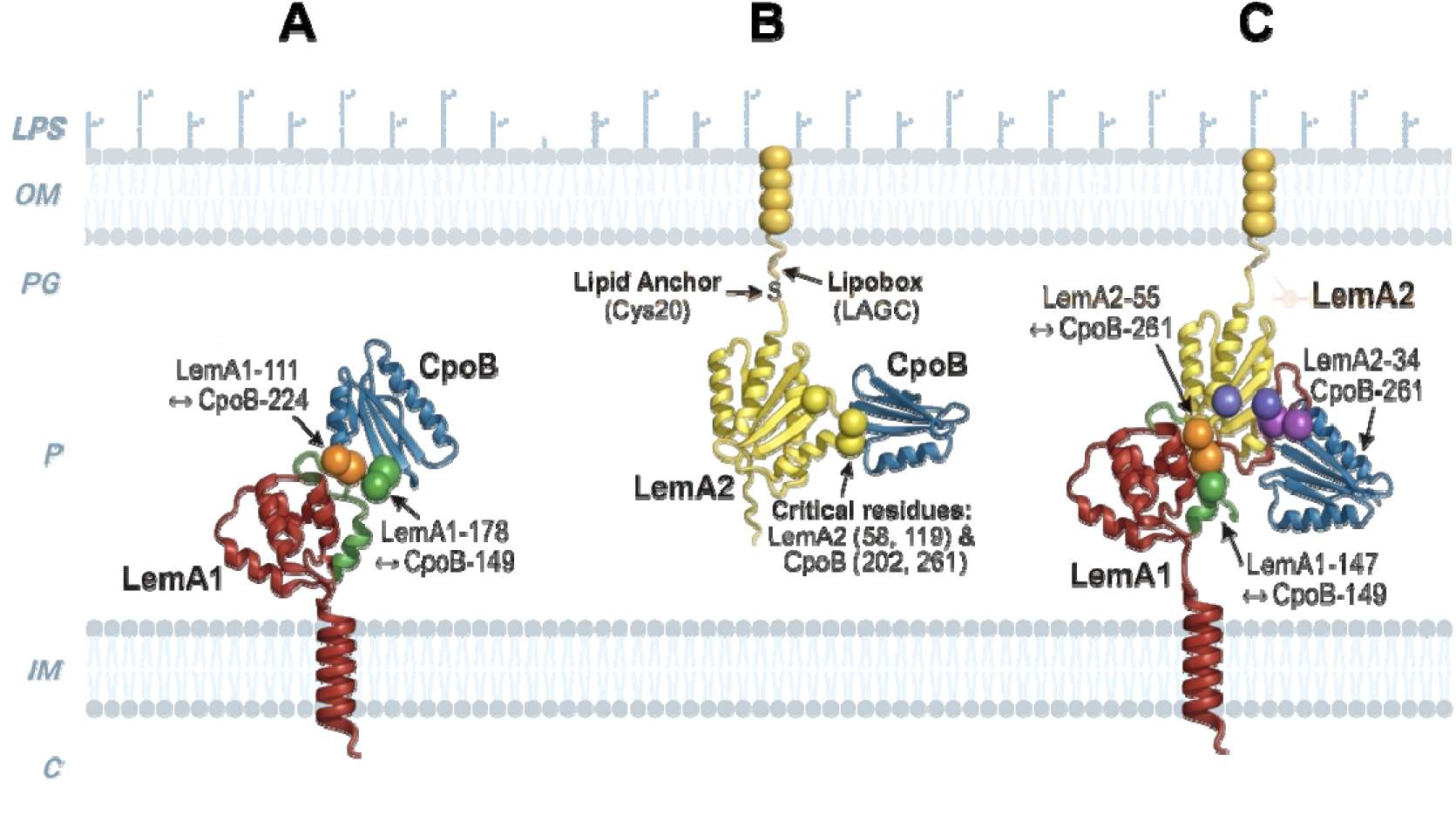
AlphaFold3-predicted interaction networks of PaLemA proteins with PaCpoB. (**A**) PaLemA1-PaCpoB binary complex. PaLemA1 (red, high confidence to light grey) and PaCpoB (blue, high confidence to light grey) are shown in cartoon representation. Key contact residues identified by distance-based analysis (<4 Å) are highlighted as spheres: Pair 1 (PaLemA1-111 _↔_ PaCpoB-224, orange) and Pair 2 (PaLemA1-178 _↔_ PaCpoB-149, green). Both contact pairs reside within high-confidence structural regions, supporting the predicted interaction interface. (**B**) PaLemA2-PaCpoB binary complex. PaLemA2 (red to light grey) interacts with PaCpoB (blue to light grey) through a discrete, minimal interface. Critical residues (PaLemA2-58, 119; PaCpoB-202, 261) are shown as yellow spheres. Interactomic distances: PaLemA2-58–PaCpoB-261 = 3.53 Å; PaLemA2-119-PaCpoB-202 = 3.76 Å. (**C**) PaLemA1-PaLemA2-PaCpoB ternary complex. Simultaneous binding of both PaLemA proteins to PaCpoB is structurally compatible. Colour scheme: PaLemA1 (red to light grey), PaLemA2 (green to light grey), PaCpoB (blue to light grey). Interaction pairs (<4 Å): PaLemA1-107 _↔_ PaCpoB-224 (orange spheres); PaLemA1-147 _↔_ PaCpoB-149 (green spheres); PaLemA2-55 _↔_ PaCpoB-261 (blue spheres); PaLemA2-34 _↔_ PaCpoB-261 (purple spheres). The spatial arrangement places PaLemA1 and PaLemA2 on distinct surfaces of PaCpoB, enabling simultaneous engagement and potentially bridging the inner and outer membranes across the periplasmic space. The PyMOL images were assembled in BioRender. The figure was created with BioRender.

### A hypothesis for LemA-mediated vesicle formation

Integrating our experimental and computational data, we propose a multi-layered model for LemA-dependent BMV biogenesis: **i) The scaffold:** PaLemA1 and PaLemA2 form a trans-envelope platform, with PaLemA1 organizing inner membrane remodelling and PaLemA2 coordinating outer membrane dynamics. PaCpoB connects both systems within the periplasm, forming a central regulatory hub; **ii) The initiators:** The ESCRT-III-like protein PaPspA is recruited to PaLemA1 and likely contributes to the initial generation of membrane curvature and sealing, analogous to its role in other membrane remodelling systems (61, 62); **iii) The construction machinery:** Core components of the cell division apparatus, including FtsZ-associated proteins, FtsEX (29), and peptidoglycan synthases (PBP1A, PBP1B, PBP3) (36, 63), are recruited to locally remodel the cell wall. This enables controlled thinning and restructuring of the peptidoglycan layer, facilitating membrane deformation and eventual BMV scission; **iv) The logistics network:** Lipid transport systems (AsmA, Mla) (64, 65) and LPS biosynthesis and transport machinery (Lpt system) (66) provide the necessary building blocks for membrane expansion, ensuring that BMV formation is coupled to envelope biogenesis **v) The regulatory module:** CpoB acts as a master regulator, modulating the activity of cell wall synthases (60, 67) and coordinating the transition between membrane stability and BMV release. Its loss leads to uncontrolled vesiculation, highlighting its role in maintaining balance within the system; and **vi) The support systems:** Protein export pathways (Sec and Tat), metabolic enzymes, and respiratory chain components ensure the supply of proteins, energy, and precursors (40, 41, 68) required for sustained membrane remodelling.

This hypothesis reframes BMV formation as a highly coordinated process that repurposes the bacterial cell division and envelope maintenance machinery, rather than relying on a dedicated, isolated pathway.

### Integration with metabolism and gene regulation

Beyond structural components, our data suggest that LemA-associated complexes - although requiring further binary interaction analysis for complete protein complex reconstruction-are deeply embedded in cellular metabolism and regulatory networks. . The enrichment of transport systems, central metabolic enzymes and respiratory chain components indicates that BMV formation occurs in metabolically active membrane regions and is coupled to energy availability. Similarly, the association with transcriptional regulators such as Anr, Crc and RpoS (48, 49, 52, 69, 70), together with the observed transcriptomic changes, demonstrates that BMV biogenesis is coordinated with cellular state and environmental conditions.

In particular, the regulation of Pa*lemA2* by the extracytoplasmic function sigma factor AlgU links BMV formation to envelope stress and stationary phase adaptation (55, 71, 72). This is consistent with previous observations that BMV production is enhanced under stress conditions and may serve protective roles, including the removal of damaged envelope components or the modulation of host-pathogen interactions (17).

### A distributed and adaptable vesicle biogenesis system

Taken together, our findings support a hypothesis in which LemA proteins act as central hubs within a distributed and adaptable membrane-remodelling network. By integrating divisome components, envelope biogenesis pathways, metabolic systems and regulatory circuits, this machinery enables bacteria to dynamically control membrane curvature and BMV production in response to physiological demands. Such an architecture provides robustness and flexibility, allowing BMV formation to be tuned without compromising cell viability.

### Implications and outlook

The identification of the LemA family as a conserved membrane-remodelling system has broad implications. First, it provides a unifying framework to understand diverse bacterial organelles and BMVs, from magnetosomes to extracellular BMVs (17, 73). Second, it highlights potential targets for manipulating BMV production in biotechnological applications, including vaccine development and drug delivery (4). Finally, given the role of BMVs in virulence, LemA-associated pathways may represent novel targets for antimicrobial intervention, although this will likely require an improved understanding of the molecular mechanism of LemA-mediated curvature generation, and thereby define the spatiotemporal dynamics of LemA complexes *in vivo*.

## Materials and Methods

### Construction of expression vectors for the production of LemA proteins in *E. coli* and *P. aeruginosa*

A full list of oligonucleotides and plasmids used in this study can be found in Extended Data Table 15. DNA fragments containing genes of interest were either generated via PCR using indicated primers or chemically synthesized as double stranded DNA (gBlocks, Integrated DNA technologies, Coralville, USA, Extended Data Table 16). The pET plasmids for protein production in *E. coli* BL21 Star (DE3) were then generated by ligating gene-containing fragments into the NdeI/SpeI site of a modified pET3a vector. The pSEVA vectors for protein production in *P. aeruginosa* were constructed by ligating the corresponding gene-containing fragments into BamHI/HindIII sites of pSEVA228 (Pa*pspA*) and the EcoRI/BamHI (Pa*lemA*1 and Pa*lemA2*) or BamHI/HindIII (Pa*cpoB*) sites of pSEVA634. Pa*lemA*2 fragment was then ligated into pSEVA634-Pa*lemA*1 BamHI/HindIII restriction sites to produce the co-expressing plasmid.

### Construction of the *P. aeruginosa* deletion and overexpression strains

The deletion of *P. aeruginosa* Pa*lemA1* and Pa*lemA2* were constructed using a scarless mutagenic system (74). The flanking regions of the *P. aeruginosa* PA14 Pa*lemA1* and Pa*lemA2* genes were PCR amplified using the corresponding primers. The resulting up- and downstream fragments of each gene were spliced by overlap extension (SOEing)-PCR (26) and cloned into pJET1.2 (ThermoFisher, Darmstadt, Germany). KpnI/XbaI and EcoRI/XbaI fragments from these plasmids were ligated into the suicide vector pSEVA612S, creating pSEVA612S-Pa*lemA*1 and pSEVA612S- Pa*lemA*2. These plasmids were subsequently introduced into *E. coli* DH5αλ*pir* for plasmid propagation. *E. coli* ST18-based biparental mating was used to transfer the vectors into *P. aeruginosa* PA14. Genomic integration was monitored by PCR. Next, the replicative plasmid pSEVA528S harbouring the I-*sceI* gene (isolated from *E. coli* DH5α) was transferred into the conjugated strains and induced by the addition of 15 mM toluic acid. Gene deletion in gentamicin sensitive colonies was confirmed by colony PCR (cPCR) before the pSEVA528S plasmid was cured.

The *P. aeruginosa* Pa*pelBCDEFG*, Pa*pspA* and Pa*cpoB* clean deletion mutants were generated using a recently developed CRISPR-Cas9-assisted recombineering methodology (75, 76). Briefly, two complementary oligonucleotides (Extended Data Table 15) were designed using the CRISPy-web server (77) to generate specific spacers targeting *P. aeruginosa*’s *pelB* (aapelB_PA14_spc1_S/aapelB_PA14_spc1_AS), *pspA* (pspA_PA14_spc1_S/pspA_PA14_spc1_AS) or *cpoB* (aaPau_cpoB_spc2-S/ aaPau_cpoB_spc2-AS). Next, the oligonucleotide pairs were phosphorylated, hybridized, and cloned into the BsaI-digested pS448·CsR according to the specifications from Pankratz and colleagues (75). The deletion was carried out in *P. aeruginosa* PA14 previously electroporated with the recombineering plasmid pSH624- *ssr*. This strain was grown overnight at 37 °C in 10 ml of LB medium supplemented with 50 µg/ml Gm. Next, 5 ml of fresh LB medium containing Gm and 1 mM IPTG were added to the cultures. After 2 h incubation at 37 °C, electrocompetent cells were prepared from 6 ml suspension of the induced cultures by consecutive washes with 0.3 M sucrose. Aliquots of 100 μl of cells were electroporated with 250 ng of the pS448·CsR-derivative cloned with the suitable spacer for each gene and 100 pmol of the pre-heated recombineering oligonucleotide (Extended Data Table 15) using the following settings: 200 Ω, 25 µF, 2.5 kV. Cells were recovered for 2 hours in 2 ml of LB medium containing Gm and 2 mM 3-*m*Bz for the induction of *cas9*. After incubation, cell dilutions were plated onto LB-Agar containing 500 µg/ml streptomycin, Gm, and 2 mM 3-*m*Bz to select for plasmids and to counter-select the wildtype cells by CRISPR-Cas9-induced DNA double-strand breaks. After 24 h of incubation at 37 °C, ΔPa*pelBCDEFG* (henceforth referred to as ΔPa*pel*), ΔPa*pspA* and ΔPa*cpoB* mutants were cPCR identified using primers flanking the genomic context of each mutation. Finally, successfully generated *P. aeruginosa* mutants were cured of plasmids by three consecutive passages in LB medium devoid of antibiotics.

### General culture conditions

The growth conditions for all experiments were conducted in biological triplicates. Unless otherwise indicated, *E. coli* and *P. aeruginosa* strains were cultured in LB at 37°C. For production of LemA proteins in *E. coli*, bacteria were grown to an OD_600nm_ of 0.5, gene expression induced using 100 μM isopropyl β-D-1-thiogalactopyranoside (IPTG) or 5 mM arabinose and further incubated at 19°C for 16 hours. *P. aeruginosa* cultures were inoculated to a starting OD_600nm_ of 0.05, grown to an OD_600nm_ of 0.5 after which protein production was induced using 100 µM IPTG or 3-*m*Bz and incubated for additional an 2 h. Strains without plasmids were inoculated equally but cultivated for 6 h without any additives. Antibiotics concentrations used were 15 μg mL^−1^ gentamicin (Gm), 150 μg mL^−1^ ampicillin (Ap), 50 μg mL^−1^ streptomycin (Sm), and 15 μg mL^−1^ tetracycline (Tc) for *E. coli*; and 50 μg mL^−1^ Gm, 500 μg mL^−1^ Sm, 500 μg mL^−1^ kanamycin (Km) and 100 μg mL^−1^ Tc for *P. aeruginosa* PA14.

### Bacterial membrane vesicles isolation from *P. aeruginosa*

All experiments were conducted in biological triplicates. For BMV isolation, *P. aeruginosa* strains of interest were inoculated in 20 mL LB, supplemented with antibiotics if needed and incubated over night at 37 °C and 160 rpm. Cells were subcultured by inoculating 100 mL LB to a final OD_600 nm_ of 0.05 and cultivated at 37 °C, 160 rpm. Plasmid-based overexpression of Pa*pspA* or Pa*lemA* or Pa*cpoB* was induced when the OD_600nm_ reached 0.5 by adding 100 µM 3-*m*Bz or IPTG, respectively. To avoid artifacts resulting from extended overproduction of proteins, bacteria were removed by centrifugation (8000 x g, 4 °C, 20 min) 2 h post induction and supernatant was sterile filtered using 0.2 µm PES filter. Deletion strains were cultured upon early stationary phase. Supernatants were concentrated using 100 kDa centrifugal concentrators [VivaSpin 20, 100 000 MWCO (Sartorius, Göttingen, Germany)] at 4 °C and 4000 x g. After resuspending the remaining material in 1 mL 1 x PBS, again 1 mL 1 x PBS was used to wash the filter. The concentrated samples were ultracentrifuged at 4 °C, 150 000 x g for 2 h [Optima L-90K Ultracentrifuge with Type T70.1 Ti Fixed-Angle Titanium Rotor (Beckman Coulter, USA)] and the supernatants were removed carefully without removing any brown BMV material. The BMV pellets were resuspended in a total volume of 1 mL 1 x PBS and sterile filtered with 0.20 µm syringe filters.

### Bacterial membrane vesicle quantification by fluorescent membrane staining

BMV yields were determined by detecting fluorescence signal emitted using membrane staining dye FM4-64 (Invitrogen, ThermoFisher Scientific, Waltham, USA). To obtain a stock solution of 50 µg mL^-1^, the dye was dissolved in water and kept at 4 °C. Before measurements, a working staining solution was prepared by diluting the stock solution in HBSS to a final concentration of 5 µg mL^-1^. Afterwards, samples were diluted 100-fold in the working staining solution. For measurements, 200 µL was transferred into a 96-well plate and fluorescence was immediately determined using a plate reader (Tecan Spark, Tecan Group, Männedorf, Switzerland) with the following settings: excitation at 588 nm, emission at 734, gain = 150, 30 flashes, Z-position = 20000 at room temperature. Each sample was measured in triplicates.

### PaLemA1 and PaLemA2 antibodies

Peptides covering the following the amino acid residues of PaLemA1 were selected to raise antibodies: amino acid residues 48-62 (DLVPNLVSTVKGYAS), 63-77 (HEASVLTQVTEARAK), 78-92 (VGSVQLNADQLDDEQ) and 101-116 (QGELSSALSRLLVVTE). For PaLemA2 they were 44 - 58 (LKQRHDELPKLVEA), 89 – 104 (NALGKAESGLRAGLGQL) and 168 - 182 (KAAELLQFSEAEKAD). Anti-LemA1 and anti-LemA2 antibodies were generated in rabbits by Metabion AG (Planegg/Steinkirchen, Germany).

### Field Emission Scanning Electron Microscopy (FESEM)

BL21 Star (DE3) colonies containing the plasmid of interest were used to inoculate 50 mL LB supplemented with 100 μg mL−1ampicillin in baffled flasks. Cells were grown at 37 °C with shaking to an OD600 ∼ 0.4, protein production was induced by addition of IPTG to a final concentration of 0.1 mM. Cultures were incubated overnight at 19 °C with shaking. *P. aeruginosa* strains were grown as specified above. Samples were fixed with 2% glutaraldehyde in growth medium, left for 30 minutes and then formaldehyde was added to a final concentration of 5%. After 1 hour on ice or standing overnight at 7°C fixed bacteria were washed twice for 10 minutes with TE buffer (20 mM TRIS, 1 mM EDTA, pH 6.9), dehydrated with a graded series of acetone (10, 30, 50, 70, 90%) for 15 minutes for each step on ice. Samples in the 100% acetone step were allowed to reach room temperature before another change to 100% acetone was applied. Samples were then subjected to critical-point drying with liquid CO2 (CPD 30, Bal-Tec, Liechtenstein). Dried samples were fixed onto aluminium stubs with a conductive adhesive tape and covered with a gold/palladium film by sputter coating (SCD 500 Bal-Tec, Liechtenstein) before examination in a field emission scanning electron microscope Zeiss Merlin (Zeiss, Oberkochen) using the Everhart Thornley HESE2 detector and the inlens SE detector in a 25:75 ratio with an acceleration voltage of 5 kV using the SEM software 5.5 and 6.06.

### Transmission electron microscopy (TEM) for negative stain and ultrathin sections

Bacteria were grown and BMVs were isolated as described above and fixed as described in the FESEM part. For the negative stain and TEM we followed the steps previously described (78). For ultrathin sections, after washing with TE buffer samples were incubated in 1% aqueous osmium tetroxide for 1 hour at room temperature. After washing with TE buffer bacteria/ BMVs were embedded into 2% agar, cut into small cubes and stepwise dehydrated with 10, 30, 50% acetone. Samples were then incubated overnight at 7 °C in 70% acetone containing 2% uranyl acetate. The following day samples were dehydrated with 90% and 100% acetone. Infiltration with LV resin was performed (1 part acetone/1 part LV resin, 1 part acetone/2 parts LV resin) before infiltration in pure resin with several changes. After polymerization at 70°C ultrathin sections of 40 - 70 nm in thickness were cut, counter-stained with uranyl acetate and lead citrate before examination in a Zeiss TEM 910 at an acceleration voltage of 80 kV. Images were recorded digitally at calibrated magnifications with a Slow-Scan CCD-Camera (ProScan, 1024×1024, Scheuring, Germany) with ITEM- Software (Olympus Soft Imaging Solutions, Münster, Germany). For negative staining a thin carbon film was floated on a sample droplet to allow adhesion of BMVs. After 1 minute, the film was taken off with a 300-mesh copper grid and washed twice with distilled water. Finally, the grid was applied to a droplet of 4% uranyl acetate (w/v), incubated for 1 minute and heat-dried after blotting the excessive liquids. Examination was performed with a Zeiss Libra 120 Plus equipped with Slow Scan CCD camera (Sharp:eye, 2048*2048; Tröndle, Germany) and at an acceleration voltage of 120kV. Images were digitally acquired at calibrated magnifications using the ITEM-Software.

### Immuno-labelling on ultrathin sections

Bacterial pellets were resuspended in 2% formaldehyde and 0.5% glutaraldehyde prepared in 100 mM sodium cacodylate buffer (pH 7.2) and incubated for 2 h with gentle rotation. Following fixation, cells were recovered by centrifugation at 6,000 x g for 2 min and washed twice for 10 min in 100 mM sodium cacodylate buffer (pH 7.2).

Samples were subsequently dehydrated through a graded ethanol series consisting of 50%, 70%, and 90% ethanol for 10 min each, followed by three washes in 100% ethanol, each lasting 15 min. Cell pellets were infiltrated with 2 mL LR white resin and incubated overnight at room temperature with continuous rotation. The resin was then replaced with fresh LR white resin and samples were incubated for an additional 6 h. Pellets were resuspended in fresh resin, transferred into 1-mL gelatin capsules, and centrifuged at 4,000 x g to concentrate the cells at the capsule tip. Polymerization was carried out at 60°C for 24 h.

Ultrathin sections (60-70 nm) were prepared using an RMC MT-XL ultramicrotome fitted with a DiATOME 45° diamond knife and collected onto 300-mesh gold grids.

Prior to immunolabelling, grids were equilibrated on a drop of TBST (20 mM Tris-HCl, pH 7.2, containing 500 mM NaCl, 0.05% Tween 20, and 0.1% BSA) and subsequently blocked in 2% BSA prepared in TBST for 30 min at room temperature. Grids were then transferred directly onto 20 μL drops containing the appropriate primary antibody (anti-PaLemA1 or anti-PaLemA2; see above) and incubated for 1 h. Following incubation, grids were washed in fresh TBST and briefly rinsed for 10 s in a stream of TBST. For secondary detection, grids were equilibrated in drops containing the corresponding gold-conjugated secondary antibodies (10 nm goat anti-rabbit IgG; Agar Scientific) and incubated for 30 min in fresh antibody solution. Excess antibody was removed by washing the grids twice in TBST, followed by rinsing in a stream of ddH2O and air-drying.

Sections were contrasted by staining with 4.5% uranyl acetate in 1% acetic acid for 15 min, followed by two washes with dH_2_O. Grids were then stained with Reynolds lead citrate for 3 min and rinsed with ddH_2_O. Imaging was performed using a JEOL-1230 transmission electron microscope equipped with a Gatan Multiscan digital camera operated at an accelerating voltage of 80 kV.

### Affinity co-purification of PaLemA1 and PaLemA2 interaction partners

Affinity chromatography coupled with mass spectrometry was employed to determine the interaction partners of PaLemA1 and PaLemA2 proteins. To that end, recombinant versions of Pa*lemA1* and Pa*lemA2* C-terminally fused to a strep tag II were used. The expression vector employed for this experiment was the IPTG-inducible pSEVA634, which was introduced in each of the corresponding background mutants. An overnight culture was grown in LB medium supplemented with 50 μg/mL gentamycin. The main culture was inoculated to an OD_600nm_ of 0.05 and grown without antibiotics for 2 hours. 1 mM IPTG was added at this time point and the culture was further cultivated for 3 hours (late exponential growth phase). At this stage, the culture was supplemented with 0.125 % (v/v) formaldehyde with a syringe and further incubated for 20 min at 37°C and 180 rpm. 135 mM glycine was added to quench formaldehyde excess and incubated for 5 min at 37°C and 150 rpm. Cells were subsequently harvested by at 7000 x g for 20 min at 4°C. The cell pellet was washed twice with phosphate buffered saline (PBS; 137 mM NaCl, 2.7 mM KCl, 4.3 mM Na_2_HPO_4_ × 7 H_2_O, and 1.4 mM KH_2_PO_4_) at 7000 x g for 20 min at 4°C. Cells were resuspended in 10 mL PBS buffer. The cell suspension was supplemented with 1.5 mL of 70-100 μm glass beads. Cells were lysed by FastPrep24 grinder system (mpbio, USA) for 30 sec with each 6.5 m/s velocity twice. Whole cell extracts were centrifuged for 20 min at 4°C and 4000 x g to remove glass beads. The supernatant was transferred into an ultracentrifugation tube (10.4 mL, polycarbonate bottle, 16 x 76 mm, Beckman Coulter) and centrifuged for 1 h at 150.000 x g (Type 70.1 Ti, Beckman Coulter) and 4°C. The supernatant was disposed the membrane pellet was resuspended in 4 mL solubilization buffer (18.8 mM K2HPO4; 1.2 mM KH2PO4 pH 8; with 1.8 % triton X-100). Solubilization of membrane-residing proteins was achieved at 4°C overnight. For the purification of Strep-tag^®^II-recombinant proteins the manufactureŕs guidelines were followed (IBA Lifesciences, Germany). The eluates were concentrated with a centrifugal filter with a MWCO of 10 kDa (Amicon Ultra 0.5 mL 10 kDa MWCO, Millipore, Germany) until a final volume of 200 μL was achieved.

### Validation of binary interactions by two-hybrid system

To investigate protein-protein interactions, we employed the bacterial two-hybrid (BACTH) system based on the reconstitution of the adenylate cyclase catalytic domain of *Bordetella pertussis*, as previously described (79). DNA sequences encoding the proteins PaLemA2, PaLemA1, PaCpoB, PaMrcB, and PaFtsN were PCR-amplified and cloned into the BACTH vectors pUT18, pUT18C, pKT25, or pKNT25 to generate N-terminal and C-terminal fusions with the T18 and T25 catalytic subdomains. When appropriate monomers interact, the catalytic subdomains restore adenylate cyclase, leading to intracellular cAMP synthesis and activation of the lac operon.

The periplasmic domain of PaLemA2 (a.a. 21-202), lacking its N-terminal signal peptide (a.a. 1-20), was amplified using primer pairs lemA2-Fw/lemA2(18C)-Rv and lemA2-Fw/lemA2(25)-Rv. The resulting amplicons were cloned into pUT18C and pKT25, respectively, generating the constructs pUT18C-LemA2 and pKT25-LemA2. The DNA sequence corresponding to the PaLemA1 cytoplasmic domain (a.a. 28-190), excluding the N-terminal periplasmic and transmembrane region (a.a. 1-27), was amplified with primers lemA1-Fw/lemA1(18C)-Rv and lemA1-Fw/lemA1(25)-Rv. These products were cloned into pUT18C and pKT25 to generate the constructs pUT18C-LemA1 and pKT25-LemA1. The periplasmic domain of PaCpoB (a.a. 20-254), excluding the predicted signal peptide (a.a. 1-19), was amplified using various primer combinations to be cloned into different BACTH vectors. The primers CpoB-Fw/CpoB(18/N25)-Rv were used for cloning into pUT18 and pKNT25, CpoB-Fw/CpoB(25)-Rv for pKT25, and CpoB-Fw/CpoB(18C)-Rv for pUT18C. This strategy resulted in the plasmids pUT18-CpoB, pUT18C-CpoB, pKT25-CpoB, and pKNT25-CpoB. The cytoplasmic domain of PaMrcB (a.a. 1-22) was amplified using MrcB-Fw and MrcB-Rv primers, and the amplicon was cloned into pKNT25 and pUT18 to generate pKNT25-MrcB and pUT18-MrcB, respectively. Finally, the periplasmic domain (a.a. 44-231) of PaFtsN was amplified using primer pairs FstN-Fw/FstN(25)-Rv and FstN-Fw/FstN(18C)-Rv and cloned into pKT25 and pUT18C to generate the constructs pKT25-FtsN and pUT18C-FtsN.

All BACTH vectors (pUT18, pUT18C, pKT25, and pKNT25) were linearized with the restriction enzyme SmaI prior to cloning. Gibson Assembly reactions were carried out according to the manufacturer’s protocol to insert the amplified DNA fragments into the corresponding vectors. The resulting plasmids were co-transformed into *Escherichia coli* BTH101, and the resulting colonies were assessed by the development of blue coloration on LB agar plates containing X-gal and by measuring β-galactosidase activity. Control experiments were carried out co-transforming pKT25-zip and pUT18C-zip. Co-transformed cells were grown in 3 ml of LB in the presence of 0.5mM IPTG and appropriate antibiotics at 30°C and vigorous agitation overnight. A volume of 15 µL was spotted onto LB agar plates supplemented with X-gal (40 µg/mL), IPTG (0.5 mM), and the appropriate antibiotics, and incubated at 30 °C for 24-48 hours. For the β-galactosidase assay, overnight cultures were diluted and grown to OD₆₀₀ ≈ 0.5-1.0. Then, 0.2 mL of culture was collected and the OD_600nm_ recorded. Cells were permeabilized by adding 10 µL of 0.1% SDS and 20 µL of chloroform, and 800 µL of Z-buffer (104 mM Na2HPO4 7 H2O, 45 mM NaH2PO4 H2O, 10 mM KCl, 1 mM MgSO4 7 H2O) followed by vortexing for 15 seconds. Then, 200 µL of ONPG (4 mg/mL in Z-buffer) was added and the reaction until a yellow colour developed. The reaction was stopped by adding 500 µL of 1 M Na₂CO₃. Units of β-galactosidase activity were calculated according to the method of Miller (80).

### RNA isolation and transcriptomic analysis

Bacterial cultures (in biological triplicates) were prepared by diluting overnight-grown cells to OD_600_ 0.05 in 20 mL of LB medium, followed by incubation at 37°C with 180 rpm shaking until mid-exponential phase (OD_600_ = 0.5-0.6). For RNA isolation, 1 mL of each sample was mixed with the same volume of RNA protect (Qiagen, Hilden, Germany), incubated for 10 min at room temperature and spun down for 10 min at 16,000 x G to collect the cell pellet. Total RNA was isolated with the RNeasy Mini Kit (Qiagen, Hilden, Germany) in combination with Qiashredder columns (Qiagen, Hilden, Germany) as described previously (81). Ribosomal RNA was depleted using the RiboZero Gold Kit (Illumina, San Diego, CA, USA) and the cDNA libraries were generated with the ScriptSeq v2 kit (Illumina). The samples were sequenced as 50 nt single-end reads on an Illumina *HiSeq 2500* instrument. Sequencing reads were mapped to the *P. aeruginosa* UCBPP-PA14 genome (NC_008463.1) with *bowtie2* (82), and the reads per gene were extracted using *featureCounts* (83). Differential gene expression analysis between the mutant strains *vs* PA14 wildtype was computed from the reads per gene using the *R* package *edgeR* (84). The obtained *p*-values for differential expression were adjusted to account for the FDR using the method by Benjamini and Hochberg (85). Only genes with an adjusted *p*-value < 0.05 were considered differentially expressed. Raw Illumina reads will be available at the Gene Expression Omnibus (GEO) after acceptance of the manuscript.

### Fluorescence measurements in cells expressing *P_lemA1_* and *P_lemA2_* transcriptional reporters

The putative promoter regions (circa 320 bp) of Pa*lemA1* and Pa*lemA2* were PCR amplified using primer pairs aaP56810Fw/aaP56810Rev and aaP06990Fw/aaP06990Rev, respectively (Extended Data Table 15). Both primer pairs include KpnI and XbaI restriction sites to facilitate each promoter insertion into the same sites of the GFP-bearing plasmid pSEVA637. The resulting plasmids were electroporated into *P. aeruginosa* strains prepared with 0.3 M sucrose as described above. To measure the activity of the promoters along the growth curve, *P. aeruginosa* cells harbouring the reporter plasmids were grown overnight at 37°C in LB medium supplemented with Gm. Following, cultures were prepared in 20 mL of LB medium devoid of antibiotics at OD_600_ 0.05 and grown at 37°C under constant agitation. Additionally, for the samples treated with cell-wall biosynthesis inhibitors, either 0.025 mg mL^-1^ meropenem or 100 mg mL^-1^ cycloserine was added to the cultures (which corresponds to 1:10 of the calculated minimal inhibitory concentration or MIC for each antibiotic, respectively). Cells were sampled every two hours, diluted in filter-sterilized PBS (0.22 nm pore size), and the GFP signal was measured with a CytoFLEX flow cytometer (Beckman Coulter, California, USA) using a 488 nm excitation and a 525 nm (40 nm BP filter) fluorescence recovery. A final GFP measurement was also recorded after 24 hours of growth.

To measure the promoter expression in colony biofilm-grown cells, the reporter strains tested above were grown overnight in LB medium and 10 ml from each culture were spotted onto tryptone-agar plates (1% tryptone, 1% agar) until dry. The colonies were incubated at 25°C and samples were taken from 2-5 days post-inoculation. The sampling was performed by scooping the biomass from the colony with a sterile inoculation loop, followed by its resuspension in filter-sterilized PBS by thorough vortexing. The OD_600nm_ of the suspension was measured and the cells were diluted in 1 ml PBS to obtain an approximate density of 10^6^ cells mL^-1^. Finally, the GFP fluorescence of 20,000 events was measured with a CyFlow^®^ Cube 8 (Partec GmbH, Münster, Germany) flow cytometer using a 488 nm excitation and a 536 nm (40 nm BP filter) fluorescence recovery.

The resulting data were processed using FlowJo v. 9.6.2 software (FlowJo LLC, Ashland, OR, USA). The mean GFP fluorescence of the population was used to calculate the promoter activity in all tested samples.

### Mass spectrometry by LC-MS/MS

For proteomic analyses using LC-MS/MS, isolated BMVs or (co-)purified PaLemA1/PaLemA2 protein fractions were solubilized using 1% SDS, reduced for 30 minutes with 5 mM tris(2-carboxyethyl)phosphine (TCEP) at 56°C and alkylated with 10 mM methyl methanethiosulfonate (MMTS) for 15 min. Thereafter, a single-pot solid-phase-enhanced sample preparation (SP3) protocol (27,28) was employed before trypsin (Promega, Fitchburg, WI, USA) was added (1:50 w/w) in 50 mM HEPES (pH 8), and samples were incubated overnight at 37°C. Digested peptides were allowed to bind to the beads by applying >95% acetonitrile (ACN) and incubation overnight, then beads were three times washed with 100% ACN and air dried. Peptides were eluted with 20 µl 2% dimethyl sulfoxide (DMSO) for 30 min, vacuum dried and dissolved in 0.1% formic acid, and allowed to resolve using an ultrasound water bath. Samples were centrifuged for 20 min at 20.000 rpm at 20°C, and supernatants were transferred to new LoBind tubes. For each sample, 200ng peptide mixture was applied to C18 Evotips (EV-2001; Evosep) according to the manufacturer’s protocol. Evotips were then loaded on an Evosep One HPLC (Evosep) connected to a TimsTOF*Pro* mass spectrometer (Bruker). The Evosep One HPLC was operated with the standard 60 sample per day method (21 min gradient at a flow rate of 1.0 µl/min; buffer A: 0.1% formic acid, buffer B: 0.1% formic acid in acetonitrile). For the TimsTOF*Pro* mass spectrometer, the standard MSMS Bruker method “DDA PASEF method for short gradients with 0.5 s cycletime” was employed. MS settings in detail were: scan begin 100 m/z; scan end 1700 m/z; ion polarity: positive; scan mode: PASEF. Tims settings were: mode custom; number of PASEF ramps: 4; charge minimum: 0; charge maximum: 5; 1/K0 start 0.75 V*s/cm2; 1/K0 end 1.4 V*s/cm2; ramp time: 100.0 ms; MS average: 1. Raw data files were analyzed using the PEAKS software (PEAKS studio version 10.6). Detailed PEAKS settings were; parent mass error tolerance: fragmentation mode: high energy CID (y and b ions); 20.0 ppm; fragment mass error tolerance: 0.03 Da; enzyme: trypsin; max. missed cleavages: 1; fixed-modifications: beta-methylthiolation C (45.99); variable modifications: oxidation M (15.99); False discovery rate (FDR): 1%; Database; PA14 (release 05.2019). Data analyses, data presentation and statistics were performed using R, Perseus (version 2.1.3.0) and GraphPad Prism 10. All interactomic experiments were conducted in biological triplicates. The mass spectrometry proteomics will be deposited to the ProteomeXchange Consortium via the PRIDE (86) partner repository upon acceptance.

### Homologous Sequence Identification, Phylogenetic and ecological distribution Analysis of LemA

A BLASTp search was performed using the *Pseudomonas aeruginosa* UCBPP-PA14 PaLemA protein (PA14_56810) as the query against the Progenomes v2.0 database, which contains representative sequences from over 12,000 prokaryotic species (24). The search identified 2,566 LemA homologs across the database with a sequence identity threshold of >30%, a coverage threshold of >75%, and an E-value cutoff of <10⁻ . LemA homolog sequences were then aligned using the MAFFT v7.525 G-INS-i algorithm (87), and poorly aligned regions were removed with TrimAl v1.5 using the *automated1* mode (88). The phylogenetic tree was constructed using FastTree v2.1 with the LG substitution model and default parameters (89). To optimize the tree topology, the SPR algorithm was applied with a maximum of 4 subtree pruning and regrafting steps *(-spr 4*). Bootstrap support values were calculated using 1000 bootstrap replicates. The resulting tree was visualized and analyzed using iTOL (90) for graphical representation.

To investigate the ecological distribution of LemA homologues, genomes containing LemA1, LemA2, or both homologues (LemA1-A2) were classified according to their annotated habitat metadata (Extended Data Table 1). Habitat assignments were obtained from the ProGenomes2 database, which provides standardized ecological categories for sequenced prokaryotic genomes. Eight habitat categories were considered: soil, aquatic, aquatic sediment, freshwater, host-associated, plant-associated, food-associated, and disease-associated environments.

For each LemA category (LemA1, LemA2 and genomes containing both homologues), the number of genomes assigned to each habitat was counted. Habitat proportions were calculated by normalizing these counts to the total number of genomes within each category that had an available habitat annotation. This normalization step was applied because not all genomes in the ProGenomes2 database include habitat metadata.

The resulting habitat distributions were compared against the global ecological distribution of genomes in the ProGenomes2 reference dataset. To test whether LemA-associated genomes were significantly enriched or depleted in particular habitats relative to the background distribution, contingency tables were constructed for each habitat and LemA category (Extended Data Table 2). Statistical significance was assessed using Fisher’s exact test and resulting P-values were corrected for multiple testing using the Benjamini–Hochberg false discovery rate (FDR) procedure (Extended Data Table 3).

### Bioinformatic Prediction of Membrane Topology and Subcellular Localization

Signal peptides were predicted using SignalP 6.0 with default parameters for Gram-negative bacteria. Lipoprotein signal peptides (Sec/SPII) were distinguished from classical Sec/SPI signals based on lipobox motif detection and predicted cleavage sites. Transmembrane helices and membrane topology were predicted using DeepTMHMM. The orientation of predicted transmembrane segments was assessed by analysing the distribution of positively charged residues flanking the helix according to the positive-inside rule. Subcellular localization was predicted using ProtT5, a protein language model trained for bacterial localization prediction (cytoplasmic, cytoplasmic membrane, periplasmic, outer membrane). Predictions generated by eukaryotic-trained models (e.g., TargetP 2.0), including ER, mitochondrial, or lysosomal/vacuolar targeting signals, were disregarded as non-applicable in the bacterial context. Non-classical (leaderless) secretion was evaluated using SecretomeP 2.0 with the recommended bacterial threshold score of 0.5. Lipoprotein predictions were independently validated using LipoP 1.0 to discriminate between signal peptides and transmembrane helices. All topology and localization analyses were interpreted within the structural framework of Gram-negative cell envelope organization (inner membrane–periplasm–outer membrane).

### Structural Alignment of LemA Homologs and MamQ

Predicted structural models were obtained from the AlphaFold Protein Structure Database for: PaLemA1 (UniProt Q93DY8); PaLemA2 (UniProt Q9I3S4); TmLemA1 (UniProt Q9X056); MamQ (Magnetospirillum gryphiswaldense; UniProt Q93DY8).

Pairwise structural alignments were performed using the PyMOL “align” algorithm (PyMOL Molecular Graphics System, version 2.5, Schrödinger LLC). Only residues with predicted Local Distance Difference Test (pLDDT) scores > 70 were included to restrict comparisons to high-confidence structural regions and exclude poorly modelled loops or terminal segments. Root-mean-square deviation (RMSD) values were calculated over aligned Cα atoms. Structural coverage was defined as the percentage of Cα atoms in the shorter structure that could be structurally aligned to the reference model. All pairwise combinations were analysed: TmLemA vs PaLemA1, TmLemA vs PaLemA2, TmLemA vs MamQ, PaLemA1 vs PaLemA2, PaLemA1 vs MamQ, and PaLemA2 vs MamQ. Structures were visualized in PyMOL with pLDDT-based confidence colouring.

### Protein Structure Prediction and Interaction Analysis

Structural models of PaLemA1, PaLemA2, and PaCpoB from *Pseudomonas aeruginosa* PA14 were generated using AlphaFold3 via the AlphaFold Server platform with default settings.

Binary complex predictions were performed for PaLemA1-PaCpoB and PaLemA2-PaCpoB using multimer mode. A ternary complex (PaLemA1-PaLemA2-PaCpoB) was predicted using AlphaFold3 multimer modelling with all three sequences as input.

For each system, the top-ranked model (rank_0) was selected based on global confidence metrics and compatibility with predicted membrane topology. Five independent prediction runs (distinct random seeds) were performed per complex to assess reproducibility of interaction interfaces. Only interaction patterns reproducible across independent runs were considered for further analysis.

Per-residue confidence was evaluated using pLDDT scores (0–100 scale) and classified as: High confidence: ≥ 90; Medium confidence: 70–89; Low confidence: 50– 69; Very low confidence: < 50. Only interaction interfaces involving residues with medium-to-high confidence (pLDDT ≥ 70) were interpreted. Intermolecular contacts were identified using a distance-based criterion. Residues were considered interacting if any heavy-atom pair between chains was within 4.0 Å. For each interaction, residue pairs and minimum interatomic distances were recorded. Interface residues were mapped onto three-dimensional models to assess clustering and surface distribution. For ternary complexes, pairwise contact analyses were performed for each chain combination.

### Structural Visualization and Schematic Representation

All structural models were visualized using PyMOL (version 2.5, Schrödinger LLC). Proteins were rendered in cartoon representation with manual per-residue colouring according to pLDDT confidence categories. Interaction residues were highlighted as spheres centred on representative atoms. Colour coding was applied consistently across binary and ternary complexes to distinguish specific contact pairs. Topology cartoons and conceptual membrane orientation schematics were generated from the combined topology and localization predictions obtained using SignalP 6.0, DeepTMHMM, ProtT5, SecretomeP 2.0, and LipoP 1.0. Structural models were visualized in PyMOL (version 2.5, Schrödinger LLC), and simplified publication-ready schematics were created using BioRender. These illustrations are conceptual representations intended to summarize predicted membrane topology and domain organization and should not be interpreted as experimentally determined structural models.

## Acknowledgments

We are grateful to Ina Brentrp (Helmholtz Centre for Infection Research, Braunschweig, Germany) for the technical support in the electron microscopy experiments. We are indebted to Roberto Balbotín Soria (Universidad de Sevilla, Spain) for his assistance, technical support and providing the equipment for conducting the flow cytometry analyses.

## References

1. Lopez-Marques RL, Perez-Castineira JR, Losada M, Serrano A. 2004. Differential regulation of soluble and membrane-bound inorganic pyrophosphatases in the photosynthetic bacterium *Rhodospirillum rubrum* provides insights into pyrophosphate-based stress bioenergetics. J Bacteriol 186:5418–26.

2. Toyofuku M, Morinaga K, Hashimoto Y, Uhl J, Shimamura H, Inaba H, Schmitt-Kopplin P, Eberl L, Nomura N. 2017. Membrane vesicle-mediated bacterial communication. ISME J 11:1504–1509.

3. Ellis TN, Kuehn MJ. 2010. Virulence and immunomodulatory roles of bacterial outer membrane vesicles. Microbiol Mol Biol Rev 74:81–94.

4. Munoz-Echeverri LM, Benavides-Lopez S, Geiger O, Trujillo-Roldan MA, Valdez-Cruz NA. 2024. Bacterial extracellular vesicles: biotechnological perspective for enhanced productivity. World J Microbiol Biotechnol 40:174.

5. Arce-Rodriguez A, De la Pena Noya J, Borrero-de Acuna JM. 2026. Bacterial Extracellular Vesicles in Biomedical Research and Clinical Translation. Microb Biotechnol 19:e70350.

6. Stewart GE, Guo Y, Cao S. 2026. Emerging bacterial membrane vesicles from microbial messengers to therapeutic nanocarriers. J Control Release 398:115250.

7. Chen Y, Yuan X, Huang Y, Wei Q, Zhang X. 2026. Recent advances in hybrid membrane vesicles as a programmable biomimetic drug delivery. J Control Release 398:115244.

8. Schwechheimer C, Kuehn MJ. 2015. Outer-membrane vesicles from Gram-negative bacteria: biogenesis and functions. Nat Rev Microbiol 13:605–19.

9. Beveridge TJ. 1999. Structures of gram-negative cell walls and their derived membrane vesicles. J Bacteriol 181:4725–33.

10. Mashburn-Warren LM, Whiteley M. 2006. Special delivery: vesicle trafficking in prokaryotes. Mol Microbiol 61:839–46.

11. Turnbull L, Toyofuku M, Hynen AL, Kurosawa M, Pessi G, Petty NK, Osvath SR, Carcamo-Oyarce G, Gloag ES, Shimoni R, Omasits U, Ito S, Yap X, Monahan LG, Cavaliere R, Ahrens CH, Charles IG, Nomura N, Eberl L, Whitchurch CB. 2016. Explosive cell lysis as a mechanism for the biogenesis of bacterial membrane vesicles and biofilms. Nat Commun 7:11220.

12. Sangiorgio G, Nicitra E, Bivona D, Bonomo C, Bonacci P, Santagati M, Musso N, Bongiorno D, Stefani S. 2024. Interactions of Gram-Positive Bacterial Membrane Vesicles and Hosts: Updates and Future Directions. Int J Mol Sci 25.

13. Wang H, Chen Y, Huang H, Chen R, Chen S, Huang X. 2026. Recent advances in membrane vesicles of *Streptococcus mutans*: biogenesis, regulations, and functions. J Oral Microbiol 18:2735102.

14. Liang S, Gao N, Fei F, Jin X, Liu J, Zhang X, Du X. 2026. Targeting Outer Membrane Vesicles of *Pseudomonas aeruginosa*: From Pathogenesis to Therapeutics. J Microbiol Biotechnol 36:e2605019.

15. Zhang Q, Zhang B, Aazmi MS, Chen L, Yahya M. 2026. Bacterial Extracellular Vesicles at the Crossroads of Immune Regulation and Biofilm Dynamics: Biogenesis, Comparative Analysis, and Translational Challenges. Biomolecules 16.

16. 16. Kim B, Son Y, Yang J, Park W. 2026. New insights into outer membrane vesicles in Gram-negative bacteria from biogenesis to applications. Commun Biol 9.

17. Toyofuku M, Schild S, Kaparakis-Liaskos M, Eberl L. 2023. Composition and functions of bacterial membrane vesicles. Nat Rev Microbiol 21:415–430.

18. Schlosser L, Sachse C, Low HH, Schneider D. 2023. Conserved structures of ESCRT-III superfamily members across domains of life. Trends Biochem Sci 48:993–1004.

19. Liu M, Liu Y, Song T, Yang L, Qi L, Zhang YZ, Wang Y, Shen QT. 2024. Three-dimensional architecture of ESCRT-III flat spirals on the membrane. Proc Natl Acad Sci U S A 121:e2319115121.

20. Komeili A, Vali H, Beveridge TJ, Newman DK. 2004. Magnetosome vesicles are present before magnetite formation, and MamA is required for their activation. Proc Natl Acad Sci U S A 101:3839–44.

21. Barber-Zucker S, Keren-Khadmy N, Zarivach R. 2016. From invagination to navigation: The story of magnetosome-associated proteins in magnetotactic bacteria. Protein Sci 25:338–51.

22. Raschdorf O, Forstner Y, Kolinko I, Uebe R, Plitzko JM, Schuler D. 2016. Genetic and Ultrastructural Analysis Reveals the Key Players and Initial Steps of Bacterial Magnetosome Membrane Biogenesis. PLoS Genet 12:e1006101.

23. Nudelman H, Zarivach R. 2014. Structure prediction of magnetosome-associated proteins. Front Microbiol 5:9.

24. Mende DR, Letunic I, Maistrenko OM, Schmidt TSB, Milanese A, Paoli L, Hernandez-Plaza A, Orakov AN, Forslund SK, Sunagawa S, Zeller G, Huerta-Cepas J, Coelho LP, Bork P. 2020. proGenomes2: an improved database for accurate and consistent habitat, taxonomic and functional annotations of prokaryotic genomes. Nucleic Acids Res 48:D621–D625.

25. 25. Muller J, Kretschmer M, Opitsch E, Holland S, Borrero-de Acuna JM, Jahn D, Neumann-Schaal M, Wegner A. 2025. Bacterial membrane vesicles of Pseudomonas aeruginosa activate adenosine monophosphate-activated protein kinase signaling through inhibition of mitochondrial complex III. PNAS Nexus 4:pgaf248.

26. Kretschmer M, Muller J, Henke P, Otto V, Rodriguez AA, Musken M, Jahn D, Borrero-de Acuna JM, Neumann-Schaal M, Wegner A. 2023. Isolation and Quantification of Bacterial Membrane Vesicles for Quantitative Metabolic Studies Using Mammalian Cell Cultures. Cells 12.

27. Morrison JJ, Camberg JL. 2024. Building the Bacterial Divisome at the Septum. Subcell Biochem 104:49–71.

28. Tisma M, Kaljevic J, Gruber S, Le TBK, Dekker C. 2024. Connecting the dots: key insights on ParB for chromosome segregation from single-molecule studies. FEMS Microbiol Rev 48.

29. Alcorlo M, Martinez-Caballero S, Li J, Sham LT, Luo M, Hermoso JA. 2024. Modulation of the lytic apparatus by the FtsEX complex within the bacterial division machinery. FEBS Lett 598:2836–2851.

30. Cameron TA, Margolin W. 2024. Insights into the assembly and regulation of the bacterial divisome. Nat Rev Microbiol 22:33–45.

31. Mehla J, Liechti G, Morgenstein RM, Caufield JH, Hosseinnia A, Gagarinova A, Phanse S, Goodacre N, Brockett M, Sakhawalkar N, Babu M, Xiao R, Montelione GT, Vorobiev S, den Blaauwen T, Hunt JF, Uetz P. 2021. ZapG (YhcB/DUF1043), a novel cell division protein in gamma-proteobacteria linking the Z-ring to septal peptidoglycan synthesis. J Biol Chem 296:100700.

32. Hirakawa H, Suzue K, Tomita H. 2022. Roles of the Tol/Pal System in Bacterial Pathogenesis and Its Application to Antibacterial Therapy. Vaccines (Basel) 10.

33. Bertsche U, Kast T, Wolf B, Fraipont C, Aarsman ME, Kannenberg K, von Rechenberg M, Nguyen-Disteche M, den Blaauwen T, Holtje JV, Vollmer W. 2006. Interaction between two murein (peptidoglycan) synthases, PBP3 and PBP1B, in *Escherichia coli*. Mol Microbiol 61:675–90.

34. Spratt BG. 1975. Distinct penicillin binding proteins involved in the division, elongation, and shape of *Escherichia coli* K12. Proc Natl Acad Sci U S A 72:2999–3003.

35. Smith JD, Kumarasiri M, Zhang W, Hesek D, Lee M, Toth M, Vakulenko S, Fisher JF, Mobashery S, Chen Y. 2013. Structural analysis of the role of *Pseudomonas aeruginosa* penicillin-binding protein 5 in beta-lactam resistance. Antimicrob Agents Chemother 57:3137–46.

36. Castanheira S, Lopez-Escarpa D, Paradela A, Garcia-Del Portillo F. 2024. *In Vivo* Cross-Linking Sheds Light on the *Salmonella* Divisome in Which PBP3 and PBP3(SAL) Compete for Occupancy. Mol Microbiol 122:797–818.

37. Herianto S, Lee HM, Chen C, Ho JA, Tu HL. 2025. Functional reconstitution of bacterial ESCRT-III protein PspA identifies key regions in membrane binding and remodeling. Protein Sci 34:e70309.

38. Hudina E, Schott-Verdugo S, Junglas B, Kutzner M, Ritter I, Hellmann N, Schneider D, Gohlke H, Sachse C. 2025. The bacterial ESCRT-III PspA rods thin lipid tubules and increase membrane curvature through helix alpha0 interactions. Proc Natl Acad Sci U S A 122:e2506286122.

39. Masilamani R, Cian MB, Dalebroux ZD. 2018. *Salmonella* Tol-Pal Reduces Outer Membrane Glycerophospholipid Levels for Envelope Homeostasis and Survival during Bacteremia. Infect Immun 86.

40. Hegde RS, Keenan RJ. 2024. A unifying model for membrane protein biogenesis. Nat Struct Mol Biol 31:1009–1017.

41. Jauss B, Petriman NA, Drepper F, Franz L, Sachelaru I, Welte T, Steinberg R, Warscheid B, Koch HG. 2019. Noncompetitive binding of PpiD and YidC to the SecYEG translocon expands the global view on the SecYEG interactome in *Escherichia coli*. J Biol Chem 294:19167–19183.

42. van Bloois E, Dekker HL, Froderberg L, Houben EN, Urbanus ML, de Koster CG, de Gier JW, Luirink J. 2008. Detection of cross-links between FtsH, YidC, HflK/C suggests a linked role for these proteins in quality control upon insertion of bacterial inner membrane proteins. FEBS Lett 582:1419–24.

43. Karmakar R. 2021. State of the art of bacterial chemotaxis. J Basic Microbiol 61:366–379.

44. Neumann-Schaal M, Acuna JMB, Jahn D. 2026. Strict Aerobic Lifestyle and Anaerobic Survival of Bacteria: Inseparable Twins? Mol Microbiol doi:10.1111/mmi.70071.

45. Ciemniecki JA, Newman DK. 2023. NADH dehydrogenases are the predominant phenazine reductases in the electron transport chain of *Pseudomonas aeruginosa*. Mol Microbiol 119:560–573.

46. Udaondo Z, Schilder KA, Blesa ARM, Tena-Garitaonaindia M, Mangana JC, Daddaoua A. 2025. Transcriptional Regulatory Systems in *Pseudomonas*: A Comparative Analysis of Helix-Turn-Helix Domains and Two-Component Signal Transduction Networks. Int J Mol Sci 26.

47. Corona F, Reales-Calderon JA, Gil C, Martinez JL. 2018. The development of a new parameter for tracking post-transcriptional regulation allows the detailed map of the *Pseudomonas aeruginosa* Crc regulon. Sci Rep 8:16793.

48. Yin L, Gong X, Li Y, Qu J, Zhang Y, Jin Y, Jin S, Wu W. 2025. The Catabolite Repression Control Protein Crc Regulates the Type III Secretion System Through the Adenylate Cyclase CyaB in *Pseudomonas aeruginosa*. Microorganisms 13.

49. Trunk K, Benkert B, Quack N, Munch R, Scheer M, Garbe J, Jansch L, Trost M, Wehland J, Buer J, Jahn M, Schobert M, Jahn D. 2010. Anaerobic adaptation in *Pseudomonas aeruginosa*: definition of the Anr and Dnr regulons. Environ Microbiol 12:1719–33.

50. Tribelli PM, Lujan AM, Pardo A, Ibarra JG, Fernandez Do Porto D, Smania A, Lopez NI. 2019. Core regulon of the global anaerobic regulator Anr targets central metabolism functions in *Pseudomonas* species. Sci Rep 9:9065.

51. Horna G, Ruiz J. 2021. Type 3 secretion system of *Pseudomonas aeruginosa*. Microbiol Res 246:126719.

52. Muriel-Millan LF, Montelongo-Martinez LF, Gonzalez-Valdez A, Bedoya-Perez LP, Cocotl-Yanez M, Soberon-Chavez G. 2024. The alternative sigma factor RpoS regulates *Pseudomonas aeruginosa quorum sensing* response by repressing the *pqsABCDE* operon and activating *vfr*. Mol Microbiol 121:291– 303.

53. Poulsen BE, Yang R, Clatworthy AE, White T, Osmulski SJ, Li L, Penaranda C, Lander ES, Shoresh N, Hung DT. 2019. Defining the core essential genome of *Pseudomonas aeruginosa*. Proc Natl Acad Sci U S A 116:10072–10080.

54. Lee SA, Gallagher LA, Thongdee M, Staudinger BJ, Lippman S, Singh PK, Manoil C. 2015. General and condition-specific essential functions of *Pseudomonas aeruginosa*. Proc Natl Acad Sci U S A 112:5189–94.

55. Chevalier S, Bouffartigues E, Tortuel D, David A, Tahrioui A, Labbe C, Barreau M, Tareau AS, Louis M, Lesouhaitier O, Cornelis P. 2022. Cell Envelope Stress Response in *Pseudomonas aeruginosa*. Adv Exp Med Biol 1386:147–184.

56. Dudek CA, Jahn D. 2022. PRODORIC: state-of-the-art database of prokaryotic gene regulation. Nucleic Acids Res 50:D295–D302.

57. Wang B, Lin YC, Vasquez-Rifo A, Jo J, Price-Whelan A, McDonald ST, Brown LM, Sieben C, Dietrich LEP. 2021. *Pseudomonas aeruginosa* PA14 produces R-bodies, extendable protein polymers with roles in host colonization and virulence. Nat Commun 12:4613.

58. Grant CR, Wan J, Komeili A. 2018. Organelle Formation in Bacteria and Archaea. Annu Rev Cell Dev Biol 34:217–238.

59. Toyofuku M, Nomura N, Eberl L. 2019. Types and origins of bacterial membrane vesicles. Nat Rev Microbiol 17:13–24.

60. Gray AN, Egan AJ, Van’t Veer IL, Verheul J, Colavin A, Koumoutsi A, Biboy J, Altelaar AF, Damen MJ, Huang KC, Simorre JP, Breukink E, den Blaauwen T, Typas A, Gross CA, Vollmer W. 2015. Coordination of peptidoglycan synthesis and outer membrane constriction during *Escherichia coli* cell division. Elife 4.

61. Junglas B, Huber ST, Heidler T, Schlosser L, Mann D, Hennig R, Clarke M, Hellmann N, Schneider D, Sachse C. 2021. PspA adopts an ESCRT-III-like fold and remodels bacterial membranes. Cell 184:3674–3688 e18.

62. Liu J, Tassinari M, Souza DP, Naskar S, Noel JK, Bohuszewicz O, Buck M, Williams TA, Baum B, Low HH. 2021. Bacterial Vipp1 and PspA are members of the ancient ESCRT-III membrane-remodeling superfamily. Cell 184:3660–3673 e18.

63. Boes A, Olatunji S, Breukink E, Terrak M. 2019. Regulation of the Peptidoglycan Polymerase Activity of PBP1b by Antagonist Actions of the Core Divisome Proteins FtsBLQ and FtsN. mBio 10.

64. 64. Sposato D, Mercolino J, Torrini L, Sperandeo P, Lucidi M, Alegiani R, Varone I, Molesini G, Leoni L, Rampioni G, Visca P, Imperi F. 2024. Redundant essentiality of AsmA-like proteins in *Pseudomonas aeruginosa*. mSphere 9:e0067723.

65. Guest RL, Lee MJ, Wang W, Silhavy TJ. 2023. A periplasmic phospholipase that maintains outer membrane lipid asymmetry in *Pseudomonas aeruginosa*. Proc Natl Acad Sci U S A 120:e2302546120.

66. Schultz KM, Schneider JR, Fischer MA, Cina NP, Riegert MO, Frank DW, Klug CS. 2023. Binding and transport of LPS occurs through the coordinated combination of an array of sites across the entire *Escherichia coli* LPS transport protein LptA. Protein Sci 32:e4724.

67. Caveney NA, Egan AJF, Ayala I, Laguri C, Robb CS, Breukink E, Vollmer W, Strynadka NCJ, Simorre JP. 2020. Structure of the Peptidoglycan Synthase Activator LpoP in *Pseudomonas aeruginosa*. Structure 28:643–650 e5.

68. Williams HD, Zlosnik JE, Ryall B. 2007. Oxygen, cyanide and energy generation in the cystic fibrosis pathogen *Pseudomonas aeruginosa*. Adv Microb Physiol 52:1–71.

69. Jorgensen F, Bally M, Chapon-Herve V, Michel G, Lazdunski A, Williams P, Stewart G. 1999. RpoS-dependent stress tolerance in *Pseudomonas aeruginosa*. Microbiology (Reading) 145 ( Pt 4):835–844.

70. Yang N, Ding S, Chen F, Zhang X, Xia Y, Di H, Cao Q, Deng X, Wu M, Wong CC, Tian XX, Yang CG, Zhao J, Lan L. 2015. The Crc protein participates in down-regulation of the Lon gene to promote rhamnolipid production and *rhl quorum sensing* in *Pseudomonas aeruginosa*. Mol Microbiol 96:526–47.

71. Wood LF, Leech AJ, Ohman DE. 2006. Cell wall-inhibitory antibiotics activate the alginate biosynthesis operon in *Pseudomonas aeruginosa*: Roles of sigma (AlgT) and the AlgW and Prc proteases. Mol Microbiol 62:412–26.

72. Chevalier S, Bouffartigues E, Bazire A, Tahrioui A, Duchesne R, Tortuel D, Maillot O, Clamens T, Orange N, Feuilloley MGJ, Lesouhaitier O, Dufour A, Cornelis P. 2019. Extracytoplasmic function sigma factors in Pseudomonas aeruginosa. Biochim Biophys Acta Gene Regul Mech 1862:706–721.

73. Taoka A, Eguchi Y, Shimoshige R, Fukumori Y. 2023. Recent advances in studies on magnetosome-associated proteins composing the bacterial geomagnetic sensor organelle. Microbiol Immunol 67:228–238.

74. Martinez-Garcia E, de Lorenzo V. 2011. Engineering multiple genomic deletions in Gram-negative bacteria: analysis of the multi-resistant antibiotic profile of *Pseudomonas putida* KT2440. Environ Microbiol 13:2702–16.

75. Pankratz D, Gomez NO, Nielsen A, Mustafayeva A, Gur M, Arce-Rodriguez F, Nikel PI, Haussler S, Arce-Rodriguez A. 2023. An expanded CRISPR-Cas9-assisted recombineering toolkit for engineering genetically intractable *Pseudomonas aeruginosa* isolates. Nat Protoc 18:3253–3288.

76. Jeske A, Arce-Rodriguez A, Thoming JG, Tomasch J, Haussler S. 2022. Evolution of biofilm-adapted gene expression profiles in *lasR*-deficient clinical *Pseudomonas aeruginosa* isolates. NPJ Biofilms Microbiomes 8:6.

77. Blin K, Pedersen LE, Weber T, Lee SY. 2016. CRISPy-web: An online resource to design sgRNAs for CRISPR applications. Synth Syst Biotechnol 1:118–121.

78. Ayala-Garcia P, Moreno-de Castro N, Jimenez-Guerrero I, Musken M, Arce-Rodriguez A, Perez-Montano F, Borrero-de Acuna JM. 2024. Isolation, Quantification, and Visualization of Extracellular Membrane Vesicles in Rhizobia Under Free-Living Conditions. Methods Mol Biol 2751:219–228.

79. Karimova G, Pidoux J, Ullmann A, Ladant D. 1998. A bacterial two-hybrid system based on a reconstituted signal transduction pathway. Proc Natl Acad Sci U S A 95:5752–6.

80. 80. Miller J. 1992. A short course in bacterial genetics: a laboratory manual and handbood for Escherichia coli and related bacteria. A short course in bacterial genetics: a laboratory manual and handbood for escherichia coli and related bacteria:143–149.

81. Kordes A, Preusse M, Willger SD, Braubach P, Jonigk D, Haverich A, Warnecke G, Haussler S. 2019. Genetically diverse *Pseudomonas aeruginosa* populations display similar transcriptomic profiles in a cystic fibrosis explanted lung. Nat Commun 10:3397.

82. Langmead B, Salzberg SL. 2012. Fast gapped-read alignment with Bowtie 2. Nat Methods 9:357–9.

83. Liao Y, Smyth GK, Shi W. 2014. featureCounts: an efficient general purpose program for assigning sequence reads to genomic features. Bioinformatics 30:923–30.

84. Robinson MD, McCarthy DJ, Smyth GK. 2010. edgeR: a Bioconductor package for differential expression analysis of digital gene expression data. Bioinformatics 26:139–40.

85. Benjamini Y, Drai D, Elmer G, Kafkafi N, Golani I. 2001. Controlling the false discovery rate in behavior genetics research. Behav Brain Res 125:279–84.

86. Perez-Riverol Y, Bai J, Bandla C, Garcia-Seisdedos D, Hewapathirana S, Kamatchinathan S, Kundu DJ, Prakash A, Frericks-Zipper A, Eisenacher M, Walzer M, Wang S, Brazma A, Vizcaino JA. 2022. The PRIDE database resources in 2022: a hub for mass spectrometry-based proteomics evidences. Nucleic Acids Res 50:D543–D552.

87. Katoh K, Misawa K, Kuma K, Miyata T. 2002. MAFFT: a novel method for rapid multiple sequence alignment based on fast Fourier transform. Nucleic Acids Res 30:3059–66.

88. Capella-Gutierrez S, Silla-Martinez JM, Gabaldon T. 2009. trimAl: a tool for automated alignment trimming in large-scale phylogenetic analyses. Bioinformatics 25:1972–3.

89. Price MN, Dehal PS, Arkin AP. 2010. FastTree 2--approximately maximum-likelihood trees for large alignments. PLoS One 5:e9490.

90. Letunic I, Bork P. 2024. Interactive Tree of Life (iTOL) v6: recent updates to the phylogenetic tree display and annotation tool. Nucleic Acids Res 52:W78–W82.

