## Supplemental Material for "From Magnetosomes to Vesicles: LemA Proteins as Drivers of Bacterial Membrane Reorganization"

**Short running title:** Proteins of vesicle and organelle formation

José Manuel Borrero-de Acuña^1,2,3^, Svea Holland^2,4^, Rokas Juodeikis^5,6^, Alejandro Arce-Rodríguez^1,3^, Paula Ayala García^3^, Mustafa Safa Karagöz^7,8^, Joaquín Giner-Lamia^9^, Gabriella Molinari^10^, Gintare Deimantaviciute^5^, Viktoria Otto^1,2^, Carlos Azogue-Palma^3^, Ian R. Brown^5^, Katrin Müller^1,2^, Martina Jahn^1^, Kim Eileen Rennhack^1^, Susanne Häussler^11,12,13,14^, Manfred Rohde^10^, Lothar Jänsch^15^, Marco van Ham^15^, Mathias Müsken^10^, Evelyne Deery^5^, C. Mark Smales^6^, Meina Neumann-Schaal^2,4^, Martin J. Warren^5,6*^; and Dieter Jahn^1,2*^

^1^Institute of Microbiology, Technische Universität Braunschweig, 38106 Braunschweig, Germany

^2^Braunschweig Integrated Centre of Systems Biology (BRICS), Technische Universität

Braunschweig, 38106 Braunschweig, Germany

^3^Departamento de Microbiología, Facultad de Biología, Universidad de Sevilla, Sevilla, Spain

^4^Leibniz-Institut DSMZ-Deutsche Sammlung von Mikroorganismen und Zellkulturen, 38124 Braunschweig, Germany.

^5^School of Biosciences, University of Kent, Canterbury, CT2 7NJ, UK

^6^Quadram Institute Bioscience, Norwich Science Park, Norwich, NR4 7UQ, UK

^7^Biochemistry and Biophysics Center, National Heart, Lung, and Blood Institute, National Institutes of Health, Bethesda, Maryland, USA

^8^Department of Medical Biochemistry and Biophysics, Karolinska Institutet, Tomtebodavägen 23A, 17165 Solna, Stockholm, Sweden

^9^Instituto de Bioquímica Vegetal y Fotosíntesis, Universidad de Sevilla, Consejo Superior de Investigaciones Científicas, Américo Vespucio 49, Sevilla, 41092, Spain

^10^Central Facility for Microscopy, Helmholtz Centre for Infection Research (HZI), 38124 Braunschweig, Germany

^11^Institute for Molecular Bacteriology, TWINCORE GmbH, Center of Clinical and Experimental Infection Research, a joint venture of the Hannover Medical School and the Helmholtz Centre for Infection Research, Hannover, Germany

^12^Department of Molecular Bacteriology, Helmholtz Centre for Infection Research (HZI), 38124, Braunschweig, Germany

^13^Department of Clinical Microbiology, Copenhagen University Hospital – Rigshospitalet, 2100 Copenhagen, Denmark

^14^Cluster of Excellence RESIST (EXC 2155), Hannover Medical School, Hannover, 30625, Germany

^15^Cellular Proteome Research, Helmholtz Centre for Infection Research (HZI), 38124 Braunschweig, Germany

*Correspondence to: Martin J Warren or Dieter Jahn

**Extended introduction**

The presence of specialised membranous structures in bacteria has been known for over 60 years with the discovery of chromatophores in photosynthetic bacteria (1). Since then, a multitude of novel bacterial organelles have been described (2, 3) including intricate intracytoplasmic membrane invaginations found in methane and ammonia oxidizing bacteria as well as magnetosomes, the specialized membrane organelles capable of the biomineralization of iron (4, 5). These bacterial organelles are essential for the specialised biochemistry and habitat adaptation of their producers. Highly related to formation of intracellular membrane-encapsulated organelles is the production of extracellular membranous vesicles by many bacteria that are commonly referred as bacterial membrane vesicles (BMVs) (6, 7). Naturally, intracellular organelles and BMVs as the result of membrane shaping processes have major implications for our understanding of microbial virulence and bacterial community communication encompassing the inter-relationships among a large variety of species that represent the backbone of the ecosystem of the planet. Consequently, the ability to control and stimulate the production of BMVs and organelles also holds potential for many biotechnological applications especially in the area of vaccine production, drug delivery and complex recombinant production processes (8-10).

Extracellularly secreted BMVs are often classified as outer membrane vesicles (OMVs), outer-inner membrane vesicles (OIMVs) and cytoplasmic membrane vesicles (CMVs). Extracellular BMVs in bacteria are important for intra-species, inter-species and inter-kingdom cell signalling and communication (11, 12). OMVs contain a single membrane and are thought to result from outer membrane blebbing of Gram-negative bacteria, a process that allows for the inclusion of some periplasmic components (13-15). OIMVs consist of two membranes and have been suggested to arise from either blebbing or explosive cell lysis (16). Finally, CMVs contain a single membrane and have been shown to be produced by Gram-positive bacteria most likely using an unknown conserved blebbing mechanism (17).

Other bacteria form OMV-related tube-shaped structures. These various BMVs have not only been shown to be metabolically active but also to be involved in bacterial communication through the trafficking of molecules such as DNA, RNA and proteins (18-21). Furthermore, BMVs play a role in a variety of bacteria-host interactions, including modulation of immune cells, and have been shown to be trafficked into the host cell allowing for delivery of signal molecules (22-25). Membranous structure formation is likely the property of most bacteria, however, the exact underlying molecular mechanisms for the formation of membrane organelles and BMVs remain unclear. Thus, almost no specialised proteins have so far been identified in the process of bacterial BMV formation although several genetic mutations that influence BMV formation have been determined (26). The only exception is cyanobacterial phage shock protein PspA which was shown via cryoelectron microscopy to adopt a canonical ESCRT-III-fold and to generate positive membrane curvature to remodel BMVs in serving as lipid transfer platform and linking previously separated lipid structures (27-29). The endosomal sorting complexes required for transport (ESCRT) were shown to be responsible for membrane remodelling in many eukaryotic cellular processes, such as cytokinetic abscission, multivesicular body biogenesis and viral budding. The most abundant ESCRT subunit ESCRT-III assembles into flat spirals and is essential to initiate membrane invagination (30).

Here, we report the discovery of the LemA family of proteins involved in BMV formation in bacteria. We show the effect of the heterologous overproduction of 13 different bacterial LemA proteins in *E. coli* using different microscopic techniques and observe that this results in large-scale membrane restructuring and vesiculation. Homologous overproduction of PaLemA1 and PaLemA2 in *P.* *aeruginosa* results in the appearance of either internal cytoplasmic or external membrane vesicles. Additionally, *P. aeruginosa* mutant strains lacking both Pa*lemA1* and Pa*lemA2* are largely defected in extracellular BMV secretion. Interactomics studies using PaLemA1 and PaLemA2 identified 256 protein interaction mostly involved in cell division and related processes of lipopolysaccharide (LPS), cell wall, and membrane formation with corresponding gene regulatory and stress response processes including the known vesicle formation protein PspA and the protein CpoB. CpoB was reported before as a periplasmic factor that coordinates peptidoglycan remodelling during cell division with distinct functional differences between *E. coli* and *P. aeruginosa* (31-33). The bacterial predator *Bdellovibrio bacteriovorus* deposits during invasion a vesicular compartment filled with CpoB into the prey´s periplasm to position chimeric fibres required for the process (34).

The formation of a large protein complex coordinating BMV formation controlled by PaLemA proteins, CpoB and PspA was elucidated. Finally, it was concluded that the complex-interacting sigma factor AlgU regulates Pa*lemA2* transcription.

**Results**

1. **Initial discovery of a protein of the vesicle formation machinery: MamQ initiates inner membrane curvature during magnetosome formation**

The MamQ protein was shown to be essential for the formation of the membranous structures of the magnetosome. We showed that removal of the predicted C-terminal and the transmembrane region of MamQ results in the aggregation of the protein and the loss of the membranous phenotype (Fig. 1b). Extended Data Fig. 1 shows a model for MamQ in the cytoplasmic membrane based on a mutational analysis of MamQ where mutations in the periplasmic region (residues 68-272) were detrimental to magnetosome formation (35).

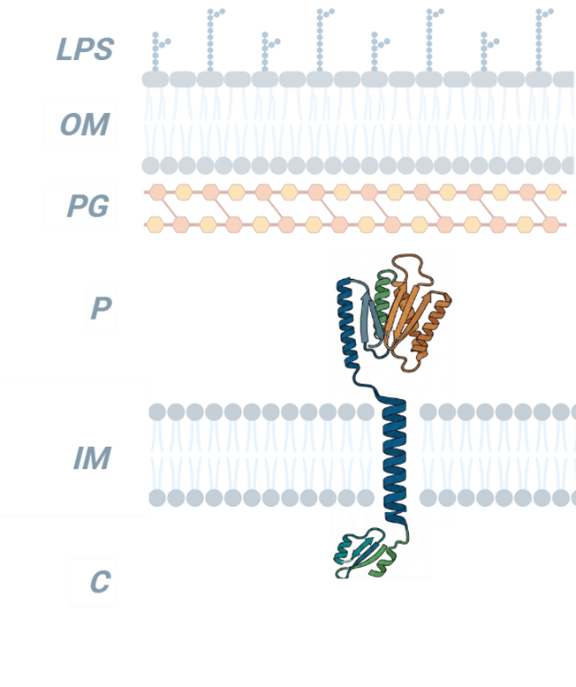

**Extended Data Fig. 1. Modelled structure of MamQ from *Magnetospirillum* *gryphiswaldense*.** The MamQ protein structure was predicted using AlphaFold3, and structural models were visualized in PyMOL (version 2.5, Schrödinger LLC). Conceptual membrane topology schematics and publication-ready illustrations were generated in BioRender based on the predicted structural and topology features. The model contains an internal transmembrane helix (residues 47–67), with the N-terminal domain (residues 1–46) oriented toward the cytoplasm and the C-terminal domain (residues 68–272) positioned toward the periplasmic/magnetosome lumenal side. The orientation was inferred from the charge distribution flanking the transmembrane helix in accordance with the positive-inside rule (36). The distribution of positively charged residues in bacterial inner membrane proteins correlates with the trans-membrane topology. Eukaryotic-trained model predictions (e.g., TargetP lysosomal/vacuolar signals) are disregarded as artifacts in the bacterial context. Abbreviations: LPS = Lipopolysaccharide; OM = outer membrane; PG = Peptidoglycan; P = periplasm; IM = inner membrane; C = cytoplasm. The PyMOL images were assembled in BioRender.

No membrane invaginations were observed during expression of the mutated *mamQ* genes in *E. coli*. These results indicate that MamQ has the ability to initiate membrane invagination, most likely through the induction of membrane curvature.

2. **MamQ is part of LemA family of bacterial protein**

MamQ is a member of the LemA protein family (PF04011). LemA was originally described as an antigenic determinant in *Listeria* *monocytogenes*. Analysis of 2,565 LemA homologues deduced from 12,221 prokaryotic reference genomes (Extended Data Tables 1 - 3) revealed a deep evolutionary split into LemA1 (n = 1,470) and LemA2 clades (n = 1,095) (Extended Data Fig. 2a).

Members of the LemA family are widely distributed across 36 bacterial phyla, with the highest representation in Proteobacteria (46%), Firmicutes (20%), Bacteroidota (16%) and Actinobacteriota (9%) (Extended Data Fig. 2b). LemA proteins are present in organisms occupying diverse ecological niches, including host-associated, soil and aquatic environments. The broad phylogenetic distribution and frequent duplication of LemA proteins suggest that they confer a conserved and potentially adaptive function, consistent with a fundamental role in bacterial membrane remodelling across diverse lifestyles.

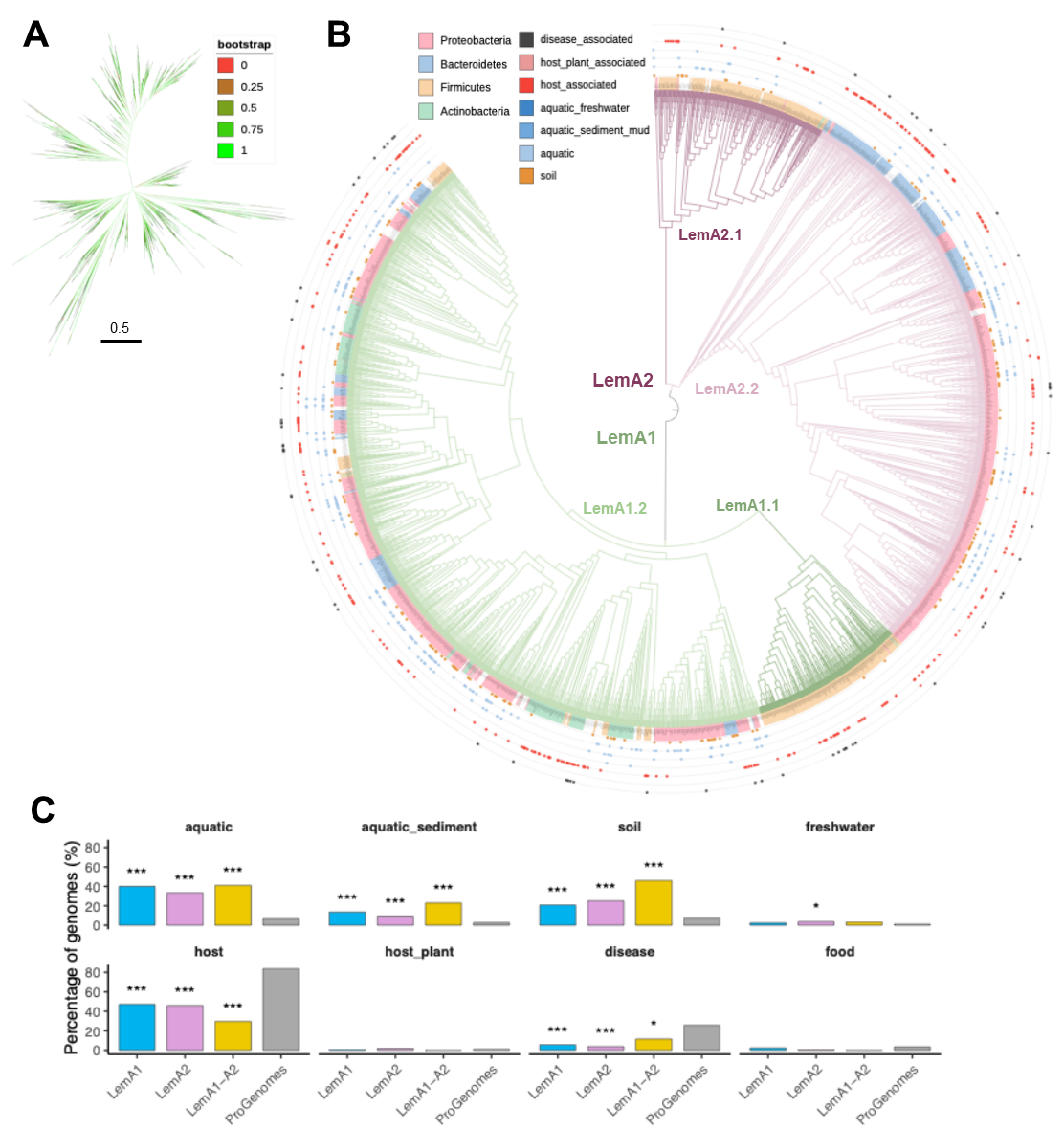

**Extended Data Fig. 2.** **Phylogenetic analysis and ecological distribution of LemA family. A)** Maximum-likelihood phylogeny inferred with FastTree (see Methods) from 2,565 LemA homologous sequences identified in the ProGenomes2 dataset. Bootstrap support is represented as a colour gradient (see legend). **B)** Circular representation of the phylogeny in a, highlighting the major phyla. Outer rings denote habitat annotations for each genome. **C)** Habitat distribution of genomes encoding LemA1, LemA2, or both homologues (LemA1-A2) compared to proGenomes2. Percentages represent the proportion of strains assigned to each ecological category relative to the total number of strains within each group for which habitat information was available (n = 56,723 annotated genomes). Statistical significance was determined using two-tailed Fisher’s exact tests, comparing the frequency of each habitat in LemA groups against the proGenomes2 background. *p*-values were adjusted for multiple testing using the Benjamini-Hochberg False Discovery Rate (*FDR*) method. Asterisks indicate significance levels: **FDR* < 0.05, ***FDR < 0.01*, ****FDR < 0.001.*

3. **LemA proteins from various bacteria restructure *E. coli* membranes**

To functionally investigate if the broader LemA protein family similarly induces membrane restructuring, we cloned and expressed thirteen different *lemA* genes in *E. coli* (Extended Data Fig. 3a). The encoded proteins (Extended Data Fig. 3b), varied in their predicted membrane topology and represented both Gram-negative (*Bacteroides*, *Brucella*, *Crinalium*, *Pseudomonas*, *Spirochaeta*, *Thioflavicoccus*) and	Gram-positive (*Anaerobacillus*, *Bacillus*, *Exiguobacterium*, *Listeria*) bacteria (Extended Data Fig. 3b). Recombinant LemA production in *E. coli* was observed for ten of these genes via SDS PAGE (Extended Data Fig. 3c). The three proteins (CreLemA2, BatLemA1A, StLemA2) that were not well-overproduced in *E. coli* still caused delayed growth and cell lysis in the host (Extended Data Fig. 3c, lanes 2, 7, & 12).

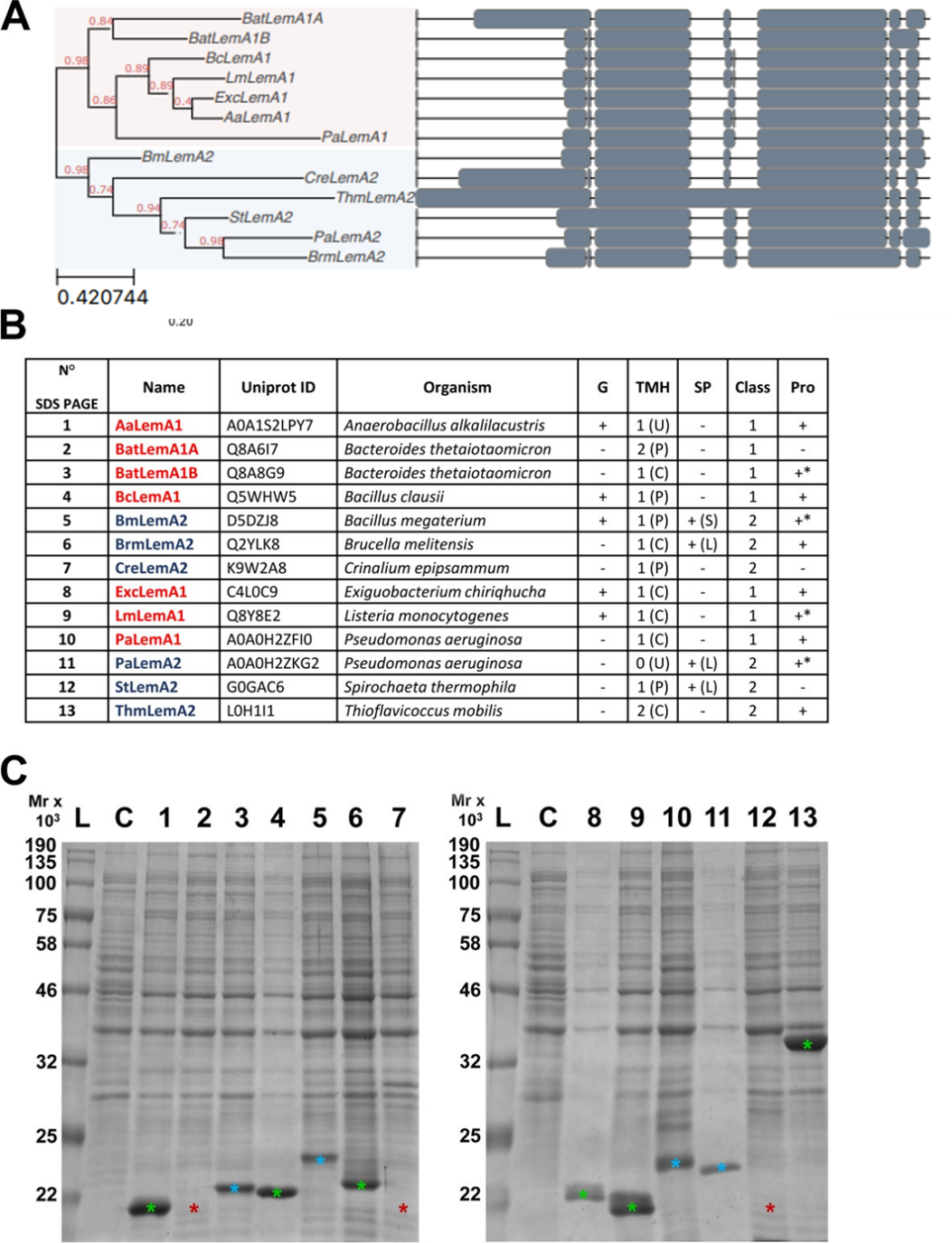

**Extended Data Fig. 3. Phylogenetic analysis, characteristics and recombinant production of the 13 LemA proteins investigated in this study.** The phylogenetic analysis (**A**) reveals the two major clades, LemA1 and LemA2. The tree was inferred from an alignment of 13 LemA protein sequences generated with MAFFT and reconstructed using the maximum-likelihood method implemented in FastTree. Branch support values are indicated at the main nodes, and branch lengths represent substitutions per site. The graphical representation of the tree was generated using the ETE Toolkit. Conserved regions across the aligned sequences are shown. The table (**B**) correlates the SDS gel lane number with the abbreviation of the various LemA proteins, their Uniprot ID and the organism from which they are derived. Further abbreviations are: G – Gram-positive [+] or Gram-negative [-]; TMHs – predicted transmembrane helices with the soluble part facing the periplasm [P], cytoplasm [C]; or it is unknown [U]; SP – signal peptide (for Sec pathway [S]/ with Lipoprotein signature [L]); Class – 1 or 2, as described in the main text Pro – obvious protein production [+] or absence [-] in *E. coli* BL21(DE3) Star as observed via SDS-PAGE. The Coomassie Brilliant Blue stained SDS PAGE (**C**) shows the recombinant production of each protein listed in the table above in *E. coli*. A green asterisk on a stained protein indicates the presence of a protein band that is consistent with the predicted molecular mass. A blue asterisk indicates an electrophoretic behaviour slightly different from the expected molecular mass. Such differences in electrophoretic mobility are often related to the hydrophobic character of some of the proteins. A red asterisk is shown when no visible band for the produced LemA protein was observed. Abbreviations: Mr x 10^3^ = relative molecular mass x 10^3^; L = ladder of marker proteins with defined molecular mass.

Thin sections of the recombinant *E. coli* that more efficiently produced the various LemA proteins were then analysed using transmission electron microscopy (TEM). Figure 2 presents the morphotypes of strains that achieved high levels of LemA expression, including variants displaying minor alterations in electrophoretic mobility, while maintaining normal host growth and cellular integrity. In contrast, the strains shown in Extended Data Fig. 4 exhibited either aberrant electrophoretic migration patterns or no protein production yields or evidence of host toxicity. Although intracellular membrane proliferation was observed across all samples, the morphotypes in Extended Data Fig. 4 were characterized by increased intracellular protein aggregation and signs of cellular disintegration

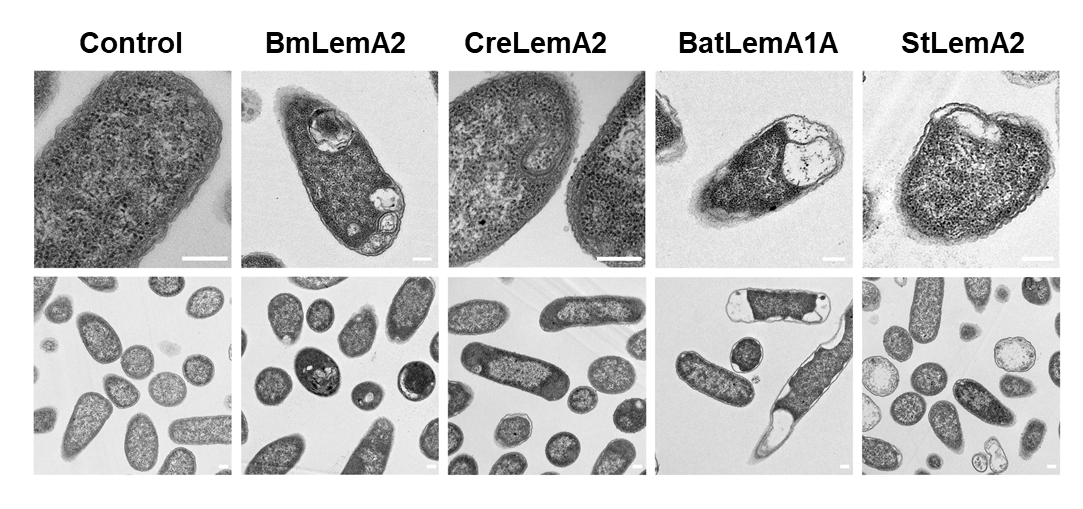

**Extended Data Fig. 4. Transmission electron micrographs of recombinant *E. coli* cells producing 4 different bacterial LemA proteins with reduced production yields or aberrant migration pattern.** The production of these proteins reveals membrane restructuring, cell disintegration and protein aggregation. Recombinant *E. coli* BL21 (DE3) Star cells producing *Bacillus* *megaterium* LemA2 (BmLemA2), *Crinalium* *epipsammum* LemA2 (CreLemA2), *Bacteroides* *thetaiotaomicron* LemA1A (BatLemA1A) and *Spirochaeta* *thermophila* LemA2 (StLemA2) were analysed by TEM and photographed after 16 hours growth. Observed phenotypes revealed membrane restructuring, cell disintegration and protein aggregation. The shown scale bars represent 200 nm.

The major phenotypes observed were the formation of (1) CMVs (AaLemA1, BcLemA1, LmLemA1, BatLemA1B), (2) intracellular tubular membrane structures (ExcLemA1, ThmLemA2) and (3) extrusion of the inner membrane into the periplasmic space (BrmLemA2). Occasionally LemA1 overproduction in *E. coli* led to cell lysis (Extended Data Fig. 5), perhaps reflecting a general stress during protein overproduction. In some instances, it appeared that extensive restructuring of the cytoplasmic membrane led to cell death, but leaving cells with internal CMVs intact which are observed as stable entities within the debris of lysed cells (Extended Data Fig. 5). Unlysed cells showed in certain cases aggregated proteins comprising the grey matter surrounding the CMVs (Extended Data Fig. 5).

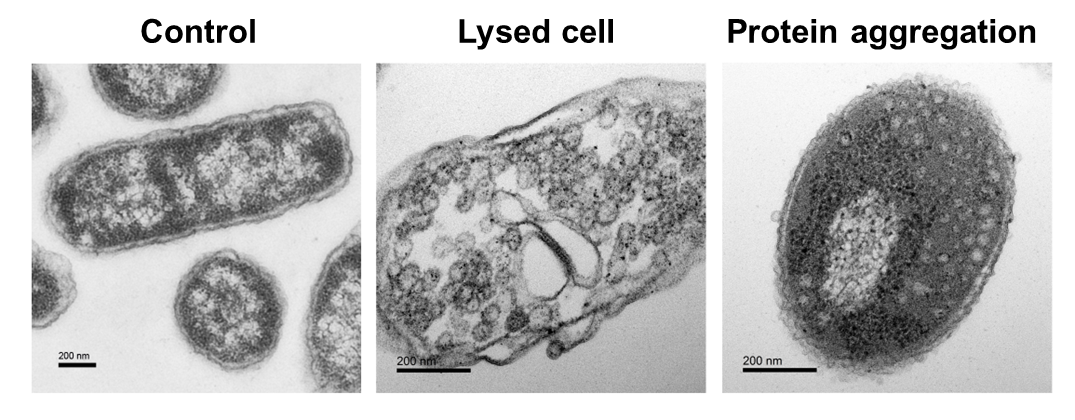

**Extended Data Fig. 5. Electron micrographs of additional phenotypes observed for *E. coli* BL21(DE3) cells producing *P. aeruginosa* LemA1 (PaLemA1).** Electron micrographs of *E. coli* BL21 (DE3) Star cells producing PaLemA1 are shown. All images were taken after 16 hours of *E. coli* growth. The control strain was *E. coli* harbouring the empty pET3a plasmid (Control). Image in the middle (Lysed cell) shows the conservation of CMVs post cell lysis. The image on the right (protein aggregation) shows aggregation observed around the membrane and the CMV (light grey areas). All scale bars are 500 nm.

In order to understand the physiological role of LemA proteins we further investigated the two proteins PaLemA1 and PaLemA2 from *P.* *aeruginosa*, a well-studied opportunistic bacterial pathogen where advanced molecular genetics methods are available. *P. aeruginosa* PA14 contains genes for both LemA1 (Pa*lemA1*) and LemA2 (Pa*lemA2*), which are located in two distinct operons. We observed that production of both proteins in *E. coli* did not result in detectable cellular toxicity, despite their electrophoretic migration differing marginally from that expected based on their predicted molecular weights. The membrane structures induced by the presence of recombinantly produced PaLemA1 in *E. coli* yielded circular CMVs rather than lamellar arrangements, as no tubular assemblies were observed (Extended Data Fig. 6 & 7a).

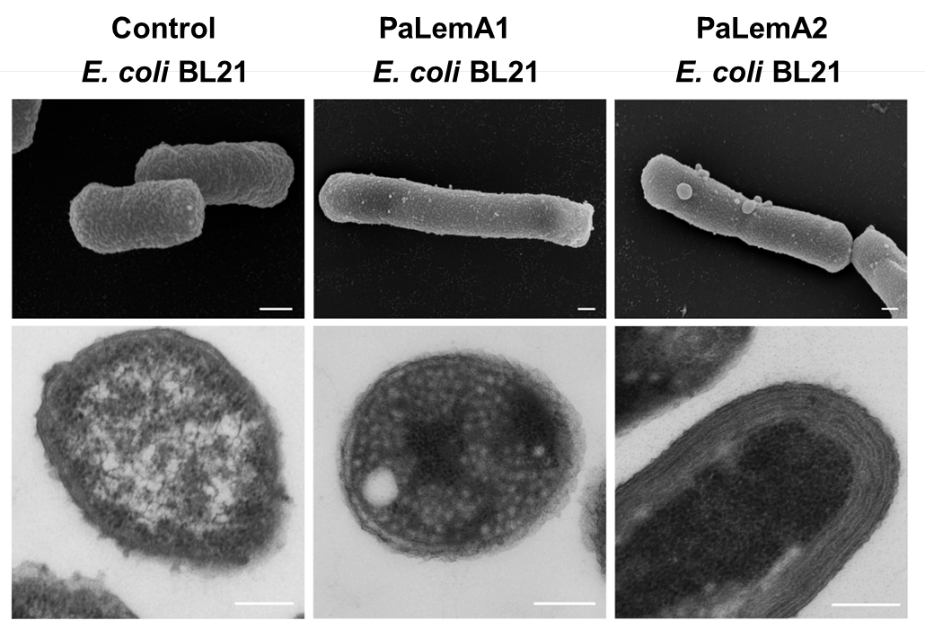

**Extended Data Fig. 6. Transmission and scanning electron micrographs of recombinant *E. coli* cells producing PaLemA1 and PaLemA2.**  The Pa*lemA*1 and Pa*lemA*2 genes were individually overexpressed in *E. coli* BL21 (DE3) Star. FESEM (upper row) and TEM (lower row) micrographs were taken after 16 hours of *E. coli* growth. Overproduction of PaLemA1 in *E. coli* led to the formation of multiple intracellular globular structures (CMVs) as visualized by TEM. Almost no OMVs were visible on the surface of these bacteria. In contrast, FESEM analysis revealed membrane blebbing for the *E. coli* strains overproducing PaLemA2, resulting in the formation of large sized OMVs on the surface. When producing PaLemA2 *E. coli* showed stacks of membranous structures below the surface. Scale bars are 200 nm.

The variety of intracellular structures observed in *E. coli* with the PaLemA2 overproducing strain were quite distinct to the PaLemA1 overproducing strain (Extended Data Fig. 6 & 7c). Stacked membrane structures close to the inner membrane were observed (Extended Data Fig. 6 & 7c). Furthermore, field emission scanning electron microscopy (FESEM) revealed large vesicular structures on the surface of the cells producing the lipoprotein, suggesting a role of PaLemA2 in OMV formation (Extended Data Fig. 6).

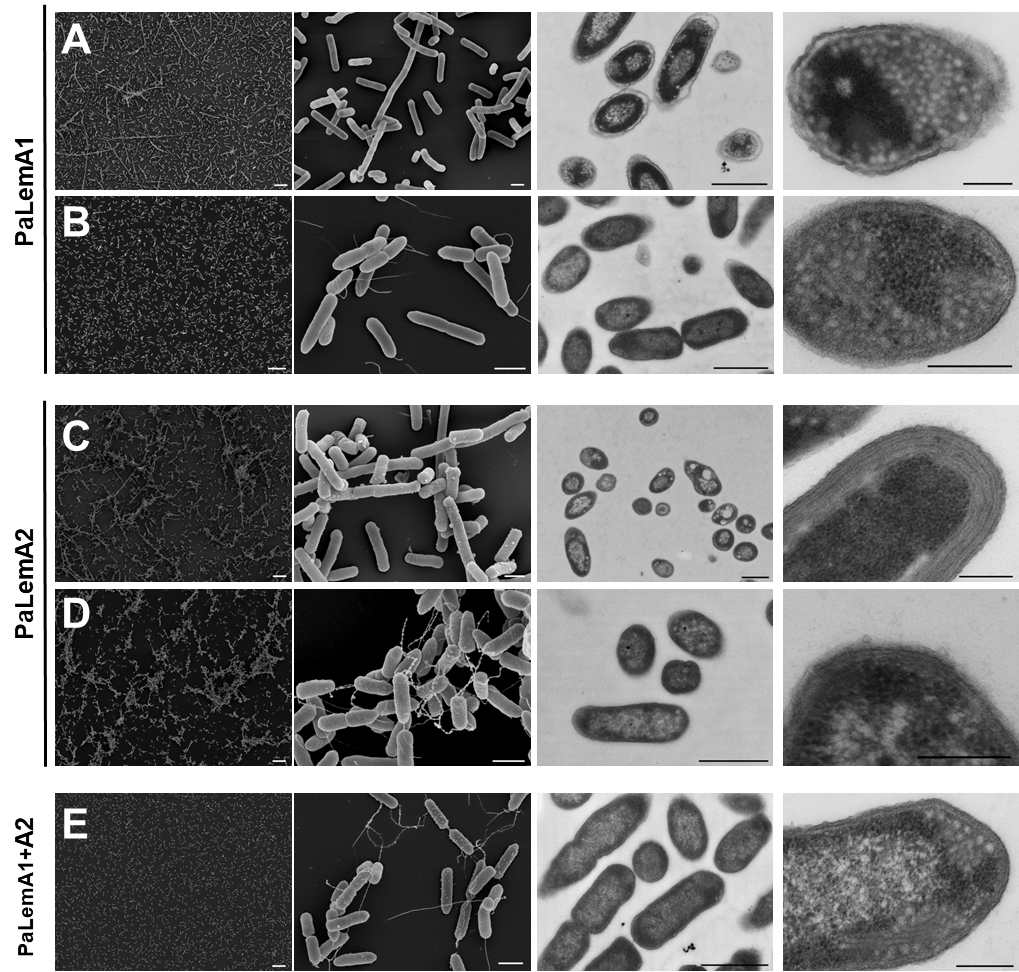

**Extended Data Fig. 7. Additional scanning and transmission electron micrographs of *E. coli* and *P. aeruginosa* overproducing PaLemA1 and PaLemA2.** All images were taken after 5 hours growth. The panels show a combination of low and high magnification scanning and transmission micrographs. Panel A shows *E. coli* BL21 (DE3) Star producing PaLemA1, Panel B shows *P. aeruginosa* ΔPa*lemA1* producing PaLemA1, Panel C shows *E. coli* producing PaLemA2, Panel D shows *P. aeruginosa* ΔPa*lemA2* producing PaLemA2, Panel E shows *P. aeruginosa* ΔPa*lemA1+A2* producing PaLemA1+A2. FESEM analyses reveal that overproduction of PaLemA1 in *E. coli* (**A**) results in some extended cells. However, no increase in vesiculation was observed on the surface. TEM micrographs show multiple intracellular globular structures (CMVs). When the production of PaLemA1 in *P. aeruginosa* (**B**) was analysed by FESEM almost no external OMVs were observed. TEM analyses reveal the presence of multiple intracellular tubular and membranous structures. Overproduction of PaLemA2 in either *E. coli* or *P. aeruginosa* yielded a hypervesiculation phenotype observable by FESEM (**C, D**) characterized by multiple OMVs on the surface and attached to the flagella. Interestingly, *E. coli OMVs* on the surface were of a much larger size. Remarkably, *E. coli* and *P. aeruginosa* overproducing PaLemA2 appear to agglutinate in this early stage of growth. This phenotype is usually observable only in the stationary phase. TEM analysis of *E. coli* strain producing PaLemA2 shows the presence of multiple stacked membranes and evidence of uncontrolled membrane proliferation. Moreover, more frequent events of OMVs being released from the surface were captured. The *P. aeruginosa* strain overexpressing Pa*lemA1* and Pa*lemA2* in tandem (**E**) shows a mixed phenotype of the PaLemA1 and PaLemA2 producing *P. aeruginosa* strains. From left to right scale bars are 10, 1, 1, and 0.2 µm, respectively.

Immunogold labelling was carried out to confirm the presence of the recombinant PaLemA proteins in the membranous structures in *E. coli*. The gold labelling was found co-localised with the membranous structures, indicating that the PaLemA proteins are associated with the restructured membranes (Extended Data Fig. 8). Moreover, PaLemA1 appeared to localise to the inner membrane and to the intracellular structures formed in *E. coli*, while PaLemA2 appeared to be present both on the inner and outer membrane (Extended Data Fig. 8). This observation is consistent with the predicted localization of these proteins. Overall, these results showed that in *E. coli* PaLemA1 associated with the inner membrane and was most likely involved in membrane curvature that led to the formation of CMVs, while PaLemA2 associated with the outer membrane and sustained the production of OMVs.

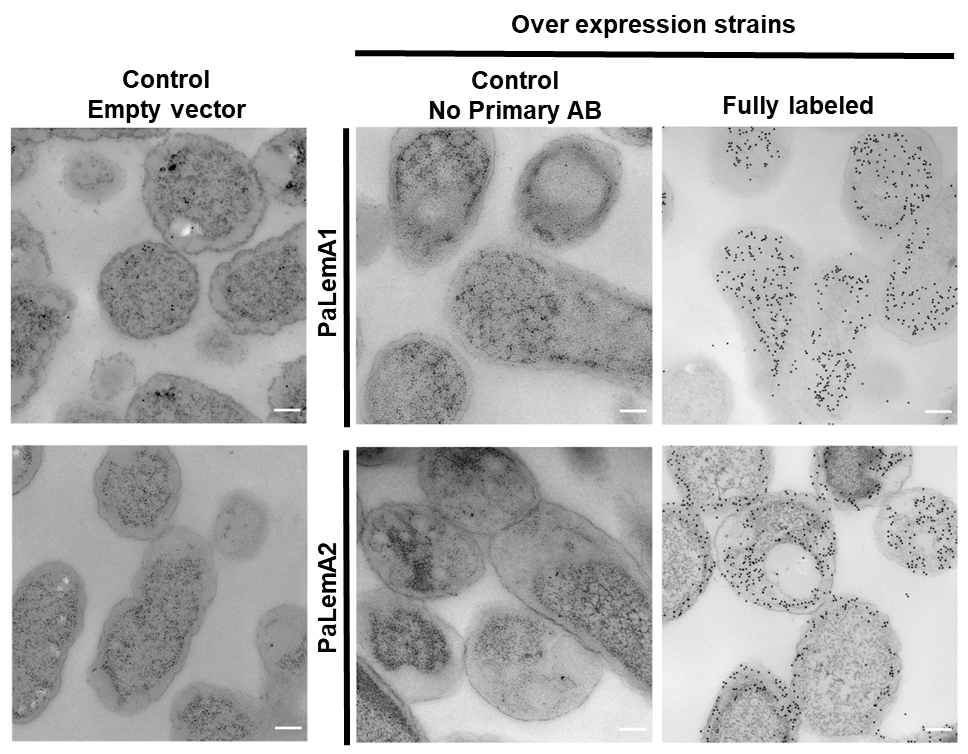

**Extended Data Fig. 8.** **Cellular localization of the PaLemA1 and PaLemA2 proteins recombinantly produced in *E. coli* using immuno-gold labelled proteins and transmission electron microscopy.** All *E. coli* BL21 (DE3) Star strains were grown for 16 hours, fixed and treated with either anti-PaLemA1 and PaLemA2 antibodies as indicated. Transmission electron micrographs of thin sections of *E. coli* cells containing either the control vector (empty vector) or Pa*lemA*1 (upper panel) or Pa*lemA*2 (lower panel) encoding plasmids were taken. Controls without the primary antibody (No Primary AB) are shown. Immuno-labelled PaLemA1 was localized in the cytoplasm to the membranous structures and to the cytoplasmic membrane, while PaLemA2 was almost exclusively found at the cell surface. PaLemA2 localization was monitored for most of the cells in the overall bacterial population. All scale bars are 1 µm.

4. ***P. aeruginosa* LemA1 (PaLemA1) and LemA2 (PaLemA2) differentially modulate membrane architecture and vesicle formation**

The membrane structures induced by the presence of recombinantly produced PaLemA1 in *E. coli* yielded circular CMVs rather than lamellar arrangements, as no tubular assemblies were observed (Extended Data Figs. 6 & 7a). The variety of intracellular structures observed in *E. coli* with the PaLemA2 overproducing strain were quite distinct to the PaLemA1 overproducing strain (Extended Data Figs. 4, 6 & 7a & 7c). Stacked membrane structures close to the inner membrane were observed (Extended Data Figs. 6 & 7c). Furthermore, field emission scanning electron microscopy (FESEM) revealed large vesicular structures on the surface of the cells producing the lipoprotein, suggesting a role of LemA2 in OMV formation (Extended Data Fig. 6). Immunogold labelling was carried out to confirm the presence of the recombinant PaLemA proteins in the membranous structures in *E. coli*. The gold labelling was found co-localized with the membranous structures, indicating that the PaLemA proteins are associated with the restructured membranes (Extended Data Fig. 8).

In order to characterize the PaLemA proteins in their native host, the corresponding genes were deleted in *P. aeruginosa* to generate two single and one double knock out strains (ΔPa*lemA*1, ΔPa*lemA*2, ΔPa*lemA*1+A2). Gene overexpression strains were then generated in these null backgrounds (resulting in ΔPalemA1+pPalemA1, ΔPa*lemA*2 + pPa*lemA*2, ΔPa*lemA*1+*A*2 + pPa*lemA*1+A2). No growth defects were observed for any of these strains. Significant PaLemA1 amounts were detected via Western blotting in the whole cellular extract of PaLemA1 overproduction strains (ΔPa*lemA*1 + pPa*lemA*1, ΔPa*lemA*1+A2 +pa*lemA*1+A2) (Extended Data Fig. 9). Analogously, PaLemA2 was detected in the whole cellular extract of PaLemA2 overproduction strains (ΔPa*lemA*2 + pPa*lemA*2, ΔPa*lemA*1+A2 + pPa*lemA*1+A2) (Extended Data Fig. 9). However, in isolated OMV/OIMV solely PaLemA2 was found in the corresponding PaLemA2 overproducing strains (Extended Data Fig. 9).

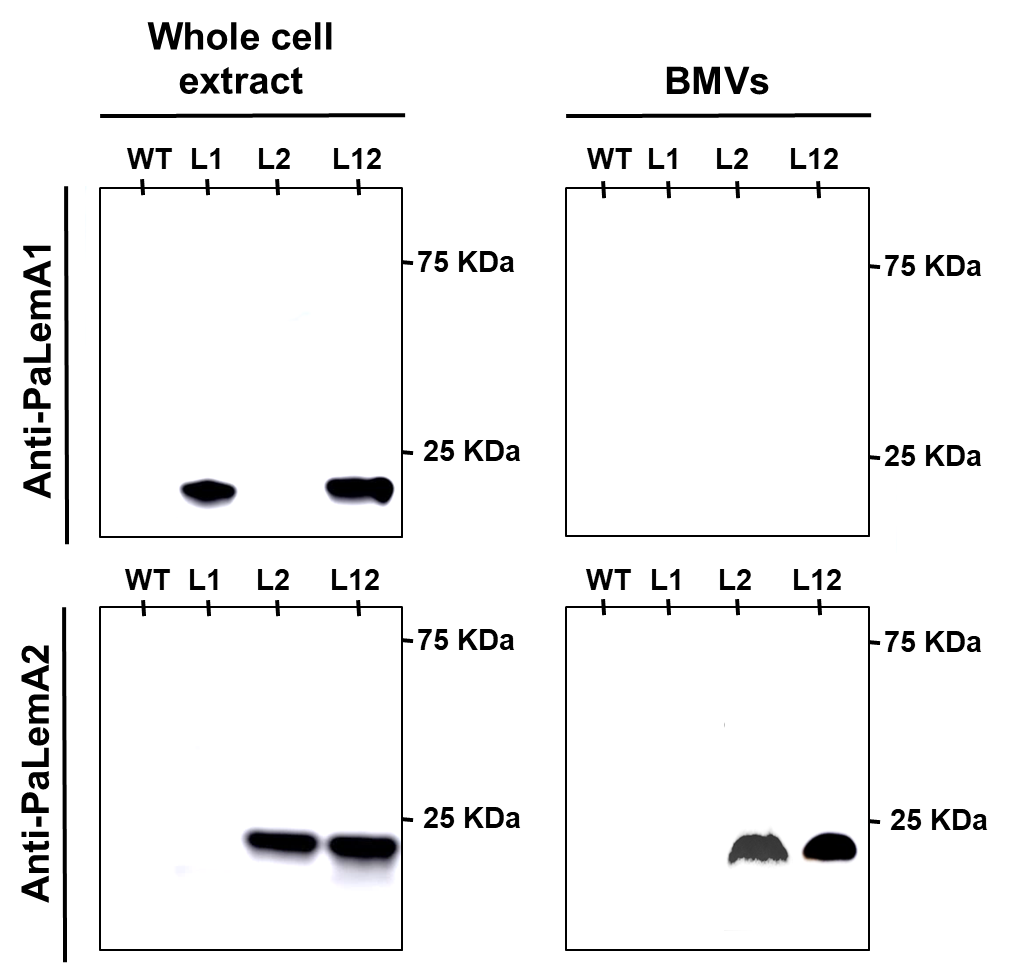

**Extended Fig. 9. The cellular and extracellular localization of the PaLemA1 and PaLemA2 proteins in *P. aeruginosa*.** The figure shows a western blot analysis using rabbit-raised anti-PaLemA1 and anti-PaLemA2 antibodies. The *P. aeruginosa* proteins were immunochemically detected in whole cellular extracts prepared from wildtype (WT), ΔPa*lemA1*+pPa*lemA1* (L1), ΔPa*lemA2*+pPa*lemA2* (L2), and ΔPa*lemA1+A2*+pPa*lemA1+A2* (L12). Additionally, isolated BMVs of bacterial cells were analysed in a similar fashion in order to detect protein in any secreted BMVs. The relative molecular mass x 10^3^ of a marker protein is given.

Thus, next we analysed the function of PaLemA1 using transmission (TEM) and scanning electron microscopy (FESEM) of the various *P. aeruginosa* overexpressing and knockout strains (Fig. 3, Extended Data Fig. 7, 10 & 11).

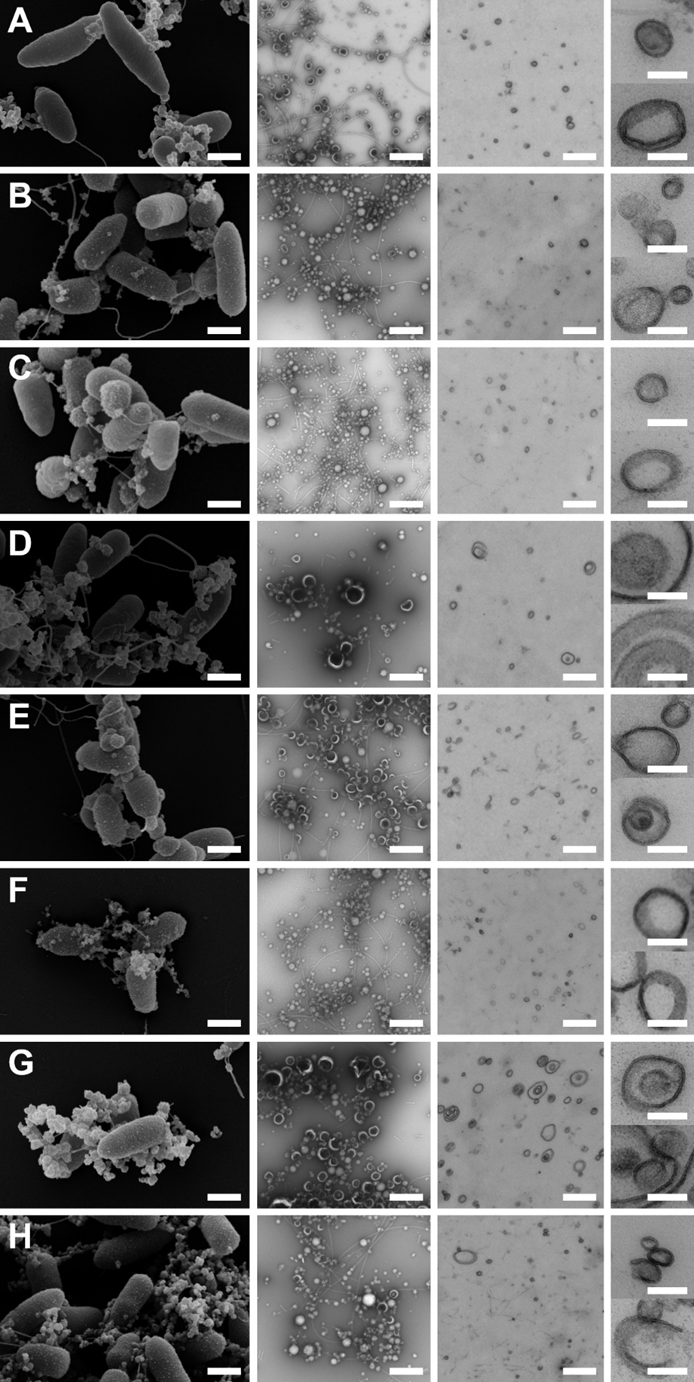

**Extended Data Fig. 10. The overall bacterial morphology of the various recombinant *P. aeruginosa* LemA strains was analysed by SEM.** Additionally, shape, size and the presence of either a single or double bilayer were determined for all strains through the use of negative staining and TEM. **A)** *P. aeruginosa* wildtype, **B)** *P. aeruginosa* wildtype + empty vector pSEVA634, **C)** *P. aeruginosa* ΔPa*lemA1* + pSEVA634, **D)** *P. aeruginosa* ΔPa*lemA1* + pPa*lemA1*, **E)** *P. aeruginosa* ΔPa*lemA2* + pSEVA634, **F)** *P. aeruginosa* ΔPa*lemA2* + pPa*lemA2*, **G)** *P. aeruginosa* ΔPa*lemA1+A2* + pSEVA634 and **H)** *P. aeruginosa* ΔPa*lemA1+A2* + pPa*lemA1+A2*. A mixture of OIMVs and OMVs was observed for most of the strains by negative staining and TEM. Larger secreted BMVs, often showing a dark interior in the negative staining samples, seem to correlate with the fraction of OIMVs. They can be found more often in the Pa*lemA2*-deficient strain compared to the Pa*lemA1*. In contrast, the Pa*lemA2*-overproducing strains showed a tendency to form almost exclusively small-size OMVs. BMVs carrying multiple membrane structures were observed for both PaLemA1 overproduction strains (ΔPa*lemA1+ p*Pa*lemA1*, ΔPa*lemA1+A2 +* pPa*lemA1+A2*) and most frequent, in the double mutant strain ΔPa*lemA1+A2*, indicating a disturbance in the BMV formation process. Scale bars are 500 nm for the first three image sets and 100 nm for two magnified single BMV sections.

No significant phenotype was observed for the knockout strains (Extended Data Fig. 10c, e, g & 11c - e). Overproduction of PaLemA1 in *P. aeruginosa* resulted in the formation of intracellular tubular and vesicular membrane structures as observed by TEM (Fig. 3 & Extended Data Fig. 7b). Vesicle-like structures were only rarely observed by FESEM on the cell surface of the wildtype and the ΔPa*lemA*1 mutant (Fig. 3 & Extended Data Fig. 10) and consequently wildtype levels of BMVs were seen for both strains (Fig 3). No significant phenotype could be observed by TEM in cell sections of PaLemA2-overproducing cells (Fig. 3). Instead, increased vesiculation of the outer membrane and cell agglutination was detected using FESEM (Fig. 3, Extended Data Fig. 7 d & 10f). A snapshot of the vesiculation process is shown in more detail in Extended Data Fig. 12. An agglutination effect was observed for the PaLemA2-overproducing strains after 5 hours of growth (3 hours induction of PaLemA2 production) and is characterized by bacterial communities with intertwined flagella interacting with a significant number of small sized vesicles (Extended Data Fig. 10f). A particular hallmark of the vesicles from the ΔPa*lemA*2 deletion mutant was their very large size and dark content, as observed by negative staining and TEM of the vesicles and FESEM of the cells (Fig. S11e). Many, but not all of the BMVs, revealed a double membrane bilayer. These phenomena increased significantly in aged cultures. Moreover, FESEM analyses revealed that deletion of Pa*lemA*2 significantly reduced the levels of observable BMVs. Overexpression of both Pa*lemA* genes in the double knockout strain (ΔPa*lemA*1+A2 + Pa*lemA*1+A2) led to a combined phenotype with abundant globular and tubular structures to the inside of the cell and bubble-like invaginations to the outside (Fig. 3, and Extended Data Fig. 7e). Interestingly, for both the PaLemA1 overproduction strain (ΔPa*lemA*1+Pa*lemA*1) and the double mutant strain (ΔPa*lemA*1+A2 + Pa*lemA*1+A2) very unusual BMVs carrying multiple membrane structures integrated into each other were observed after 16 hours of growth, indicating that the vesicle formation process had been disrupted (Extended Data Fig. 10d, g & h).

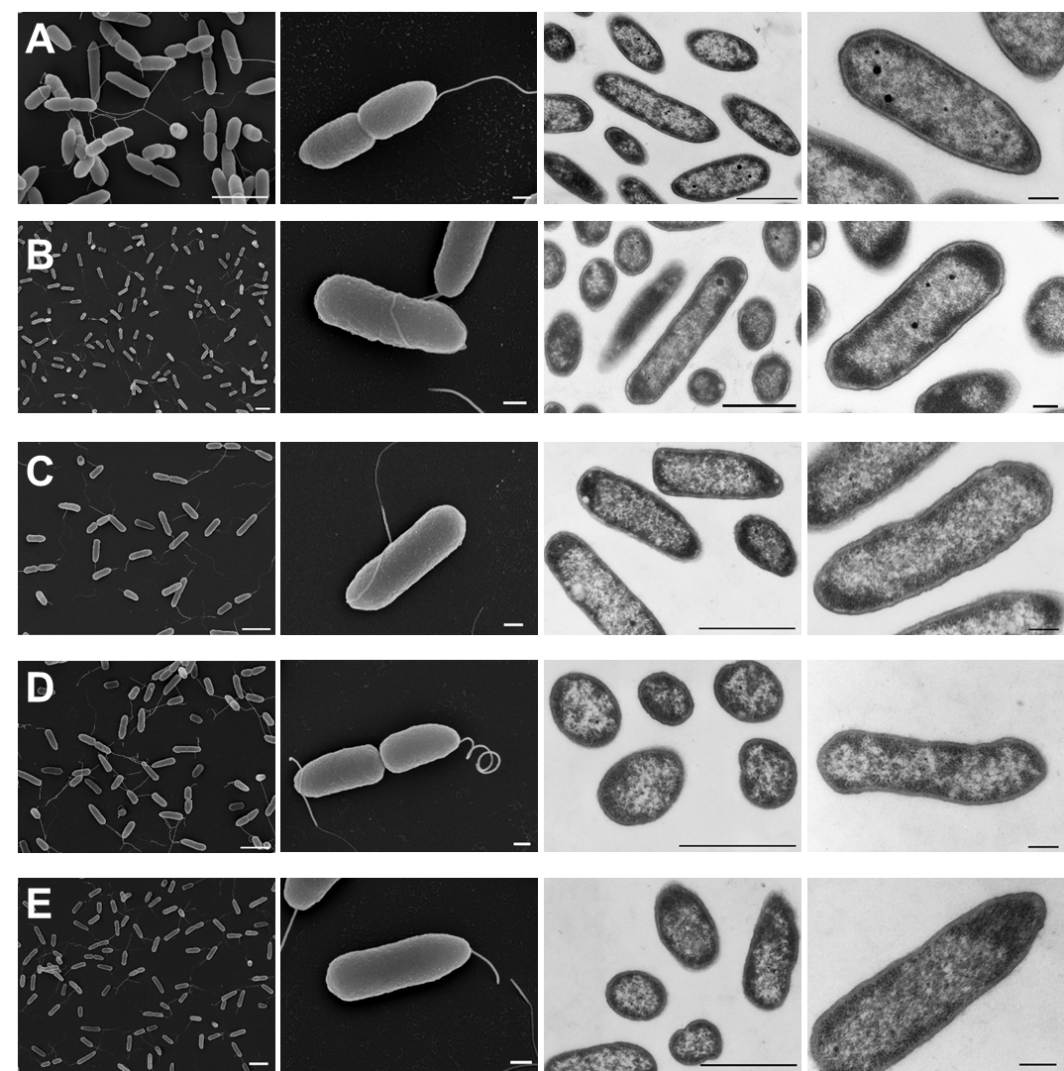

**Extended Data Fig. 11. Scanning and transmission electron micrographs of *P. aeruginosa* deletion strains harbouring the empty vector.** The figure shows, in 5 panels (**A-E**), scanning and transmission electron micrographs of control strains of *P. aeruginosa* used in this study. All images were taken after 5 hours growth. Panel A shows wildtype *P. aeruginosa* PA14: Panel B shows wildtype (PA14) + empty vector pSEVA634: Panel C shows *P. aeruginosa* ΔPa*lemA1* + pSEVA634: Panel D shows *P. aeruginosa* ΔPa*lemA2* + pSEVA634: Panel E shows *P. aeruginosa* ΔPa*lemA1+A2* + pSEVA634. No unusual phenotypes were observed for these strains. From left to right scale bars are 2, 0.2, 1 and 0.2 µm, respectively.

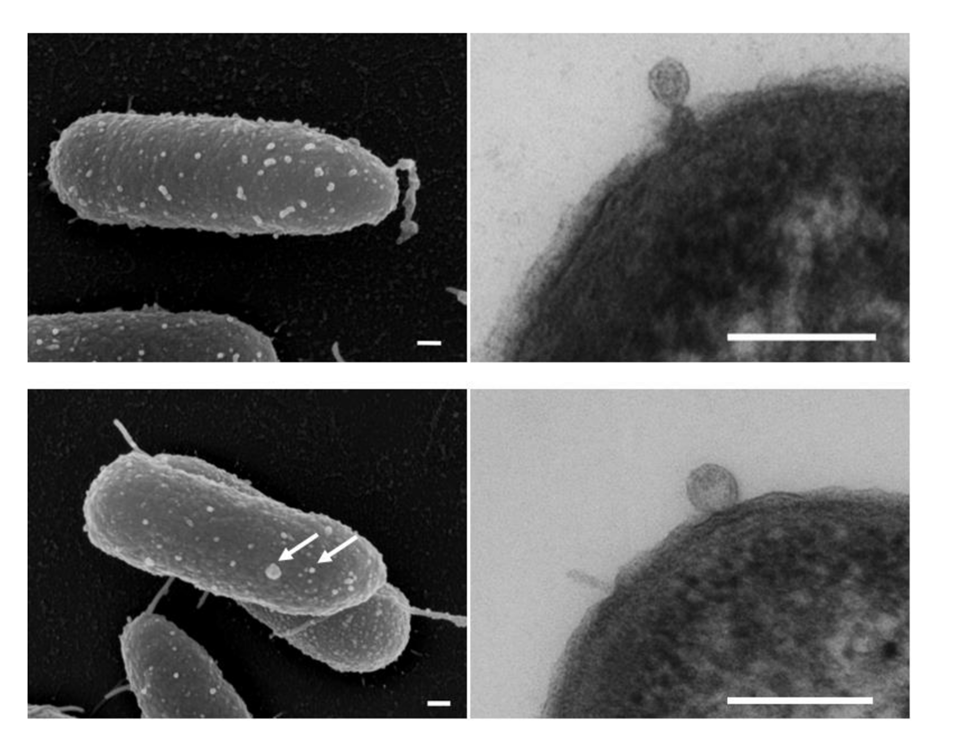

**Extended Data Fig. 12. Scanning and transmission electron micrographs of OMV formation by *P. aeruginosa* overproducing PaLemA2.** Snapshot of the vesiculation process of BMV formation in the hypervesiculating *P. aeruginosa* ΔPa*lemA*2+pPa*lemA*2 strain can be seen. OMVs/OIMVs are indicated by white arrows. Scale bars are 100 nm.

5. **Elucidation of the protein machinery for BMV formation – initial biochemical studies**

Our next aim was to get first biochemical insights into LemA protein function. First experiments included the recombinant production of all our various cloned LemA1 and LemA2 proteins (Fig. 2) to test them for their physical influence on lipid double layer structures as methodically outlined before for proteins like for example for the ESCRT-III-protein family proteins(37, 38). We even produced the *Thermatoga* *maritima* LemA of which a known crystal structure (without membrane helix) exists (PDB 2ETD) since 2005. All of the LemA1 proteins naturally containing a membrane spanning helix were additionally produced without their helix. With almost 30 different produced proteins we observed almost identical biochemical behaviours. Some of the proteins, as expected, especially the membrane helix-containing LemA1 proteins were insoluble and hardly soluble in the multiple tested detergents. For the residual soluble LemA proteins we observed a unique unspecific aggregation behaviour even in the presence of detergents. These observations led us to the assumption, that LemA protein show their specific biochemical function, as they do it in *in vivo* tests, only the concert with other proteins, most likely in a form of a dynamic membrane shaping complex composed of various different exchangeable components involved processes like cell division and intracellular organelle formation. Various components might be stable permanent parts, other are exchangeable and mediate process specificity. Next, an interactomics analyses using PaLemA1 and PaLemA2 were performed to elucidate the protein machinery of BMV formation.

6. **Elucidation of the protein machinery for BMV formation – an interactome study**

The bioinformatic interpretation of the data revealed a distinct group of proteins interacting with PaLemA1 and PaLemA2 as clearly visible in Extended Figure 13. In total 119 candidate proteins among the 1403 identified proteins differentially interact with PaLemA1, 133 out of 2259 with PaLemA2, and of which 13 interact with both proteins of interest (Extended Data Tables 4 & 5; the raw proteomic data will be deposited in the ProteomeXchange Consortium via the PRIDE (39) partner repository upon acceptance). As background control a PA14 WT proteome was analysed and used for normalization. The volcano plots shown in Extended Data Fig. 13 reveal the clear-cut separate clustering of these 119 proteins (in red) for PaLemA1 and 133 proteins for PaLemA2 (labelled in red). These proteins were further investigated for their potential function.

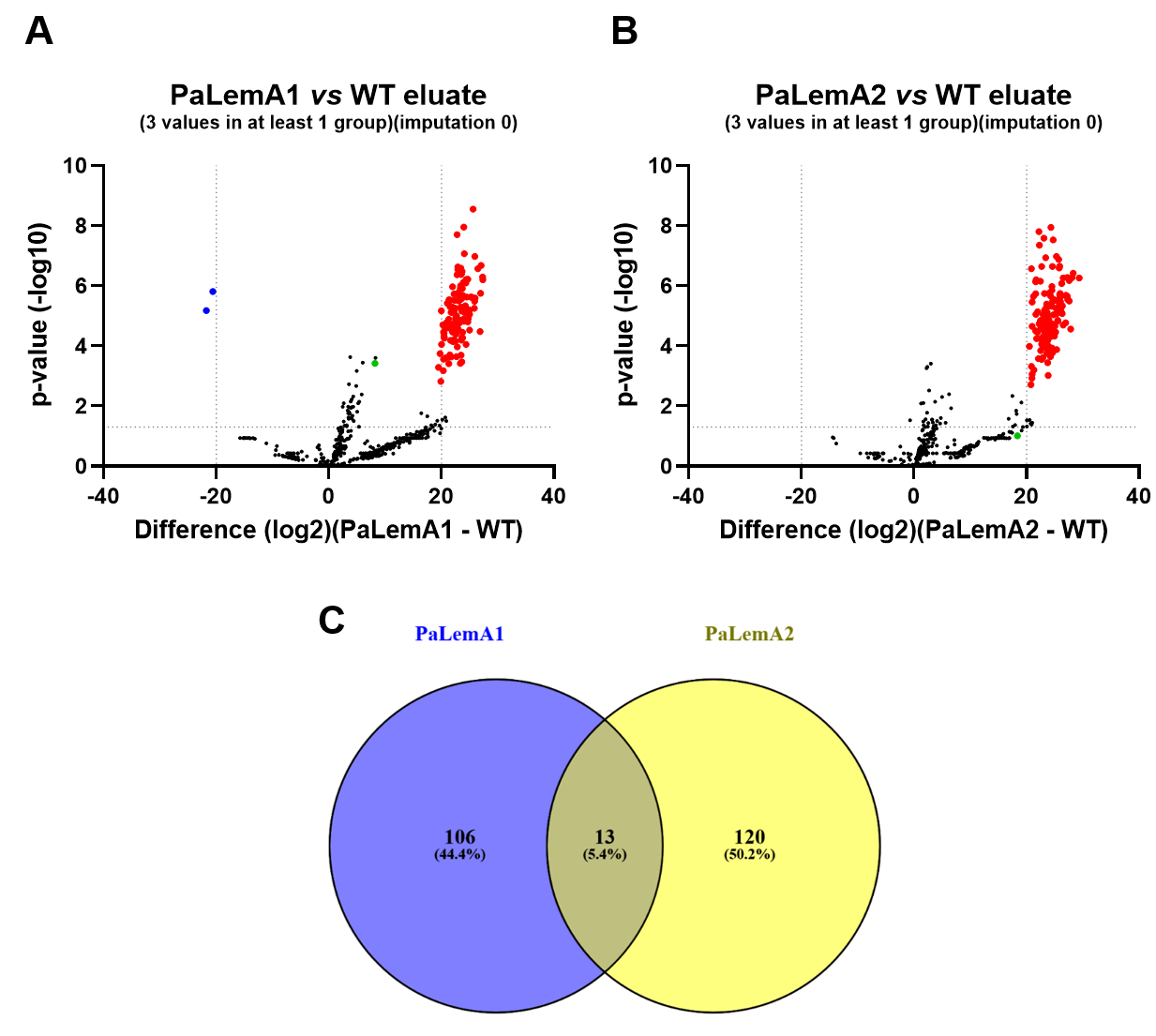

**Extended Data Fig. 13: Proteomic analyses of PaLemA1 and PaLemA2 interactomes**. Affinity-tagged PaLemA1 and PaLemA2 were purified and co-purified proteins (interactomes) were analysed by high resolution LC-MS/MS. Volcano plots show the distribution of PaLemA1 (A) and PaLemA2 (B) interacting proteins as compared to the wildtype lysates. Proteins selected for further analyses and discussion are given in red, PaLemA1 (**A**) and PaLemA2 (**B**) are indicated as green dots. (**C**) Venn diagram showing the number of PaLemA1 (119) and PaLemA2 (133) interacting proteins, of which 13 bind to both PaLemA1 and PaLemA2. (Co-)purification and proteomic analyses were performed in three independent biological replicates (n=3).

7. **The PaLemA1 and PaLemA2 interactome: proteins of cell division as structural basis for BMV and organelle formation**

The interactome study identified over 250 interaction partners for PaLemA1 and PaLemA2. These partners can be grouped into functional modules that collaborate to build vesicles. Firstly, the LemA complex co-opts core components of the divisome, the protein complex responsible for cell division, to build vesicles. This suggests vesicle formation is a "hijacked" or specialized form of cell division (40). This connects cell wall synthesis to outer membrane (OM) dynamics. It links the peptidoglycan (PG) synthase PBP1B to the OM via TolA, both PaLemA- bound (41). PaLemA1-bound FtsZ, FtsA and ZipA form the Z-ring, the cytoskeletal scaffold that defines the division plane (42). ZipA tethers FtsZ to the membrane. This machinery could be repurposed to create the force needed for membrane invagination during vesicle formation. Z-ring-associated protein ZapG, bound to PaLemA1, helps in stabilizing the membrane curvature machinery at the Z-ring (43). PaLemA1-bound FtsE as part of an ABC transporter-like complex regulates septal PG synthesis and Z-ring constriction (44). It links the Z-ring to downstream PG synthesis enzymes, coordinating constriction with new wall synthesis. FtsQLB, FtsW, FtsI (PBP3) are core divisome components (45). FtsI (PBP3) is the septal PG transpeptidase that builds the division wall. FtsW is its cognate flipase. These are the primary enzymes building the new cell wall at the division site. Their recruitment directly links vesicle formation to active peptidoglycan synthesis (46). PBP1A (PonA) was found bound to both PaLemA1 and PaLemA2 (47). It is a major peptidoglycan synthase. Its presence links general PG synthesis to the vesicle formation complex. PaLemA1- bound PBP1B (MrcB) is a key divisome-associated PG synthase, further solidifying the link between division and vesicle formation. Pa LemA2-bound PBP5 (DacC) is a DD-carboxypeptidase that trims PG peptides, regulating the level of cross-linking. PaLemA1- bound FtsN is a crucial activator of the division machinery, and its presence here suggests it might also activate vesicle formation (48) (Extended Data Table 6).

8. **The PaLemA interactome links vesicle formation to cell envelope biogenesis and lipid metabolism**

Extended Data Table 6 lists all enzymes of cell wall metabolism found bound to the PaLemA proteins. **UTP-glucose-1-phosphate uridylyltransferase (GalU) catalyses the general activation of glucose for further polymerization reactions including cell wall biosynthesis, glycogen, sucrose, starch, cellulose and pectin formation. For this purpose, glucose-1-phosphate** and UTP are converted to **UDP-glucose** and diphosphate. GalU-dependent LPS biosynthesis is responsible for the sensitivity to certain phages and antibiotics (49-52). **Here, a central enzyme of the O-antigen formation is recruited to the PaLemA organized super-complex.** The essential enzyme phosphoglucosamine mutase (GlmM) converts glucosamine-6-phosphate to glucosamine-1-phosphate during cell wall and LPS biosynthesis (53, 54). The 4-hydroxy-tetrahydrodipicolinate synthase (DapA) performs an aldol condensation of aspartate-semialdehyde and pyruvate yielding hydroxy-2,3,4,5-tetrahydrodipicolinate, an essential precursor of Lys and peptidoglycan biosynthesis. The **4-hydroxy-tetrahydrodipicolinate reductase DapB is** catalysing the conversion of 4-hydroxy-tetrahydrodipicolinate to tetrahydrodipicolinate, the product released by the DapA-catalysed reaction (55). *meso-*Diaminopimelate is required for cell wall biosynthesis and that makes DapA and DapB essential enzymes and antibiotic target in many bacteria (56). Additionally, several enzymes involved in peptidoglycan cross-linking were detected in the interactome, FtsI, MrcA and MrcB and DccA, all exhibiting DD-transpeptidase activity as the major step for cross-linking two peptidoglycan pentapeptide chains. FtsI is part of the division septum and localized in the septal ring, has an active domain accommodating two peptides to be joined and is also involved in the recruitment of FtsN (57). MrcA and MrcB catalyse additionally the transglycoyslation reaction, elongation both the glycan chain and forming cross-links in the growing peptidoglycan structure (58). DccA has been shown to play a role under specific stress conditions (59). Obviously, interaction of PaLemA1 with the rate-limiting step enzyme and at the same time with recycling enzyme allows for the control of this obviously important pathway (60). Finally, cell division does not only require *de novo* cell wall synthesis, but also cell wall recycling. The N-acetylglucosaminidase NagZ is a key enzyme of peptidoglycan recycling in Gram negative bacteria. It is cleaving the terminal 1,4-linked N-acetylglucosamine from peptide-linked peptidoglycan fragments, giving rise to free N-acetylglucosamine, anhydro-N-acetylmuramic acid and anhydro-N-acetylmuramic acid-linked peptides (61). Latter acts as signal molecules for the transcriptional regulator AmpR which in turn induces antibiotic resistance genes (62). Thus *P. aeruginosa* NagZ is valuable drug target for antibiotic development (63, 64). Additionally, we detected the diaminopimelate decarboxylase (LysA) which catalyses the last step of Lys biosynthesis, the decarboxylation of meso-diaminopimelate of the DAP pathway or released from peptidoglycan to form Lys and CO_2_.

**Extended Data Table 6:** **Complete** **annotated interaction partners of the PaLemA1 and PaLemA2 proteins involved in cell division, LPS, cell wall, and membrane metabolism.** Accession, locus tag, gene name, description and enrichment factor are given. Detailed protein information was added according to Uniprot database (https://uniprot.org) and the Pseudomonas database (https://pseudomonas.com). Analysis was performed with RStudio software, n = 3. Essential genes after Poulsen *et al*., 2019 (65) and Lee *et al*., 2015 (66) (accession and locus tag in red), only in Poulsen *et al*., 2019 (pink) and only in Lee *et al*., 2015 (green) are labelled.

| Accession | Locus tag | Gene | Description | Interactor of LemA1 (A1)/  LemA2 (A2).  Enrichment Factor |
| --- | --- | --- | --- | --- |
| *Cell division, membrane, cell wall, and LPS maturation* | | | | |
| Q02K11 | *PA14_44670* | *zipA* | Cell division protein ZipA, stabilizes the FtsZ protofilaments | A1 27.30 |
| A0A0H2ZFN5 | *PA14_57690* | *zapG* | Cell division protein ZapG (YhcB), links Z-ring to septal peptidoglycan synthesis | A1 26.86 |
| A0A0H2ZL67 | *PA14_04910* | *ftsE* | Cell division ATP-binding protein FtsE | A1 23.56 |
| A0A0H2ZC79 | *PA14_30290* | *ftsK* | DNA translocase FtsK | A1 23.64 |
| A0A0H2ZGU2 | *PA14_62860* | *ftsH* | ATP-dependent zinc metalloprotease FtsH | A2 24.78 |
| A0A0H2ZIC7 | *PA14_65270* | *hflC* | HflC regulates the protease FtsH | A1 25.94 |
| A0A0H2ZEZ2 | *PA14_14860* | *-* | **HTH cro/C1-type domain-containing protein, similiar to cytoskeleton protein binding protein RodZ** | A1 25.91 |
| A0A0H2ZIQ4 | *PA14_66760* | *-* | **Putative cell division protein, cell wall binding SPOR domain-containing, divisome targeting, FtsN-like** | A2 22.52 |
| A0A0H2Z7H9 | *PA14_51690* | *cpoB* | Cell division coordinator CpoB | A1 23,42  A2 26,34 |
| A0A0H2ZES6 | *PA14_16150* | *pspA* | Phage shock protein, PspA (ESCRT-III-type protein), BMV formation | A1 24,47 |
| A0A0H2ZLI8 | *PA14_57425* | *ftsI* | Bifunctional peptidoglycan synthase (transglycosylase/ transpeptidase) Penicillin-binding protein 3, PBP3 at the division septum | A2 23.40 |
| A0A0H2ZIP9 | *PA14_66670* | *ponA*  *mrcA* | Bifunctional peptidoglycan synthase (transglycosylase/ transpeptidase) penicillin-binding protein 1A, PBP1A | A1 22,24  A2 22,26 |
| A0A0H2ZHP9 | *PA14_62200* | *mrcB* | Bifunctional peptidoglycan synthase (transglycosylase/ transpeptidase) Penicillin-binding protein 1B, PBP1B | A1 25.01 |
| A0A0H2ZFH3 | *PA14_12100* | *dacC* | Bifunctional peptidoglycan synthase (transglycosylase/ transpeptidase) Penicillin-binding protein 5/PBP5 | A2 28.08 |
| A0A0H2ZAA4 | *PA14_38350* | *galU* | **UTP-glucose-1-phosphate uridylyltransferase** (peptidoglycan and LPS biosynthesis) | A2 20.49 |
| Q02FS3 | *PA14_62840* | *glmM* | Phosphoglucosamine mutase (LPS + peptidoglycan synthesis) | A1 22.26 |
| A0A0H2Z7V1 | *PA14_51270* | *dapA* | 4-hydroxy-tetrahydrodipicolinate synthase, meso-DAP biosynthesis for peptidoglycan | A1 21.64 |
| Q02FR3 | *PA14_62940* | ***dapB*** | **4-hydroxy-tetrahydrodipicolinate reductase,** meso-DAP biosynthesis for peptidoglycan | A2 20.94 |
| A0A0H2ZJ02 | *PA14_69670* | *lysA* | Diaminopimelate decarboxylase,  L-lysine biosynthesis via DAP pathway | A1 21.31 |
| A0A0H2Z802 | *PA14_49280* | *-* | Lytic murein transglycosylase family | A2 23.02 |
| Q02PG9 | *PA14_25195* | *nagZ* | Beta-N-acetylhexosaminidase, peptidoglycan recycling, NagZ | A1 22,46 |
| Q02Q76 | *PA14_22010* | *minE* | Cell division topological specificity factor MinE | A1 23,53 |
| A0A0H2ZHJ7 | *PA14_61990* | ***parA*** | **Chromosome partitioning protein ParA** | A1 20.38 |
| A0A0H2ZJW9 | *PA14_73330* | *parB* | Chromosome-partitioning protein ParB | A1 22.92 |
| Q02GB1 | *PA14_60445* | *obg* | GTPase Obg controls via ppGpp HokB toxin which causes membrane potential colapse | A1 24.66 |
| A0A0H2ZHS5 | *PA14_62120* | *pssA* | CDP-diacylglycerol--serine O-phosphatidyltransferase, phospholipid biosynthesis | A1 23.74 |
| Q02PB3 | *PA14_25900* | *fabV* | NADH-dependent enoyl-acyl-carrier-protein reductase FabV (fatty acid biosynthesis) | A1 23.46 |
| Q02RB1 | *PA14_17270* | ***accA*** | **Acetyl-coenzyme A carboxylase carboxyl transferase subunit Alpha** (fatty acid biosynthesis) | A2 20.99 |
| Q02RE5 | *PA14_1686* | *plsB* | **Glycerol-3-phosphate acyltransferase,** (diacylglycerol biosynthesis) | A1 21.97 |
| A0A0H2ZGS4 | *PA14_57840* | ***ttg2D***  ***mlaC*** | General glycerophospholipid transporter across the periplasm | A2 23.71 |
| A0A0H2ZFP5 | *PA14_57850* | *mlaD* | Outer membrane lipid asymmetry maintenance protein | A2 23.76 |
| A0A0H2Z976 | *PA14_44080* | *-* | START domain-containing protein, polyketide cyclase / dehydrase and lipid transport family protein | A2 24.62 |
| A0A0H2ZCS3 | *PA14_27730* | *yafH* (*fadE*) | Acyl-coenzyme A dehydrogenase (fatty acid degradation) | A1 23,49 |
| Q02PH8 | *PA14_25080* | ***fadB*** | **Fatty acid oxidation complex subunit alpha** | A2 20.86 |
| A0A0H2ZLD3 | *PA14_05860* | ***-*** | **Long-chain acyl-CoA thioester hydrolase family protein** | A2 21.80 |
| A0A0H2ZGK8 | *PA14_56850* | *-* | Putative lipoprotein, neighbour of LemA1 encoding gene | A2 24.76 |
| A0A0H2ZJN1 | *PA14_71840* | *yfjO* | **Outer membrane assembly lipoprotein YfiO** | A2 22.90 |
| A0A0H2ZID3 | *PA14_66240* | *waaC* | Lipopolysaccharide heptosyltransferase I (LPS biosynthesis) | A1 22.98 |
| A0A0H2ZHE9 | *PA14_66250* | *waaF* | Heptosyltransferase II (LPS biosynthesis) | A1 22.61 |
| A0A0H2Z7E1 | *PA14_52150* | *lpxO2* | Lipopolysaccharide biosynthetic protein LpxO2 | A1 22.55 |
| A0A0H2ZGK2 | *PA14_57920* | *lptA* | Lipopolysaccharide export system protein LptA | A2 24.31 |
| A0A0H2ZIU6 | *PA14_69090* | *yhiI* | Putative membrane fusion protein | A2 23.80 |
| A0A0H2Z7R1 | *PA14_51730* | *tolA* | Cell envelope integrity protein TolA | A2 23.61 |
| Q02NA6 | *PA14_30310* | *lolA* | Outer-membrane lipoprotein carrier protein | A2 23.19 |
| A0A0H2ZFA6 | *PA14_13140* | *-* | Lipoprotein LolA-like, potential outer membrane lipoprotein-sorting protein | A2 23.06 |
| A0A0H2ZC15 | *PA14_24300* | *-* | Lipoprotein LolA-like, potential outer membrane lipoprotein-sorting protein | A2 24.23 |
| A0A0H2Z614 | *PA14_53500* | *-* | **OmpA-like domain-containing protein** | A1 20.25 |
| A0A0H2ZKV8 | PA14_67975 | - | AsmA family outer membrane biogenesis protein | A1 22.46 |
| A0A0H2Z6T6 | *PA14_50140* | ***fliF*** | **Flagellar M-ring protein** | A2 22.56 |
| A0A0H2ZC68 | *PA14_23830* | ***fimV*** | **Motility hub protein,** type IV pilus- assembly, twitching motility, cAMP-dep. virulence gene expression | A1 21.86 |
| A0A0H2ZIN8 | *PA14_66630* | ***pilP*** | **Type 4 fimbrial biogenesis protein,** recruited to sites of cell division | A2 23.84 |

9. **The PaLemA1 and PaLemA2 interactome: proteins of cytoplasmic membrane biogenesis and recycling**

**Various proteins of cytoplasmic membrane turnover were found attached to the PaLemAs (Extended Data Table 6). Glycerol-3-phosphate 1-*O*-acyltransferase PlsB (PaLemA1) is catalysing the initial step of phospholipid biosynthesis, the ligation of** *sn*-glycerol-3-phosphate with an acyl ACP to form 1-acyl-sn-glycerol-3-phosphate (67, 68). PlsB was found involved in multidrug resistance, biofilm formation and virulence (69, 70). Similarly, the alpha subunit (AccA) of the essential initial key enzyme of fatty acid biosynthesis **acetyl-coenzyme A carboxyltransferase was found attached to PaLemA2.** First, biotin carboxylase catalyses the carboxylation of biotin on its carrier protein and then hydrogencarbonate is transferred by the carboxyltransferase with its subunits alpha (AccA) and beta (AccD) to acetyl-CoA to form malonyl-CoA, the general precursor of fatty acid biosynthesis (71, 72). Next, PaLemA1 interacting essential enoyl-[acyl-carrier-protein] reductase FabV catalyses the last reduction of trans-2-enoyl-ACP to the fully saturated acyl-ACP species in the fatty acid elongation cycle (73). FabV is a potent target of novel antibiotics (74). Like the cell wall also membrane lipids are subject to continuous turnover (75). Thus, central enzymes of β-oxidation were found attached to PaLemA proteins. Long-**chain acyl-CoA thioester hydrolases** catalyse the hydrolysis of acyl-CoAs to the free fatty acid and CoASH. Next, FadE (Acyl-coenzyme A dehydrogenase) catalyses the FAD-dependent oxidation of acyl-CoAs to 2-enoyl-CoAs. Then FadB, the α-subunit of the multienzyme complex FadBA with dodecanoyl-CoA delta-isomerase, enoyl-CoA hydratase, 3-hydroxybutyryl-CoA epimerase and 3-hydroxyacyl-CoA dehydrogenase activities continues fatty acid degradation (76, 77). Essential PssA (CDP-diacylglycerol-serine O-phosphatidyltransferase) involved in stress-induced membrane remodelling were found all attached to PaLemA1 (78).

10. **PaLemA1 and PaLemA2 change BMV lipid composition**

We found several enzymes of lipid, cell wall and LPS biogenesis bound to PaLemA1 and PaLemA2 **(Extended Data Table 6)**. In order to test the influence of PaLemA1 and PaLemA2 on the lipid composition of BMV we first measured changes in lipid composition of the cells and OMV/OIMV over time (Extended Data Fig. 14). Surprisingly, we observed almost no changes in lipid composition for the cells over time while the OMV/OIMV showed a decrease of long-chain and unsaturated lipids with an increase in mid-chain and especially hydroxy fatty acids with time (Extended Data Fig. 14). Mid-chain and hydroxy fatty acids are constituents of the lipid A of the LPS. These observations indicate that there is a higher amount of LPS present in the vesicle fraction which indicates that there are more OMVs present in the vesicle preparation. Next, we followed up the question, do PaLemA1 and PaLemA2 have something to do with these changes, Thus, we compared the lipid composition of OMV/OIMV after 6 hours of growth of the wildtype, the single and the double mutant of Pa*lemA1* and Pa*lemA2*. Overall, the major trends observed for wildtype OMV/OIMV over time are more or less disrupted in the various mutants (Extended Data Fig. 15). Mid chain lipid essentially remained constant, while the decrease of long-chain and unsaturated lipids with an increase of hydroxy lipids was stopped. Obviously, the PaLemA1 and PaLemA2 mediated protein-protein contacts contribute to the restructuring of the vesicle composition indicating an altered LPS share in the lipid composition.

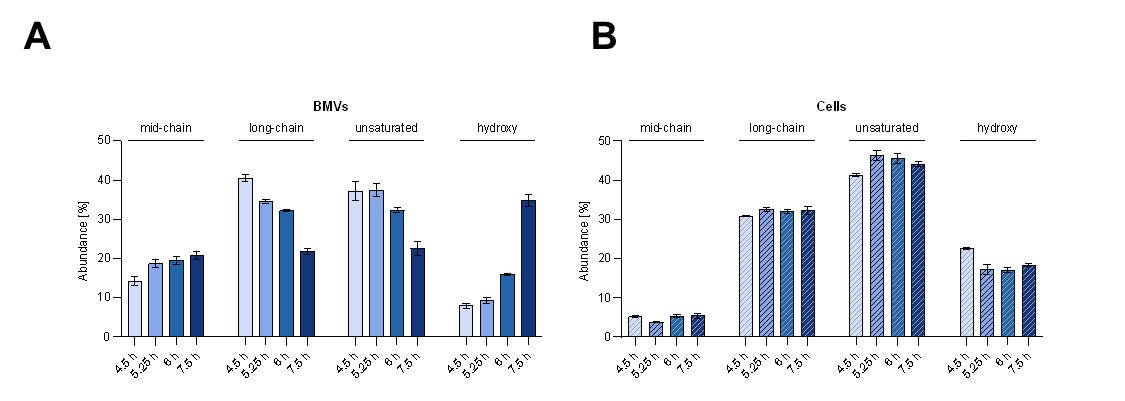

**Extended Data Fig. 14: Characterization of the fatty acid composition of *P. aeruginosa* cells and derived BMVs over time.** To analyse fatty acid composition of BMVs (**A**) and corresponding cells (**B**) in the course of time, BMVs were isolated after 4.5, 5.25, 6 and 7.5 hours and analysed along with the corresponding cells regarding their fatty acid composition. Fatty acids were identified using GC-MS and GC-FID and categorized as follows: mid-chain (≤ 12 carbons), long-chain (13–21 carbons), unsaturated (≥ one carbon-carbon double bond or cyclopropane group) and hydroxy (≥ one hydroxyl group). For each category, the sum of all relevant fatty acids was calculated as a percentage of the total fatty acid content per biological replicate. Data represent the mean ± SEM of three biological replicates.

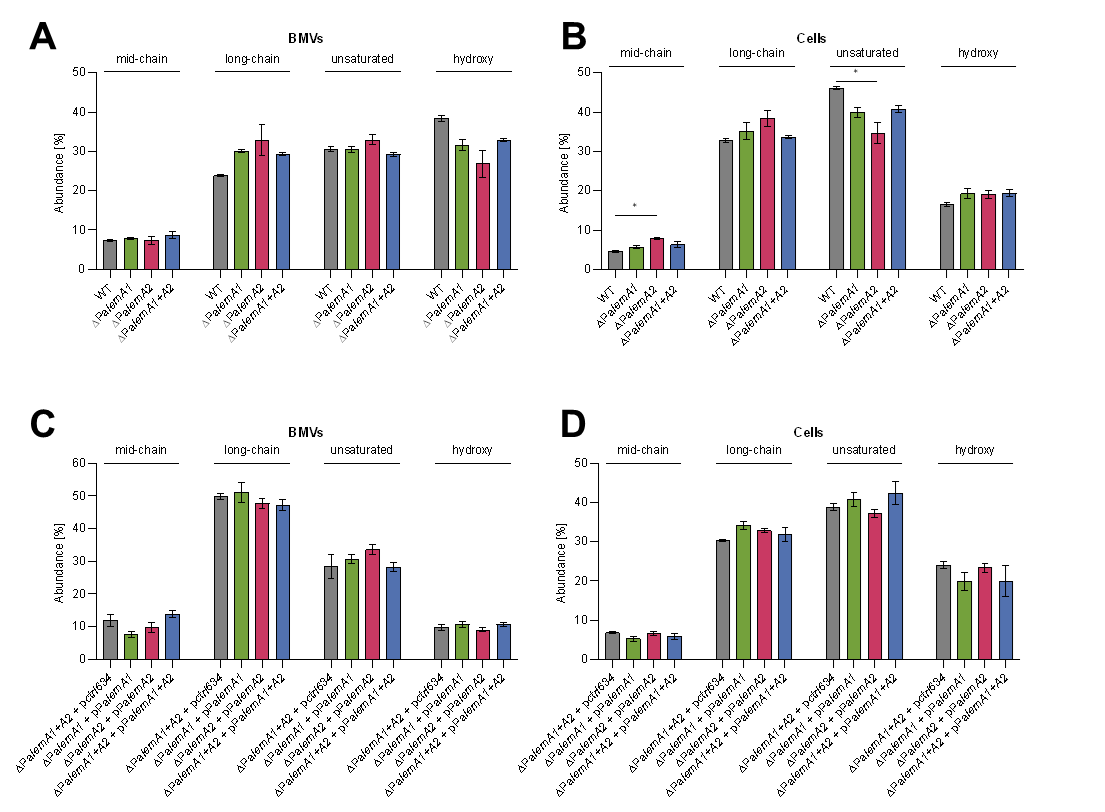

**Extended Data Fig.15. Characterization of the fatty acid composition of *P. aeruginosa lemA* mutant cells and derived BMVs.** This figure shows the impact of Pa*lemA* knockout (**A, B**) and overexpression (**C, D**) on the fatty acid composition of cells and derived BMVs. For knockout strains, samples were harvested and isolated in early stationary phase (6 h) while cells and BMVs from the recombinantly overproducing strains were isolated 2 hours post induction with 100 µM IPTG. Fatty acids were identified using GC-MS and GC-FID and categorized as follows: mid-chain (≤ 12 carbons), long-chain (13–21 carbons), unsaturated (≥ one carbon-carbon double bond or cyclopropane group) and hydroxy (≥ one hydroxyl group). For each category, the sum of all relevant fatty acids was calculated as a percentage of the total fatty acid content per biological replicate. Data represent the mean ± SEM of three biological replicates.

11. **The PaLemA1 and PaLemA2 interactome: proteins of outer membrane and LPS biogenesis and recycling**

The outer membrane itself is composed of an inner leaflet with glycerophospholipids and an outer leaflet made of LPS with inserted integral β-barrel proteins and lipoproteins. Enzymes and transporters involved in the corresponding metabolism were interaction partners of the PaLemA proteins **(Extended Data Table 6)**. The transport of glycerophospholipids through the periplasm to the outer membrane is mediated by PaLemA-bound phospholipid binding protein Ttg2D (MlaC) and the outer membrane lipid asymmetry maintenance protein (MlaD) as part of the Ttg2ECBA (MlaBDEF) transport system (79, 80). Similarly, the family of AsmA proteins mediate the transport of the glycerophospholipids from the inner membrane to the outer membrane (81). One AsmA protein and one Asm family protein was found bound to PaLemA1. LPS is composed of a hydrophobic domain called lipid A, a core oligosaccharide and a distal polysaccharide named *O*-antigen. Both heptosyltransferases WaaF und WaaC (both bound by PaLemA1) are responsible for the synthesis of the inner core of the core oligosaccharide (82, 83). The dioxygenase LpxO2 (bound by PaLemA1) performs a site-specific lipid A hydroxylation which is important for the adaptation to the cystic fibrosis lung (84). LptA (bound by PaLemA2) is the only entirely periplasmic subunit of the seven proteins containing LPS transport system. The protein protects the hydrophobic acyl chains of LPS during their passage through the hydrophilic periplasm. Thus, LptA is essential for LPS formation and cell viability (85). The periplasmic chaperone LolA (bound by the lipoprotein PaLemA2) is essential for outer membrane lipoprotein formation. It accepts triacylated lipoproteins from the inner membrane transporter LolCDE and carries them across the periplasm to the corresponding outer membrane receptor LolB (86). Without LolA no lipoproteins can be transported to the outer membrane. Further, two LolA-like proteins were found bound to PaLemA2. The essential integral inner membrane protein TolA is a central component of the Tol-Pal system which ensures integrity of the outer membrane and is central to cell division (41, 87). The protein interacts with TolB, TolQ, TolR and Pal proteins. TolA uses a long α-helix to span the periplasm forming a bridge between inner and outer membrane. The protein is recruited to the septum during cell division and uses the proton motive force via TolQ/R to perform conformational changes to ensure membrane integrity (88). The essential FliF protein forms the MS-ring of the flagellar motor and serves as assembly platform for the whole flagellum (89). FimV is a polar landmark protein composed a peptidoglycan-binding LysM-like domain in the periplasm, a transmembrane part, a cytoplasmic “FimV domain” composed of tetrapeptide repeats (90). Protein is essential for the proper polar placement of the chemotaxis system (91, 92). Finally, PilP is a periplasmic lipoprotein playing a key role in the assembly and function of type-V-pili. It connects the inner membrane PilMNO complex with the outer membrane secretion pore PilQ forming another bridge between inner and outer membrane (93). FimV serves as one of the hub proteins in this process.

12. **The PaLemA1 interactome identifies the ESCRT-III-like protein PaPspA as a modulator of vesicle formation**

We have shown that extracellular BMV biogenesis rates are highly dependent on the action of both, PaLemA1 and PaLemA2 proteins since deletion of each gene independently does not lead to significant decrease in BMV formation but disruption of both impairs the process (Fig. 3). Our interactome studies identified a known player of bacterial vesicle formation PaPspA (94, 95) as interaction partner of PaLemA1. To investigate a potential coordinated action of PaLemA1 and PaLemA2 with PaPspA we assessed the impact of the deletion and overexpression of the Pa*pspA*-gene on extracellular BMV formation by fluorescence quantification. Moreover, we monitored whether overexpression of Pa*lemA* genes in a Pa*pspA*-lacking strain impacted overall extracellular BMV generation. The deletion or overexpression of Pa*pspA* did not show significant bacterial surface alterations as observed by electron microscopy (Extended Data Fig. 16a). The Pa*pspA* gene lacking strain was affected slightly in extracellular vesiculation rates in comparison to the wildtype control strain (Extended Data Fig. 16b). Interestingly, Pa*pspA* overexpression in the mutant background strain led to diminishment in extracellular BMV formation (Extended Data Fig. 16b). This possibly indicates a stoichiometry-impairment of the PaLemA-PaPspA multiprotein complex. This observation was supported by the fact that overexpression of each Pa*lemA* genes in the *P. aeruginosa* Pa*pspA* knockout mutant also impaired vesiculation rates (Extended Data Fig. 16b). This effect was magnified when both Pa*lemA1* and Pa*lemA2* genes were concurrently overexpressed in the Pa*pspA* mutant (Extended Data Fig. 16b). Interestingly, Pa*pspA*-deficient strains overexpressing the Pa*lemA* genes accumulated large amounts of extracellular cellular material and debris unrelated to BMVs (Extended Data Fig. 16a). This phenotype may reflect an impaired ability of the background strain to reseal the membrane following BMV detachment, a function previously attributed to PspA. Additionally, PaPspA does not influence the lipid composition of the OMV/OIMV (Extended Data Fig. 16c). Overall, similar to the PaLemA1 and PaLemA2 PaPspA is a known, but not essential part of the vesiculation complex indicating a certain degree of plasticity in complex composition. However, once present the stoichiometry of the different partners in the complex is essential for efficient BMV formation.

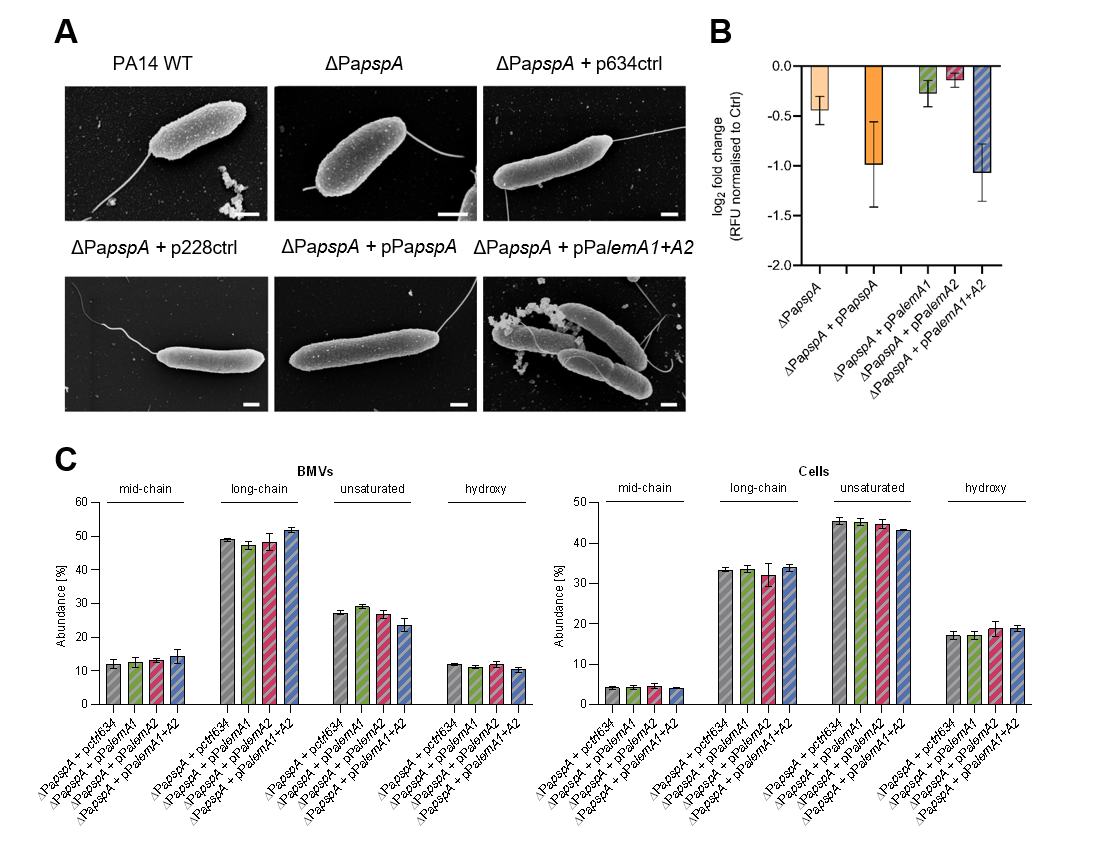

**Extended Data Fig. 16. Impact of disrupting the PaLemA-PaPspA multiprotein complex in *P. aeruginosa*.** (**A**) FESEM revealed that overproduction of both PaLemA1 and PaLemA2 in a Pa*pspA*-deficient background led to an accumulation of extracellular matrix. (**B**) Quantification of BMVs from Pa*pspA* knockout and Pa*pspA*-deficient strains overexpressing Pa*pspA*, Pa*lemA1*, Pa*lemA2* or both Pa*lemAs* collectively was performed using the fluorescent membrane binding dye FM4-64 to determine lipid content. BMVs derived from ∆Pa*pspA* and its corresponding control (PA14 WT) were isolated in early stationary phase. BMVs from plasmid-bearing strains were isolated 2 hours post induction with 100 µM IPTG (Pa*lemA* overexpression) or 100 µM 3-*m*Bz (Pa*pspA* overexpression). Strains harbouring the empty vectors (p*ctrl634* and p*ctrl228*) served as controls. All BMV samples were normalized to the mean relative fluorescence (RFU) of their respective control. Data are represented as the log_2_ fold-change and show the mean ± SEM of three biological replicates. (**C**) Fatty acid profiling of BMVs and cells from Pa*pspA* mutants overexpressing Pa*lemAs (*∆Pa*pspA+*pPa*lemA*) was conducted 2 hours post induction with 100 µM IPTG. Fatty acids were identified using GC-MS and GC-FID and categorized as follows: mid-chain (≤ 12 carbons), long-chain (13–21 carbons), unsaturated (≥ one carbon-carbon double bond or cyclopropane group) and hydroxy (≥ one hydroxyl group). For each category, the sum of all relevant fatty acids was calculated as a percentage of the total fatty acid content per biological replicate. Data represent the mean ± SEM of three biological replicates.

13. **The PaLemA interactome identifies PaCpoB as a key regulator of vesicle formation**

The key regulatory protein of bacterial cell division CpoB was found bound to PaLemA1 and PaLemA2 during the interactomics studies proteins (Extended Data table 6). To validate the protein-protein interactions identified from the interatomic data, we performed *in vivo* experiments using the bacterial adenylate cyclase two-hybrid (BACTH) system. Specifically, we investigated whether the CpoB protein of *P. aeruginosa* (PaCpoB) is able to interact with PaLemA1 and PaLemA2. The cytoplasmic domain of PaLemA1, lacking the predicted transmembrane helix and periplasmic domain (amino acids 28-190), was fused to the C-terminus of T25 and T18 fragments of adenylate cyclase (LemA1_25 and LemA1_18C). Similarly, the PaLemA2 protein lacking the signal peptide (amino acids 20-202) was fused to the T25 and T18 subdomains, leaving free the PaLemA2 C-terminal region (LemA2_25 and LemA2_18C). For PaCpoB, the leader sequence was excluded (20-274) and fusions to T25 and T18 subdomains were generated at both the N- and C-termini (CpoB_25, CpoB_25N, CpoB_18 and CpoB_18C). After generating the constructs of interest, β-galactosidase activity was assessed by growth on LB agar plates supplemented with X-gal and by quantitative β-galactosidase assays. The results showed that the cytoplasmic region of PaLemA1 fused to T18 interacted with PaCpoB fused to the C-terminus of T25 (CpoB_25 / LemA1_18C), as evidenced by blue colony formation and high β-galactosidase activity (Fig. 5). The PaLemA1–PaCpoB interaction was also detected when PaCpoB was fused to the N-terminus of T25 (CpoB_25N / LemA1_18C), although with lower β-galactosidase activity, as revealed by the quantitative assays (Fig. 5). In contrast, no interaction between PaLemA1 and PaCpoB was detected, either on indicator plates or in β-galactosidase assays, when the cytoplasmic fragment of LemA1 was fused to T25, regardless of whether PaCpoB was fused to the N- or C-terminus of T18 (CpoB_18 / LemA1_25 and CpoB_18C / LemA1_25) (Fig. 5). In the case of PaLemA2, blue colonies and high β-galactosidase activity were observed for all tested combinations, indicating that PaLemA2 interacts strongly with PaCpoB, independently of whether PaCpoB was fused to the N- or C-terminus of T25 or T18 (Fig. 5).

In addition, interactions between PaLemA1 and PaLemA2 and the proteins FstN and MrcB were investigated. For this purpose, the periplasmic domain of FstN, excluding the cytoplasmic transmembrane domain and the leader peptide (amino acids 44–231), was fused to T25 and T18. In the case of MrcB, T25 and T18 were fused to the N-terminal region corresponding to its cytoplasmic domain (amino acids 1–22). All combinations tested (FstN-PaLemA1, FstN-PaLemA2, MrcB-PaLemA1, and MrcB-PaLemA2) indicated weak or no interaction, as evidenced by almost white colonies and basal levels of β-galactosidase activity (Extended Data Fig. 17).

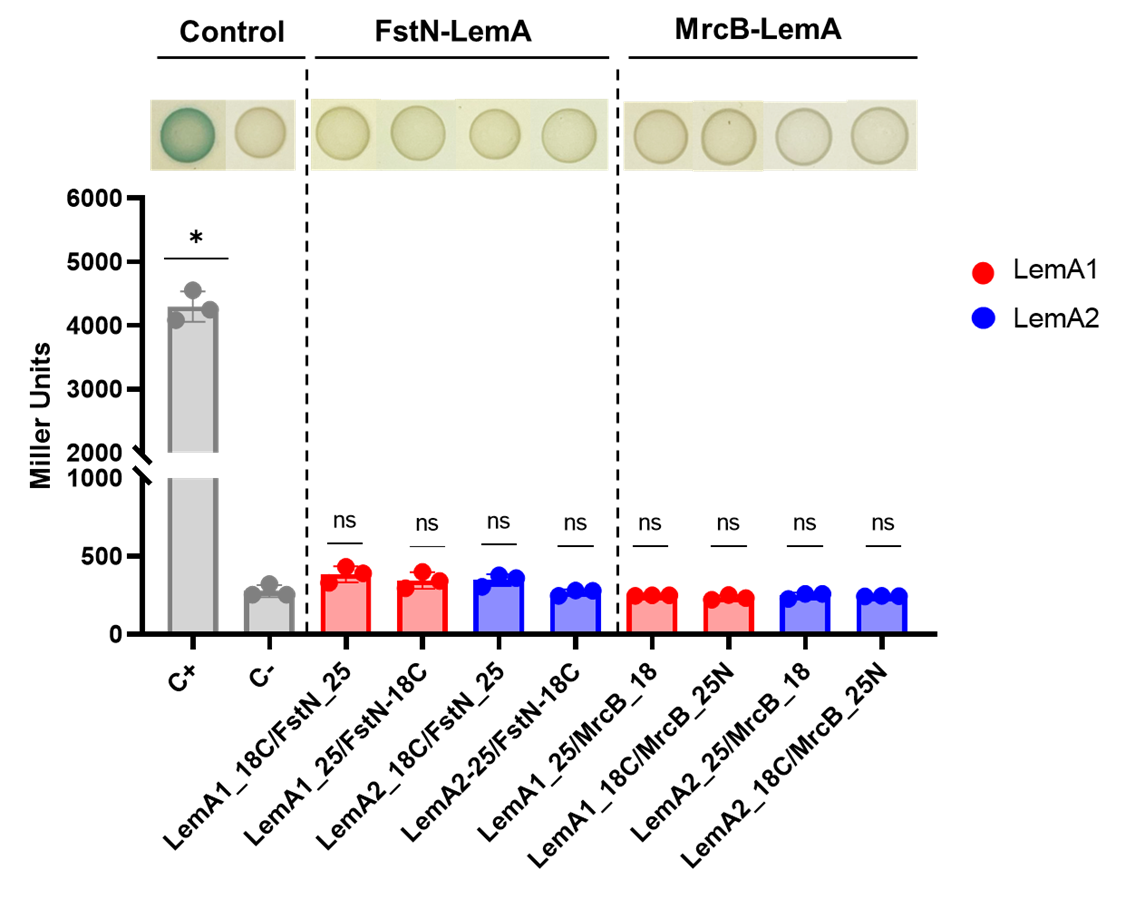

**Extended Data Fig. 17. Bacterial two-hybrid results for PaFstN-PaLemA and PaMrcB-PaLemA interactions.** β-galactosidase activity was measured in *E. coli* BTH101 cells co-transformed with the indicated T18 and T25 fusion constructs. Leucine zipper fusions were used as a positive control (+), while empty T18 and T25 vectors served as a negative control (-). β-galactosidase activity values were statistically compared to the empty vector using a Mann-Whitney nonparametric test. Asterisks (*) denote statistically significant differences at *p*-value = 0.05. Representative colonies from each co-transformation are shown above the graph; blue coloration (LacZ activity) indicates protein-protein interaction.

14. **The PaLemA links vesicle biogenesis to protein export and cargo loading**

Sec- and TAT-dependent protein transport is essential for almost all enzymatic, regulatory and structural protein activities of the IM, periplasmic space and outer membrane (96). First of all, SecY was found of one of the rare proteins to be bound to both PaLemA1 and PaLemA2 (Extended Data, Table 7). Further, the proteins SecA, SecB, SecD, SecF, the signal peptidase LepB and the Sec-associated proteins PpiD, YajC and PA14_14900 with the periplasmic thiol:disulfide interchange proteins DsbC and DsbG and the periplasmic chaperone SurA were interaction partners of PaLemA2 (Extended Data Table 6). The LemA functional picture gets further complete when one looks at the SecYEG interactome (97). In agreement the interactome showed the interaction of the SecYEG translocon to the PaLemA1 and 2 interaction partners FtsH, SecA, SecD, and the before mentioned HflC. Finally, The TAT translocase TatA is part of the PaLemA1 interactome (Extended Data Fig. 18). Overall, PaLemA interaction partners involved in protein export and modification are important for the periplasmic protein complex sustaining BMV formation.

**Extended Data Table 7:** **Annotated interaction partners of the PaLemA1 and PaLemA2 involved in protein export.** Accession, locus tag, gene name, description and enrichment factor are given.

Detailed protein information was added according to Uniprot database (https://uniprot.org) and the Pseudomonas database (https://pseudomonas.com). Analysis was performed with RStudio software, n = 3. Essential genes after Poulsen *et al*., 2019 and Lee *et al*., 2015 (accession and locus tag in red), only in Poulsen *et al*., 2019 (pink) and only in Lee *et al*., 2015 (green) are labelled.

| Accession | Locus tag | Gene | Description | Interactor of LemA1 (A1) /LemA2 (A2).  Enrichment Factor |
| --- | --- | --- | --- | --- |
| Protein export | | | | |

| Q02H37 | *PA14_57220* | *secA* | Protein translocase subunit SecA | A2 26.02 |
| --- | --- | --- | --- | --- |
| Q02EN8 | *PA14_67720* | *secB* | Protein-export protein SecB | A2 23.96 |
| A0A0H2ZF84 | *PA14_14630* | *secD* | Protein translocase subunit SecD | A2 28.27 |
| A0A0H2ZFD4 | *PA14_14650* | *secF* | Protein-export membrane protein SecF | A2 26.14 |
| A0A0H2ZGC8 | *PA14_09050* | *secY* | Protein translocase subunit SecY | A1 22.68  A2 24.18 |
| A0A0H2ZA00 | *PA14_41190* | *ppiD* | Periplasmic chaperone, associated with periplasmic part of Sec system | A2 27.84 |
| A0A0H2ZE27 | *PA14_14900* | *-* | Ancillary SecYEG translocon subunit | A2 26.51 |
| A0A0H2ZF92 | *PA14_14610* | *yajC* | Sec translocon accessory complex subunit YajC | A2 24.52 |
| A0A0H2Z5X4 | *PA14_54350* | *lepB* | Signal peptidase I | A2 22.27 |
| A0A0H2ZEZ9 | *PA14_16050* | *dsbC* | Thiol:disulfide interchange protein | A2 25.37 |
| A0A0H2ZBN7 | *PA14_32610* | *dsbG* | Thiol:disulfide interchange protein | A2 23.33 |
| A0A0H2ZLB3 | *PA14_07760* | *surA* | Periplasmic chaperone SurA | A2 23.17 |
| Q02FR0 | *PA14_62990* | *grpE* | Nucleotide exchange factor for Hsp70 chaperones | A2 25.31 |
| A0A0H2ZHB4 | *PA14_60190* | *clpB* | Chaperone protein ClpB | A2 21.69 |
| A0A0H2ZCK1 | *PA14_21880* | *tsp* | Periplasmic tail-specific protease | A2 26,33 |
| A0A0H2ZA00 | *PA14_41190* | *ppiD* | Associated with periplasmic part of Sec system | A2 27.84 |
| +Q02EU9 | *PA14_66960* | *tatA* | Sec-independent protein translocase protein TatA | A1 25.64 |
| A0A0H2ZID1 | *PA14_66580* | *-* | **Putative Type II secretory pathway protein** | A2 23.93 |

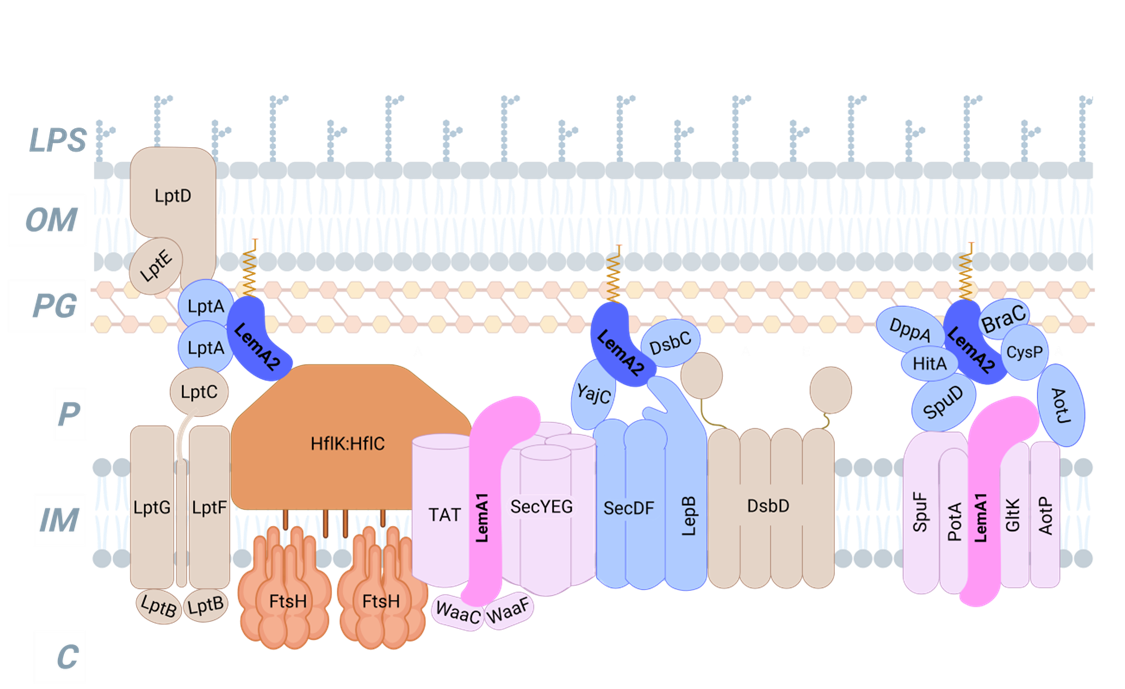

**Extended Data Fig. 18: Protein complex model showing the major components of the Protein translocase systems and transporters found attached to the PaLemAs.** PaLemA1 and PaLemA2 interactome data for the interaction of various protein transport and modification proteins in conjunction with other metabolite and dipeptide transport proteins were visualized as a mega-complex. PaLemA1 is coloured in magenta while PaLemA2 is in blue. Observed protein-protein interaction do not have to be necessarily of direct nature due to the crosslinking procedure. They rather represent the observed overall mega-complex. Abbreviations: LPS = Lipopolysaccharide; OM = Outer membrane; PG = Peptidoglycan; IM = Inner Membrane; C = Cytoplasm. The figure was created with Biorender.

The interactome data suggested a coordination of the BMV-related protein transport via the PaLemA1 and 2. In order to test the influence of both proteins on the OMV/OIMV protein composition, corresponding vesicles were isolated and purified from the wildtype, ΔPa*lemA*1, Pa*lemA*2, and Pa*lemA*1+A2 strains using sucrose gradient centrifugation and subjected to a proteomic analysis (Extended Data Table 8; the raw proteomic data will be deposited in the ProteomeXchange Consortium via the PRIDE (39) partner repository upon acceptance). Comparison of the PaLemA1 and 2 interactomes with the isolated vesicle proteomes revealed that 10 out of 106 proteins of the PaLemA1 interactome (SecY, PhzD1, CysE, PcnB, RpoS, FadE, HflC, PhhB) were also found in the wildtype vesicle proteome (Extended Data Table 8). Similarly, 19 out of 121 proteins of the PaLemA2 interactome (RecA, MexAB, SecCY, CysE, DapB, DnaN, CcoNP, MdoG, SdhC) were found in wildtype OMV/OIMV (Extended Data Table 8). Mutations in both Pa*lemA*s abolished almost completely the presence of these interactome proteins in isolated vesicles (Extended Data Table 8). Interestingly, SecY of the Sec protein export system found in the interactome of both PaLemA1 and 2 was also found in the OMV/OIMVs in a PaLemA-dependent manner. The five most abundant proteins in wildtype BMVs were PA14_26020 (aminopeptidase Lap), PA14_16100 (PET hydrolase/cutinase-like domain-containing protein), PA14_27100 (triacylglycerol lipase LipA), PA14_50290 (flagellin FliC), and PA14_08070 (putative prophage major tail sheath protein GpFl). PA14_26020 (Lap) and PA14_16100 were consistently detected in all knockout strains while PA14_27100 (LipA) was found absent in both the ΔPa*lemA*1 and ΔPa*lemA*2 mutants. PA14_50290 (FliC) was missing in ΔPa*lemA*2 and in the double knockout strain ΔPa*lemA1*+*A2*, while PA14_08070 (GpFI) was found absent in BMVs isolated from all Pa*lemA* mutants. Both PaLemA proteins significantly influence the isolated vesicle protein composition

15. **The PaLemA interactome links vesicle formation to nutrient transport and metabolic adaptation**

Besides proteins required for BMV formation, additional compounds including various metabolites and metals serving as energy source, building block or signal molecule have to traffic the inner membrane. Involved genes have to be regulated accordingly, required proteins have to be synthesized including cofactors, localized, assembled and protected. All these processes require energy. To dedicate appropriate proteins to BMV formation, we found multiple proteins of membrane transport, amino acid metabolism, chemotaxis (Extended Data table 9), energy metabolism (Extended Data table 10), and gene expression with DNA/RNA turnover and stress response (Extended Data table 11) attached to the PaLemA proteins.

**N-containing compound transport (amino acids, polyamines, dipeptides)**

Dipeptides, also in their cyclic form, and polyamines are well known essential carbon- and nitrogen sources, signal molecules and structural components of *P. aeruginosa* (98, 99). Surprisingly, PaLemA2 was found associated with almost the complete set of di/tripeptide transporter substrate binding proteins (SBP) with DppA1, DppA3, DppA4, DppA5, and the ATP-binding protein DppF (100). Their substrate profile in *P. aeruginosa* showed that DppA1 and DppA3 preferentially bind dipeptides, while DppA4 favours tripeptides. The ligand of DppA5 is unknown (101, 102). *P. aeruginosa* can use polyamines as sole carbon and nitrogen source (103). Spermidine regulates gene expression via translational control and riboswitches, acts as antioxidant and is known to regulate biofilm formation via quorum sensing, and to sustain antimicrobial tolerance (104, 105). The periplasmic polyamine spermidine-SBP SpuD and SpuE and the corresponding transporter proteins with the potential ATP-binding proteins PotA and SpuF forming a type III secretion system were found attached to PaLemAs (103, 104). Obviously, this T3SS for polyamines is important for BMV formation and controlled by spermidine (106). Closely interconnected to polyamine transport and catabolism are the processes of arginine metabolism (107, 108). Together they promote c-di-GMP and biofilm formation (109). The arginine/ornithine binding proteins AotJ (PaLemA2) and the subunit AotP (PaLemA1) of the corresponding antiporter system is necessary for the arginine amino acid metabolism and anaerobic arginine fermentation (102, 107). As expected from the presence arginine and polyamine transporters additional amino acid transporters were discovered. The high affinity branched-chain amino acids transport SBP BraC and the corresponding ATP-binding protein BraF were found bound to PaLemA2 (110). BraC was demonstrated to transport Thr, Leu, Ile, Val, Ala and homoserine (102). Several SBP including MetQ and YbeJ predicted to be involved in methionine and polar amino acid transport were found bound PaLemA2.

**Chemotaxis**

Interestingly, corresponding chemotaxis proteins (PctA, PctB, PctC + potential methyl-accepting chemotaxis proteins MCPs) were found attached mainly to PALemA1. PctA was shown to bind most amino acids except for Asp, Glu and Gln while PctB is specific for Arg, Lys, Ala, Met, and Gln (111). PctC binds His, Pro, and GABA (112).  After proteins of amino acids chemotaxis, transport proteins for their successful uptake, also enzymes of the corresponding amino acid metabolism were found attached to the PaLemAs (Extended Data Table 9).

**Amino acid metabolism**

The Serine *O*-acetyltransferase CysE catalysing the first step of Cys biosynthesis as part of the cysteine synthase complex is an attractive drug target for antibiotic development (113-116). Also, proteins involved in degradation of Ser can be found. The serine hydroxymethyltransferase GlyA2 and the aminomethyltransferase GcvT2 are both part of the Gly cleavage system (GCS) for the conversion of glycine into pyruvate (117).

**The 3-isopropylmalate dehydrogenase LeuB catalyses the** irreversible oxidative decarboxylation of **3-isopropylmalate** to 2-oxoisocaproate that is finally converted into Leu by a branched-chain aminotransferase (118, 119). **In contrast, we found also enzymes of branched chain amino acid degradation pathways: The CoA-acylating methylmalonate semialdehyde dehydrogenase MmsA catalyses the conversion of methylmalonate semialdehyde with CoA-SH and NAD^+^ to propionyl-CoA, CO2 and NADH + H^+^ during Val catabolism (120). During Leu and terpene degradation isovaleryl-CoA dehydrogenase** LiuA catalyses the conversion of 3-methylbutanoyl-CoA with an oxidized electron-transfer flavoprotein and one proton to 3-methylbut-2-enoyl-CoA and the reduced electron-transfer flavoprotein (121).

**The bifunctional chorismate mutase/prephenate dehydratase** PheA **is a key enzyme of** Phe biosynthesis via the shikimate pathway underlying strict allosteric control by end product Phe. The **chorismate mutase activity catalyses the conversion of chorismate into prephenate which in turn is converted by the prephenate dehydratase activity into phenylpyruvate, H_2_O and CO_2_ (122). A *pheA* mutant revealed a strong persister phenotype with high antibiotic tolerance (123). The 3‑dehydroquinate dehydratase** AroQ1, catalysing the dehydration of 3-dehydroquinate to 3‑dehydroshikimate and H_2_O, is another key enzyme of the shikimate pathway for aromatic amino acid biosynthesis (124). The shikimate pathway is not present in mammals and represents an important target for antibiotic development. Phe and Tyr catabolism requires the pyridoxal phosphate-dependent aromatic acid transaminase PhhC which transfers the nitrogenous group of an aromatic amino acid to 2-oxoglutarate to form an aromatic oxo acid and Glu (125). Phenylalanine hydroxylase PhhA catalyses the important conversion of Phe into Tyr. Again, the interactome shows also enzymes of aromatic amino acid degradation pathways: **The flavin enzyme tryptophan 2-monooxygenase IaaM catalyses** oxidation of Trp with O_2_ to indole-3-acetamide, CO_2_ and H_2_O during the biosynthesis of the plant hormone auxin (126, 127).

The pyridoxal phosphate-dependent **arginine decarboxylase AdiA catalyses the first step of Arg degradation, the conversion of Arg into agmatine and CO_2_. In several bacteria the protein is the adaptation to acidic conditions and the synthesis of putrescine (128-130) and the relevance of polyamines for *P. aeruginosa* has been already discussed above. Pyridoxine/pyridoxamine phosphate is the essential cofactor of aminotransferases which are essential for amino acid anabolism and catabolism. The crucial flavoenzyme pyridoxine/pyridoxamine 5'-phosphate oxidase PdxH catalyses the conversion of** pyridoxine 5'-phosphate (PNP) or pyridoxamine 5'-phosphate (PMP) into pyridoxal 5'-phosphate (PLP), representing the active coenzyme form (131).

**Other transport proteins**

Surprisingly, only two proteins of ABC sugar transporters (GltF, GltJ) were found bound by PaLemA1. WspA, a homologue of MCPs senses surface contact and activates c-di-GMP signaling (132, 133). A subunit of the thiosulfate ABC transporter (CysP) was found bound to PaLemA2 (102). Interestingly, investigations of *P. aeruginosa* fuels cells showed an increase in power density by a *cysP* mutant indicating the importance of the transporter for coordination of phenazine versus classical electron transport-based energy generation. The ferric iron and magnesium uptake systems HitA, FbpA and MgtE (134) were found bound to PaLemA2 and PaLemA1, respectively. Interestingly, vaccination of mice with recombinant HitA or with *Yersinia pseudotuberculosis* OMVs containing a *P. aeruginosa* PcrV-HitA_T_ (PH) fusion antigen provided protection against subsequent *P. aeruginosa* challenges (134, 135). The periplasmic ferric binding protein FbpA from *Neisseria* and *Haemophilus influenzae* is considered as a naked iron and Fe-chelate carrier (136). A *H. influenzae fbpA* mutant was found deficient in using human transferrin as sole iron source (137). The magnesium transporter MgtE is an important virulence regulator with cytotoxicity-regulating functions (138). MgtE inhibits structural T3SS gene expression by stimulation *rsmYZ* transcription (139, 140). The periplasmic phosphate (Pi)-binding protein Pst was found bound PaLemA1. Its production is highly induced upon phosphate starvation (141). Under these conditions *P. aeruginosa* forms unusual appendages incorporating the PstS protein which are visually distinct from pili and flagella (142). They promote adhesion to intestinal epithelial cells (143). RND multidrug efflux membrane fusion proteins MexAB interacted with PaLemA2 interlacing BMV formation with carbapenem resistance (144). The turgor pressure reactive small-conductance mechanosensitive channel MscK (145) interacted with PaLemA 2, while the YbdG (146) did so with PaLemA1.

**Extended Data Table 9:** **Annotated interaction partners of the PaLemA1 and PaLemA2 proteins involved in** **transport processes, catabolic and biosynthetic pathways.** Accession, locus tag, gene name, description and enrichment factor are given. Detailed protein information was added according to Uniprot database (https://uniprot.org) and the Pseudomonas database (https://pseudomonas.com). Analysis was performed with RStudio software, n = 3. Essential genes after Poulsen *et al*., 2019 and Lee *et al*., 2015 (accession and locus tag in red) only in Poulsen *et al*., 2019 (pink) and only in Lee *et al*., 2015 (green) are labelled.

| Accession | Locus tag | Gene | Description | Interactor of LemA1 (A1) /LemA2 (A2).  Enrichment Factor |
| --- | --- | --- | --- | --- |
| Transport | | | | |

| A0A0H2ZDE6 | *PA14_23010* | *gltK* | ATP-binding component of glucose ABC transporter | A1 24.98 |
| --- | --- | --- | --- | --- |
| A0A0H2Z8S4 | *PA14_46920* | *gltJ* | Permease of glucose ABC transporter | A1 23.41 |
| A0A0H2ZDK3 | *PA14_22990* | *gltF* | **Permease of ABC sugar transporter** | A1 21.49 |
| A0A0H2Z7A1 | *PA14_52790* | *aotJ* | Arginine/ornithine binding protein AotJ | A2 27.31 |
| A0A0H2Z7B3 | *PA14_52750* | *aotP* | Arginine/ornithine transport protein AotP | A1 24.33 |
| Q02DZ3 | *PA14_70860* | ***pstS*** | Phosphate ABC transporter periplasmic phosphate-binding protein **PstS** | A1 21.96 |
| A0A0H2ZEE8 | *PA14_13010* | *metQ* | **Periplasmic methionine ABC transporter substrate-binding lipoprotein** | A2 23.95 |
| A0A0H2ZIR4 | *PA14_72640* | *-* | **Periplasmic (methionine) ABC transporter substrate-binding lipoprotein** | A2 25.18 |
| A0A0H2Z9N7 | *PA14_36200* | *-* | **Periplasmic amino acid ABC transporter substrate-binding protein** | A2 26.35 |
| A0A0H2ZIL8 | *PA14_68070* | *-* | **Periplasmic amino acid ABC transporter substrate-binding protein** | A2 23.58 |
| A0A0H2ZE55 | *PA14_14390* | *-* | **Periplasmic ABC transporter substrate-binding protein, lipoprotein** | A2 24.04 |
| A0A0H2ZDK0 | *PA14_16890* | *-* | Auxiliary component of ABC transporter | A2 22.10 |
| Q02UB7 | *PA14_03920* | *spuD* | **Putrescine-binding periplasmic protein** | A2 27.00 |
| A0A0H2ZL00 | *PA14_03930* | *spuE* | Polyamine transport protein | A2 25,90 |
| A0A0H2ZKB1 | *PA14_03940* | *spuF* | Spermidine/putrescine import ATP-binding protein PotA | A1 24,02 |
| Q02R79 | *PA14_17640* | *potA* | **Spermidine/putrescine import ATP-binding protein PotA** | A1 20,53 |
| A0A0H2ZKK4 | *PA14_07860* | ***potA*** | **Spermidine/putrescine import ATP-binding protein PotA** | A1 24,23 |
| A0A0H2ZGV2 | *PA14_58350* | *dppA1* | Di/tripeptide-binding protein 1 | A2 24,32 |
| A0A0H2ZGN2 | *PA14_58390* | *dppA3* | Di/tripeptide-binding protein 3 | A2 27,56 |
| A0A0H2ZGV7 | *PA14_58420* | *dppA4* | Di/tripeptide-binding protein 4 | A2 27,27 |
| A0A0H2ZI72 | *PA14_70200* | ***dppA5*** | Di/tripeptide-binding protein 5 | A2 22,24 |
| A0A0H2ZH52 | *PA14_58490* | ***dppF*** | **Di/tripeptide transport ATP-binding protein DppF** | A1 27.34 |
| A0A0H2Z7T0 | *PA14_50520* | *braC* | Branched-chain amino acid transport protein BraC | A2 27,56 |
| A0A0H2Z7J0 | *PA14_50550* | *braF* | **Branched-chain amino acid transport protein BraF** | A1 22.32 |
| A0A0H2ZLM2 | *PA14_21175* | - | Putative ABC-type phosphonate transport system (PnhD -like) | A2 26,93 |
| A0A0H2Z953 | *PA14_45110* | *cysP* | Sulfate-binding protein of ABC transporter | A2 25,99 |
| A0A0H2Z6Z0 | *PA14_54540* | *tctC* | **Tripartite tricarboxylate transporter substrate binding protein** | A2 24,61 |
| A0A0H2ZKX6 | *PA14_05530* | *mexA* | RND multidrug efflux membrane fusion protein MexA | A2 26,10 |
| A0A0H2ZLB4 | *PA14_05540* | *mexB* | Efflux pump membrane transporter | A2 25,05 |
| A0A0H2ZHY9 | *PA14_62000* | *hitA* | Ferric iron-binding periplasmic protein HitA | A2 24,96 |
| A0A0H2Z6Q0 | *PA14_53890* | *-* | DUF3649 domain-containing protein, potential iron uptake | A1 19.87 |
| A0A0H2ZIS4 | *PA14_57110* | ***fbpA*** | Iron (III) transport system substrate-binding protein | A2 23.58 |
| A0A0H2Z7D2 | *PA14_52460* | ***mgtE*** | **Magnesium transporter MgtE** | A1 21.63 |
| A0A0H2ZGM3 | *PA14_57110* | *-* | **Small-conductance mechanosensitive channel, osmo-regulation** | A2 22.70 |
| A0A0H2ZIM0 | *PA14_66400* | *mscK* | Putative small-conductance mechanosensitive channel | A2 25,57 |
| A0A0H2ZEI9 | *PA14_19230* | ***ybdG*** | **Mechanosensing system component YbdG** | A1 21.71 |
| **Chemotaxis** |  |  |  |  |
| A0A0H2ZG92 | *PA14_56000* | *pctA* | **Chemotactic transducer PctA,** methyl-accepting chemotaxis, inner membrane protein | A2 22.85 |
| A0A0H2ZGN3 | *PA14_56010* | *pctB* | **Chemotactic transducer PctB** | A2 25.70 |
| A0A0H2ZGG1 | *PA14_55960* | *pctC* | **Chemotactic transducer PctC** | A2 23.66 |
| A0A0H2ZGH1 | *PA14_61300* | *-* | **Putative methyl-accepting chemotaxis protein (MCP)** | A2 21.92 |
| A0A0H2ZKI7 | *PA14_29800* | *pilJ* | Type IV pili methyl-accepting chemotaxis transducer PilJ | A2 23.84 |
| A0A0H2ZC84 | *PA14_29800* | *-* | **Putative chemotaxis transducer** | A2 25.78 |
| **Amino acid metabolism** |  |  |  |  |
| A0A0H2ZF79 | *PA14_14700* | ***cysE*** | **Serine O-acetyltransferase**, first step of cysteine biosynthesis | A1 25.98  A2 21,21 |
| A0A0H2ZK38 | *PA14_05060* | ***-*** | Putative [homoserine O-acetyltransferase](https://www.uniprot.org/uniprotkb/A0A3S4RSA9/entry), first step of methionine biosynthesis | A2 23.51 |
| A0A0H2Z9Q9 | *PA14_41020* | ***adiA*** | **Degradative arginine decarboxylase** | A2 22.94 |
| A0A0H2ZDG4 | *PA14_23790* | ***leuB*** | **3-isopropylmalate dehydrogenase,**  L-leucine biosynthesis | A1 22.50 |
| A0A0H2ZAG9 | *PA14_38440* | ***liuA*** | **Isovaleryl-CoA dehydrogenase,** L-leucine degradation | A2 24.74 |
| A0A0H2ZKJ4 | *PA14_05480* | ***iaaM*** | **Tryptophan 2-monooxygenase,**  auxin biosynthesis | A2 22.58 |
| A0A0H2ZDL6 | *PA14_23280* | ***pheA*** | **Bifunctional chorismate mutase/prephenate dehydratase,** L‑phenylalanine biosynthesis | A1 21.53 |
| Q02I31 | *PA14_53000* | ***phhB*** | **Pterin-4-alpha-carbinolamine dehydratase,** tetrahydrobiopterin biosynthesis | A1 23.75 |
| A0A0H2Z637 | *PA14_53010* | ***phhC*** | Aromatic amino acid transaminase | A1 23.77 |
| A0A0H2ZHW1 | *PA14_64090* | ***aroQ1*** | **3-dehydroquinate dehydratase**, shikimate pathway | A1 22.44 |
| A0A0H2ZAB7 | *PA14_33010* | ***glyA2*** | **Serine hydroxymethyltransferase,** reversible interconversion of Ser + Gly | A1 20.34 |
| A0A0H2Z5W5 | *PA14_54620* | ***mmsA*** | [methylmalonate-semialdehyde dehydrogenase (CoA acylating)](https://www.uniprot.org/uniprotkb/A6VAN0/entry), **valine catabolism** | A1 22.00 |
| A0A0H2ZBK9 | *PA14_33040* | ***gcvT2*** | **Aminomethyltransferase of the** glycine cleavage system (GCS) | A1 21.35 |
| A0A0H2ZES7 | *PA14_17690* | ***-*** | **Putative glutamate synthase, large subunit, archaeal type** | A1 23.67 |
| Q02IL5 | *PA14_50800* | ***pdxH*** | **Pyridoxine/pyridoxamine 5'-phosphate oxidase** | A1 22.26 |

16. **The PaLemA interactome links vesicle formation to cellular energy metabolism**

Biosynthetic processes like BMV formation require energy. Thus, it was not surprising, that the PaLemA coordination platform attracts an almost complete respiratory chain with key components of the corresponding actors of the central metabolism and cofactor formation (147). Major components of the primary proton-pumping and electron donating dehydrogenase NADH dehydrogenase (NuoB, NuoC, NuoE, NuoI bound to PaLemA1) (148, 149), a second primary electron donating dehydrogenase malate:quinone oxidoreductase MqoB (PaLemA1 bound) (150), a third enzyme, the glucose dehydrogenase Gcd (PaLemA1), the major molybdenum-containing subunit of formate dehydrogenase-O (FdnG, PaLemA1 bound) and finally the subunits of succinate dehydrogenase (SdhC, PaLemA1 bound, SdhD, bound to both PaLemA 1 and PaLemA2) were identified in the PaLemA interactome (151). MqoB and FdnG sustain anaerobic respiration while Gcd prefers the coupling to aerobic electron transfer chains. Phenazine of *P. aeruginosa* including pyocyanin (PYO), phenanzine-1-carboxylic acid (PCA) and phenanzine-1-carboxamide, act as electron sink for anaerobic glucose catabolism via extracellular electron transfer (152). In this context the NADH dehydrogenase Nuo acts as phenazine reductase channelling electrons into the ubiquinone pool (153). Not only several Nuo subunits are interacting with PaLemA1, but also the phenazine synthesizing isoenzymes of isochorismatase (PhzD1/D2) are interaction partners of PaLemA1 (154). Furthermore, periplasmic electron transferring proteins including azurin (Azu) and the iron-sulphur protein FixG were found PaLemA2-attached. The small periplasmic blue copper protein Azu facilitates the anaerobic respiratory chains of denitrification (155). Furthermore, a 28 amino acid long peptide of Azu is successfully used in cancer therapy (156, 157). FixG proteins belong to a protein family characterized by the binding of a [4Fe-4S] cluster mediating electron transfer (158). Finally, the subunits CcoP and CcoN of the oxygen-dependent terminal oxidase cbb_3_-type cytochrome c oxidase were found as interaction partners of PaLemA1. Correspondingly, the cytochrome c-type biogenesis protein CycH was found bound to both PaLemA 1 and PaLemA2 (159).

**Cofactor formation**

The membrane protein CycH is an apocytochrome chaperone during c-type cytochrome formation. Similarly, the putative uroporphyrin-III C-methyltransferase HemX was found attached to both PaLemA 1 and PaLemA2 (160), which is known to be is encoded in the *hemCDX* operon in *E. coli* (161). This potential enzyme catalyses a reaction at the branch point of tetrapyrrole biosynthesis towards vitamin B_12_ and siroheme. Other PaLemA-attached cofactor biosynthesis enzymes were **C-methyltransferase (UbiE) of ubiquinone biosynthesis and thiazole synthase ThiG involved in thiamine biosynthesis.** *P. aeruginosa* has been shown to use mainly ubiquinone-9 also during anaerobic metabolism (162). During ubiquinone biosynthesis the **C-methyltransferase UbiE** methylates 6-methoxy-2-nonaprenyl-1,4-benzoquinol using S-adenosyl-L-methionine to form 6-methoxy-3-methyl-2-octaprenyl-1,4-benzoquinol, S-adenosyl-L-homocysteine and H^+^ (163, 164). Thiazole synthase ThiG catalyses the crucial 3 substrate condensation using 1-deoxy-D-xylulose-5-phosphate, glycine-derived 2-iminoacetate and ThiS-thiocarboxylate to form a stable thiazole tautomer which gets converted into the aromatic thiazole phosphate during the next step (165).The PaLemA1-bound **pterin-4-alpha-carbinolamine dehydratase (PhhB) is relevant for** tetrahydrobiopterin biosynthesis, e.g. important for Tyr biosynthesis.

**Extended Data Table 10:** **Complete** **annotated interaction partners of the PaLemA1 and PaLemA2 proteins involved in energy metabolism and cofactor biosynthesis.** Accession No., locus tag, gene name, description and enrichment factor are given. Detailed protein information was added according to Uniprot database (https://uniprot.org) and the Pseudomonas database (https://pseudomonas.com). Analysis was performed with RStudio software, n = 3. Essential genes after Poulsen *et al*., 2019 and Lee *et al*., 2015 (accession and locus tag in red), only in Poulsen *et al*., 2019 (pink) and only in Lee *et al*., 2015 (green) are labelled.

| Accession | Locus tag | Gene | Description | Interactor of LemA1 (A1) /LemA2 (A2).  Enrichment Factor |
| --- | --- | --- | --- | --- |
| Energy metabolism | | | | |

| Q02ND0 | *PA14_30010* | *nuoB* | NADH-quinone oxidoreductase subunit B | A1 25.24 |
| --- | --- | --- | --- | --- |
| Q02ND1 | *PA14_29990* | *nuoC* | NADH-quinone oxidoreductase subunit C/D | A1 26.93 |
| A0A0H2ZC94 | *PA14_29980* | *nuoE* | NADH dehydrogenase I subunit E | A1 24.07 |
| Q02ND6 | *PA14_29920* | *nuoI* | NADH-quinone oxidoreductase subunit I | A1 24.66 |
| A0A0H2ZHI8 | *PA14_61400* | *mqoB* | Malate:quinone oxidoreductase | A1 25.85 |
| A0A0H2ZB70 | *PA14_34970* | *gcd* | Glucose dehydrogenase | A2 24.33 |
| A0A0H2ZL50 | *PA14_63605* | *fdnG* | Formate dehydrogenase-O major subunit | A2 21.62 |
| A0A0H2Z8W2 | *PA14_44060* | *sdhC* | Succinate dehydrogenase cytochrome *b*_556_ subunit | A2 22.78 |
| A0A0H2Z9F0 | *PA14_44050* | *sdhD* | Succinate dehydrogenase hydrophobic membrane anchor | A1 23.27  A2 21.02 |
| A0A0H2ZI65 | *PA14_65000* | *azu* | Azurin | A2 24.59 |
| A0A0H2Z8I7 | *PA14_45280* | *cycH* | Cytochrome c-type biogenesis protein | A1 23.39  A2 23.87 |
| A0A0H2Z954 | *PA14_44400* | *ccoP* | Cbb_3-_type cytochrome c oxidase subunit | A2 25.86 |
| A0A0H2Z9C8 | *PA14_44370* | *ccoN* | Cytochrome-c oxidase | A2 21,62 |
| A0A0H2ZG66 | *PA14_10540* | *fixG* | Respiratory iron-sulphur cluster-binding protein | A2 25.51 |
| A0A0H2ZHI5 | *PA14_61140* | *-* | [2Fe-2S ferredoxin-type domain-containing protein](https://www.uniprot.org/uniprotkb/Q9HVH1/entry) | A2 20.98 |
| Q02DF0 | *PA14_73290* | *atpF* | ATP synthase subunit b | A2 24,61 |
| Q02DE8 | *PA14_73310* | *atpB* | ATP synthase subunit a | A2 23,13 |
| A0A0H2ZDQ9 | *PA14_22910* | *edd* | Phosphogluconate dehydratase (Entner-Doudoroff pathway) | A1 24.36 |
| Q02KR1 | *PA14_41670* | ***ppsA*** | **Phosphoenolpyruvate synthase,** (gluconeogénesis) | A2 23.73 |
| A0A0H2Z7C9 | *PA14_53940* | ***prpB*** | 2-methylisocitrate lyase **(**methylcitrate cycle, catabolism of propanoate) | A1 21.33 |
| A0A0H2ZKF2 | *PA14_01730* | ***ppk2*** | **ADP/GDP-polyphosphate phosphotransferase** | A1 22.82 |
| A0A0H2ZFK1 | *PA14_11690* | ***ppa*** | **Inorganic pyrophosphatase PpA** | A1 23.45 |
| *A0A0H2ZF69* | *PA14_09450* | *phzD1* | Phenazine biosynthesis protein PhzD (Pyocyanin formation) | A1 23.23 |
| *A0A0H2ZKU3* | *PA14_39925* | *phzD2* | Phenazine biosynthesis protein PhzD (Pyocyanin formation) | A1 23.23 |
| Q02EV4 | *PA14_66900* | ***ubiE*** | **Ubiquinone biosynthesis C-methyltransferase** | A1 22.53 |
| Q02U32 | *PA14_04980* | ***thiG*** | **Thiazole synthase,** thiamine biosynthesis | A1 22.07 |
| A0A0H2ZJ90 | *PA14_69430* | *hemX* | Uroporphyrin-III C-methyltransferase | A1 23.81  A2 23.22 |

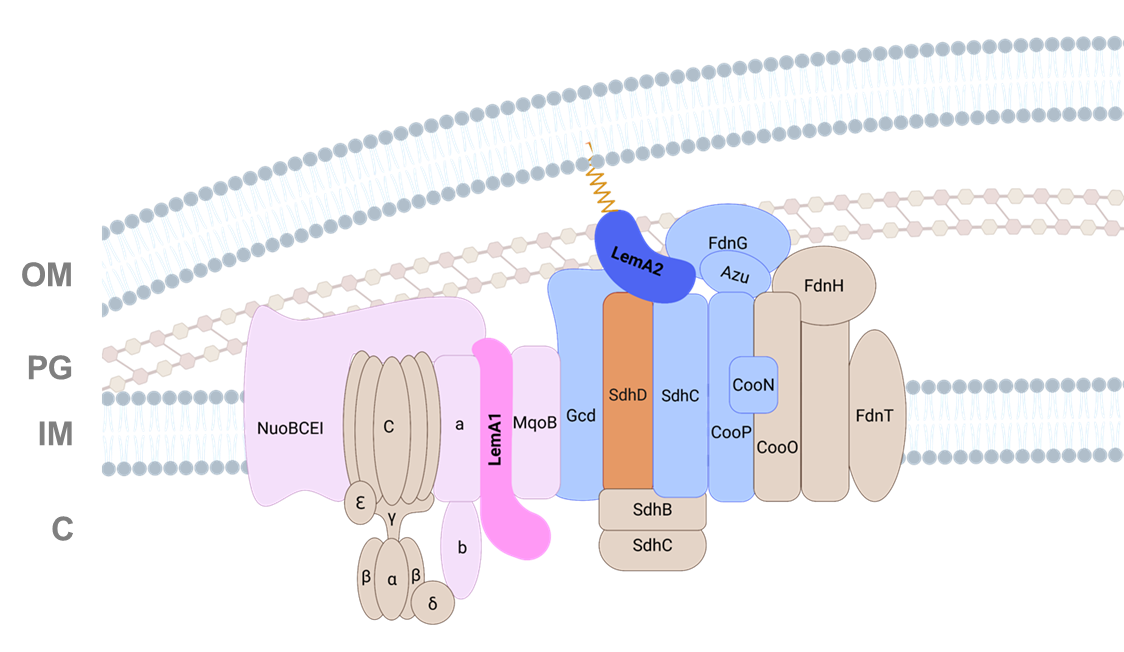

**Extended Data Fig. 19: Protein complex model showing the major components of the energy metabolism found attached to the PaLemAs.** PaLemA1 and PaLemA2 interactome data for the interaction of various proteins of the energy metabolism were visualized as a mega-complex. PaLemA1 is coloured in magenta while PaLemA2 is in blue. Observed protein-protein interaction do not have to be necessarily of direct nature due to the crosslinking procedure. They rather represent the observed overall mega-complex. Abbreviations: OM = Outer membrane; PG = Peptidoglycan; IM = Inner Membrane; C = Cytoplasm. Abbreviations: OM = Outer membrane; PG = Peptidoglycan; IM = Inner Membrane; C = Cytoplasm. The figure was created with Biorender.

17. **The PaLemA interactome links vesicle formation to gene regulation and stress response**

Finally, central components of the gene expression process, including DNA/RNA metabolism, replication, transcription, translation, degradation and corresponding stress responses were found PaLemA-bound.

**DNA/RNA metabolism**

During purine metabolism glutamine-hydrolyzing GMP synthase GuaA catalyses the crucial step towards GTP, the conversion of xanthine monophosphate (XMP) with Glu, ATP and H_2_O to form GMP, AMP, PP_i_ and 2H^+^ (166, 167). UMP biosynthesis via a salvage pathway relies on the **uracil phosphoribosyltransferase Upp which catalyses the conversion of** uracil and 5-phospho-alpha-D-ribose 1-diphosphate to UMP and PP_i_ (168). *P. aeruginosa* requires for growth on adenosine the quorum-sensing controlled nucleoside hydrolase Nuh (169, 170). The enzyme hydrolyzes the N-glycosidic bond of nucleosides with the release of the free base (purine or pyrimidine) and the sugar (ribose or deoxyribose) (169).

**Replication**

During DNA replication, the homodimeric β-sliding clamp, composed of two subunits DnaN forming a doughnut shaped ring with a central channel, interacts preferentially with the negatively charged phosphate backbone of the DNA, allowing the clamp to slide along the DNA sustaining vital DNA replication, which is clearly distinct of that of other bacteria including *E. coli* (171).

**Transcription**

During transcription the dynamic modulator NusG (Spt5 in eukaryotes/Archaea, attached to PaLemA2) regulates throughout all cellular life forms RNA polymerase pausing to allow for RNA folding. It also interacts with ribosomes and terminations factors to control the transcriptional process (172, 173). RapA (HepA), associated with PaLemA2, is an ATPase that removes the post-termination complex of RNA polymerase from the DNA strand to finally terminate transcription (174). RpoS (attached to PaLemA1) is an alternative sigma factor known to be involved in the general stress and stationary phase response including iron-related radical oxygen species in many bacteria (175). In *P. aeruginosa* RpoS is also involved in the control of pyocyanin production, biofilm formation via quorum sensing by the regulation of the regulatory proteins LasR and RhlR (176, 177). In this context RpoS and LasR coordinate the production of phenazine (178) of which the two isoenzymes of isochorismatase PhzD1/2 are interacting with PaLemA1. Pyocyanin is involved in the anaerobic extracellular electron transport. In agreement the major anaerobic regulator Anr, found attached to PaLemA1, coordinates the expression of genes of denitrification, arginine fermentation, and anaerobic survival (179-181). Mutation of the gene for the catabolite repression protein Crc or corresponding regulation of the Crc-antagonist small RNA CrcZ allows for biofilm expansion by the induced production of the biofilm matric components Pel exopolysaccharides and CupBC fimbriae. The mediator of this molecular convergence point for nutrient – and quorum sensing inputs Crc was found bound to PaLemA2 (182). The copper uptake regulator CueR was discovered originally as target of the quorum-sensing regulator LasR (183). A co-regulation of the copper uptake systems COPA1/2 and of azurin by CueR with RpoS and Anr was observed before (184). The histidine kinase PhoQ is part of the essential PhoPQ two component regulatory system which functions as sensor for environmental conditions including low pH, low Mg^2+^ concentrations, and contact with epithelium cells and mediates cellular responses in *P. aeruginosa* like virulence, biofilm formation and resistance against aminoglycosides, tetracycline, antimicrobial peptides (CAMPs) and polymyxin B (185-192). The poly(A) polymerase PcnB polyadenylates RNA molecules at their 3´ends and plays a central role for mRNA stability via the control of RNA degradation (193, 194). Furthermore, PcnB controls the copy number of plasmids. The enzyme modulates virulence and resistance in many pathogenic bacteria and is controlled by Tyr phosphorylation (195-197). **The ATP-dependent** DEAD-box **RNA helicase** RhlE2 binds and unwinds double-stranded RNA. RhlE1 is solely required for growth in the cold. In contrast RhlE2 is a global post-transcriptional regulator controlling the level of hundreds of transcripts. RhlE2 carries an unique C-terminal extension supporting RNA unwinding activity of the N-terminal domain and a RNA-dependent interaction with RNase E and components of the cellular RNA degradation apparatus (198). Due to its essential cellular function RhlE2 represents an attractive drug target.

**Translation**

The non-essential 50S ribosomal protein L29 RpmC stabilizes the structure of 23S rRNA, is part of the polypeptide exit tunnel, and makes direct contact to the ribosome-associated chaperone trigger factor. Ribosomal protein L33 RpmG also binds to 23S rRNA and can be post-translationally methylated (199, 200). GTPase EngA (YihI) is a double Era-like (Der) GTPase that plays a role in 50S ribosome subunit maturation via ensuring stable 23S rRNA integration (201). The EngA protein is subject to Lys polyphosphate modification which allows for efficient biofilm formation and pyoverdine and pyocyanin production. Corresponding **ADP/GDP-polyphosphate phosphotransferase** Ppk2 is also a PaLemA protein bound protein (Extended Data Tables 11). **The ribosome maturation factor RimP is crucial for 30S subunit assembly via 16S rRNA processing and controlling S5 and S12 protein-RNA interaction with the 5´end of the RNA (202-204).** The energy-dependent translational throttle protein EttA binds to the ribosome E-site and controls the progression of the 70S ribosomal initiation complex into elongation phase in response to ATP/ADP ratios. Dipeptide formation is stimulated in the presence of ATP and inhibited by ADP, like in the stationary phase (205-207). Ribosome recycling factor RRF, in *P. aeruginosa* encoded by the *frr* gene, is needed for release of 70S ribosomes from the mRNA upon reaching the stop codon (208, 209). Mutation of *frr* intervenes in the dissociation of ribosomal complexes, thus preventing ribosome stalling and enhancing ribosome recycling (210). Several aminoacyl-tRNA synthetases required for the ligation of amino acids to their cognate tRNAs were found bound to PaLemA proteins (Extended Data Table 11), including the n**atural miss-charging aspartate-tRNA (Asp/Asn) ligase AspS (211-213), the α-subunit GlyQ of glycine-tRNA ligase, methionine-tRNA ligase MetG, glutamate-tRNA ligase GltX and tryptophan-tRNA ligase TrpS. Several of the enzymes were used for the development of antibiotic targets (214, 215). Besides its role in protein biosynthesis, glutamyl-tRNA synthetase GltX catalyses the first step of tetrapyrrole biosynthesis, the formation of Glu-tRNA^Glu^, which gets subsequently converted into 5-aminolevulinic acid, the general precursor molecule of all tetrapyrroles in *P. aeruginosa* (216).** The flavoprotein **tRNA-uridine-5-carboxymethyl-aminomethyl modification enzyme MnmG** (GidA) is involved in the addition of a carboxymethylaminomethyl (cmnm) group at the wobble position uracil (U34) of certain tRNAs, forming tRNA-cmnm_5_s_2_U34 (217, 218). MnmG is also called glucose-inhibited division protein A **(GidA) since it** **promotes cell division via control of the expression of the division septum component YmgF (219).**

**Stress response**

Interestingly, a manifold of proteins reactive to multiple cellular stresses interacted with the PaLemA proteins. The ubiquitous catalase KatA (220), known to cope with hydrogen peroxide stress, was found attached to PaLemA1. Peroxide stress-induced DNA damage leads to an activation of the SOS response (221). We found RecA, a protein central to DNA repair and recombination after genotoxic damage (222), particularly during the SOS response to interact with PaLemA2. Importantly, to prevent damaged DNA propagation, RecA-activated SOS response induces arrest of cell division, maintaining the functions of the divisome proteins via interaction with FtsZ (223). SOS-response and cell cytokinesis have been shown to modulate BMV formation(7, 224-226). In fact, BMVs have been observed to originate at the septum during cell division and contain specific DNA fragments (226). The close interaction with proteins of the cell division machinery might explain its observed interaction with PaLemA2. Additionally, iron availability was shown to impact oxygen stress and to effect BMV formation. BMVs were proven to participate in iron acquisition and delivery to neighbouring cells (13, 227). Moreover, iron acquisition is intertwined with cell cycle control and proper cell division (228). Hence, the importance of having the iron-sequester bacterioferritin BfrA protein attached to PaLemA1, which can thereby modulate iron homeostasis. CspA and CapB are the ma**jor cold shock proteins of bacteria involved in the** cold-shock acclimatization by ensuring rRNA transcription (229, 230), preserving RNA stability by prevention of secondary structure formation (231), stimulation of translation in the cold to prevent ribosome stalling at low temperatures (232), maintaining membrane fluidity, preserving enzyme integrity and ultimately protecting DNA (230). **In *E. coli*** the *fxsA* **gene encodes the phage T7 F exclusion suppressor and is controlled by a heat shock response (233, 234). In *Pseudomonas pergamininesis* the membrane protein FxsA is involved in acid-tolerance (235).** The γ-glutamyltranspeptidase **Ggt** is a heterodimeric enzyme that is autocatalytically generated from **an in**active precursor to form the active heterodimeric mature enzyme. The enzyme catalyses the hydrolysis of gamma-glutamyl bonds in gamma-glutamyl compounds such as the antioxidant glutathione. **Thus, Ggt is also a glutathione hydrolase involved in glutathione recycling (236, 237). Glutathione biosynthesis plays an important role during multiple stress responses, virulence, quorum sensing and biofilm formation (238-241). The periplasmic stress protein OsmY is** significantly upregulated in response to hypertonicity and in the stationary phase by RpoS (242, 243)**. The presence of** OsmY helps to overcome plasmolysis-induced structural changes and shortens the time to growth resumption after osmotic upshift. Prediction for its exact molecular function range from chaperone function to physical stabilization of the periplasmic space during inward shrinking (244). The **glucan biosynthesis protein G OpgG is important for the low osmolarity-induced biosynthesis of periplasmic glucans (245).** In *P. aeruginosa* PA14 the *opgGH* locus mediates the biosynthesis of β-1,2-linked linear glucosyl polymers with a few β-1,6 glucosyl residues and succinyl residues attached to this glucose backbone (246). **The** **cyclophilin type peptidyl-prolyl cis-trans isomerase PpiA is a periplasmic, slow folding enzyme catalysing** the proline imidic peptide bond *cis*-*trans* isomerization in proteins (247). In *E. coli* it is involved in biofilm formation and swimming motility (248).

**Protein degradation**

Several proteases including three zinc metallopeptidases including the **M18 family aminopeptidase ApeB**, one cysteine protease and the endopeptidase La (Lon) were found attached to the PaLemA proteins (Extended Table 11). The aspartyl amino peptidase ApeB catalyses the removal of Asp and Glu from the N-terminus of peptides. The ATP-dependent cytoplasmic protease Lon is crucial for protein quality and cellular function control in organisms across all domains of life. In *P. aeruginosa*, *lon* mutants revealed multiple phenotypic defects in virulence, motility, biofilm formation and antibiotic tolerance (249-251). Interestingly, Lon is degrading the cell division inhibitor protein SulA to ensure proper cell division (252).

**Host response**

Finally, several proteins of *P. aeruginosa* involved in the response to host defence systems were bound by the PaLemAs. A typical example is lysozyme which is cleaving the β-1,4-glycosidic bonds between N-acetylmuramic acid and N-acetyl-D-glucosamine residues of peptidoglycan, leading to bacterial lysis. The lipoprotein MliC (membrane bound lysozyme inhibitor of c-type lysozyme) of *P. aeruginosa* possess lysozyme inhibitory activity (253, 254). The crystal structure of the *P. aeruginosa* MliC protein in complex with lysozyme showed the insertion of an invariant loop of the MiliC protein into the active site cleft of lysozyme (254). The biofilm matrix associated Ecotin is a strong inhibitor of the bactericidal enzyme neutrophil elastase which is produced by the host immune system during *P. aeruginosa* infections (255, 256). Moreover, the human serine proteases MASP-1 and MASP-2, the two key activator proteases of the complement lectin pathway are inhibited by ecotin (257). Antimicrobial peptides are small, charged molecules with antimicrobial activity (258, 259). The Mig-14-like protein of *P. aeruginosa* is required for recognition of the CAMPs colistin and Novispirin G10 (260).

18. **Control of BMV-formation related gene expression by PaLemA1 and PaLemA2 protein interaction partners Anr, Crc and RpoS**

To test the influence of PaLemA protein-bound transcription factors on the gene expression during BMV formation in *P. aeruginosa* the transcriptomes of ΔPa*lemA*1, ΔPa*lemA*2 and ΔPa*lemA*1+A2 were compared to wildtype gene expression (Extended Data Tables 12-14). To perform the transcriptome analyses, bacterial cultures of the aforementioned strains were prepared in LB broth and samples were taken during the mid-exponential phase of growth. The RNA was isolated, processed and sequenced as described in the methods section, and the differential gene expression between samples was obtained using *edgeR*. Surprisingly, in the ΔPa*lemA*1 mutant multiple genes for transporters (*dppA1*, *dppA3*, *dppA4*, *dppC*, *dppF*, *gltF*, *gltK*, *braCF*), the glycine cleavage system (*gcvP2H2*), phenylalanine hydroxylase (*phhA*), all attached to PaLemAs were found induced (Extended Data Tables 12-14). Additional genes for transporter of ribose (*rbsABCR*), glucose (*oprB*, *gltB*), ribokinase (*rbsK*), branched chain amino acids (*bkdAB*), aromatic amino acids (*aroP2*) iron (fbpA), heme (PhuR) behaved identical. Moreover, several genes for enzyme including dihydrolipoamide dehydrogenase (*lpdV*), fumarylacetoacetase (*fahA*), esterase (estA), D-amino acid dehydrogenase (*dadA*), alanine racemase (*dadX*), glycerol kinase (*glpK*) and the corresponding regulator (*glyR*) showed increased transcription in the ΔPa*lemA*1 mutant. All listed genes (labelled with yellow background in Extended Data Tables 12-14 belong to the catabolite regulator Crc regulon (261). Crc was found attached to PaLemA2 as part of the PaLemA coordinated complex. Obviously, deletion of the gene for PaLemA1 restructures the complex and probably leads to the release of Crc from the complex and the subsequent observed gene regulation. Interestingly, the complete *pqsABCDE* operon (light red background) encoding proteins involved in the quinolone formation required for quorum sensing and biofilm formation were also found induced (262). The PaLemA1 attached alternative sigma factor RpoS is involved in these processes (176). Repressed in the ΔPa*lemA*1 mutant were multiple genes of the type III secretion system (*pscNO*, *pcr12DHV*, *exoUY*, *popBDN* with blue background) which is also under the control of Crc (263). Furthermore, several genes of the energy metabolism (*atpDEH*, *fbr*, *aceEF*, *cyoABCE*) were found repressed (brown background). The pyruvate responsive regulator GavR represses most genes of the type III secretion system and the *aceEF* operon (264). The ΔPa*lemA*2 strain revealed the strong induction of around 30 genes of the *P. aeruginosa* Anr regulon (i.e. *arcABD*, *adhA*, *ccoDN*, *oprG*, *hemN*, *nirS*, *dnr*, *uspA,* *narXL*, Extended Data Tables 12-14, pink background) (179, 265). Anr was found before attached to PaLemA1 as part of the PaLemA-coordinated mega-complex. Again, all genes of the quorum sensing operon (*pqsABCDE,* light red background) were again found induced (Extended Data Tables 12-14). Out of approximately 160 repressed genes in the ΔPa*lemA*2 mutant 30 genes encoded transcriptional regulators (i.e. *nusG*, *rpoH*, *mvaT*, *chpD*, *phoB*, *prtR*, *wspF*, *ntrB*, *dsmR*, *feaR*, *cueR*, brown background). Similarly, 16 genes of replication (i.e. *repC*, *smpB*, *soj*, *dnaQ*, *dnaG*, *alkA*) and translation (i.e. *rrnF*, cat, *rpmB,* grey background). The transcriptome of the double mutant ΔPa*lemA*1 + 2 mainly represented a mixture of the outlined transcriptomes of the single mutants. Only the repression of the type II secretion system observed for the ΔPa*lemA*1 mutant was missing. However, Obviously, PaLemA mediated protein interactions to transcriptional regulators mediate BMV formation-related gene regulation.

**Extended Data Table 11:** **Annotated interaction partners of the PaLemA1 and PaLemA2 proteins involved in** **DNA metabolism, gene regulation, protein biosynthesis, stress and host responses.** Accession No., locus tag, gene name, description and enrichment factor are given. Detailed protein information was added according to Uniprot database (https://uniprot.org) and the Pseudomonas database (https://pseudomonas.com). Analysis was performed with RStudio software, n = 3. Essential genes after Poulsen *et al*., 2019 and Lee *et al*., 2015 (accession and locus tag in red), only in Poulsen *et al*., 2019 (pink) and only in Lee *et al*., 2015 (green) are labelled.

| Accession | Locus tag | Gene | Description | Interactor of LemA1 (A1) /LemA2 (A2).  Enrichment Factor |
| --- | --- | --- | --- | --- |
| DNA metabolism, gene regulation, protein biosynthesis and stress response | | | | |

| A0A0H2ZIW1 | *PA14_00020* | ***dnaN*** | **Beta sliding clamp, DNA replication** | A2 23,12 |
| --- | --- | --- | --- | --- |
| Q02R89 | *PA14_17530* | ***recA*** | Recombinase A | A2 24,00 |
| Q02RS2 | *PA14_15340* | *guaA* | GMP synthase, purine metabolism | A2 22,20 |
| Q02G28 | PA14_61470 | *upp* | **Uracil phosphoribosyltransferase,** pyrimidine metabolism | A2 21,97 |
| A0A0H2Z6T8 | *PA14_54410* | *mucB* | Negative regulator for alginate biosynthesis MucB | A2 23,96 |
| A0A0H2Z5X2 | *PA14_54430* | *algU* | RNA polymerase sigma factor AlgU | A1 23,61  A2 21,62 |
| A0A0H2ZGS0 | *PA14_57760* | *algW* | Periplasmic serine endoprotease, DegP-like, AlgW | A1 21,44  A2 24,50 |
| A0A0H2ZEU1 | *PA14_17480* | *rpoS* | RNA polymerase sigma factor RpoS | A1 24,89 |
| [A0A0H2ZCQ4](https://www.uniprot.org/uniref/UniRef100_A0A0H2ZCQ4) | *PA14_21230* | *hepA* | RNA polymerase-associated protein RapA | A1 24,06 |
| A0A0H2Z8S5 | *PA14_44490* | *anr* | Transcriptional regulator Anr (anaerobic gene regulation) | A1 20,71 |
| A0A0H2ZI82 | *PA14_70390* | *crc* | Global metabolic regulator (catabolite repression control, Crc) | A2 24,00 |
| A0A0H2Z8C1 | *PA14_49170* | *phoQ* | **Histidine kinase PhoQ of two component PhoPQ system, Mg^2+^ deficiency, acidic pH, CAMPs** | A1 21.74 |
| A0A0H2ZKB4 | *PA14_04010* | *-* | **Transcriptional regulator** of **unknown function** | A2 24,64 |
| A0A0H2ZF93 | *PA14_08710* | *nusG* | Transcription termination/ antitermination protein NusG | A2 25,60 |
| A0A0H2ZHV5 | *PA14_62560* | ***pcnB*** | **Poly(A) polymerase I** | A1 21.49 |
| A0A0H2ZJ79 | *PA14_01760* | *nuh* | Nonspecific ribonucleoside hydrolase | A2 24.25 |
| Q02RV3 | *PA14_14930* | *engA/*  *yihI* | **Der-type GTPase,** late steps of [ribosome biogenesis](https://www.uniprot.org/keywords/KW-0690) | A1 22.68 |
| Q02T72 | *PA14_08930* | ***rpmC*** | **Large ribosomal subunit protein uL29** | A1 23.68 |
| A0A0H2ZJ31 | *PA14_70180* | ***rpmG*** | **Large ribosomal subunit protein bL33** | A1 21.38 |
| Q02RC5 | *PA14_17100* | *frr* | Ribosome-recycling factor RRF | A2 24,29 |
| A0A0H2ZK67 | *PA14_05560* | *rhlE2* | **ATP-dependent** DEAD-box **RNA helicase** involved in ribosome assembly | A1 23.97 |
| Q02FS6 | *PA14_62780* | ***rimP*** | **Ribosome maturation factor RimP** | A1 20.38 |
| A0A0H2ZHF4 | *PA14_60800* | *ettA* | Energy-dependent translational throttle protein EttA | A1 21.04 |
| Q02IC5 | *PA14_51820* | ***aspS*** | **Natural miss-charging Aspartate--tRNA(Asp/Asn) ligase AspS** | A2 21.30 |
| Q02V72 | *PA14_00100* | ***glyQ*** | **Glycine--tRNA ligase alpha subunit** | A1 23.35 |
| Q02QX1 | *PA14_19050* | ***metG*** | **Methionine--tRNA ligase** | A1 19.77 |
| Q02PU7 | *PA14_23560* | ***gltX*** | **Glutamate--tRNA ligase** | A1 22.99 |
| A0A0H2ZGQ0 | *PA14_57670* | ***trpS*** | **Tryptophan--tRNA ligase** | A1 22.76 |
| Q02DE3 | *PA14_73370* | ***mnmG/*** *gidA* | **tRNA uridine 5-carboxymethyl-aminomethyl modification enzyme,**  glucose-inhibited division protein A | A1 22.98 |
| A0A0H2ZDX1 | *PA14_21760* | *capB* | Cold acclimation protein B | A2 21,63 |
| A0A0H2Z7Q7 | *PA14_51840* | *cspA* | **Ma****jor cold shock protein CspA** | A1 22.69 |
| A0A0H2ZGF2 | *PA14_57030* | *fxsA* | **Phage T7 F exclusion suppressor, acid and heat resistance** | A1 23.01 |
| A0A0H2ZGK4 | *PA14_09150* | *katA* | Catalase | A1 24,02 |
| A0A0H2Z8J7 | PA14_46960 | ***ggt*** | **Glutathione hydrolase proenzyme** | A2 25.33 |
| Q02ET9 | PA14_67090 | ***opgG*** | **Gl****ucans biosynthesis protein G,**  osmoregulated periplasmic glucans | A2 26.51 |
| 0A0H2ZHN3 | *PA14_62690* | ***osmY*** | **Os****mo-induced periplasmic stress protein binding to inner and outer membrane** | A2 23.70 |
| A0A0H2ZDH5 | *PA14_22450* | ***ppiA*** | **Pe****ptidyl-prolyl cis-trans isomerase PpiA, cyclophilin type** | A2 23.29 |
| *A0A0H2ZG72* | *PA14_09160* | *bfrA* | Bacterioferritin | A1 23.90 |
| A0A0H2ZAC4 | *PA14_38080* | *-* | **Putative cysteine protease** | A2 23.48 |
| A0A0H2ZL01 | *PA14_04890* | *-* | **Putative zinc protease** | A2 23.90 |
| 0A0H2ZGS5 | *PA14_62420* | *-* | **Putative zinc metallopeptidase,** transmembrane anchor | A2 21.68 |
| Q02Q78 | *PA14_21990* | ***apeB*** | **M18 family aminopeptidase 2, zinc metallopeptidase** | A2 20.85 |
| A0A0H2ZH67 | *PA14_60550* | *lon* | **Endopeptidase La (Lon)** | A1 20.46 |
| **Host response** |  |  |  |  |
| Q02NQ8 | *PA14_28450* | *eco* | Ecotin, inhibitor of human serine proteases MASP-1 and MASP-2, the two key activator proteases of the complement lectin pathway | A2 22.50 |
| A0A0H2Z780 | *PA14_53040* | ***mliC*** | Membrane-bound lysozyme inhibitor of C-type lysozyme | A2 25.53 |
| A0A0H2ZI96 | *PA14_66150* | ***mig14*** | Mig-14-like protein involved in antimicrobial peptide recognition | A1 22.03 |

19. **The alternative sigma factor AlgU controls Pa*lemA2* expression in response to envelope stress and growth phase**

The interactomics studies showed an interaction of both PaLemA1 and PaLemA2 with AlgU and AlgW, while MucB is solely interacting with PaLemA2. These proteins are all compounds of the regulatory cascade for the activation of the extracytoplasmic function (ECF) sigma factor AlgU. AlgU (also named AlgT or σ^22^) is widely conserved among bacteria, typically inducing the expression of genes coping with tolerance to osmotic, oxidative, thermal and cell envelope-related stresses (266). Moreover, it is important to activate the transcription of genes involved in the synthesis and export of alginate, an exopolysaccharide that promotes the adhesion to abiotic surfaces and the formation of mucoid biofilms (267). The regulation of AlgU activity is very complex. The cytoplasmic ECF sigma factor AlgU in its inactive form is sequestered by the transmembrane anti-sigma factor MucA which in turn bound and stabilized by the periplasmic MucB. AlgU activation is mediated via proteolysis of its inner-membrane anchored anti-sigma factor MucA part in response to cell envelope stress. This activation of AlgU occurs when envelope stress activates MucE which in turn triggers the AlgW protease. AlgW in concert with the MucP protease cleaves and inactivates the anti-sigma factor MucA which leads to the release of the sigma factor AlgU to initiate the transcription of its regulon. Additionally, a second activation mechanism is triggered by the inactivation of MucB part. MucB interacts with the anti-sigma factor MucA and restricts AlgW protease access to the MucA cleavage site. Under cell envelope stress conditions, the MucB undergoes a conformational change to release MucA. As a result, the frequency of MucA cleavage and sigma factor AlgU release is a function of factor MucB removal and protease AlgW activation (266).

After observing a clear effect of PaLemA1 and PaLemA2 overproduction in vesiculation, we sought to identify the conditions where both genes are expressed under standard laboratory conditions. Taking into account our results of interactomics, the activation of AlgU could be a key feature in the regulation of the transcription of Pa*lemA1* or/and Pa*lemA2*. Furthermore, after inspecting the promoter region of Pa*lemA*2 (i.e. 300 bp upstream of the start codon), we found a putative AlgU binding site (268) located 60 nucleotides upstream of its start codon (Extended Data Fig. 20).

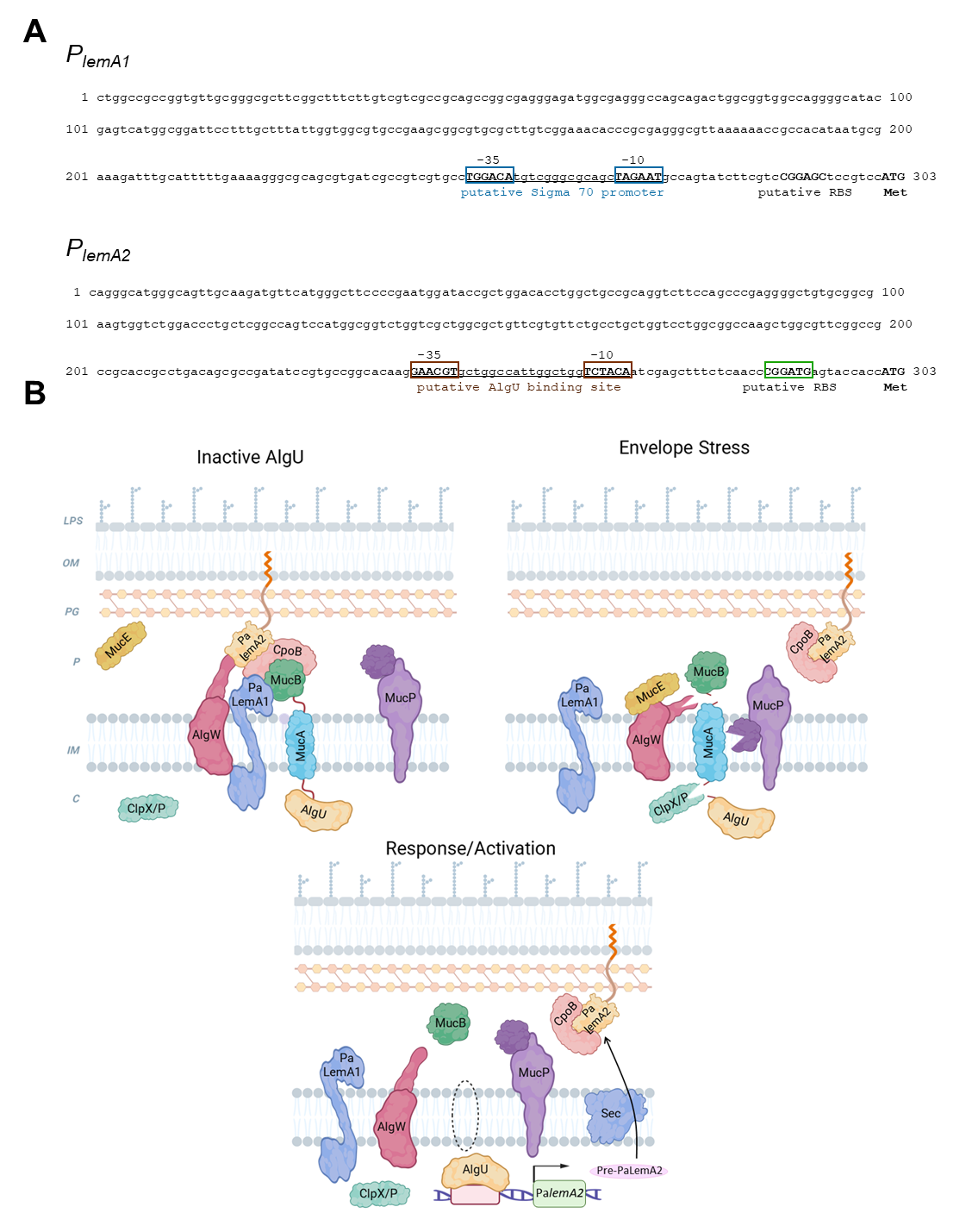

**Extended Data Fig. 20.** (**A**) Predicted promoter sequence of (upper panel) *PalemA1* and (lower panel) *PalemA2*. Shown are the start codons and the putative ribosome binding sites (RBS) for both genes. Note also the putative sigma 70 promoter depicted for *P_lemA1_* and the clear AlgU (sigma E) binding site present in *P_lemA2_.* (**B**) Mechanism of AlgU activation via Regulated Intramembrane Proteolysis (RIP). Under homeostatic conditions (upper left), correctly folded outer membrane proteins (OMPs) prevail, and AlgU is sequestered in the cytosol by the anti-σ factor MucA r(RseA). In this context the PaLemA1-CpoB-PaLemA2 complex may contribute to the stability of the overall complex via binding some of its components as shown by the interactomics data in tables 4-5. Under cell wall/envelope stress (upper right), the periplasm accumulates over-produced MucE, unassembled OMPs, (also truncated type IV pilin and unassembled LPS; not shown). This accumulated LPS binds to MucB, antagonizing its protective effect on MucA. Simultaneously, unfolded proteins bind the AlgW protease, which alongside MucP cleaves MucA to release AlgU (σE). Under these conditions the PaLemA1-CpoB-PaLemA2 complex might dissociate from the involved proteins. The degradation of MucA is finalized by the cytoplasmic protease ClpX, allowing AlgU to associate with the core RNA polymerase (RNAP). This process is regulated by the protease MucD, which represses RIP by chaperoning or degrading misfolded periplasmic proteins, thereby limiting AlgW activation.

Adapted from (269). The figure was created with Biorender.

To gain a deeper insight into this hypothesis, we created in-plasmid promoter fusions with the fluorescent reporter GFP and checked the activity of *P_PalemA1_* (pSB637-PalemA1) and *P_PalemA2_* (pSB637-PalemA2) along the growth curve in LB medium testing different conditions. Initial fluorescence measurements obtained by flow cytometry showed that transcription of Pa*lemA*2 in *P. aeruginosa* PA14 occurs during the transition to the stationary phase (Extended Data Fig. 21). Conversely, we failed to detect the expression of Pa*lemA*1 at any timepoint tested (Extended Data Fig. 21). Following, we tested our reporter strains under cell envelope stress-inducing conditions. For this endeavour, were treated these cultures with subinhibitory doses of either beta-lactam antimicrobials or with the peptidoglycan synthesis inhibitor D-cycloserine and we quantified the GFP fluorescence using flow cytometry. Both types of substances are reported to cause cell wall stress while activating the AlgU regulon (270, 271). We observed that treatment with all cell-envelope-compromising chemicals tested increased the GFP fluorescence triggered by *P_PalemA2_* (Extended Data Fig. 21). To validate that the activation of *PalemA2* was indeed promoted by AlgU, we transferred our reporter plasmids into *P. aeruginosa* PA14 ΔPa*algU* and checked the fluorescence production under the same stressors tested previously. As expected, we observed that the activation of *P_PalemA2_* induced in the WT strain by the antimicrobials and the D-cycloserine was abolished in the strain lacking AlgU, corroborating the essentiality of this ECF sigma factor to fully promote the transcription of *PalemA2* (Extended Data Fig. 21). Yet again, the conditions in which *PalemA1* expression becomes activated remained elusive, as we could not detect fluorescence production through its promoter in our experimental setup (Extended Data Fig. 21).

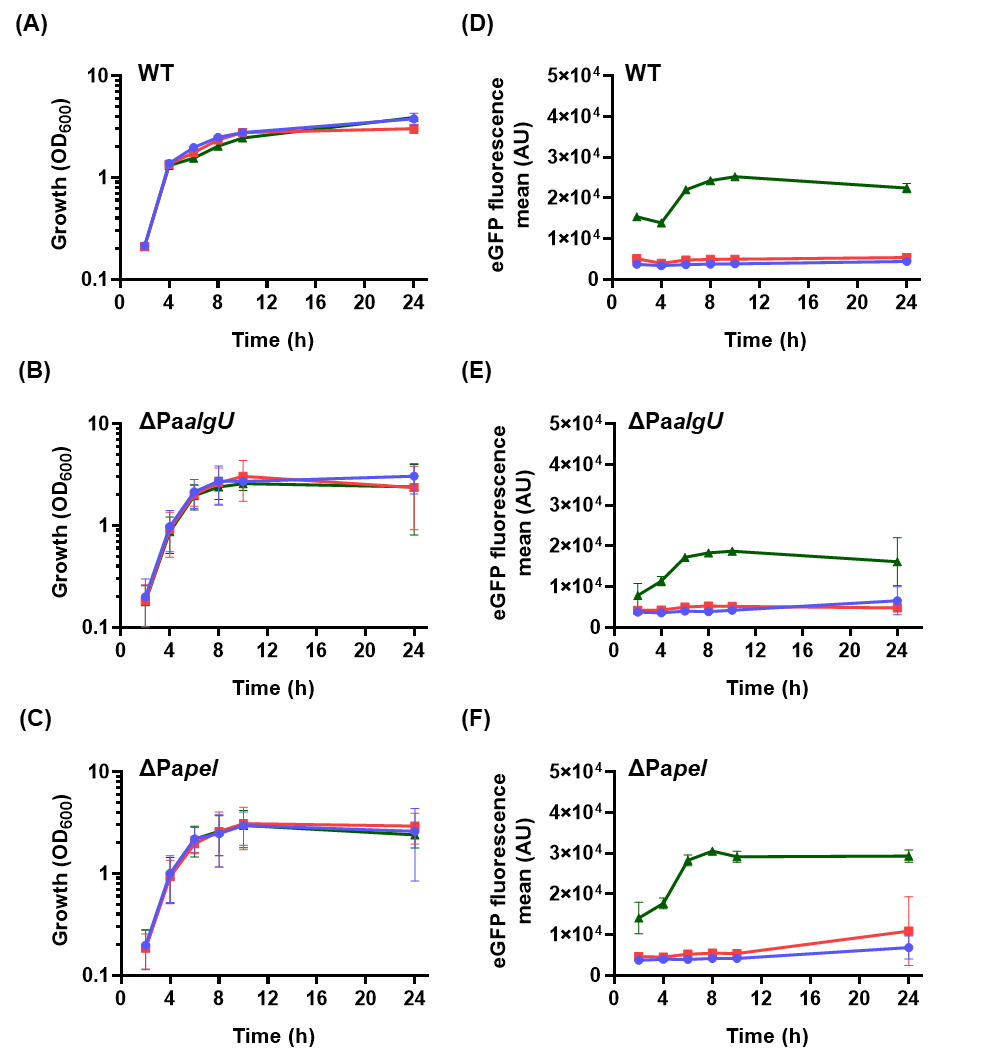

**Extended Data Fig. 21.** Transcriptional activation of *P_PalemA1_* and *P_PalemA2_* in *P. aeruginosa* cells grown under planktonic conditions. Growth curves (**A-C**) and their corresponding eGFP activity (**D-F**) of PA14 wildtype, ΔPa*algU* and ΔPa*pel* strains transformed with the empty plasmid (blue lines) or the vectors carrying the *P_PalemA1_* (red) and *P_PalemA2_* (green) reporter fusions. The arbitrary eGFP units were extracted from the mean fluorescence obtained by analysing the sampled cell populations by flow cytometry**.**

Noteworthy, Wang and colleagues have recently observed that PaLemA2 could be found amongst the most abundant proteins in SDS-insoluble material obtained from PA14 biofilms (272). Moreover, its abundance was strongly increased in biofilms obtained from a mutant in the biosynthesis of the exopolysaccharide (EPS) Pel, the main EPS produced by *P. aeruginosa* PA14. Thus, we constructed a Pa*pel* strain and inspected the activity of *P_lemA1_* and *P_lemA2_* in planktonic cells of this mutant transformed with both reporter plasmids. As shown in Extended Data Fig. 22, the expression through *P_lemA2_* was higher in the *P. aeruginosa* ΔPa*pel* mutant compared to *P. aeruginosa* PA14 wt and the ΔPa*algU* strains. Following this, we also quantified the activity of both promoters in cells recovered from colony biofilms grown in tryptone agar plates (i.e. reproducing the colony growth conditions used by Wang and collaborators).

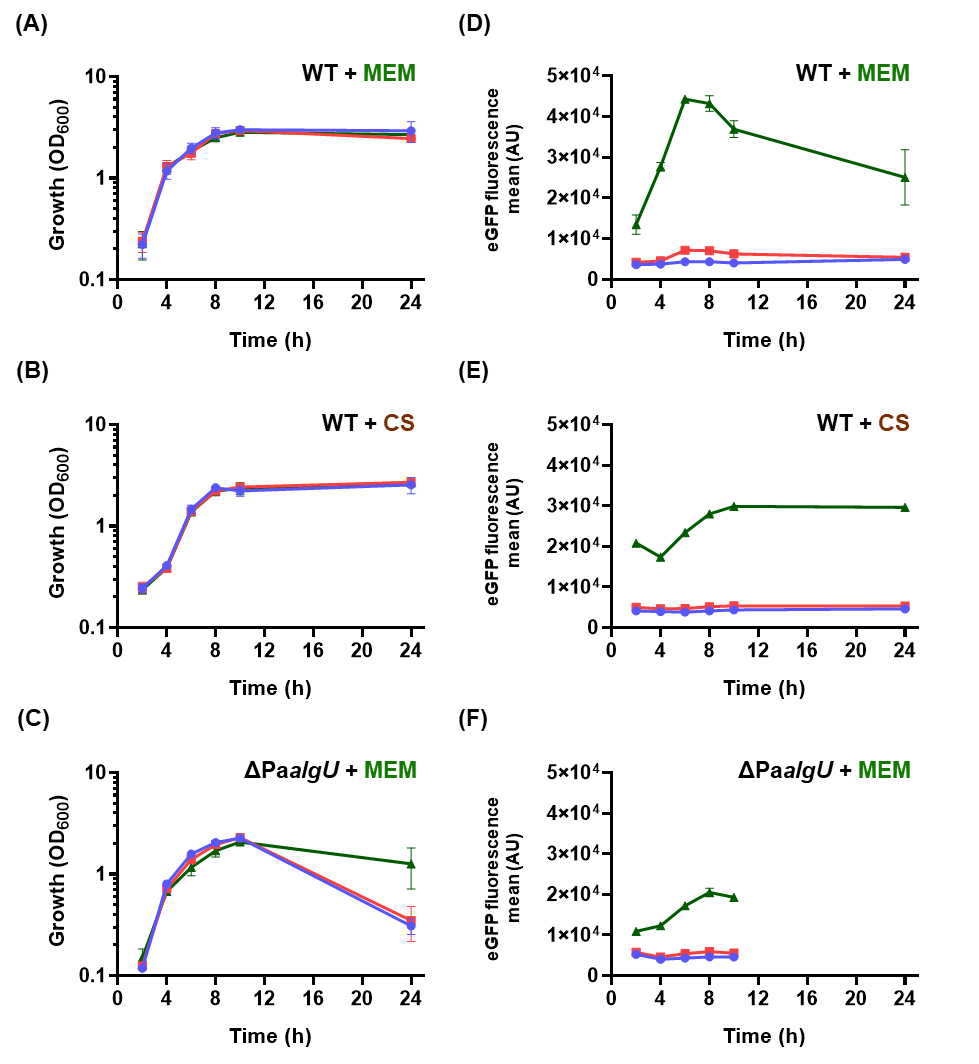

**Extended Data Figure 22. Transcriptional activation of *P_PalemA1_* and *P_PalemA2_* in *P. aeruginosa* cells upon treatment with cell wall stressors.** Growth curves (**A-C**) and their corresponding eGFP activity (**D-F**) of PA14 wildtype and ΔPa*algU* strains transformed with the empty plasmid (blue lines) or the vectors carrying the *P_PalemA1_* (red) and *P_PalemA2_* (green) reporter fusions. The bacterial cultures were treated with a 1/10 MIC concentration of meropenem (MEM; 0.025 mg mL^-1^) or cycloserine (CS; 100 mg mL^-1^) at time point 0. The arbitrary eGFP units were extracted from the mean fluorescence obtained by analysing the sampled cell populations by flow cytometry**.**

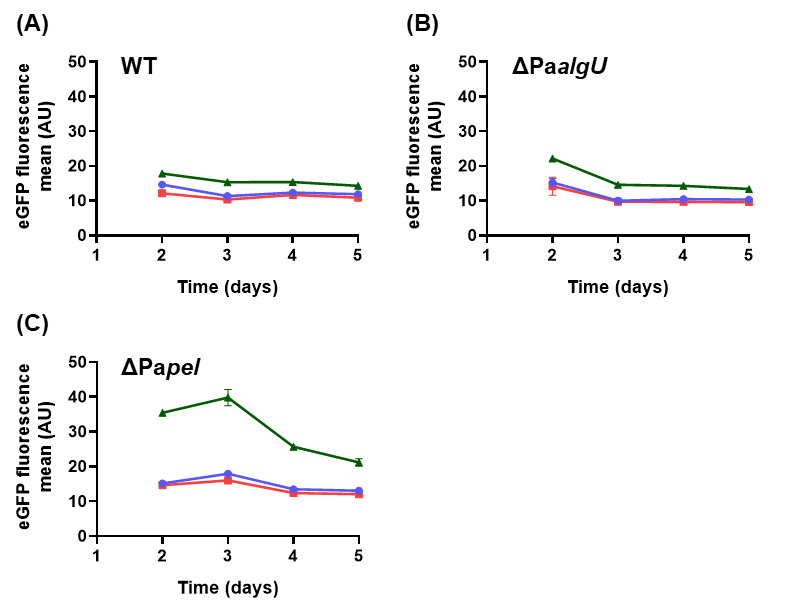

**Extended Data Fig. 23. Transcriptional activation of *P_PalemA1_* and *P_PalemA2_* in *P. aeruginosa* PA14 wt** (**A**), ΔPa*algU* (**B**) and ΔPa*pel* (**C**) strains grown under biofilm conditions on tryptone agar plates. The bacterial strains were transformed with the empty plasmid (blue lines) or the vectors carrying the *P_PalemA1_* (red) and *P_PalemA2_* (green) reporter fusions. Mean eGFP reporter activity was quantified by flow cytometry in individual cells derived from biofilm colonies, which were subsequently resuspended and diluted in PBS.

The colonies were resuspended in PBS and the GFP fluorescence of bacteria was analysed by flow cytometry. In the case of *P_lemA1_*, no expression was observed in biofilm-grown cells, as was also the case for planktonic-grown cultures. Meanwhile, colonies from the ΔPa*pel* mutant reported higher *P_lemA2_*-stemming fluorescence reads than PA14 wt and ΔPa*algU* strains Altogether, these results indicate that *P_lemA2_* activation occurs during stationary phase and requires the alternative sigma factor AlgU for its full expression. Moreover, the influence of AlgU and the impact of Pel exopolysaccharide production on the expression of *P_PalemA2_*, suggests that PaLemA2 is produced upon cell wall stress-inducing insults, such as the damage elicited by the antibiotics MEM and CS. From the obtained results a regulatory model was deduced Extended Data Fig. 20).

20. ***In silico* structural analysis of PaLemAs-PaCpoB interactions in *P. aeruginosa***

The LemA protein family has been implicated in envelope-associated processes in Gram-negative bacteria, yet its molecular interaction landscape remains incompletely defined. To explore potential structural relationships in *P. aeruginosa* PA14, AlphaFold3 multimer modelling was performed for PaLemA1, PaLemA2, and the outer membrane-associated protein PaCpoB.

Membrane topology predictions were generated using SignalP 6.0, DeepTMHMM, ProtT5, and SecretomeP. PaLemA1 is predicted to be a single‑pass inner‑membrane protein with a transmembrane helix spanning residues 6–22 (DeepTMHMM). Residues 1–5 are located in the cytoplasm, while residues 23–190 form a periplasmic domain, while no cleavable Sec signal peptide is found. The enrichment of positively charged residues N‑terminal to the helix supports cytoplasmic retention according to the positive‑inside rule, and ProtT5 assigns it to the cytoplasmic membrane with 0.94 probability. TmLemA1 shows an equivalent architecture with a helix at residues 11–31, a short cytoplasmic N‑terminus (1–10), and a periplasmic C‑terminus (32–193), also lacking a cleavable signal peptide and predicted to localize to the cytoplasmic membrane (ProtT5 probability 0.969). Thus, PaLemA1 and TmLemA1 share a conserved signal‑anchor topology. In contrast, PaLemA2 is identified as a lipoprotein. SignalP detects a canonical Sec/SPII signal peptide with a lipobox and a cleavage site between residues 19-20, placing a lipidated cysteine at position +1, and no additional transmembrane helices are predicted. ProtT5 predicts outer‑membrane localization (0.72 probability), indicating that PaLemA2 is membrane‑anchored via lipid modification and exposed to the periplasm. MamQ displays an opposite orientation with an internal helix (residues 47-67) places residues 1-46 in the periplasm and residues 68-272 in the cytoplasm, with no signal peptide detected, and ProtT5 again predicts cytoplasmic‑membrane localization (0.95 probability). Overall, LemA proteins share a periplasm‑exposed folded domain but differ in their membrane‑targeting mechanisms, whereas MamQ possesses a distinct, inverted topology.

**Structural conservation among LemA homologs**

Pairwise structural alignments of AlphaFold3 models were performed using PyMOL, restricted to high-confidence regions (pLDDT > 70; Extended Data Fig. 24). Strong structural conservation was observed between TmLemA1 and PaLemA2 (RMSD 4.176 Å over 184 Cα atoms; 91.1% coverage). Alignment of TmLemA1 and PaLemA1 shows high coverage (92.6%) but increased deviation (RMSD 12.861 Å). The two *P. aeruginosa* paralogs (PaLemA1 vs PaLemA2) exhibit moderate divergence (RMSD 8.244 Å; 91.1% coverage). In contrast, MamQ is structurally distinct from all LemA homologs (RMSD 56–60 Å; ~59.6% coverage), indicating absence of the conserved LemA core fold. Collectively, these analyses indicate conservation of a LemA structural scaffold across species and membrane-targeting strategies, whereas MamQ represents a distinct protein family. (Extended Data Fig. 24)

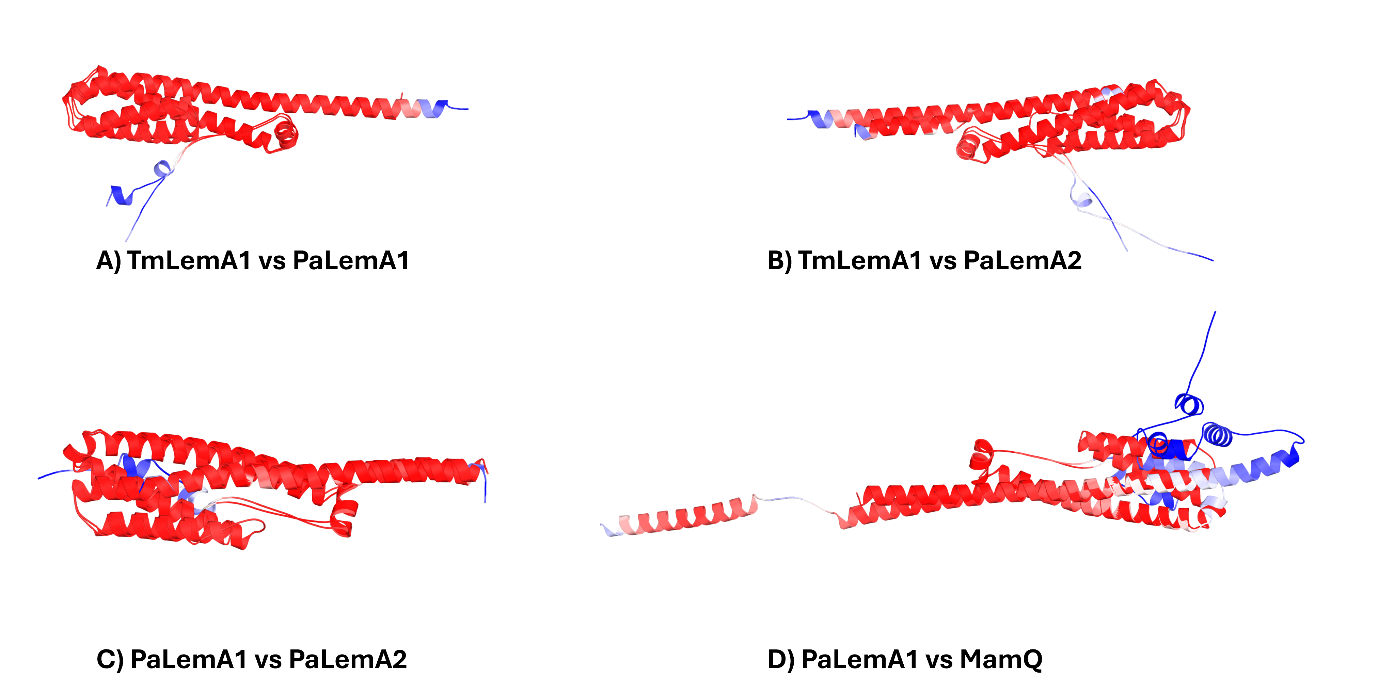

**Extended Data Fig. 24. Structural analysis of LemA homologs and MamQ by AlphaFold prediction and pairwise alignment.** AlphaFold3-predicted structures coloured by per-residue confidence (pLDDT). Colour scheme: red (≥90, very high confidence), pink (70–89, high confidence), grey (50–69, low confidence), light gray (<50, very low confidence). Models display well-folded regions (red/pink) with high prediction accuracy, while disordered regions (grey) indicate lower confidence. **Pairwise structural alignments of LemA homologs and MamQ.** Structures were aligned using PyMOL based on high-confidence regions (pLDDT > 70). RMSD values and aligned Cα atoms are indicated. Colouring reflects pLDDT confidence as described above. A) *Thermotoga maritima* LemA (TmLemA) vs PaLemA1. RMSD 12.861 Å; 176 aligned Cα atoms; 92.6% coverage of PaLemA1. B) *Thermotoga maritima* LemA vs PaLemA2. RMSD 4.176 Å; 184 aligned Cα atoms; 91.1% coverage of PaLemA2. C)PaLemA1 vs PaLemA2. RMSD 8.244 Å; 184 aligned Cα atoms; 91.1% coverage of PaLemA2. D)PaLemA1 vs MamQ. RMSD 59.855 Å; 162 aligned Cα atoms; 59.6% coverage of MamQ.

**PaLemA proteins interact with PaCpoB through distinct binding modes**

AlphaFold3 multimer modelling was used to predict PaLemA1-PaCpoB, PaLemA2–PaCpoB, and PaLemA1–PaLemA2–PaCpoB complexes (Extended Data Fig. 25). Interaction interfaces were defined using a heavy-atom distance threshold of <4 Å. All reported contact residues are located within high-confidence regions (pLDDT ≥ 70). The PaLemA1–PaCpoB model predicts a multi-contact interface. Two primary contact pairs were identified (PaLemA1-111 and PaCpoB-224 (3.64 Å); PaLemA1-178 and PaCpoB-149 (3.78 Å)). Additional residues (107, 110, 189) cluster within the same interfacial region. The spatial clustering of contacts suggests an extended interaction surface. In contrast, PaLemA2 engages PaCpoB through a more restricted interface comprising two contact pairs (PaLemA2-58 and PaCpoB-261 (3.53 Å); PaLemA2-119 and PaCpoB-202 (3.76 Å)). No additional residues were consistently observed within the <4 Å threshold across prediction runs, indicating a comparatively discrete interaction surface. The different interaction patterns of PaLemA1 and PaLemA2 with PaCpoB indicate functional diversification within the LemA family. PaLemA1 uses an extended surface that contacts many residues across its folded domain, creating a high‑avidity, long‑lived complex that is ideal for structural or scaffolding roles. Its inner‑membrane anchoring, with the C‑terminal domain in the periplasm, positions it as a stable platform for recruiting or retaining PaCpoB at specific membrane sites. In contrast, PaLemA2 contacts PaCpoB through two small, spatially separated residue pairs. This minimal interface permits rapid, reversible binding, suggesting a regulatory function that can respond to cellular signals or compete with other PaCpoB partners. As a lipid‑anchored lipoprotein, PaLemA2’s modest binding surface is consistent with a dynamic, modulatory role rather than a structural one.

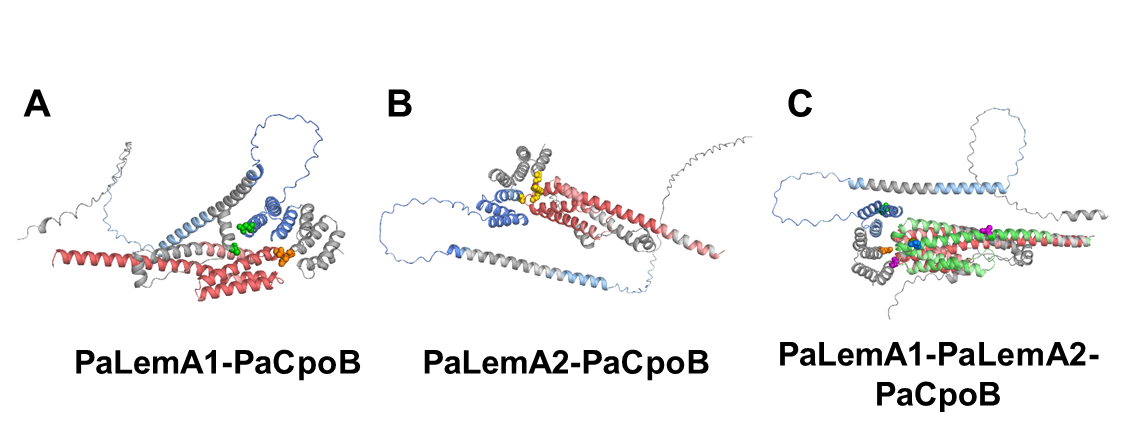

**Extended Data Fig. 25. AlphaFold3 confidence assessment of PaLemA-PaCpoB interaction complexes.** Models are coloured by per-residue confidence (pLDDT) following standard AlphaFold conventions: red (high confidence, pLDDT ≥ 90), pink (medium confidence, 70–90), grey (low confidence, 50–70), and light grey (very low confidence, <50). For each complex, the top-ranked model (rank_0) is shown in cartoon representation with transparent surface rendering to visualize interaction interfaces. (**A**) PaLemA1-PaCpoB binary complex. PaLemA1 (left) and PaCpoB (right) are shown with confidence colouring. The majority of both proteins fold with high to medium confidence, particularly in structured core regions. The interaction interface (boxed region, enlarged in inset) involves residues predominantly within high-confidence areas (red/pink), supporting the reliability of the predicted contact pairs PaLemA1-111–PaCpoB-224 and PaLemA1-178–PaCpoB-149. Low-confidence regions are limited to flexible termini and loop segments distant from the binding interface. (**B**) PaLemA2–PaCpoB binary complex. PaLemA2 (left) and PaCpoB (right) displayed with confidence colouring. Both critical contact pairs (PaLemA2-58–PaCpoB-261 and PaLemA2-119–PaCpoB-202) involve residues with high confidence scores (pLDDT > 88, red/pink), as shown in the enlarged interface view (inset). The structured core of PaLemA2 is well-modelled, while PaCpoB exhibits high confidence throughout the regions involved in PaLemA2 binding. (**C**) PaLemA1–PaLemA2–PaCpoB ternary complex. Simultaneous binding of both PaLemA proteins to PaCpoB. Colour scheme: PaLemA1 (chain A), PaLemA2 (chain B), and PaCpoB (chain C) are each coloured by per-residue confidence as described above. All four identified contact pairs -PaLemA1-107–PaCpoB-224, PaLemA1-147–PaCpoB-149, PaLemA2-55–PaCpoB-261, and PaLemA2-34–PaCpoB-261- involve residues within high-confidence structural contexts (red/pink). Enlarged views (insets) show each interaction interface with confidence colouring, demonstrating that predicted contacts occur in well-folded regions of all three proteins. The consistent localization of interaction interfaces to high-confidence structural regions across all three complexes supports the biological plausibility of the predicted binding modes and indicates that the observed contacts are not artifacts of poorly modelled regions.

**Distinct binding modes suggest functional diversification**

The fact that PaLemA1 and PaLemA2 bind PaCpoB with markedly different architectures reveals functional diversification within the PaLemA family. Although the two paralogs share a common fold (RMSD 8.2 Å over 184 Cα atoms; Supplementary Fig. S23), PaLemA1 uses an extensive, multi‑contact interface that provides broad surface complementarity and likely forms a high‑affinity, stoichiometric complex. Such a scaffold‑like interaction is consistent with a constitutive role, anchoring PaCpoB‑containing assemblies at the inner membrane–periplasm interface, as expected for a single‑pass inner‑membrane protein with a periplasmic C‑terminus. In contrast, PaLemA2 interacts with PaCpoB through a small, spatially separated patch of contacts, a hallmark of transient, regulatory bindings that allow rapid association‑dissociation cycles and conformational flexibility. Its classification as a lipoprotein suggests lateral mobility in the outer leaflet of the inner membrane, enabling dynamic repositioning in response to envelope cues. Together, these divergent interaction modes indicate that PaLemA1 functions primarily as a structural stabilizer, whereas PaLemA2 serves a more modulatory, context‑dependent role.

**Implications for envelope-associated complex assembly**

Both PaLemA proteins can bind PaCpoB at the same time, implying the formation of a ternary complex that spans the periplasm and physically links the inner and outer membranes. PaLemA1 is an inner‑membrane protein whose C‑terminal domain resides in the periplasm, whereas PaLemA2 is a lipid‑anchored periplasmic protein; together they could create a bridge that mechanically couples inner‑ and outer‑membrane dynamics during processes such as cell division, transmits conformational or stress signals across the envelope, and/or reinforces regions of curvature or mechanical stress. Comparison of binary and ternary interaction models shows subtle shifts in residue contacts, e.g. the PaLemA2‑119–PaCpoB‑202 interaction present in the binary complex moves beyond 4 Å in the ternary assembly, suggesting either allosteric rearrangements upon higher‑order assembly or intrinsic flexibility that allows multiple quasi‑stable binding states, a hypothesis that will require targeted biochemical and structural tests. Structurally, LemA proteins retain a conserved core fold despite disparate membrane‑targeting strategies. TmLemA and PaLemA2 align with an RMSD of 4.2 Å, indicating that the periplasmic domain alone can mediate PaCpoB binding without transmembrane helices. In contrast, PaLemA1 and PaLemA2 diverge (RMSD 8.2 Å), consistent with subfunctionalization after gene duplication PaLemA1 may act as a stabilizing scaffold while PaLemA2 provides a more dynamic, modulatory interface. MamQ, however, is far more divergent (RMSD > 56 Å), confirming it belongs to a separate protein family and that the PaLemA-PaCpoB interaction network is specific to LemA homologs.

Despite methodological limitations, the present *in silico* analysis provides the first structural framework for understanding PaLemA-PaCpoB interaction networks in *P. aeruginosa*. The data reveal clearly distinguishable binding architectures for PaLemA1 and PaLemA2 and demonstrate the structural compatibility of simultaneous CpoB engagement within a ternary assembly that could, in principle, bridge inner and outer membrane compartments. When interpreted in the context of membrane topologies and structural comparisons, these findings support a model in which functional diversification within the LemA family extends to interaction geometry and complex assembly behaviour.

**Materials and Methods Tables**

**Extended Table 15.** Bacterial strains, vectors and oligonucleotides employed throughout this study.

| **Bacterial strains** | **Relevant features ^a^** | **Source/Reference** |
| --- | --- | --- |
| ***Pseudomonas aeruginosa*** |  |  |
| PA14 | Wildtype strain | DSMZ |
| ΔPa*lemA1* | *P. aeruginosa* PA14Δ*lemA1* knockout mutant strain | This study |
| ΔPa*lemA2* | *P. aeruginosa* PA14Δ*lemA2* knockout mutant strain | This study |
| ΔPa*lemA1+A2* | *P. aeruginosa* double knockout mutant strain deficient in the expression of *lemA1* and *lemA2* | This study |
| ΔPa*lemA1*+pPa*lemA1* | ΔPa*lemA1* complemented with the pSEVA634 plasmid harbouring Pa*lemA1* | This study |
| ΔPa*lemA2*+pPa*lemA2* | ΔPa*lemA2* complemented with the pSEVA634 plasmid harbouring Pa*lemA2* | This study |
| ΔPa*lemA1+A2*+pPa*lemA1+A2* | ΔPa*lemA1+A2* complemented with the  pSEVA634 plasmid harbouring both Pa*lemA* genes | This study |
| ΔPa*lemA1+A2*+p*ctrl63*4 | ΔPa*lemA1*+*A2* complemented with the  pSEVA634 plasmid | This study |
| ΔPa*algU* | *P. aeruginosa* PA14 carrying a full deletion of *algU* | (273) |
| ΔPa*pel* | *P. aeruginosa* PA14 carrying a deletion of *pelBCDEFG* | This study |
| ΔPa*cpoB* | *P. aeruginosa* PA14 carrying a deletion of *cpoB* | This study |
| ΔPa*cpoB*+p*ctrl634* | ΔPa*cpoB* complemented with pSEVA634 | This study |
| ΔPa*cpoB*+pPa*cpoB* | ΔPa*cpoB* complemented with pSEVA634 harbouring Pa*cpoB* | This study |
| ΔPaPa*pspA* | *P. aeruginosa* PA14 Δ*pspA* knockout mutant strain | This study |
| ΔPa*pspA*+pPa*pspA* | ΔPa*pspA* complemented with pSEVA228 harbouring Pa*pspA* | This study |
| ΔPa*pspA*+p*ctrl228* | ΔPa*pspA* complemented with pSEVA228 | This study |
| ΔPa*pspA*+pPa*lemA1* | ΔPa*pspA* complemented with pSEVA634 harbouring Pa*lemA1* | This study |
| ΔPa*pspA*+pPa*lemA2* | ΔPa*pspA* complemented with pSEVA634 harbouring Pa*lemA2* | This study |
| ΔPa*pspA*+pPa*lemA+A2* | ΔPa*pspA* complemented with the  pSEVA634 plasmid harbouring both Pa*lemA* genes | This study |
| ΔPa*pspA*+p*ctrl634* | ΔPa*pspA* complemented with pSEVA634 | This study |
| ***Escherichia coli*** |  |  |
| DH5α | F– Φ80*lacZ*ΔM15 Δ(*lacZYA*-*arg*F) U169 *rec*A1 *end*A1 *hsd*R17 (rK–,  mK+) *pho*A *sup*E44 λ– *thi*-1 *gyr*A96 *rel*A1 | ThermoFisher, Scientific, Darmstadt, Germany |
| DH5αλpir | *sup* E44, Δ*lacU169* (Φ*lacZ*ΔM15), *recA1*, *endA1*, *hsdR17*, *thi*-1, *gyrA96*, *relA1*, λpir phage lysogen | Biomedal, Seville, Spain |
| ST18 | *pro thi hsdR1* Tpr Sm^R^; chromosome::RP4-2 Tc^R^::Mu-Kan::Tn7/lpir Δ*hemA* | (274) |
| BL21(DE3) Star | *F - ompT hsdS B (r B - m B - ) gal dcm rne131 (DE3)* | ThermoFisher Scientific, Darmstadt, Germany |
| BTH101 | *F-, cya-99, araD139, galE15, galK16, rpsL1 (Str r), hsdR2, mcrA1, mcrB1* | (275) |
| **Vectors** |  |  |
| pET3a.BcLemA1 | pET3a* vector containing Bc*lemA1* | This study |
| pET3a.BmLemA2 | pET3a* vector containing Bm*lemA2* | This study |
| pET3a.BrmLemA2 | pET3a* vector containing Brm*lemA2* | This study |
| pET3a.CreLemA2 | pET3a* vector containing Cre*lemA2* | This study |
| pET3a.ExcLemA1 | pET3a* vector containing Exc*lemA1* | This study |
| pET3a.LmLemA1 | pET3a* vector containing Lm*lemA1* | This study |
| pET3a.PaLemA1 | pET3a* vector containing Pa*lemA1* | This study |
| pET3a.PaLemA2 | pET3a* vector containing Pa*lemA2* | This study |
| pET3a.StLemA2 | pET3a* vector containing St*lemA2* | This study |
| pET3a.ThmLemA2 | pET3a* vector containing Thm*lemA2* | This study |
| pSEVA634 | Gm^R^; *oriV* (pBBR1), *lacI^Q^/Ptrc*. Broad host-range replicative plasmid inducible upon IPTG addition | (276) |
| pSEVA634-Pa*lemA1* | pSEVA634 vector containing Pa*lemA1* | This study |
| pSEVA634-Pa*lemA2* | pSEVA634 vector containing Pa*lemA2* | This study |
| pSEVA634-Pa*lemA1+A2* | pSEVA634 vector containing Pa*lemA1* and Pa*lemA2* | This study |
| pSEVA634-Pa*cpoB* | pSEVA634 vector containing Pa*cpoB* | This study |
| pSEVA228 | Km^R^, *ori V* (RK2), xylS/Pm. Broad-host-range low-copy replicative plasmid inducible upon 3-*m*Bz addition | (276) |
| pSEVA228-Pa*pspA* | pSEVA228 vector containing Pa*pspA* | This study |
| pSEVA637 | Gm^R^, *oriV* pBBR1, *oriT*, promoter-less cloning vector including *gfp* as a reporter gene | (276) |
| pSEVA637-*PlemA1* | Gm^R^, pSEVA637 cloned with the promoter sequence of Pa*lemA1* | This study |
| pSEVA637-*PlemA2* | Gm^R^, pSEVA637 cloned with the promoter sequence of Pa*lemA2* | This study |
| pS448∙CsR | Sm^R^*, oriV pRO1600/ColE1*, *oriT*, *xylS*, *Pm*à*cas9*, *P_EM7_*àsgRNA | (277, 278) |
| pS448∙CsR-PA14-pelB1 | Sm^R^*,* pS448∙CsR derivative expressing the specific sgRNA targeting *pelB* | This study |
| pS448∙CsR-PA14-pspA1 | Sm^R^*,* pS448∙CsR derivative expressing the specific sgRNA targeting *pspA* | This study |
| pS448∙CsR-PA14-cpoB2 | Sm^R^*,* pS448∙CsR derivative expressing the specific sgRNA targeting *cpoB* | This study |
| pSH624-*ssr* | Gm^R^*, oriV* *RK2, oriT*, *lacI^q^*,*P_trc_*à*ssr* | (278, 279) |
| pKT25 | Encoding the T25 fragment (1-224 amino acids of CyaA). MCS at the 3' end of T25 to  allow construction of in-frame fusions at the C-terminal end of the T25. Km^R^ | (275) |
| pKNT25 | Harbouring the T25 sequence fused in frame downstream from a MCS. Km^R^ | (275) |
| pUT18 | Encoding the T18 fragment (225 - 399 amino acids of CyaA) downstream of a MCS. Amp^R^ | (275) |
| pUT18C | MCS is located at the 3’ end of the T18 open reading frame. Amp^R^ | (275) |
| pKT25-zip | Leucine zipper motif fused to T25 | (275) |
| pUT18C-zip | Leucine zipper motif fused to T25 | (275) |
| pKT25-*lemA1* | *lemA1* gene fragment encoding its cytoplasmic domain fused to the N-terminus of T25 | This study |
| pUT18C-*lemA1* | *lemA1* gene fragment encoding its cytoplasmic domain fused to N-terminus of T18 | This study |
| pKT25-*lemA2* | *lemA2* gene fragment encoding its periplasmic domain lacking the signal peptide and fused to N-terminus of T25 | This study |
| pUT18C-*lemA2* | *lemA2* gene fragment encoding its periplasmic domain lacking the signal peptide and fused to N-terminus of T18 | This study |
| pKT25-*cpoB* | *cpoB* gene fragment encoding its periplasmic domain lacking the signal peptide and fused to N-terminus of T25 | This study |
| pKNT25-*cpoB* | *cpoB* gene fragment encoding its periplasmic domain lacking the signal peptide and fused to C-terminus of T25 | This study |
| pUT18-*cpoB* | *cpoB* gene fragment encoding its periplasmic domain lacking the signal peptide and fused to C-terminus of T18 | This study |
| pUT18C-*cpoB* | *cpoB* gene fragment encoding its periplasmic domain lacking the signal peptide and fused to N-terminus of T18 | This study |
| pKNT25-*mrcB* | *mrcB* gene fragment encoding its cytoplasmic region fused to C-terminus of T25 | This study |
| pUT18-*mrcB* | *mrcB* gene fragment encoding its cytoplasmic region fused to C-terminus of T18 | This study |
| pKT25-*ftsN* | *ftsN* gene fragment encoding its periplasmic domain fused to C-terminus of T25 | This study |
| pUT18C-*ftsN* | *ftsN* gene fragment encoding its periplasmic domain fused to C-terminus of T18 | This study |
| **Oligos (5´ 🡪** **3´)** |  |  |
| MgMamQase | GCTGATTAATATGGCAGTAAGCGATGC | ThermoFisher Scientific, Schwerte, Germany |
| MgMamQspe | GTGCACTAGTCAATTCTTGGATTCCTG | ThermoFisher Scientific, Schwerte, Germany |
| MgMamQ∆72 | GTCGAGCTCATTCTTGGATTCCTGCGAATG | ThermoFisher Scientific, Schwerte, Germany |
| FwBmLemA2nde | GAGCATATGAAGAAAAGAGGTAGTACATTG | ThermoFisher Scientific, Schwerte, Germany |
| RvBmLemA2spe | CATACTAGTCACTTATCGCTCCCAAAATCA AC | ThermoFisher Scientific, Schwerte, Germany |
| FwBrmLemA2nde | GTACATATGACCGCTCAGACGGTTGCCAC | ThermoFisher Scientific,  Schwerte, Germany |
| RvBrmLemA2spe | GACACTAGTCAGTTGAAGTTGACCTTAG | ThermoFisher Scientific, Schwerte, Germany |
| FwPaLemA1nde | GTACATATGAGTCTGACCGCTATCGCTTTC | ThermoFisher Scientific, Schwerte, Germany |
| RvPaLemA1spe | GTAACTAGTCAGCCGAACAGGGCCTTGAG | ThermoFisher Scientific, Schwerte, Germany |
| RvPaLemA1xho | GTACTCGAGCCGAACAGGGCCTTGAG | ThermoFisher Scientific, Schwerte, Germany |
| FwPaLemA2nde | GTACATATGCCTCGATTCCTCAG | ThermoFisher Scientific, Schwerte, Germany |
| RvPaLemA2xho | GTACTCGAGCTGCGCCGGTTGCGGGTTG | ThermoFisher Scientific, Schwerte, Germany |
| 56810UpFw | GGTACCTCCTTGACGTCGATCTTGTCT | ThermoFisher Scientific, Schwerte, Germany |
| 56810UpRv | AGCGCAGGTCGAAGTCCATCGGACGGAGC TCCGGACGAAGA | ThermoFisher Scientific, Schwerte, Germany |
| 56810DwFw | GATGGACTTCGACCTGCGCT | ThermoFisher Scientific, Schwerte, Germany |
| 56810DwRv | TCTAGAATTCGCGCTCGTCCAACT | ThermoFisher Scientific, Schwerte, Germany |
| 06990UpFw | GAATTCTTAATCTTCGGGCAATCCTG | ThermoFisher Scientific, Schwerte, Germany |
| 06990UpRv | AGTAGCAGCCGGCGCATGGCGGTGGTACT CATCCGGGTTG | ThermoFisher Scientific, Schwerte, Germany |

**References**

1.	Lopez-Marques RL, Perez-Castineira JR, Losada M, Serrano A. 2004. Differential regulation of soluble and membrane-bound inorganic pyrophosphatases in the photosynthetic bacterium *Rhodospirillum* *rubrum* provides insights into pyrophosphate-based stress bioenergetics. *J* *Bacteriol* 186:5418–26.

2.	Cornejo E, Abreu N, Komeili A. 2014. Compartmentalization and organelle formation in bacteria. *Curr* *Opin* *Cell* *Biol* 26:132–8.

3.	Bobik TA, Lehman BP, Yeates TO. 2015. Bacterial microcompartments: widespread prokaryotic organelles for isolation and optimization of metabolic pathways. *Mol* *Microbiol* 98:193–207.

4.	Grant CR, Wan J, Komeili A. 2018. Organelle Formation in Bacteria and Archaea. *Annu* *Rev* *Cell* *Dev* *Biol* 34:217–238.

5.	Taoka A, Eguchi Y, Shimoshige R, Fukumori Y. 2023. Recent advances in studies on magnetosome-associated proteins composing the bacterial geomagnetic sensor organelle. *Microbiol* *Immunol* 67:228–238.

6.	Toyofuku M, Morinaga K, Hashimoto Y, Uhl J, Shimamura H, Inaba H, Schmitt-Kopplin P, Eberl L, Nomura N. 2017. Membrane vesicle-mediated bacterial communication. *ISME* *J* 11:1504–1509.

7.	Toyofuku M, Nomura N, Eberl L. 2019. Types and origins of bacterial membrane vesicles. *Nat* *Rev* *Microbiol* 17:13–24.

8.	Munoz-Echeverri LM, Benavides-Lopez S, Geiger O, Trujillo-Roldan MA, Valdez-Cruz NA. 2024. Bacterial extracellular vesicles: biotechnological perspective for enhanced productivity. *World* *J* *Microbiol* *Biotechnol* 40:174.

9.	Meidaninikjeh S, Mohammadi P, Elikaei A. 2024. Bacteriophages and bacterial extracellular vesicles, threat or opportunity? *Life* *Sci* 350:122749.

10.	Ho MY, Liu S, Xing B. 2024. Bacteria extracellular vesicle as nanopharmaceuticals for versatile biomedical potential. *Nano* *Converg* 11:28.

11.	Xiu L, Wu Y, Lin G, Zhang Y, Huang L. 2024. Bacterial membrane vesicles: orchestrators of interkingdom interactions in microbial communities for environmental adaptation and pathogenic dynamics. *Front* *Immunol* 15:1371317.

12.	Zhao G, Jones MK. 2023. Role of Bacterial Extracellular Vesicles in Manipulating Infection. *Infect* *Immun* 91:e0043922.

13.	Schwechheimer C, Kuehn MJ. 2015. Outer-membrane vesicles from Gram-negative bacteria: biogenesis and functions. *Nat* *Rev* *Microbiol* 13:605–19.

14.	Beveridge TJ. 1999. Structures of gram-negative cell walls and their derived membrane vesicles. *J* *Bacteriol* 181:4725–33.

15.	Mashburn-Warren LM, Whiteley M. 2006. Special delivery: vesicle trafficking in prokaryotes. *Mol* *Microbiol* 61:839–46.

16.	Turnbull L, Toyofuku M, Hynen AL, Kurosawa M, Pessi G, Petty NK, Osvath SR, Carcamo-Oyarce G, Gloag ES, Shimoni R, Omasits U, Ito S, Yap X, Monahan LG, Cavaliere R, Ahrens CH, Charles IG, Nomura N, Eberl L, Whitchurch CB. 2016. Explosive cell lysis as a mechanism for the biogenesis of bacterial membrane vesicles and biofilms. *Nat* *Commun* 7:11220.

17.	Sangiorgio G, Nicitra E, Bivona D, Bonomo C, Bonacci P, Santagati M, Musso N, Bongiorno D, Stefani S. 2024. Interactions of Gram-Positive Bacterial Membrane Vesicles and Hosts: Updates and Future Directions. *Int* *J* *Mol* *Sci* 25.

18.	Grande R, Di Marcantonio MC, Robuffo I, Pompilio A, Celia C, Di Marzio L, Paolino D, Codagnone M, Muraro R, Stoodley P, Hall-Stoodley L, Mincione G. 2015. *Helicobacter* *pylori* ATCC 43629/NCTC 11639 Outer Membrane Vesicles (OMVs) from Biofilm and Planktonic Phase Associated with Extracellular DNA (eDNA). *Front* *Microbiol* 6:1369.

19.	Orench-Rivera N, Kuehn MJ. 2016. Environmentally controlled bacterial vesicle-mediated export. *Cell* *Microbiol* 18:1525–1536.

20.	Bitto NJ, Chapman R, Pidot S, Costin A, Lo C, Choi J, D'Cruze T, Reynolds EC, Dashper SG, Turnbull L, Whitchurch CB, Stinear TP, Stacey KJ, Ferrero RL. 2017. Bacterial membrane vesicles transport their DNA cargo into host cells. *Sci* *Rep* 7:7072.

21.	Sjostrom AE, Sandblad L, Uhlin BE, Wai SN. 2015. Membrane vesicle-mediated release of bacterial RNA. *Sci* *Rep* 5:15329.

22.	Kaparakis-Liaskos M, Ferrero RL. 2015. Immune modulation by bacterial outer membrane vesicles. *Nat* *Rev* *Immunol* 15:375–87.

23.	Manning AJ, Kuehn MJ. 2011. Contribution of bacterial outer membrane vesicles to innate bacterial defense. *BMC* *Microbiol* 11:258.

24.	Kwaku GN, Ward RA, Vyas JM, Harding HB. 2024. Host innate immune systems gather intel on invading microbes via pathogen-derived extracellular vesicles. *Extracell* *Vesicle* 3.

25.	Peregrino ES, Castaneda-Casimiro J, Vazquez-Flores L, Estrada-Parra S, Wong-Baeza C, Serafin-Lopez J, Wong-Baeza I. 2024. The Role of Bacterial Extracellular Vesicles in the Immune Response to Pathogens, and Therapeutic Opportunities. *Int* *J* *Mol* *Sci* 25.

26.	Nevermann J, Silva A, Otero C, Oyarzun DP, Barrera B, Gil F, Calderon IL, Fuentes JA. 2019. Identification of Genes Involved in Biogenesis of Outer Membrane Vesicles (OMVs) in *Salmonella* *enterica* Serovar Typhi. *Front* *Microbiol* 10:104.

27.	Schlosser L, Sachse C, Low HH, Schneider D. 2023. Conserved structures of ESCRT-III superfamily members across domains of life. *Trends* *Biochem* Sci 48:993–1004.

28.	Liu J, Tassinari M, Souza DP, Naskar S, Noel JK, Bohuszewicz O, Buck M, Williams TA, Baum B, Low HH. 2021. Bacterial Vipp1 and PspA are members of the ancient ESCRT-III membrane-remodeling superfamily. *Cell* 184:3660–3673 e18.

29.	Junglas B, Huber ST, Heidler T, Schlosser L, Mann D, Hennig R, Clarke M, Hellmann N, Schneider D, Sachse C. 2021. PspA adopts an ESCRT-III-like fold and remodels bacterial membranes. *Cell* 184:3674–3688 e18.

30.	Liu M, Liu Y, Song T, Yang L, Qi L, Zhang YZ, Wang Y, Shen QT. 2024. Three-dimensional architecture of ESCRT-III flat spirals on the membrane. *Proc* *Natl* *Acad* *Sci* U S A 121:e2319115121.

31.	Hale CA, Persons L, de Boer PAJ. 2022. Recruitment of the TolA Protein to Cell Constriction Sites in *Escherichia* *coli* via Three Separate Mechanisms, and a Critical Role for FtsWI Activity in Recruitment of both TolA and TolQ. *J* *Bacteriol* 204:e0046421.

32.	Caveney NA, Egan AJF, Ayala I, Laguri C, Robb CS, Breukink E, Vollmer W, Strynadka NCJ, Simorre JP. 2020. Structure of the Peptidoglycan Synthase Activator LpoP in *Pseudomonas* *aeruginosa*. *Structure* 28:643–650 e5.

33.	Egan AJF, Maya-Martinez R, Ayala I, Bougault CM, Banzhaf M, Breukink E, Vollmer W, Simorre JP. 2018. Induced conformational changes activate the peptidoglycan synthase PBP1B. *Mol* *Microbiol* 110:335–356.

34.	Caulton SG, Lambert C, Tyson J, Radford P, Al-Bayati A, Greenwood S, Banks EJ, Clark C, Till R, Pires E, Sockett RE, Lovering AL. 2024. *Bdellovibrio* *bacteriovorus* uses chimeric fibre proteins to recognize and invade a broad range of bacterial hosts. *Nat* *Microbiol* 9:214–227.

35.	Raschdorf O, Forstner Y, Kolinko I, Uebe R, Plitzko JM, Schuler D. 2016. Genetic and Ultrastructural Analysis Reveals the Key Players and Initial Steps of Bacterial Magnetosome Membrane Biogenesis. *PLoS* *Genet* 12:e1006101.

36.	Heijne G. 1986. The distribution of positively charged residues in bacterial inner membrane proteins correlates with the trans-membrane topology. *EMBO* *J* 5:3021–7.

37.	Bajorek M, Schubert HL, McCullough J, Langelier C, Eckert DM, Stubblefield WM, Uter NT, Myszka DG, Hill CP, Sundquist WI. 2009. Structural basis for ESCRT-III protein autoinhibition. *Nat* *Struct* *Mol* *Biol* 16:754–62.

38.	Carlton JG, Baum B. 2023. Roles of ESCRT-III polymers in cell division across the tree of life. *Curr Opin Cell Biol* 85:102274.

39.	Perez-Riverol Y, Bai J, Bandla C, Garcia-Seisdedos D, Hewapathirana S, Kamatchinathan S, Kundu DJ, Prakash A, Frericks-Zipper A, Eisenacher M, Walzer M, Wang S, Brazma A, Vizcaino JA. 2022. The PRIDE database resources in 2022: a hub for mass spectrometry-based proteomics evidences. *Nucleic* *Acids* *Res* 50:D543–D552.

40.	Cameron TA, Margolin W. 2024. Insights into the assembly and regulation of the bacterial divisome. *Nat* *Rev* *Microbiol* 22:33–45.

41.	Hirakawa H, Suzue K, Tomita H. 2022. Roles of the Tol/Pal System in Bacterial Pathogenesis and Its Application to Antibacterial Therapy. *Vaccines* (Basel) 10.

42.	Morrison JJ, Camberg JL. 2024. Building the Bacterial Divisome at the Septum. *Subcell* *Biochem* 104:49–71.

43.	Mehla J, Liechti G, Morgenstein RM, Caufield JH, Hosseinnia A, Gagarinova A, Phanse S, Goodacre N, Brockett M, Sakhawalkar N, Babu M, Xiao R, Montelione GT, Vorobiev S, den Blaauwen T, Hunt JF, Uetz P. 2021. ZapG (YhcB/DUF1043), a novel cell division protein in gamma-proteobacteria linking the Z-ring to septal peptidoglycan synthesis. *J Biol Chem* 296:100700.

44.	Alcorlo M, Martinez-Caballero S, Li J, Sham LT, Luo M, Hermoso JA. 2024. Modulation of the lytic apparatus by the FtsEX complex within the bacterial division machinery. *FEBS* *Lett* 598:2836–2851.

45.	Bertsche U, Kast T, Wolf B, Fraipont C, Aarsman ME, Kannenberg K, von Rechenberg M, Nguyen-Disteche M, den Blaauwen T, Holtje JV, Vollmer W. 2006. Interaction between two murein (peptidoglycan) synthases, PBP3 and PBP1B, in *Escherichia* *coli*. *Mol* *Microbiol* 61:675–90.

46.	Castanheira S, Lopez-Escarpa D, Paradela A, Garcia-Del Portillo F. 2024. *In* *Vivo* Cross-Linking Sheds Light on the *Salmonella* Divisome in Which PBP3 and PBP3(SAL) Compete for Occupancy. *Mol* *Microbiol* 122:797–818.

47.	Spratt BG. 1975. Distinct penicillin binding proteins involved in the division, elongation, and shape of *Escherichia* *coli* K12. *Proc* *Natl* *Acad* *Sci* U S A 72:2999–3003.

48.	Smith JD, Kumarasiri M, Zhang W, Hesek D, Lee M, Toth M, Vakulenko S, Fisher JF, Mobashery S, Chen Y. 2013. Structural analysis of the role of *Pseudomonas* *aeruginosa* penicillin-binding protein 5 in beta-lactam resistance. *Antimicrob* *Agents* *Chemother* 57:3137–46.

49.	Choudhury B, Carlson RW, Goldberg JB. 2005. The structure of the lipopolysaccharide from a *galU* mutant of *Pseudomonas* *aeruginosa* serogroup-O11. *Carbohydr* *Res* 340:2761–72.

50.	Islam S, Oh H, Jalal S, Karpati F, Ciofu O, Hoiby N, Wretlind B. 2009. Chromosomal mechanisms of aminoglycoside resistance in *Pseudomonas* *aeruginosa* isolates from cystic fibrosis patients. *Clin* *Microbiol* *Infect* 15:60–6.

51.	Le S, Yao X, Lu S, Tan Y, Rao X, Li M, Jin X, Wang J, Zhao Y, Wu NC, Lux R, He X, Shi W, Hu F. 2014. Chromosomal DNA deletion confers phage resistance to *Pseudomonas* *aeruginosa*. *Sci* *Rep* 4:4738.

52.	Nakamura K, Fujiki J, Nakamura T, Furusawa T, Gondaira S, Usui M, Higuchi H, Tamura Y, Iwano H. 2021. Fluctuating Bacteriophage-induced *galU* Deficiency Region is Involved in Trade-off Effects on the Phage and Fluoroquinolone Sensitivity in *Pseudomonas* *aeruginosa*. *Virus* *Res* 306:198596.

53.	Tavares IM, Jolly L, Pompeo F, Leitao JH, Fialho AM, Sa-Correia I, Mengin-Lecreulx D. 2000. Identification of the *Pseudomonas* *aeruginosa* *glmM* gene, encoding phosphoglucosamine mutase. *J* *Bacteriol* 182:4453–7.

54.	Jolly L, Ferrari P, Blanot D, Van Heijenoort J, Fassy F, Mengin-Lecreulx D. 1999. Reaction mechanism of phosphoglucosamine mutase from *Escherichia* *coli*. *Eur J Biochem* 262:202–10.

55.	Gruber CC, Babu VMP, Livingston K, Joisher H, Walker GC. 2021. Degradation of the *Escherichia* *coli* Essential Proteins DapB and Dxr Results in Oxidative Stress, which Contributes to Lethality through Incomplete Base Excision Repair. *mBio* 13:e0375621.

56.	Martin C, Cami B, Yeh P, Stragier P, Parsot C, Patte JC. 1988. *Pseudomonas* *aeruginosa* diaminopimelate decarboxylase: evolutionary relationship with other amino acid decarboxylases. *Mol* *Biol* *Evol* 5:549–59.

57.	Wissel MC, Weiss DS. 2004. Genetic analysis of the cell division protein FtsI (PBP3): amino acid substitutions that impair septal localization of FtsI and recruitment of FtsN. *J Bacteriol* 186:490–502.

58.	Vollmer W, Bertsche U. 2008. Murein (peptidoglycan) structure, architecture and biosynthesis in *Escherichia* *coli*. *Biochim* *Biophys* *Acta* 1778:1714–34.

59.	Choi U, Park SH, Lee HB, Son JE, Lee CR. 2023. Coordinated and Distinct Roles of Peptidoglycan Carboxypeptidases DacC and DacA in Cell Growth and Shape Maintenance under Stress Conditions. *Microbiol* *Spectr* 11:e0001423.

60.	Zheng LY, Liu NH, Zhong S, Yu Y, Zhang XY, Qin QL, Song XY, Zhang YZ, Fu H, Wang M, McMinn A, Chen XL, Li PY. 2022. Diaminopimelic Acid Metabolism by *Pseudomonadota* in the Ocean. *Microbiol* *Spectr* 10:e0069122.

61.	Acebron I, Mahasenan KV, De Benedetti S, Lee M, Artola-Recolons C, Hesek D, Wang H, Hermoso JA, Mobashery S. 2017. Catalytic Cycle of the N-Acetylglucosaminidase NagZ from *Pseudomonas* *aeruginosa*. *J* *Am* *Chem* *Soc* 139:6795–6798.

62.	Dhar S, Kumari H, Balasubramanian D, Mathee K. 2018. Cell-wall recycling and synthesis in *Escherichia* *coli* and *Pseudomonas* *aeruginosa* - their role in the development of resistance. *J* *Med* *Microbiol* 67:1–21.

63.	Fragata JI, Rebocho MJ, Melo JQ, Macedo MM. 1985. [Prognostic value of simple chest radiography for the establishment of risk factors in open mitral valve surgery]. *Acta* *Med* *Port* 6:115–20.

64.	Ho LA, Winogrodzki JL, Debowski AW, Madden Z, Vocadlo DJ, Mark BL, Stubbs KA. 2018. A mechanism-based GlcNAc-inspired cyclophellitol inactivator of the peptidoglycan recycling enzyme NagZ reverses resistance to beta-lactams in *Pseudomonas* *aeruginosa*. *Chem* *Commun* (Camb) 54:10630–10633.

65.	Poulsen BE, Yang R, Clatworthy AE, White T, Osmulski SJ, Li L, Penaranda C, Lander ES, Shoresh N, Hung DT. 2019. Defining the core essential genome of *Pseudomonas* *aeruginosa*. *Proc* *Natl* *Acad* *Sci* U S A 116:10072–10080.

66.	Lee SA, Gallagher LA, Thongdee M, Staudinger BJ, Lippman S, Singh PK, Manoil C. 2015. General and condition-specific essential functions of *Pseudomonas* *aeruginosa*. *Proc* *Natl* *Acad* *Sci* U S A 112:5189–94.

67.	Noga MJ, Buke F, van den Broek NJF, Imholz NCE, Scherer N, Yang F, Bokinsky G. 2020. Posttranslational Control of PlsB Is Sufficient To Coordinate Membrane Synthesis with Growth in *Escherichia* *coli*. *mBio* 11.

68.	Yao J, Rock CO. 2013. Phosphatidic acid synthesis in bacteria. *Biochim* *Biophys* *Acta* 1831:495–502.

69.	Masoumi N, Keshavarzi F. 2024. The pattern of antibiotic resistance and distribution of the biofilm-producing *Pseudomonas* *aeruginosa* (PelD, PslB) isolated from infectious hospital departments. *SAGE* *Open* *Med* 12:20503121241298826.

70.	Asadpour L. 2018. Antimicrobial resistance, biofilm-forming ability and virulence potential of *Pseudomonas* *aeruginosa* isolated from burn patients in northern Iran. *J* *Glob* *Antimicrob* *Resist* 13:214–220.

71.	Best EA, Knauf VC. 1993. Organization and nucleotide sequences of the genes encoding the biotin carboxyl carrier protein and biotin carboxylase protein of *Pseudomonas* *aeruginosa* acetyl coenzyme A carboxylase. *J* *Bacteriol* 175:6881–9.

72.	Jeschke M, Ayyolath A, Maniyeri A, Raina S, Klein G. 2025. Coordinated Biosynthesis of Essential Cell Envelope Components: Lipopolysaccharide and Fatty Acids Requires LapD, Acyl Carrier Protein, and Fully Hexaacylated Lipid A. *Int* *J* *Mol* *Sci* 26.

73.	Massengo-Tiasse RP, Cronan JE. 2009. Diversity in enoyl-acyl carrier protein reductases. *Cell* *Mol* *Life* *Sci* 66:1507–17.

74.	Bulai RG, Van Eynde W, Heylen S, Cremonini M, Van Olmen F, Wicik R, Lescrinier E, Rozenski J, Voet A, Masschelein J, Verwilst P. 2025. Investigating the structure-activity-relationship of diaryl ether-based paFabV inhibitors: A path to novel antibacterials*. Eur J Med Chem* 296:117782.

75.	Yuan Y, Leeds JA, Meredith TC. 2012. *Pseudomonas* *aeruginosa* directly shunts beta-oxidation degradation intermediates into de novo fatty acid biosynthesis. *J* *Bacteriol* 194:5185–96.

76.	Prado LG, Camara NOS, Barbosa AS. 2023. Cell lipid biology in infections: an overview. *Front* *Cell* *Infect* *Microbiol* 13:1148383.

77.	Fiedler S, Steinbuchel A, Rehm BH. 2002. The role of the fatty acid beta-oxidation multienzyme complex from *Pseudomonas* *oleovorans* in polyhydroxyalkanoate biosynthesis: molecular characterization of the *fadBA* operon from *P.* *oleovorans* and of the enoyl-CoA hydratase genes *phaJ* from *P*. *oleovorans* and *Pseudomonas* *putida*. *Arch* *Microbiol* 178:149–60.

78.	Bock LJ, Ferguson PM, Clarke M, Pumpitakkul V, Wand ME, Fady PE, Allison L, Fleck RA, Shepherd MJ, Mason AJ, Sutton JM. 2021. *Pseudomonas* *aeruginosa* adapts to octenidine via a combination of efflux and membrane remodelling. *Commun* *Biol* 4:1058.

79.	Yero D, Diaz-Lobo M, Costenaro L, Conchillo-Sole O, Mayo A, Ferrer-Navarro M, Vilaseca M, Gibert I, Daura X. 2021. The *Pseudomonas* *aeruginosa* substrate-binding protein Ttg2D functions as a general glycerophospholipid transporter across the periplasm. *Commun* *Biol* 4:448.

80.	Guest RL, Lee MJ, Wang W, Silhavy TJ. 2023. A periplasmic phospholipase that maintains outer membrane lipid asymmetry in *Pseudomonas aeruginosa*. *Proc Natl Acad Sci* U S A 120:e2302546120.

81.	Sposato D, Mercolino J, Torrini L, Sperandeo P, Lucidi M, Alegiani R, Varone I, Molesini G, Leoni L, Rampioni G, Visca P, Imperi F. 2024. Redundant essentiality of AsmA-like proteins in *Pseudomonas* *aeruginosa*. *mSphere* 9:e0067723.

82.	de Kievit TR, Lam JS. 1997. Isolation and characterization of two genes, *waaC* (*rfaC*) and *waaF* (*rfaF*), involved in *Pseudomonas* *aeruginosa* serotype O5 inner-core biosynthesis. *J* *Bacteriol* 179:3451–7.

83.	Wang Z, Wang J, Ren G, Li Y, Wang X. 2015. Influence of Core Oligosaccharide of Lipopolysaccharide to Outer Membrane Behavior of *Escherichia* *coli*. *Mar* *Drugs* 13:3325–39.

84.	Hofstaedter CE, Chandler CE, Met CM, Gillespie JJ, Harro JM, Goodlett DR, Rasko DA, Ernst RK. 2024. Divergent *Pseudomonas* *aeruginosa* LpxO enzymes perform site-specific lipid A 2-hydroxylation. *mBio* 15:e0282323.

85.	Schultz KM, Schneider JR, Fischer MA, Cina NP, Riegert MO, Frank DW, Klug CS. 2023. Binding and transport of LPS occurs through the coordinated combination of an array of sites across the entire *Escherichia* *coli* LPS transport protein LptA. *Protein* *Sci* 32:e4724.

86.	Kaplan E, Greene NP, Jepson AE, Koronakis V. 2022. Structural basis of lipoprotein recognition by the bacterial Lol trafficking chaperone LolA. *Proc* *Natl* *Acad* *Sci* U S A 119:e2208662119.

87.	Lloubes R, Cascales E, Walburger A, Bouveret E, Lazdunski C, Bernadac A, Journet L. 2001. The Tol-Pal proteins of the *Escherichia* *coli* cell envelope: an energized system required for outer membrane integrity? *Res* *Microbiol* 152:523–9.

88.	Celia H, Botos I, Ghirlando R, Duche D, Beach BM, Lloubes R, Buchanan SK. 2025. Cryo-EM structures of the *E. coli* Ton and Tol motor complexes. *Nat* *Commun* 16:5506.

89.	Einenkel R, Halte M, Sawant SA, Erhardt M, Wadhwa N, Popp PF. 2025. Building the bacterial flagellum: coordinating regulation, dynamic assembly, and function. *Microbiol* *Mol* *Biol* *Rev* 89:e0009222.

90.	Schmidt LM, Pulido-Sanchez M, Treuner-Lange A, Zehner L, Lopez-Sanchez A, Cava F, Govantes F, Thormann KM. 2026. Functional characterization of the polar organizer protein FimV in *Pseudomonas* *putida*. *J* *Bacteriol* 208:e0049725.

91.	Pulido-Sanchez M, Leal-Morales A, Lopez-Sanchez A, Cava F, Govantes F. 2025. Spatial, temporal and numerical regulation of polar flagella assembly in *Pseudomonas* *putida*. *Microbiol* *Res* 292:128033.

92.	Dornes A, Schmidt LM, Mais CN, Hook JC, Pane-Farre J, Kressler D, Thormann K, Bange G. 2024. Polar confinement of a macromolecular machine by an SRP-type GTPase. *Nat* *Commun* 15:5797.

93.	Yarrington KD, Limoli DH. 2025. The type IV pilus steering committee: how Pil-Chp controls directional motility. *J* *Bacteriol* 207:e0039624.

94.	Herianto S, Lee HM, Chen C, Ho JA, Tu HL. 2025. Functional reconstitution of bacterial ESCRT-III protein PspA identifies key regions in membrane binding and remodeling. *Protein* *Sci* 34:e70309.

95.	Herianto S, Tu HL. 2026. Recent insights into bacterial ESCRT-III-mediated membrane remodeling. *Crit* *Rev* *Biochem* *Mol* *Biol* doi:10.1080/10409238.2025.2606239:1–19.

96.	Hegde RS, Keenan RJ. 2024. A unifying model for membrane protein biogenesis. *Nat Struct Mol Biol* 31:1009–1017.

97.	Jauss B, Petriman NA, Drepper F, Franz L, Sachelaru I, Welte T, Steinberg R, Warscheid B, Koch HG. 2019. Noncompetitive binding of PpiD and YidC to the SecYEG translocon expands the global view on the SecYEG interactome in *Escherichia* *coli*. *J* *Biol* *Chem* 294:19167–19183.

98.	Ahator SD, Zhang L. 2019. Small Is Mighty-Chemical Communication Systems in *Pseudomonas* *aeruginosa*. *Annu* *Rev* *Microbiol* 73:559–578.

99.	Khan MA, Zhu L, Zhu H. 2026. Peptide-based approaches to quorum-sensing disruption: emerging trends and applications in antimicrobial therapy. *Bioorg* *Med* *Chem* 133:118496.

100.	Pletzer D, Lafon C, Braun Y, Kohler T, Page MG, Mourez M, Weingart H. 2014. High-throughput screening of dipeptide utilization mediated by the ABC transporter DppBCDF and its substrate-binding proteins DppA1-A5 in *Pseudomonas* *aeruginosa*. *PLoS* *One* 9:e111311.

101.	Plochl K, Bottcher T. 2025. The solute-binding proteins DppA1-5 of *Pseudomonas* *aeruginosa* have distinct substrate profiles. *Sci* *Rep* 15:44346.

102.	Fernandez M, Rico-Jimenez M, Ortega A, Daddaoua A, Garcia Garcia AI, Martin-Mora D, Torres NM, Tajuelo A, Matilla MA, Krell T. 2019. Determination of Ligand Profiles for *Pseudomonas* *aeruginosa* Solute Binding Proteins. *Int* *J* *Mol* *Sci* 20.

103.	Luengo JM, Olivera ER. 2020. Catabolism of biogenic amines in *Pseudomonas* species. *Environ* *Microbiol* 22:1174–1192.

104.	Morgan RN, El-Behery RR. 2025. *Pseudomonas* *aeruginosa* *Quorum* Signals; Associations with Virulence, and Impact of Therapeutic Gamma Radiation Doses. *Curr* *Microbiol* 82:255.

105.	Hasan CM, Pottenger S, Green AE, Cox AA, White JS, Jones T, Winstanley C, Kadioglu A, Wright MH, Neill DR, Fothergill JL. 2022. *Pseudomonas* *aeruginosa* utilizes the host-derived polyamine spermidine to facilitate antimicrobial tolerance. *JCI* *Insight* 7.

106.	Lin Q, Wang H, Huang J, Liu Z, Chen Q, Yu G, Xu Z, Cheng P, Liang Z, Zhang LH. 2022. Spermidine Is an Intercellular Signal Modulating T3SS Expression in *Pseudomonas* *aeruginosa*. *Microbiol* *Spectr* 10:e0064422.

107.	Nishijyo T, Park SM, Lu CD, Itoh Y, Abdelal AT. 1998. Molecular characterization and regulation of an operon encoding a system for transport of arginine and ornithine and the ArgR regulatory protein in *Pseudomonas* *aeruginosa*. *J* *Bacteriol* 180:5559–66.

108.	Scribani Rossi C, Barrientos-Moreno L, Paone A, Cutruzzola F, Paiardini A, Espinosa-Urgel M, Rinaldo S. 2022. Nutrient Sensing and Biofilm Modulation: The Example of L-arginine in *Pseudomonas*. *Int* *J* *Mol* *Sci* 23.

109.	Liu Z, Hossain SS, Morales Moreira Z, Haney CH. 2022. Putrescine and Its Metabolic Precursor Arginine Promote Biofilm and c-di-GMP Synthesis in *Pseudomonas* *aeruginosa*. *J* *Bacteriol* 204:e0029721.

110.	Hoshino T, Kose-Terai K, Sato K. 1992. Solubilization and reconstitution of the *Pseudomonas* *aeruginosa* high affinity branched-chain amino acid transport system. *J* *Biol* *Chem* 267:21313–8.

111.	Reyes-Darias JA, Yang Y, Sourjik V, Krell T. 2015. Correlation between signal input and output in PctA and PctB amino acid chemoreceptor of *Pseudomonas* *aeruginosa*. *Mol* *Microbiol* 96:513–25.

112.	Rico-Jimenez M, Munoz-Martinez F, Garcia-Fontana C, Fernandez M, Morel B, Ortega A, Ramos JL, Krell T. 2013. Paralogous chemoreceptors mediate chemotaxis towards protein amino acids and the non-protein amino acid gamma-aminobutyrate (GABA). *Mol* *Microbiol* 88:1230–43.

113.	Chen C, Yan Q, Tao M, Shi H, Han X, Jia L, Huang Y, Zhao L, Wang C, Ma X, Ma Y. 2019. Characterization of serine acetyltransferase (CysE) from methicillin-resistant *Staphylococcus* *aureus* and inhibitory effect of two natural products on CysE. *Microb* *Pathog* 131:218–226.

114.	Oldham KEA, Prentice EJ, Summers EL, Hicks JL. 2022. Serine acetyltransferase from *Neisseria* *gonorrhoeae*; structural and biochemical basis of inhibition. *Biochem* *J* 479:57–74.

115.	Toyomoto T, Ono K, Shiba T, Momitani K, Zhang T, Tsutsuki H, Ishikawa T, Hoso K, Hamada K, Rahman A, Wen L, Maeda Y, Yamamoto K, Matsuoka M, Hanaoka K, Niidome T, Akaike T, Sawa T. 2023. Alkyl gallates inhibit serine O-acetyltransferase in bacteria and enhance susceptibility of drug-resistant Gram-negative bacteria to antibiotics. *Front* *Microbiol* 14:1276447.

116.	Benoni R, De Bei O, Paredi G, Hayes CS, Franko N, Mozzarelli A, Bettati S, Campanini B. 2017. Modulation of *Escherichia* *coli* serine acetyltransferase catalytic activity in the cysteine synthase complex. *FEBS* *Lett* 591:1212–1224.

117.	Sarwar Z, Lundgren BR, Grassa MT, Wang MX, Gribble M, Moffat JF, Nomura CT. 2016. GcsR, a TyrR-Like Enhancer-Binding Protein, Regulates Expression of the Glycine Cleavage System in *Pseudomonas* *aeruginosa* PAO1. *mSphere* 1.

118.	Miggiano R, Martignon S, Minassi A, Rossi F, Rizzi M. 2020. Crystal structure of *Haemophilus* *influenzae* 3-isopropylmalate dehydrogenase (LeuB) in complex with the inhibitor O-isobutenyl oxalylhydroxamate. *Biochem* *Biophys* *Res* *Commun* 524:996–1002.

119.	Hoang TT, Schweizer HP. 1997. Identification and genetic characterization of the *Pseudomonas* *aeruginosa* *leuB* gene encoding 3-isopropylmalate dehydrogenase. *Mol* *Gen* *Genet* 254:166–70.

120.	Wanders RJ, Duran M, Loupatty FJ. 2012. Enzymology of the branched-chain amino acid oxidation disorders: the valine pathway. *J Inherit Metab Dis* 35:5–12.

121.	Forster-Fromme K, Jendrossek D. 2008. Biochemical characterization of isovaleryl-CoA dehydrogenase (LiuA) of *Pseudomonas* *aeruginosa* and the importance of *liu* genes fora functional catabolic pathway of methyl-branched compounds. *FEMS* *Microbiol* *Lett* 286:78–84.

122.	Panina EM, Vitreschak AG, Mironov AA, Gelfand MS. 2001. Regulation of aromatic amino acid biosynthesis in gamma-proteobacteria. *J Mol Microbiol Biotechnol* 3:529–43.

123.	De Groote VN, Verstraeten N, Fauvart M, Kint CI, Verbeeck AM, Beullens S, Cornelis P, Michiels J. 2009. Novel persistence genes in *Pseudomonas* *aeruginosa* identified by high-throughput screening. *FEMS* *Microbiol* *Lett* 297:73–9.

124.	Nishikura-Imamura S, Matsutani M, Insomphun C, Vangnai AS, Toyama H, Yakushi T, Abe T, Adachi O, Matsushita K. 2014. Overexpression of a type II 3-dehydroquinate dehydratase enhances the biotransformation of quinate to 3-dehydroshikimate in *Gluconobacter* *oxydans*. *Appl* *Microbiol* *Biotechnol* 98:2955–63.

125.	Gu W, Song J, Bonner CA, Xie G, Jensen RA. 1998. PhhC is an essential aminotransferase for aromatic amino acid catabolism in *Pseudomonas* *aeruginosa*. *Microbiology* (Reading) 144 ( Pt 11):3127–3134.

126.	Oetiker JH, Lee DH, Kato A. 1999. Molecular analysis of a tryptophan-2-monooxygenase gene (*IaaM*) of *Agrobacterium* *vitis*. *DNA* *Seq* 10:349–54.

127.	Ralph EC, Anderson MA, Cleland WW, Fitzpatrick PF. 2006. Mechanistic studies of the flavoenzyme tryptophan 2-monooxygenase: deuterium and 15N kinetic isotope effects on alanine oxidation by an L-amino acid oxidase. *Biochemistry* 45:15844–52.

128.	Wang L, Ding B, Hu X, Li G, Deng Y. 2024. Rationally Engineering pH Adaptation of Acid-Induced Arginine Decarboxylase from *Escherichia* *coli* to Alkaline Environments to Efficiently Biosynthesize Putrescine. *Adv* *Sci* (Weinh) 11:e2307779.

129.	Richard HT, Foster JW. 2003. Acid resistance in *Escherichia* *coli*. *Adv* *Appl* *Microbiol* 52:167–86.

130.	Chan YC, Blaschek HP. 2005. Comparative analysis of *Shigella* *boydii* 18 foodborne outbreak isolate and related enteric bacteria: role of *rpoS* and *adiA* in acid stress response. *J* *Food* *Prot* 68:521–7.

131.	Zhao G, Winkler ME. 1995. Kinetic limitation and cellular amount of pyridoxine (pyridoxamine) 5'-phosphate oxidase of *Escherichia* *coli* K-12. *J* *Bacteriol* 177:883–91.

132.	O'Neal L, Baraquet C, Suo Z, Dreifus JE, Peng Y, Raivio TL, Wozniak DJ, Harwood CS, Parsek MR. 2022. The Wsp system of *Pseudomonas* *aeruginosa* links surface sensing and cell envelope stress. *Proc* *Natl* *Acad* *Sci* U S A 119:e2117633119.

133.	Kessler C, Mhatre E, Cooper V, Kim W. 2021. Evolutionary Divergence of the Wsp Signal Transduction Systems in Beta- and Gammaproteobacteria. *Appl* *Environ* *Microbiol* 87:e0130621.

134.	Elhosary MA, Bahey-El-Din M, AbdelBary A, El Guink N, Aboushleib HM. 2019. Immunization with the ferric iron-binding periplasmic protein HitA provides protection against *Pseudomonas* *aeruginosa* in the murine infection model. *Microb* *Pathog* 131:181–185.

135.	Li P, Wang X, Sun X, Guan Z, Sun W. 2021. Outer Membrane Vesicles Displaying a Heterologous PcrV-HitA Fusion Antigen Promote Protection against Pulmonary *Pseudomonas* *aeruginosa* Infection. *mSphere* 6:e0069921.

136.	Parker Siburt CJ, Mietzner TA, Crumbliss AL. 2012. FbpA--a bacterial transferrin with more to offer. *Biochim* *Biophys* *Acta* 1820:379–92.

137.	Kirby SD, Gray-Owen SD, Schryvers AB. 1997. Characterization of a ferric-binding protein mutant in *Haemophilus* *influenzae*. *Mol* *Microbiol* 25:979–87.

138.	Coffey BM, Akhand SS, Anderson GG. 2014. MgtE is a dual-function protein in *Pseudomonas* *aeruginosa*. *Microbiology* (Reading) 160:1200–1213.

139.	Chakravarty S, Melton CN, Bailin A, Yahr TL, Anderson GG. 2017. *Pseudomonas* *aeruginosa* Magnesium Transporter MgtE Inhibits Type III Secretion System Gene Expression by Stimulating *rsmYZ* Transcription. *J* *Bacteriol* 199.

140.	Anderson GG, Yahr TL, Lovewell RR, O'Toole GA. 2010. The *Pseudomonas* *aeruginosa* magnesium transporter MgtE inhibits transcription of the type III secretion system. *Infect* *Immun* 78:1239–49.

141.	Madhusudhan KT, McLaughlin R, Komori N, Matsumoto H. 2003. Identification of a major protein upon phosphate starvation of Pseudomonas aeruginosa PAO1. J Basic Microbiol 43:36–46.

142.	Zaborina O, Holbrook C, Chen Y, Long J, Zaborin A, Morozova I, Fernandez H, Wang Y, Turner JR, Alverdy JC. 2008. Structure-function aspects of PstS in multi-drug-resistant *Pseudomonas* *aeruginosa*. *PLoS* *Pathog* 4:e43.

143.	Shah M, Zaborin A, Alverdy JC, Scott K, Zaborina O. 2014. Localization of DING proteins on PstS-containing outer-surface appendages of *Pseudomonas* *aeruginosa*. *FEMS* *Microbiol* *Lett* 352:54–61.

144.	Llanes C, Hocquet D, Vogne C, Benali-Baitich D, Neuwirth C, Plesiat P. 2004. Clinical strains of *Pseudomonas* *aeruginosa* overproducing MexAB-OprM and MexXY efflux pumps simultaneously. *Antimicrob* *Agents* *Chemother* 48:1797–802.

145.	Li C, Edwards MD, Jeong H, Roth J, Booth IR. 2007. Identification of mutations that alter the gating of the *Escherichia* *coli* mechanosensitive channel protein, MscK. *Mol* *Microbiol* 64:560–74.

146.	Edwards MD, Black S, Rasmussen T, Rasmussen A, Stokes NR, Stephen TL, Miller S, Booth IR. 2012. Characterization of three novel mechanosensitive channel activities in *Escherichia* *coli*. *Channels* (Austin) 6:272–81.

147.	Neumann-Schaal M, Acuna JMB, Jahn D. 2026. Strict Aerobic Lifestyle and Anaerobic Survival of Bacteria: Inseparable Twins? *Mol* *Microbiol* doi:10.1111/mmi.70071.

148.	Hreha TN, Foreman S, Duran-Pinedo A, Morris AR, Diaz-Rodriguez P, Jones JA, Ferrara K, Bourges A, Rodriguez L, Koffas MAG, Hahn M, Hauser AR, Barquera B. 2021. The three NADH dehydrogenases of *Pseudomonas* *aeruginosa*: Their roles in energy metabolism and links to virulence. *PLoS* *One* 16:e0244142.

149.	Torres A, Kasturiarachi N, DuPont M, Cooper VS, Bomberger J, Zemke A. 2019. NADH Dehydrogenases in *Pseudomonas* *aeruginosa* Growth and Virulence. *Front* *Microbiol* 10:75.

150.	Kretzschmar U, Ruckert A, Jeoung JH, Gorisch H. 2002. Malate:quinone oxidoreductase is essential for growth on ethanol or acetate in *Pseudomonas* *aeruginosa*. *Microbiology* (Reading) 148:3839–3847.

151.	Williams HD, Zlosnik JE, Ryall B. 2007. Oxygen, cyanide and energy generation in the cystic fibrosis pathogen *Pseudomonas* *aeruginosa*. *Adv* *Microb* *Physiol* 52:1–71.

152.	Glasser NR, Saunders SH, Newman DK. 2017. The Colorful World of Extracellular Electron Shuttles. *Annu* *Rev* *Microbiol* 71:731–751.

153.	Ciemniecki JA, Newman DK. 2023. NADH dehydrogenases are the predominant phenazine reductases in the electron transport chain of *Pseudomonas* *aeruginosa*. *Mol* *Microbiol* 119:560–573.

154.	Parsons JF, Calabrese K, Eisenstein E, Ladner JE. 2003. Structure and mechanism of *Pseudomonas* *aeruginosa* PhzD, an isochorismatase from the phenazine biosynthetic pathway. *Biochemistry* 42:5684–93.

155.	Riegerova P, Horvath M, Sebesta F, Sykora J, Sulc M, Vlcek A. 2024. Single-step purification and characterization of *Pseudomonas* *aeruginosa* azurin. *Protein* *Expr* *Purif* 224:106566.

156.	Yaghoubi A, Khazaei M, Avan A, Hasanian SM, Cho WC, Soleimanpour S. 2020. p28 Bacterial Peptide, as an Anticancer Agent. *Front* *Oncol* 10:1303.

157.	Huang F, Shu Q, Qin Z, Tian J, Su Z, Huang Y, Gao M. 2020. Anticancer Actions of Azurin and Its Derived Peptide p28. *Protein* *J* 39:182–189.

158.	Preisig O, Zufferey R, Hennecke H. 1996. The *Bradyrhizobium* *japonicum* *fixGHIS* genes are required for the formation of the high-affinity cbb3-type cytochrome oxidase. *Arch* *Microbiol* 165:297–305.

159.	Deshmukh M, May M, Zhang Y, Gabbert KK, Karberg KA, Kranz RG, Daldal F. 2002. Overexpression of *ccl1*-2 can bypass the need for the putative apocytochrome chaperone CycH during the biogenesis of c-type cytochromes. *Mol* *Microbiol* 46:1069–80.

160.	Geng Z, Ge J, Cui W, Zhou H, Deng J, Xu B. 2022. Efficient *De* *Novo* Biosynthesis of Heme by Membrane Engineering in *Escherichia* *coli*. *Int* *J* *Mol* *Sci* 23.

161.	Sasarman A, Echelard Y, Letowski J, Tardif D, Drolet M. 1988. Nucleotide sequence of the *hemX* gene, the third member of the Uro operon of *Escherichia* *coli* K12. *Nucleic* *Acids* *Res* 16:11835.

162.	Vo CD, Michaud J, Elsen S, Faivre B, Bouveret E, Barras F, Fontecave M, Pierrel F, Lombard M, Pelosi L. 2020. The O(2)-independent pathway of ubiquinone biosynthesis is essential for denitrification in *Pseudomonas* *aeruginosa*. *J* *Biol* *Chem* 295:9021–9032.

163.	Lee PT, Hsu AY, Ha HT, Clarke CF. 1997. A C-methyltransferase involved in both ubiquinone and menaquinone biosynthesis: isolation and identification of the *Escherichia* *coli* *ubiE* gene. *J* *Bacteriol* 179:1748–54.

164.	Abby SS, Kazemzadeh K, Vragniau C, Pelosi L, Pierrel F. 2020. Advances in bacterial pathways for the biosynthesis of ubiquinone. *Biochim* *Biophys* *Acta* *Bioenerg* 1861:148259.

165.	Jurgenson CT, Ealick SE, Begley TP. 2009. Biosynthesis of Thiamin Pyrophosphate. *EcoSal* *Plus* 3.

166.	Nemoto N, Baba S, Kawai G, Sampei GI. 2024. Crystal structure of guanosine 5'-monophosphate synthetase from the thermophilic bacterium *Thermus* *thermophilus* HB8. *Acta* *Crystallogr* *F* *Struct* *Biol* *Commun* 80:278–285.

167.	Goncheva MI, Chin D, Heinrichs DE. 2022. Nucleotide biosynthesis: the base of bacterial pathogenesis. *Trends Microbiol* 30:793–804.

168.	Villela AD, Sanchez-Quitian ZA, Ducati RG, Santos DS, Basso LA. 2011. Pyrimidine salvage pathway in *Mycobacterium* *tuberculosis*. *Curr* *Med* *Chem* 18:1286–98.

169.	Toussaint JP, Farrell-Sherman A, Feldman TP, Smalley NE, Schaefer AL, Greenberg EP, Dandekar AA. 2017. Gene Duplication in *Pseudomonas* *aeruginosa* Improves Growth on Adenosine. *J* *Bacteriol* 199.

170.	Heurlier K, Denervaud V, Haenni M, Guy L, Krishnapillai V, Haas D. 2005. Quorum-sensing-negative (*lasR*) mutants of *Pseudomonas* *aeruginosa* avoid cell lysis and death. *J* *Bacteriol* 187:4875–83.

171.	Yee TW, Smith DW. 1990. *Pseudomonas* chromosomal replication origins: a bacterial class distinct from *Escherichia* *coli*-type origins. *Proc* *Natl* *Acad* *Sci* U S A 87:1278–82.

172.	Mooney RA, Zhu J, Saba J, Landick R. 2025. NusG-Spt5 Transcription Factors: Universal, Dynamic Modulators of Gene Expression. *J* *Mol* *Biol* 437:168814.

173.	Delbeau M, Froom R, Landick R, Darst SA, Campbell EA. 2024. The yin and yang of the universal transcription factor NusG. *Curr* *Opin* *Microbiol* 81:102540.

174.	Brewer JJ, Inlow K, Mooney RA, Bosch B, Olinares PDB, Marcelino LP, Chait BT, Landick R, Gelles J, Campbell EA, Darst SA. 2025. RapA opens the RNA polymerase clamp to disrupt post-termination complexes and prevent cytotoxic R-loop formation. *Nat* *Struct* *Mol* *Biol* 32:639–649.

175.	Jorgensen F, Bally M, Chapon-Herve V, Michel G, Lazdunski A, Williams P, Stewart G. 1999. RpoS-dependent stress tolerance in *Pseudomonas* *aeruginosa*. *Microbiology* (Reading) 145 ( Pt 4):835–844.

176.	Muriel-Millan LF, Montelongo-Martinez LF, Gonzalez-Valdez A, Bedoya-Perez LP, Cocotl-Yanez M, Soberon-Chavez G. 2024. The alternative sigma factor RpoS regulates *Pseudomonas* *aeruginosa* *quorum* *sensing* response by repressing the *pqsABCDE* operon and activating *vfr*. *Mol* *Microbiol* 121:291–303.

177.	Montelongo-Martinez LF, Hernandez-Mendez C, Muriel-Millan LF, Hernandez-Estrada R, Fabian-Del Olmo MJ, Gonzalez-Valdez A, Soberon-Chavez G, Cocotl-Yanez M. 2023. Unraveling the regulation of pyocyanin synthesis by RsmA through MvaU and RpoS in *Pseudomonas* *aeruginosa* ID4365. *J* *Basic* *Microbiol* 63:51–63.

178.	He Q, Feng Z, Wang Y, Wang K, Zhang K, Kai L, Hao X, Yu Z, Chen L, Ge Y. 2019. LasR Might Act as an Intermediate in Overproduction of Phenazines in the Absence of RpoS in *Pseudomonas* *aeruginosa*. *J* *Microbiol* *Biotechnol* 29:1299–1309.

179.	Trunk K, Benkert B, Quack N, Munch R, Scheer M, Garbe J, Jansch L, Trost M, Wehland J, Buer J, Jahn M, Schobert M, Jahn D. 2010. Anaerobic adaptation in *Pseudomonas* *aeruginosa*: definition of the Anr and Dnr regulons. *Environ* *Microbiol* 12:1719–33.

180.	Schreiber K, Krieger R, Benkert B, Eschbach M, Arai H, Schobert M, Jahn D. 2007. The anaerobic regulatory network required for *Pseudomonas* *aeruginosa* nitrate respiration. *J* *Bacteriol* 189:4310–4.

181.	Eschbach M, Schreiber K, Trunk K, Buer J, Jahn D, Schobert M. 2004. Long-term anaerobic survival of the opportunistic pathogen *Pseudomonas* *aeruginosa* via pyruvate fermentation. *J* *Bacteriol* 186:4596–604.

182.	Chen G, Fanouraki G, Anandhi Rangarajan A, Winkelman BT, Winkelman JT, Waters CM, Mukherjee S. 2024. Combinatorial control of *Pseudomonas* *aeruginosa* biofilm development by *quorum*-*sensing* and nutrient-sensing regulators. *mSystems* 9:e0037224.

183.	Thaden JT, Lory S, Gardner TS. 2010. Quorum-sensing regulation of a copper toxicity system in *Pseudomonas* *aeruginosa*. *J* *Bacteriol* 192:2557–68.

184.	Raimunda D, Padilla-Benavides T, Vogt S, Boutigny S, Tomkinson KN, Finney LA, Arguello JM. 2013. Periplasmic response upon disruption of transmembrane Cu transport in *Pseudomonas* *aeruginosa*. *Metallomics* 5:144–51.

185.	Alarcon Rios AC, Zwep LB, Vriesendorp B, Knibbe CAJ, Ciofu O, Liakopoulos A, Aulin LBS, Rozen DE, van Hasselt JGC. 2025. Colistin resistance dynamics in *Pseudomonas* *aeruginosa* under biofilm and planktonic growth. *Antimicrob* *Agents* *Chemother* 69:e0042125.

186.	Yang B, Liu C, Pan X, Fu W, Fan Z, Jin Y, Bai F, Cheng Z, Wu W. 2021. Identification of Novel PhoP-PhoQ Regulated Genes That Contribute to Polymyxin B Tolerance in *Pseudomonas* *aeruginosa*. *Microorganisms* 9.

187.	Miryala SK, Anbarasu A, Ramaiah S. 2019. Systems biology studies in *Pseudomonas* *aeruginosa* PA01 to understand their role in biofilm formation and multidrug efflux pumps. *Microb* *Pathog* 136:103668.

188.	Qadi M, Izquierdo-Rabassa S, Mateu Borras M, Domenech-Sanchez A, Juan C, Goldberg JB, Hancock REW, Alberti S. 2017. Sensing Mg(2+) contributes to the resistance of *Pseudomonas* *aeruginosa* to complement-mediated opsonophagocytosis. *Environ* *Microbiol* 19:4278–4286.

189.	Chen L, Duan K. 2016. A PhoPQ-Regulated ABC Transporter System Exports Tetracycline in *Pseudomonas* *aeruginosa*. *Antimicrob* *Agents* *Chemother* 60:3016–24.

190.	Wilton M, Charron-Mazenod L, Moore R, Lewenza S. 2016. Extracellular DNA Acidifies Biofilms and Induces Aminoglycoside Resistance in *Pseudomonas* *aeruginosa*. *Antimicrob* *Agents* *Chemother* 60:544–53.

191.	Gooderham WJ, Gellatly SL, Sanschagrin F, McPhee JB, Bains M, Cosseau C, Levesque RC, Hancock REW. 2009. The sensor kinase PhoQ mediates virulence in *Pseudomonas* *aeruginosa*. *Microbiology* (Reading) 155:699–711.

192.	Gellatly SL, Needham B, Madera L, Trent MS, Hancock RE. 2012. The *Pseudomonas* *aeruginosa* PhoP-PhoQ two-component regulatory system is induced upon interaction with epithelial cells and controls cytotoxicity and inflammation. *Infect* *Immun* 80:3122–31.

193.	Roux C, Ramos-Hue M, Audonnet M, Duviau MP, Nouaille S, Carpousis AJ, Laguerre S, Hajnsdorf E, Cocaign-Bousquet M, Girbal L. 2025. RNA stability is regulated by both RNA polyadenylation and ATP levels, linking RNA and energy metabolisms in *Escherichia* *coli*. *mBio* 16:e0268024.

194.	Maes A, Gracia C, Innocenti N, Zhang K, Aurell E, Hajnsdorf E. 2017. Landscape of RNA polyadenylation in *E. coli*. *Nucleic* *Acids* *Res* 45:2746–2756.

195.	Zhang L, Zhang S, Wang J, Zhang Z, Xu Y, Zhang C. 2025. The poly (A) polymerase *pcnB* modulates virulence and resistance in *Klebsiella* *pneumoniae* by differentially regulating chromosomal mRNA stability and plasmid copy number. *Front* *Microbiol* 16:1709161.

196.	Frisch T, Geiser P, Komi M, Karlsson PA, Bhetwal A, Jenniches L, Barquist L, Holmqvist E, Mateus A, Sellin ME, Di Martino ML. 2025. The *pcnB* gene sustains *Shigella* *flexneri* virulence. *PLoS* *Pathog* 21:e1013727.

197.	Francis N, Behera MR, Natarajan K, Laishram RS. 2023. Tyrosine phosphorylation controlled poly(A) polymerase I activity regulates general stress response in bacteria. *Life* *Sci* *Alliance* 6.

198.	Hausmann S, Gonzalez D, Geiser J, Valentini M. 2021. The DEAD-box RNA helicase RhlE2 is a global regulator of *Pseudomonas* *aeruginosa* lifestyle and pathogenesis. *Nucleic* *Acids* *Res* 49:6925–6940.

199.	Wilson DN, Nierhaus KH. 2007. The weird and wonderful world of bacterial ribosome regulation. *Crit* *Rev* *Biochem* *Mol* *Biol* 42:187–219.

200.	Shen YL, Xu L, Zhou Y, Ye BC. 2025. Bacterial ribosome heterogeneity facilitates rapid response to stress. *J Bacteriol* 207:e0005825.

201.	Hwang J, Inouye M. 2010. A bacterial GAP-like protein, YihI, regulating the GTPase of Der, an essential GTP-binding protein in *Escherichia* *coli*. *J* *Mol* *Biol* 399:759–72.

202.	Garaeva NS, Bikmullin AG, Fatkhullin BF, Validov SZ, Keiffer B, Yusupov MM, Usachev KS. 2022. Backbone and side chain NMR assignments for the ribosome maturation factor P (RimP) from *Staphylococcus* *aureus*. *Biomol* *NMR* *Assign* 16:373–377.

203.	Maksimova E, Kravchenko O, Korepanov A, Stolboushkina E. 2022. Protein Assistants of Small Ribosomal Subunit Biogenesis in Bacteria. *Microorganisms* 10.

204.	Shajani Z, Sykes MT, Williamson JR. 2011. Assembly of bacterial ribosomes. Annu Rev Biochem 80:501–26.

205.	Chen B, Boel G, Hashem Y, Ning W, Fei J, Wang C, Gonzalez RL, Jr., Hunt JF, Frank J. 2014. EttA regulates translation by binding the ribosomal E site and restricting ribosome-tRNA dynamics. *Nat* *Struct* *Mol* *Biol* 21:152–9.

206.	Ousalem F, Ngo S, Oiffer T, Omairi-Nasser A, Hamon M, Monlezun L, Boel G. 2024. Global regulation via modulation of ribosome pausing by the ABC-F protein EttA. *Nat* *Commun* 15:6314.

207.	Boel G, Smith PC, Ning W, Englander MT, Chen B, Hashem Y, Testa AJ, Fischer JJ, Wieden HJ, Frank J, Gonzalez RL, Jr., Hunt JF. 2014. The ABC-F protein EttA gates ribosome entry into the translation elongation cycle. *Nat* *Struct* *Mol* *Biol* 21:143–51.

208.	Ohnishi M, Janosi L, Shuda M, Matsumoto H, Hayashi T, Terawaki Y, Kaji A. 1999. Molecular cloning, sequencing, purification, and characterization of *Pseudomonas* *aeruginosa* ribosome recycling factor. *J* *Bacteriol* 181:1281–91.

209.	Kashimori H, Yoshida T, Kijima H, Shimahara H, Uchiyama S, Ishino T, Shuda M, Nakano H, Shibata Y, Saihara Y, Ohkubo T, Yoshida T, Kaji A, Kobayashi Y. 1999. Backbone NMR assignment and secondary structure of ribosome recycling factor (RRF) from *Pseudomonas* *aeruginosa*. *J* *Biomol* *NMR* 15:341–2.

210.	Janosi L, Mori H, Sekine Y, Abragan J, Janosi R, Hirokawa G, Kaji A. 2000. Mutations influencing the frr gene coding for ribosome recycling factor (RRF). *J Mol Biol* 295:815–29.

211.	Suzuki T, Nakamura A, Kato K, Soll D, Tanaka I, Sheppard K, Yao M. 2015. Structure of the *Pseudomonas* *aeruginosa* transamidosome reveals unique aspects of bacterial tRNA-dependent asparagine biosynthesis. *Proc* *Natl* *Acad* *Sci* U S A 112:382–7.

212.	Bernard D, Akochy PM, Bernier S, Fisette O, Brousseau OC, Chenevert R, Roy PH, Lapointe J. 2007. Inhibition by L-aspartol adenylate of a nondiscriminating aspartyl-tRNA synthetase reveals differences between the interactions of its active site with tRNA(Asp) and tRNA(Asn). *J Enzyme Inhib Med Chem* 22:77–82.

213.	Bernard D, Akochy PM, Beaulieu D, Lapointe J, Roy PH. 2006. Two residues in the anticodon recognition domain of the aspartyl-tRNA synthetase from *Pseudomonas* *aeruginosa* are individually implicated in the recognition of tRNAAsn. *J* *Bacteriol* 188:269–74.

214.	Fereshteh S, Noori Goodarzi N, Kalhor H, Rahimi H, Barzi SM, Badmasti F. 2023. Identification of Putative Drug Targets in Highly Resistant Gram-Negative Bacteria; and Drug Discovery Against Glycyl-tRNA Synthetase as a New Target. *Bioinform* *Biol* *Insights* 17:11779322231152980.

215.	Corona A, Palmer SO, Zamacona R, Mendez B, Dean FB, Bullard JM. 2018. Discovery and Characterization of Chemical Compounds That Inhibit the Function of Aspartyl-tRNA Synthetase from *Pseudomonas* *aeruginosa*. *SLAS* *Discov* 23:294–301.

216.	Jahn D, Verkamp E, Soll D. 1992. Glutamyl-transfer RNA: a precursor of heme and chlorophyll biosynthesis. *Trends* *Biochem* *Sci* 17:215–8.

217.	Maes L, Mares-Mejia I, Martin E, Bickel D, Claeys S, Vranken W, Fislage M, Galicia C, Versees W. 2025. Cryo-EM structures of the MnmE-MnmG complex reveal large conformational changes and provide new insights into the mechanism of tRNA modification. *Nucleic* *Acids* *Res* 53.

218.	Bommisetti P, Bandarian V. 2023. Insights into the Mechanism of Installation of 5-Carboxymethylaminomethyl Uridine Hypermodification by tRNA-Modifying Enzymes MnmE and MnmG. *J Am Chem Soc* 145:26947–26961.

219.	Lies M, Visser BJ, Joshi MC, Magnan D, Bates D. 2015. MioC and GidA proteins promote cell division in *E. coli. Front Microbiol* 6:516.

220.	Brown SM, Howell ML, Vasil ML, Anderson AJ, Hassett DJ. 1995. Cloning and characterization of the *katB* gene of *Pseudomonas* *aeruginosa* encoding a hydrogen peroxide-inducible catalase: purification of KatB, cellular localization, and demonstration that it is essential for optimal resistance to hydrogen peroxide. *J* *Bacteriol* 177:6536–44.

221.	Kaushik V, Tiwari M, Tiwari V. 2022. Interaction of RecA mediated SOS response with bacterial persistence, biofilm formation, and host response. *Int J Biol Macromol* 217:931–943.

222.	Gao B, Liang L, Su L, Wen A, Zhou C, Feng Y. 2023. Structural basis for regulation of SOS response in bacteria. *Proc* *Natl* *Acad* *Sci* U S A 120:e2217493120.

223.	Misra HS, Rajpurohit YS. 2024. DNA damage response and cell cycle regulation in bacteria: a twist around the paradigm. *Front* *Microbiol* 15:1389074.

224.	Mozaheb N, Mingeot-Leclercq MP. 2020. Membrane Vesicle Production as a Bacterial Defense Against Stress. *Front* *Microbiol* 11:600221.

225.	Briaud P, Carroll RK. 2020. Extracellular Vesicle Biogenesis and Functions in Gram-Positive Bacteria. *Infect* *Immun* 88.

226.	Wang H, Beier N, Boedeker C, Sztajer H, Henke P, Neumann-Schaal M, Mansky J, Rohde M, Overmann J, Petersen J, Klawonn F, Kucklick M, Engelmann S, Tomasch J, Wagner-Dobler I. 2021. *Dinoroseobacter* *shibae* Outer Membrane Vesicles Are Enriched for the Chromosome Dimer Resolution Site *dif*. *mSystems* 6.

227.	Lin J, Zhang W, Cheng J, Yang X, Zhu K, Wang Y, Wei G, Qian PY, Luo ZQ, Shen X. 2017. A *Pseudomonas* T6SS effector recruits PQS-containing outer membrane vesicles for iron acquisition. *Nat* *Commun* 8:14888.

228.	Santos TMA, Lammers MG, Zhou M, Sparks IL, Rajendran M, Fang D, De Jesus CLY, Carneiro GFR, Cui Q, Weibel DB. 2018. Small Molecule Chelators Reveal That Iron Starvation Inhibits Late Stages of Bacterial Cytokinesis. *ACS* *Chem* *Biol* 13:235–246.

229.	Panoff JM, Corroler D, Thammavongs B, Boutibonnes P. 1997. Differentiation between cold shock proteins and cold acclimation proteins in a mesophilic gram-positive bacterium, *Enterococcus* *faecalis* JH2-2. *J* *Bacteriol* 179:4451–4.

230.	Li H, Giuliodori AM, Wang X, Tian S, Su Z, Gualerzi CO, Sun Z, Fei M, Sun D, Ma H, He C. 2025. Cold Shock Proteins Mediate Transcription of Ribosomal RNA in *Escherichia* *coli* Under Cold-Stress Conditions. *Biomolecules* 15.

231.	Jiang W, Hou Y, Inouye M. 1997. CspA, the major cold-shock protein of *Escherichia* *coli*, is an RNA chaperone. *J* *Biol* *Chem* 272:196–202.

232.	Giuliodori AM, Belardinelli R, Duval M, Garofalo R, Schenckbecher E, Hauryliuk V, Ennifar E, Marzi S. 2023. *Escherichia* *coli* CspA stimulates translation in the cold of its own mRNA by promoting ribosome progression. *Front* *Microbiol* 14:1118329.

233.	Rodrigues JL, Sousa M, Prather KL, Kluskens LD, Rodrigues LR. 2014. Selection of *Escherichia* *coli* heat shock promoters toward their application as stress probes. *J* *Biotechnol* 188:61–71.

234.	Cheng X, Wang W, Molineux IJ. 2004. F exclusion of bacteriophage T7 occurs at the cell membrane. *Virology* 326:340–52.

235.	Wang Y, Dong W, Chu L, Zhao H, He L, Sheng X. 2024. A combination of proteomics, genetics, and physiology provides insights into the acid-tolerance phenotype of *Pseudomonas* *pergaminensis* F77. *Microbiol* *Res* 278:127545.

236.	Nonomura Y, Wang X, Kikuchi T, Matsui D, Toyotake Y, Takagi K, Wakayama M. 2023. Characterization of three gamma-glutamyltranspeptidases from *Pseudomonas* *aeruginosa* PAO1. J Gen Appl Microbiol 69:150–158.

237.	Okada T, Suzuki H, Wada K, Kumagai H, Fukuyama K. 2006. Crystal structures of gamma-glutamyltranspeptidase from *Escherichia* *coli*, a key enzyme in glutathione metabolism, and its reaction intermediate. *Proc* *Natl* *Acad* *Sci* U S A 103:6471–6.

238.	Wongsaroj L, Saninjuk K, Romsang A, Duang-Nkern J, Trinachartvanit W, Vattanaviboon P, Mongkolsuk S. 2018. *Pseudomonas* *aeruginosa* glutathione biosynthesis genes play multiple roles in stress protection, bacterial virulence and biofilm formation. *PLoS* *One* 13:e0205815.

239.	Van Laar TA, Esani S, Birges TJ, Hazen B, Thomas JM, Rawat M. 2018. *Pseudomonas* *aeruginosa* *gshA* Mutant Is Defective in Biofilm Formation, Swarming, and Pyocyanin Production. *mSphere* 3.

240.	Michie KL, Dees JL, Fleming D, Moustafa DA, Goldberg JB, Rumbaugh KP, Whiteley M. 2020. Role of *Pseudomonas* *aeruginosa* Glutathione Biosynthesis in Lung and Soft Tissue Infection. *Infect* *Immun* 88.

241.	Zhou H, Wang M, Smalley NE, Kostylev M, Schaefer AL, Greenberg EP, Dandekar AA, Xu F. 2019. Modulation of *Pseudomonas* *aeruginosa* *Quorum* *Sensing* by Glutathione. *J* *Bacteriol* 201.

242.	Iyer A, Frallicciardi J, le Paige UBA, Narasimhan S, Luo Y, Sieiro PA, Syga L, van den Brekel F, Tran BM, Tjioe R, Schuurman-Wolters G, Stuart MCA, Baldus M, van Ingen H, Poolman B. 2024. The Structure and Function of the Bacterial Osmotically Inducible Protein Y. *J Mol Biol* 436:168668.

243.	Zheng X, Ji Y, Weng X, Huang X. 2015. RpoS-dependent expression of OsmY in *Salmonella* *enterica* serovar typhi: activation under stationary phase and SPI-2-inducing conditions. *Curr* *Microbiol* 70:877–82.

244.	Yan Z, Hussain S, Wang X, Bernstein HD, Bardwell JCA. 2019. Chaperone OsmY facilitates the biogenesis of a major family of autotransporters. *Mol* *Microbiol* 112:1373–1387.

245.	Motouchi S, Kobayashi K, Nakai H, Nakajima M. 2023. Identification of enzymatic functions of osmo-regulated periplasmic glucan biosynthesis proteins from *Escherichia* *coli* reveals a novel glycoside hydrolase family. *Commun* *Biol* 6:961.

246.	Lequette Y, Rollet E, Delangle A, Greenberg EP, Bohin JP. 2007. Linear osmoregulated periplasmic glucans are encoded by the *opgGH* locus of *Pseudomonas* *aeruginosa*. *Microbiology* (Reading) 153:3255–3263.

247.	Kumawat M, Chaudhary D, Nabi B, Kumar M, Sarma DK, Shubham S, Karuna I, Ahlawat N, Ahlawat S. 2021. Purification and characterization of Cyclophilin: a protein associated with protein folding in *Salmonella* *Typhimurium*. *Arch* *Microbiol* 203:5509–5517.

248.	Skagia A, Vezyri E, Grados K, Venieraki A, Karpusas M, Katinakis P, Dimou M. 2017. Structure-Function Analysis of the Periplasmic *Escherichia* *coli* Cyclophilin PpiA in Relation to Biofilm Formation. *J* *Mol* *Microbiol* *Biotechnol* 27:228–236.

249.	Fernandez L, Breidenstein EBM, Taylor PK, Bains M, de la Fuente-Nunez C, Fang Y, Foster LJ, Hancock REW. 2016. Interconnection of post-transcriptional regulation: The RNA-binding protein Hfq is a novel target of the Lon protease in *Pseudomonas* *aeruginosa*. *Sci* *Rep* 6:26811.

250.	Yang N, Ding S, Chen F, Zhang X, Xia Y, Di H, Cao Q, Deng X, Wu M, Wong CC, Tian XX, Yang CG, Zhao J, Lan L. 2015. The Crc protein participates in down-regulation of the Lon gene to promote rhamnolipid production and *rhl* *quorum* *sensing* in *Pseudomonas* *aeruginosa*. *Mol* *Microbiol* 96:526–47.

251.	Breidenstein EB, Janot L, Strehmel J, Fernandez L, Taylor PK, Kukavica-Ibrulj I, Gellatly SL, Levesque RC, Overhage J, Hancock RE. 2012. The Lon protease is essential for full virulence in *Pseudomonas* *aeruginosa*. *PLoS* *One* 7:e49123.

252.	Kallazhi A, Rahman A, Romling U, Jonas K. 2025. Direct and indirect pathways linking the Lon protease to motility behaviors in the pathogen *Pseudomonas* *aeruginosa*. *PLoS* *Pathog* 21:e1013288.

253.	Callewaert L, Aertsen A, Deckers D, Vanoirbeek KG, Vanderkelen L, Van Herreweghe JM, Masschalck B, Nakimbugwe D, Robben J, Michiels CW. 2008. A new family of lysozyme inhibitors contributing to lysozyme tolerance in gram-negative bacteria. *PLoS* *Pathog* 4:e1000019.

254.	Yum S, Kim MJ, Xu Y, Jin XL, Yoo HY, Park JW, Gong JH, Choe KM, Lee BL, Ha NC. 2009. Structural basis for the recognition of lysozyme by MliC, a periplasmic lysozyme inhibitor in Gram-negative bacteria. *Biochem* *Biophys* *Res* *Commun* 378:244–8.

255.	Tseng BS, Reichhardt C, Merrihew GE, Araujo-Hernandez SA, Harrison JJ, MacCoss MJ, Parsek MR. 2018. A Biofilm Matrix-Associated Protease Inhibitor Protects *Pseudomonas* *aeruginosa* from Proteolytic Attack. *mBio* 9.

256.	Fantone KM, Nothaft H, Son Y, Stecenko AA, Szymanski CM, Rada B. 2024. The bacterial serine protease inhibitor ecotin inhibits neutrophil elastase enzymatic activity in cystic fibrosis sputa. *Heliyon* 10:e38895.

257.	Nagy ZA, Szakacs D, Boros E, Heja D, Vigh E, Sandor N, Jozsi M, Oroszlan G, Dobo J, Gal P, Pal G. 2019. Ecotin, a microbial inhibitor of serine proteases, blocks multiple complement dependent and independent microbicidal activities of human serum. *PLoS* *Pathog* 15:e1008232.

258.	Zeb S, Nazir A, Hameed MF, Ikram S, Haider Naqvi SZ, Shoaib M, Butaye P, Wang Z, Li R, Lu X. 2026. Colistin Resistance in Gram-Negative Bacteria: Mechanisms, Transmission, and Novel Intervention Strategies. *Microorganisms* 14.

259.	Jacobsen F, Mohammadi-Tabrisi A, Hirsch T, Mittler D, Mygind PH, Sonksen CP, Raventos D, Kristensen HH, Gatermann S, Lehnhardt M, Daigeler A, Steinau HU, Steinstraesser L. 2007. Antimicrobial activity of the recombinant designer host defence peptide P-novispirin G10 in infected full-thickness wounds of porcine skin. *J Antimicrob Chemother* 59:493–8.

260.	Jochumsen N, Liu Y, Molin S, Folkesson A. 2011. A Mig-14-like protein (PA5003) affects antimicrobial peptide recognition in *Pseudomonas* *aeruginosa*. *Microbiology* (Reading) 157:2647–2657.

261.	Corona F, Reales-Calderon JA, Gil C, Martinez JL. 2018. The development of a new parameter for tracking post-transcriptional regulation allows the detailed map of the *Pseudomonas* *aeruginosa* Crc regulon. *Sci* *Rep* 8:16793.

262.	Horna G, Ruiz J. 2021. Type 3 secretion system of *Pseudomonas* *aeruginosa*. *Microbiol* *Res* 246:126719.

263.	Yin L, Gong X, Li Y, Qu J, Zhang Y, Jin Y, Jin S, Wu W. 2025. The Catabolite Repression Control Protein Crc Regulates the Type III Secretion System Through the Adenylate Cyclase CyaB in *Pseudomonas* *aeruginosa*. *Microorganisms* 13.

264.	Zhou N, Yu J, Liu X, Li C, Tang H, Lyu L, Wu C, Chen Y, Zhang J, Ni J, Wang D, Tao J, Wu W, Zhang Y, Feng Y, Chao Y, Lu J, He P, Yao YF. 2025. Within-host evolution of a transcriptional regulator contributes to the establishment of chronic *Pseudomonas* *aeruginosa* infection. *Cell* *Rep* 44:115214.

265.	Tribelli PM, Lujan AM, Pardo A, Ibarra JG, Fernandez Do Porto D, Smania A, Lopez NI. 2019. Core regulon of the global anaerobic regulator Anr targets central metabolism functions in *Pseudomonas* species. *Sci* *Rep* 9:9065.

266.	Wang H, Yang Z, Swingle B, Kvitko BH. 2021. AlgU, a Conserved Sigma Factor Regulating Abiotic Stress Tolerance and Promoting Virulence in *Pseudomonas* *syringae*. *Mol* *Plant* *Microbe* *Interact* 34:326–336.

267.	Deretic V, Martin DW, Schurr MJ, Mudd MH, Hibler NS, Curcic R, Boucher JC. 1993. Conversion to mucoidy in *Pseudomonas* *aeruginosa*. *Biotechnology* (N Y) 11:1133–6.

268.	Firoved AM, Boucher JC, Deretic V. 2002. Global genomic analysis of AlgU (sigma(E))-dependent promoters (sigmulon) in *Pseudomonas* *aeruginosa* and implications for inflammatory processes in cystic fibrosis. *J* *Bacteriol* 184:1057–64.

269.	Chevalier S, Bouffartigues E, Bazire A, Tahrioui A, Duchesne R, Tortuel D, Maillot O, Clamens T, Orange N, Feuilloley MGJ, Lesouhaitier O, Dufour A, Cornelis P. 2019. Extracytoplasmic function sigma factors in *Pseudomonas* *aeruginosa*. *Biochim* *Biophys* *Acta* *Gene* *Regul* *Mech* 1862:706–721.

270.	Wood LF, Ohman DE. 2009. Use of cell wall stress to characterize sigma 22 (AlgT/U) activation by regulated proteolysis and its regulon in *Pseudomonas* *aeruginosa*. *Mol* *Microbiol* 72:183–201.

271.	Wood LF, Leech AJ, Ohman DE. 2006. Cell wall-inhibitory antibiotics activate the alginate biosynthesis operon in *Pseudomonas* *aeruginosa*: Roles of sigma (AlgT) and the AlgW and Prc proteases. *Mol* *Microbiol* 62:412–26.

272.	Wang B, Lin YC, Vasquez-Rifo A, Jo J, Price-Whelan A, McDonald ST, Brown LM, Sieben C, Dietrich LEP. 2021. *Pseudomonas* *aeruginosa* PA14 produces R-bodies, extendable protein polymers with roles in host colonization and virulence. *Nat* *Commun* 12:4613.

273.	Schulz S, Eckweiler D, Bielecka A, Nicolai T, Franke R, Dotsch A, Hornischer K, Bruchmann S, Duvel J, Haussler S. 2015. Elucidation of sigma factor-associated networks in *Pseudomonas* *aeruginosa* reveals a modular architecture with limited and function-specific crosstalk. *PLoS* *Pathog* 11:e1004744.

274.	Thoma S, Schobert M. 2009. An improved *Escherichia* *coli* donor strain for diparental mating. *FEMS* *Microbiol* Lett 294:127–32.

275.	Karimova G, Pidoux J, Ullmann A, Ladant D. 1998. A bacterial two-hybrid system based on a reconstituted signal transduction pathway. *Proc* *Natl* *Acad* *Sci* U S A 95:5752–6.

276.	Silva-Rocha R, Martinez-Garcia E, Calles B, Chavarria M, Arce-Rodriguez A, de Las Heras A, Paez-Espino AD, Durante-Rodriguez G, Kim J, Nikel PI, Platero R, de Lorenzo V. 2013. The Standard European Vector Architecture (SEVA): a coherent platform for the analysis and deployment of complex prokaryotic phenotypes. *Nucleic* *Acids* *Res* 41:D666–75.

277.	Wirth NT, Kozaeva E, Nikel PI. 2020. Accelerated genome engineering of *Pseudomonas* *putida* by I-SceI-mediated recombination and CRISPR-Cas9 counterselection. *Microb* *Biotechnol* 13:233–249.

278.	Pankratz D, Gomez NO, Nielsen A, Mustafayeva A, Gur M, Arce-Rodriguez F, Nikel PI, Haussler S, Arce-Rodriguez A. 2023. An expanded CRISPR-Cas9-assisted recombineering toolkit for engineering genetically intractable *Pseudomonas* *aeruginosa* isolates. *Nat* *Protoc* 18:3253–3288.

279.	Jeske A, Arce-Rodriguez A, Thoming JG, Tomasch J, Haussler S. 2022. Evolution of biofilm-adapted gene expression profiles in *lasR*-deficient clinical *Pseudomonas* *aeruginosa* isolates. *NPJ* *Biofilms* *Microbiomes* 8:6.
